# Accelerating the clock: Interconnected speedup of energetic and molecular dynamics during aging in cultured human cells

**DOI:** 10.1101/2022.05.10.491392

**Authors:** Gabriel Sturm, Natalia Bobba-Alves, Robert A. Tumasian, Jeremy Michelson, Luigi Ferrucci, Martin Picard, Christopher P. Kempes

## Abstract

To understand how organisms age, we need reliable multimodal molecular data collected at high temporal resolution, in specific cell types, across the lifespan. We also need interpretative theory that connects aging with basic mechanisms and physiological tradeoffs. Here we leverage a simple cellular replicative aging system combined with mathematical theory to address organismal aging. We used cultured primary human fibroblasts from multiple donors to molecularly and energetically profile entire effective lifespans of up to nine months. We generated high-density trajectories of division rates, telomere shortening, DNA methylation, RNAseq, secreted proteins/cytokines and cell-free DNA, in parallel with bioenergetic trajectories of ATP synthesis rates derived from both mitochondrial oxidative phosphorylation and glycolysis, reflecting total cellular mass-specific metabolic rate (MR). By comparing our cell culture data to data from cells in the body we uncover three fundamental speedups, or rescalings, of MR and molecular aging markers. To explain these rescalings we deploy the allometric theory of metabolism which predicts that the rate of biological aging is related to an organism’s size, MR, and the partitioning of energetic resources between growth and maintenance processes. Extending this theory we report three main findings: 1) human cells isolated from the body with faster rates of growth allocate a substantially smaller fraction of their energy budget to maintenance, and correspondingly age 50-300x faster based on multiple molecular markers. 2) Over the course of the cellular lifespan, primary human fibroblasts acquire a >100-fold hypermetabolic phenotype characterized by increased maintenance costs, and associated with increased mtDNA genome density, upregulation of senescence-associated extracellular secretion, and induction of maintenance-related transcriptional programs. 3) Finally, manipulating MR with mitochondria-targeted metabolic, genetic, and pharmacological perturbations predictably altered the molecular rate of aging, providing experimental evidence for the interplay of MR and aging in a human system. These data highlight the key role that the partitioning of energetic resources between growth and maintenance/repair processes plays in cellular aging, and converge with predictions of cross-species metabolic theory indicating that energy metabolism governs how human cells age.

**Significance Statement:** How cells age is of fundamental importance to understanding the diversity of mammalian lifespans and the wide variation in human aging trajectories. By aging primary human fibroblasts over several months in parallel with multi-omics and energetic profiling, we find that as human cells age and progressively divide more slowly, surprisingly, they progressively consume energy *faster*. By manipulating cellular metabolic rates, we confirm that the higher the cellular metabolic rate, the faster cells experience telomere shortening and epigenetic aging – a speedup phenotype consistent with allometric scaling theory. By modeling robust energetic and molecular aging trajectories across donors and experimental conditions, we find that independent of cell division rates, molecular aging trajectories are predicted by the partitioning of the energy budget between growth and maintenance processes. These results integrate molecular and energetic drivers of aging and therefore have important long-term implications to understand biological aging phenomena ranging from cellular senescence to human longevity.

## INTRODUCTION

Recent progress has been made to uncover the fundamental mechanisms of aging using quantitative molecular signatures (Wiley and Campisi 2021; López-Otín et al. 2013). These signatures include telomere length, DNA methylation, and cytokine secretion, each with the ability to track aging signatures in living organisms and cultured cells (Horvath 2013; Sayed et al. 2021; Belsky et al. 2022; A. A. Johnson et al. 2020; Sturm et al. 2019). However, much remains to be uncovered about the connection between aging mechanisms occurring at the organismal level and those at the isolated cellular level. Furthermore, current models of aging have yet to explain why different biological systems age at different rates and what fundamentally constrains the timescales of lifespan (Rando and Wyss-Coray 2021).

To dynamically examine these mechanisms over time we use *in vitro* manipulation of cultured human cells where biological rates can be dramatically adjusted. It is well-known that compared to whole organisms, isolated proliferating cells undergo replicative senescence (Hayflick and Moorhead 1964) and exhibit shortened lifespan, meaning that replicating cells maintain a self-sustaining system over months, whereas cells in the body from which they are derived can self-sustain over several decades. Whether this absolute difference is an artifact of rapid genomic replication or reflects fundamentally different timescales between *in vitro* and *in vivo* remains unclear. By using genetic, pharmacological, and metabolic manipulations, we modify mass-specific metabolic rates and document corresponding changes in aging rates using multiple aging biomarkers.

To understand the fundamental cellular mechanisms underlying these aging processes we then employ metabolic scaling theory. Metabolic scaling provides a systematic framework for understanding the causes and consequences of vastly different rates and timescales across species. Scaling laws relating body mass to metabolic rate, organism growth, and lifespan are among the most universal quantitative patterns in biology. These patterns are found in all major taxonomic groups from prokaryotes to metazoans, although with some exceptions (e.g., birds and domesticated dogs) (G. West 2018; J. H. Brown et al. 2004; DeLong et al. 2010; Kempes, Dutkiewicz, and Follows 2012; Hatton et al. 2019). Notably, differences in the “internal clock” that determine developmental rates in mouse (faster) and human (slower) are driven by bioenergetic differences linked to mitochondrial oxidative phosphorylation (OxPhos) (Matsuda et al. 2020; Diaz-Cuadros et al. 2023). We show how the underlying principles of this theory can be applied to the lifespans of cultured primary human cells and that this theory explains the interconnected rates and timescales when isolating from the body to a cultured system.

A key strength of our approach is that it: i) relies on direct multi-omic measures of molecular dynamics across the entire life course of a cell population, rather than on punctual comparisons of specific cellular states (e.g., replicative state vs senescence), and ii) links canonical features of cell senescence with novel metabolic profiling across the replicative lifespan. Furthermore, we combine these results with a theoretical framework capable of predicting the observed rate rescaling from fundamental energetic perspectives. This work is of broad importance because it provides continuous molecular and metabolic profiles across the replicative and senescence portion of the cellular lifespan, and supports the notion that the rates of aging can be altered in a single species by manipulating metabolic rates.

## RESULTS

### Age-related cytometric and bioenergetic characterization of the cellular lifespan

We performed parallel multimodal measurements of molecular and metabolic rates in primary cultured human dermal fibroblasts for up to 260 days of cellular lifespan from five healthy donors (3 females and 2 males, 0-36 years old, **Extended Data Figure 1A**). As expected, cells show stable changes over the first few months, followed by a precipitous change associated with quiescence and senescence. Specifically, over the early phase of the cellular lifespan, division rate, cell volume, mtDNA copy number, and metabolic rate (MR) are all roughly constant (**Figure 1A-F, Supporting Information i-iii**). Beyond the expected cellular features of cellular aging, monitoring MR longitudinally across the lifespan revealed a striking age-related rise in MR occurring in conjunction with the loss of cell growth (i.e. senescence), indicating a hypermetabolic phenotype of aged cells consistent with recent proteomic results in models of senescence (Kim et al. 2023) (**Figure 1G-H**). Collectively, these measures define a normal cellular lifespan where physiology is roughly homeostatic and comparable to adult physiology. These comparisons will next lead us to identify three molecular rescaling signatures (i.e., speedups) when moving cells from the human body (*in vivo*) to cell culture (*in vitro*).

**Figure 1.**
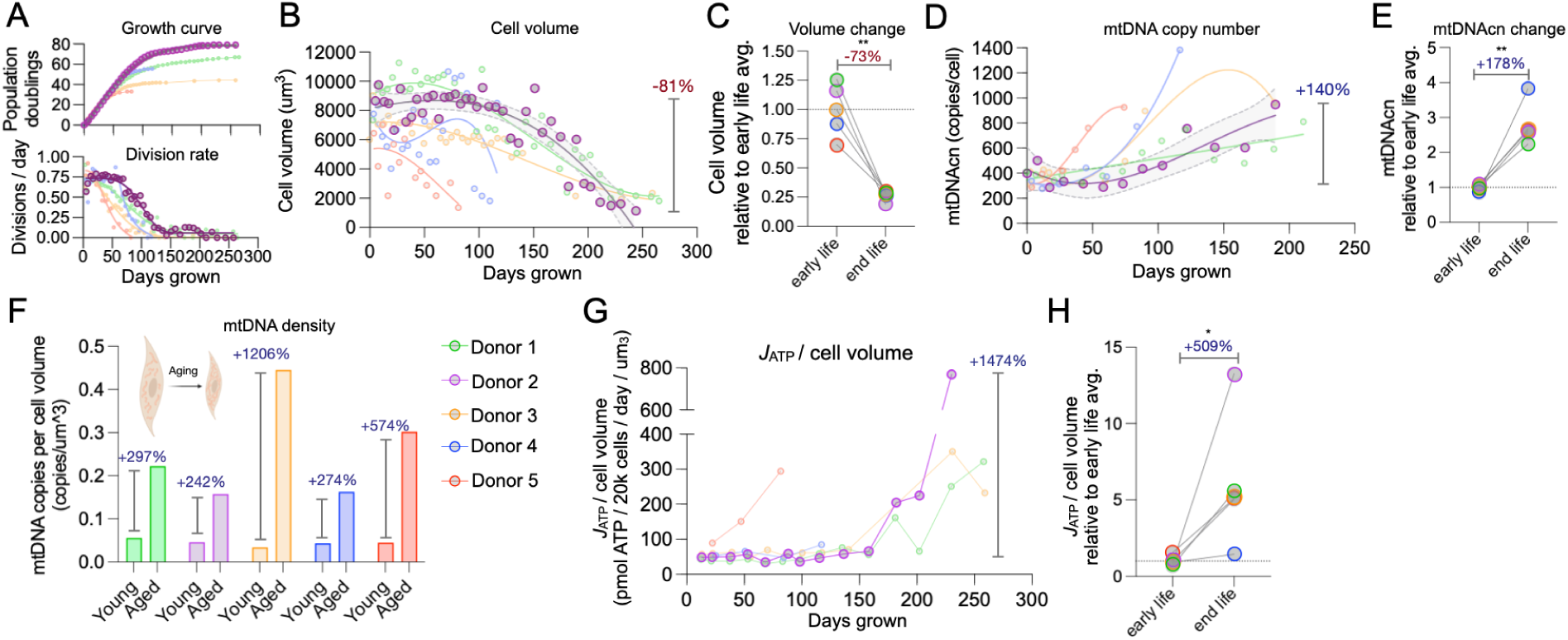
Lifespan trajectories of cell cytometry and bioenergetics reveal late-life hypermetabolism. (**A**) Cumulative population doubling (*top*) and doubling rates (*bottom*) were measured across the replicative lifespan for all fibroblast lines (n=5 donors). The longest lived cell line, Donor 2 (*purple*), is highlighted for clarity. (**B**) Absolute cell volume change (polynomial fit, 95% confidence interval) and (**C**) percent loss from early to late life (n = 7-23 timepoints per cell line). (**D**) Longitudinal trajectories for the absolute number of mitochondrial DNA copies per cell (mtDNAcn), (**E**) average percent change from early to late life, and (**F**) intracellular mtDNA density (normalized to cell volume) for young and aged fibroblasts. (**G**) Longitudinal trajectories of total ATP consumption/production rate per cell volume (*J_ATP-Total_* /cell volume) in each donor (**H**) Average change in *J_ATP-Total_* /cell volume (total MR) from early life to end of life across the 5 donors.

### Rescaling #1: Metabolic rates in vivo to in vitro

In humans, the basic metabolic cost of life measured as the resting metabolic rate is roughly ∼1,500 kcal/day, which is generated from oxygen consumption at a rate of 3.5-3.8 mlO_2_/min/kg of body weight, equivalent to 1.34 watts/kg (Taivassalo et al. 2003; Jeppesen et al. 2009; Davies 1961). During maximal exercise (VO_2max_), humans on average achieve an MR of 43 mL O_2_/min/kg or 13.67 watts/kg (Kaminsky et al. 2021), representing a ∼10-fold dynamic range at the whole-body level. The highest ever recorded VO_2max_ is 96 mL O_2_/min/kg or 32.50 watts/kg (Haugen et al. 2018), corresponding to a dynamic range of 25-fold from rest. These resting and maximal human MRs provide a baseline from which to compare MR in substantially smaller, isolated cells of the same species. We initially restricted these analyses to the first 2-5 weeks in culture to avoid the effects of replicative aging while maintaining the largest sample size possible. The resting MR of primary human fibroblasts was 1.44 watts/kg, equivalent to the amount of energy the whole body uses at rest (**Figure 2A**). If we consider that cells with high MR (e.g., beating heart cardiomyocytes, brain neurons) and low MR (e.g., skin fibroblasts) unequally contribute to whole body MR, based on tissue-specific MRs (Wang et al. 2010; Davies 1961) a better estimate of resting fibroblast MR *in vivo* would be 0.85 watts/kg (**Supplemental File 1**). According to this estimate, the isolation of skin fibroblasts increased cell-specific MR by ∼1.8-fold of *in vivo* rates. However, more sensitive estimates of *in vivo* fibroblast rates in dedicated studies are needed and could shift this factor. For instance, pulse-chase estimates of tissue-specific MR in mice indicate *in vivo* skin cells are closer to 2% of the average metabolic rate, putting *in vitro* MR at 48-fold *in vivo* rates (Bartman et al. 2021). Regardless of the baseline estimate, our human fibroblasts at least maintained but likely increased their resting MR.

**Figure 2.**
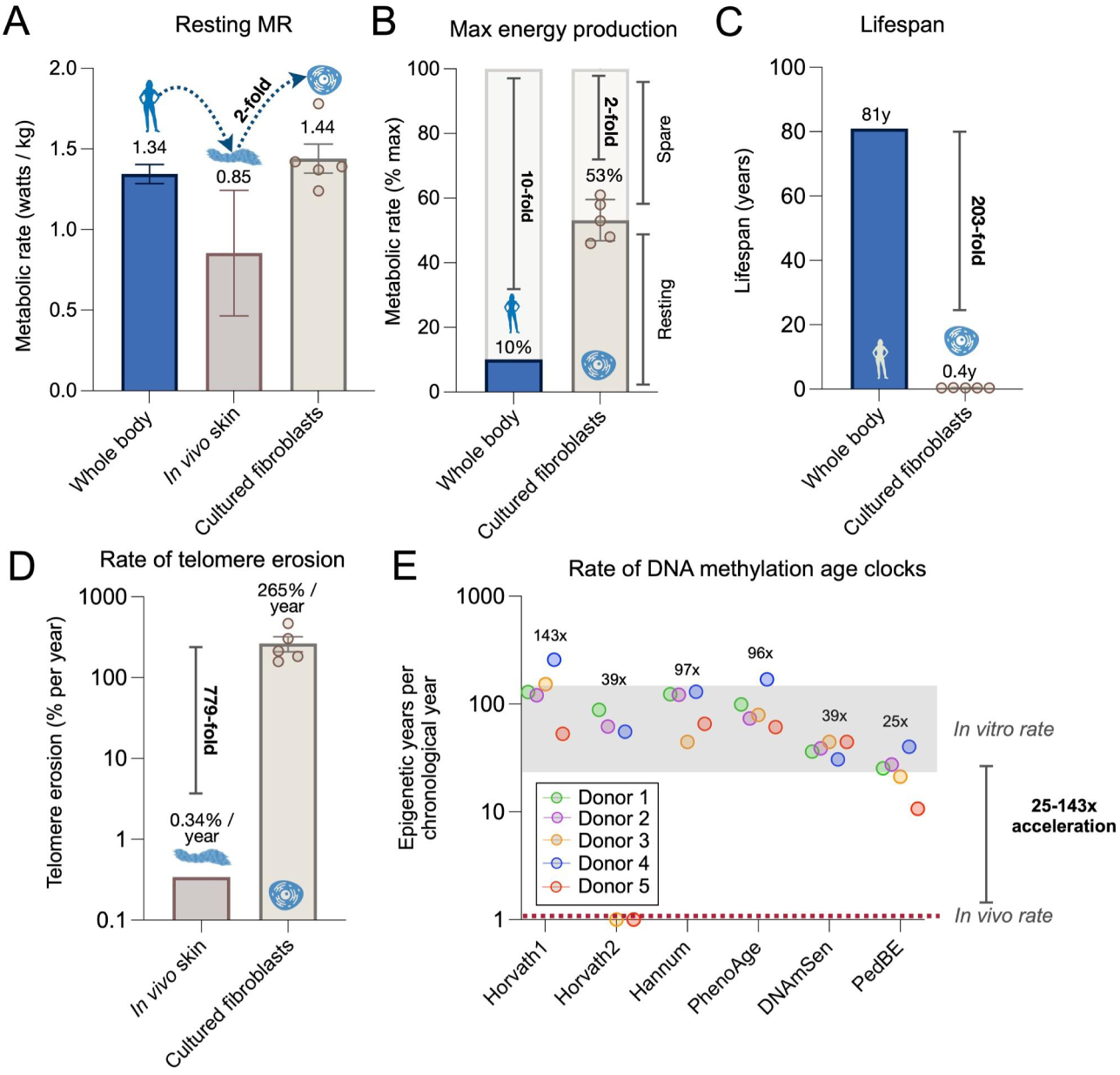
Comparison of *in vivo* and *in vitro* metabolic rates, and rates of biological aging. (**A**) Average MR for the human body, skin tissue (derived values), and primary cultured fibroblasts. Whole body and skin values derived from indirect calorimetry (n=2-3 studies ± S.E.M for independent sources per tissue, see *Methods* and *Supplemental File 1* for details). *In vitro* cellular MR was measured in five primary human fibroblast lines from healthy donors ages 0-36 years using extracellular flux analysis (Seahorse, n=5 donors ± S.E.M, using transformation into metabolic rates from (Mookerjee et al., 2018)). (**B**) Spare oxidative capacity of the whole body and cultured fibroblasts. Values are shown as the percentage of max MR determined by indirect calorimetry at maximal exercise intensity for whole body, and with the inner mitochondrial membrane proton ionophore FCCP for cultured fibroblasts. The reserve capacity of the system is highlighted as fold-change above resting values. (**C**) Average lifespan of humans (defined as average mortality across industrialized countries) and in cultured fibroblasts (defined as the time required to reach the Hayflick limit). Both definitions reflect the time over which the system (body or cell population) is self-sustainable. (**D**) *In vivo* telomere shortening rates in n=200 GTEx samples of 20-70 years old sun-exposed skin samples (Demanelis et al. 2020) compared to telomere shortening rates in cultured fibroblasts longitudinally quantified during the linear growth phase (days 10-70, 4-5 timepoints per Donor). (**E**) Rate of epigenetic aging in cultured fibroblasts (days 10-70, 4-5 timepoints per Donor) quantified with six validated epigenetic clocks trained to predict the rate of human aging *in vivo* (linear slope = 1 biological year per chronological year, red dotted line). Annotations indicate the average rate of biological aging across all donors for each clock, and the average *in vivo*-to-*in vitro* acceleration across all clocks. n=5 donors ± S.E.M for human fibroblasts.

To compare mass-specific *maximal* MR between cells and the whole body, we next measured maximal MR by uncoupling the inner mitochondrial membrane (FCCP) and blocking the mitochondrial ATP synthase (oligomycin). This combination maximizes the rate of oxygen consumption and elevates glycolytic ATP production close to its maximal capacity. Analyzing maximal relative to resting MR showed that isolated fibroblasts normally operate at approximately half of their maximal energy production capacity (**Figure 2B**). This represents a 2-fold spare capacity *in vitro*, compared to the substantially larger 10-25-fold spare capacity of the organism *in vivo*. Together, the relatively high mass-specific resting MR and reduced dynamic energetic range of primary human fibroblasts suggest that isolated fibroblasts exhibit a markedly elevated resting energy expenditure (REE) relative to *in vivo* tissue.

### Rescaling #2: Aging biomarkers in vivo to in vitro

*In vivo*, lifespan is defined as the length of time an organism remains a self-sustaining unit, after which death occurs and the system breaks down (Jones et al. 2014). Similarly, for *in vitro* replicating fibroblasts, lifespan can be conceptualized as the Hayflick limit, quantified as the total number of cell divisions achieved before senescence occurs. In both contexts, the lifespan ends when the system loses its ability to self-sustain. As expected in replicating fibroblasts, near complete growth arrest (i.e., quiescence or senescence) occurred after 148.0±4.9 days, representing approximately 0.4 years of life (**Figure 2C & Extended Data Figure 1B**). Compared to the average human lifespan (81 years) (Bank 2017), this *in vivo* to *in vitro* transition represents a ∼203-fold reduction in the lifespan of the system.

Consistent with well-established molecular features of replicative senescence (Campisi 1997), we observed linear telomere shortening over the first 100 days of growth. Compared to physiological aging of skin where telomeres erode at a rate of 0.00093% per day (Demanelis et al. 2020), relative telomere length of cultured skin fibroblasts decreased by 0.73% per day (**Figure 2D & Extended Data Figure 1H**). Relative to *in vivo* rates, this represents a ∼779-fold acceleration of telomere erosion *in vitro*, a difference that, in part, is attributable to more frequent genome replication *in vitro*.

We also measured genome-wide DNA methylation (DNAm, EPIC platform) longitudinally at multiple timepoints during the lifespan of each cell line to estimate epigenetic age using algorithms or “clocks” trained to accurately predict age in the human body (Sturm et al. 2019). In human tissues, based on the underlying training approach, DNAm clocks predict 1 year of epigenetic age for each calendar year (i.e., a slope of 1) with high accuracy (r>0.95) (Levine et al. 2018; Horvath 2013; Hannum et al. 2013; Horvath et al. 2018; McEwen et al. 2020; Levine et al. 2019). However, in cultured fibroblasts during the early portion of the lifespan, DNAm clocks return slopes ranging between 25-143 biological years per chronological year (**Figure 2E**). Epigenetic aging rates using more robust principal component (PC)-adjusted clocks (Higgins-Chen et al. 2022) revealed a more stable and consistent acceleration of 44-58 biological years per chronological year *in vitro* relative to *in vivo* (**Extended Data Figure 3A-C**). Thus, the rate of epigenetic aging is accelerated by an average factor of ∼50x in isolated cells. Put simply, a week of aging *in vitro* is approximately equivalent to a year of aging *in vivo*. We next confirm this same age-related speed up is apparent across the entire molecular landscape of the cell.

### Rescaling #3: Global transcriptional and methylation in vivo to in vitro

Next we quantified the degree of global rescaling occurring across the functional genome when moving cells from the body to cell culture. *In vivo* transcriptomic and methylome data was aggregated from publicly available cross-sectional studies of skin, the tissue of origin for our fibroblast. The datasets used include GTEx postmortem skin biopsies (RNAseq, n=520, ages 20-70) (Lonsdale et al. 2013) and TwinsUK Biobank skin punch biopsies (DNAm arrays, n=322, ages 40-85) (Moayyeri et al. 2013). *In vitro* RNAseq and DNAm data were both generated from our cultured fibroblasts across the cellular lifespan (n=3 donors, 7-14 timepoints per cell line, 3-265 days grown). Since we are primarily interested in the overall rates of change in gene expression or DNAm levels in noisy data, to impose the least number of assumptions we fit the data using linear regressions. In these regressions, slopes establish the rates of global change for each gene per unit of time, both *in vivo* and *in vitro*. Genes and methylation sites were filtered to include those that significantly change with age (false discovery rate, fdr<0.05), and slopes obtained across the entire lifespan to reflect the rate of change with chronological age. Using the ratio of the median slope of age-related in vivo and in vitro markers then yields rescaling factors reflecting the difference between isolated cells vs the human body (**Extended Data Figure 4A**).

To account for the high variability of cross-sectional *in vivo* data, we quantified rescaling factors using increasingly stringent models (**Extended Data Figure 4B**), which included linear regression, min/max confidence intervals (min-max CI), linear mixed-effects (LMER), and permutation modeling (see methods for further details). RNAseq- or DNAm-based rescaling factors ranged from ∼100-300x. DNA methylation data showed an average ∼63x higher rescaling factors than gene expression data. Thus, regardless of the underlying model and modality (gene expression or DNAm), our analysis reveals a robust speedup of global molecular processes in cells growing in culture, in the same order of magnitude as telomere shortening rates and DNAm clocks.

A critical consideration for these results, which connects with the explanatory theory below, is to assess how much of the speedup was due to cell divisions opposed to hypermetabolism. Thus we estimated RNAseq and DNAm rescaling factors at discrete portions of the lifespan with altered rates of cell division (**Extended Data Figure 4C-D**; **Extended Data Figure 5, *Supporting Information iv*.**) and found that cell division contributes to approximately half of the *in vitro* speedup of biological aging. In other words, even at the end of the cellular lifespan when there is a near absence of cell division, fibroblasts exhibit a speedup of molecular processes of at least ∼100x relative to age-related rates observed in the human body.

### Interpretive theory of interspecies rescaling

So far we have quantified three key rescaling signatures emerging from the culture of isolated cultured cells compared to the whole body: in vitro, cells operate at a mass-specific MR ∼2-48x higher than in vivo, exhibiting a remarkable late-life increase in energetic demand resulting in hypermetabolism, ii) based on canonical human aging markers, cultured cells age ∼50x faster than in the human body, and iii) global age-related molecular recalibrations speed up by 100x, independent of the rate of cell division. To understand these seemingly interconnected results we need a theoretical framework that accounts for aging under vastly different metabolic rates.

An effective framework to consider the systematic relationships governing these rates is that of metabolic scaling theory which stems from comparative biology and macroecology (G. B. West, Brown, and Enquist 1997; J. H. Brown et al. 2004; G. West 2018). This theory explains a variety of interconnected physiological timescales that systematically vary by several orders of magnitude amongst species (G. B. West, Brown, and Enquist 1997; J. H. Brown et al. 2004; G. West 2018). These cross-species scaling relationships can often be connected with fundamental mechanisms and tradeoffs which over evolutionary time are optimized for efficiency (Kempes, Koehl, and West 2019). For example, the nonlinear relationship between metabolic rate and body size has been derived from fundamental transport limitations of vascular systems (G. B. West, Brown, and Enquist 1997; Geoffrey B. West, Brown, and Enquist 1999; G. West 2018) within what is known as the metabolic theory of ecology. The central metabolic theory in turn underpins a variety of physiological rate predictions. For example, smaller mammals are predicted to have greater vascular delivery rates to individual cells which leads to higher cellular metabolic rates, and this relationship is proposed to lead to faster rates of various damage processes and aging (Geoffrey B. West, Woodruff, and Brown 2002; G. West 2018; Hou 2013). This evidence supports the rate of living hypothesis (i.e., animals with faster metabolic rates age faster) and matches longstanding observations indicating that the lifespan of mammals systematically increases with body size following a power law relationship (Speakman 2005; Schmidt-Nielsen and Knut 1984; Economos 1980).

Since the theory is built around interspecific comparisons of mammals it may seem odd to use it for making single cell predictions. However, a single-cell perspective is built into the theory as the terminal units of resource consumption and physiological function, and the theory can be thought of as coupling single cells to a whole-body vascular delivery system. As such, the theory treats whole bodies as the sum of single cells, and makes a variety of cellular rate predictions, including how cells should behave in isolation *in vitro (Geoffrey B. West, Woodruff, and Brown 2002; Savage et al. 2007)*. Here we extend that theory to explicitly predict aging under a variety of metabolic rate shifts for cells in culture. Because this is based around cross-species perspectives where rates naturally shift by several orders of magnitude, it is ideal for interpreting the aging of cells in culture, which exhibit rates roughly two orders of magnitude higher than those found in adult human tissues.

### Maintenance and growth as predictors of lifespan

Across mammals of different adult mass, *M*, metabolic rates, *B*, have been shown to follow *B* ∝ *M*^3/4^ (West, 2018). A key result of the metabolic theory of ecology is that ontogenetic growth can be derived from the partitioning of total metabolism into two main processes: biosynthesis and maintenance. Maintenance stems from the bioenergetic literature and is defined as all cellular processes that do not directly contribute to biosynthesis including repair and turnover processes (Buttgereit and Brand 1995; Kempes, van Bodegom, et al. 2017). The total energy budget of an organism is partitioned between growth and maintenance processes:

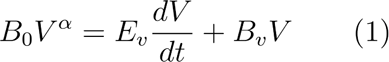

where *E_υ_* is the energy required to synthesize a unit of new biomass (here measured in volume units *V* which is proportional to mass *M*), and *B_υ_* is the metabolic rate required to maintain an existing unit of biomass. This equation can be solved for the growth trajectory of a single organism (Geoffrey B. West, Woodruff, and Brown 2002; Kempes, Dutkiewicz, and Follows 2012; Moses et al. 2008), and it can also be used for finding the growth rate of an organism. This equation can be rewritten in terms of the growth dynamics as

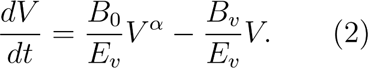

Because α = 3/4, this equation shows that the amount of energy devoted to maintenance, *B_υ_V*, will increase in time and eventually compose the entire energetic budget of a mature organism. This point corresponds to the metabolic homeostasis observed during the initial growth of cells placed in culture and provides a baseline against which to compare late-life departures.

These equations further highlight that the rate at which organisms grow, their eventual adult size, and overall lifespan depend on the fractional investment in maintenance processes (**Extended Data Figure 6A**). More fractional energy investment in growth reduces available resources for to prevent the accumulation of damage, accounting for the observations that smaller animals with more rapid ontogenetic trajectories exhibit predictably higher somatic mutation rate (Cagan et al. 2022) and telomere shortening rates (Whittemore et al. 2019). Development in different organisms can be viewed as the rescaling of a universal curve (G. B. West, Brown, and Enquist 2001; Swovick et al. 2021) the same developmental sequences occur at different rates in different-sized mammals (faster in smaller species) (Matsuda et al. 2020; Diaz-Cuadros et al. 2023), consistent with instructive role of metabolic rates on developmental and aging rates (**Figure 3A**).

**Figure 3.**
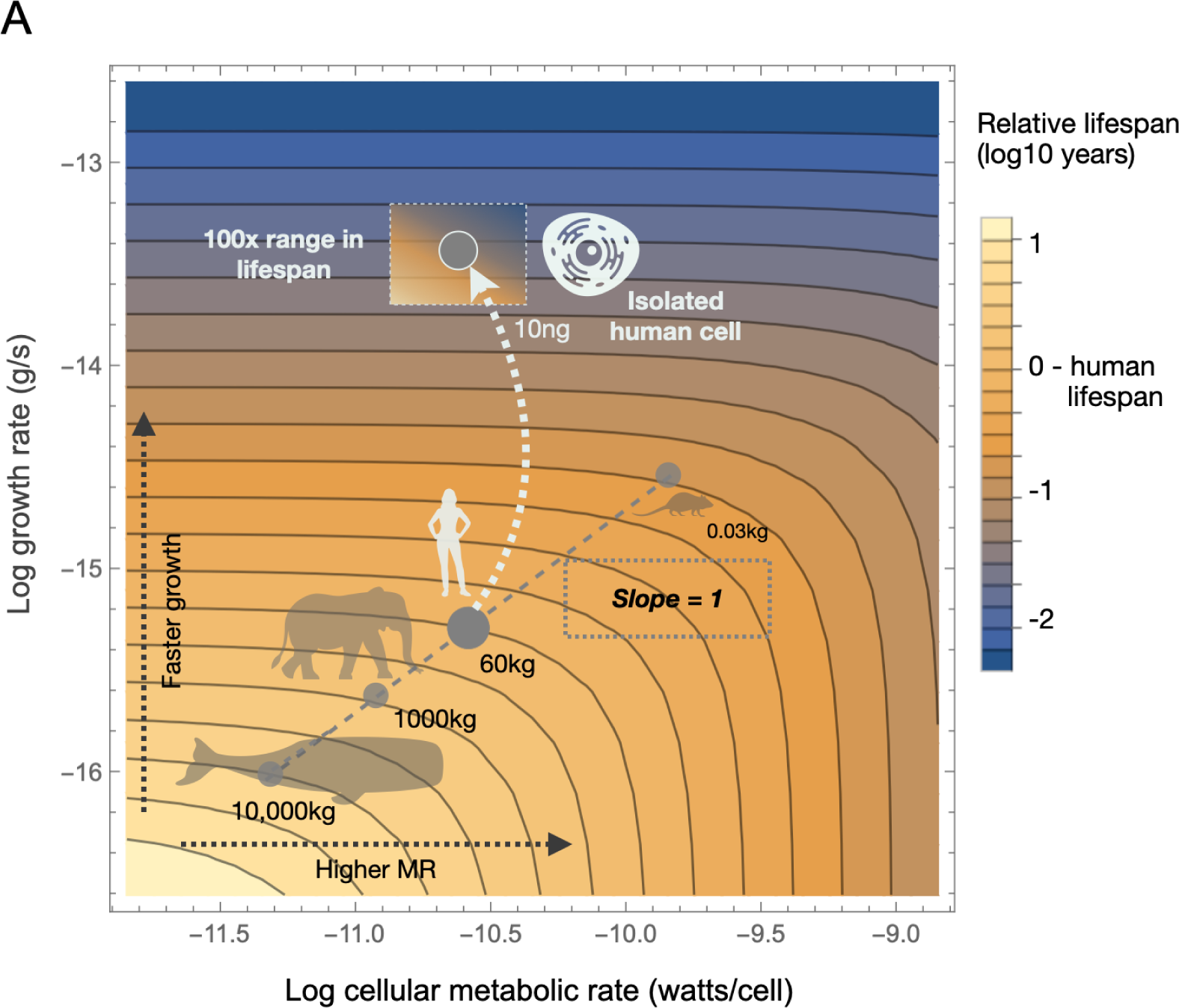
Allometric scaling laws predict accelerated aging in cultured human cells. (**A**) Allometric scaling properties of metabolic rate vs growth rate across mammals of different sizes. Contour plot is log10 cellular metabolic rate (watts/cell) against log10 growth rate (g/s) with the contours representing relative lifespan normalized to a human lifespan in years. Growth rates are estimated from maximum growth rate during development. Dashed gray line represents the range for all mammalian cells *in vivo* of different sizes, spanning mice to whales. White line indicates transition from whole body (*in vivo*) to cultured cells (*in vitro*). Inset box shows ranges of cellular lifespan derived from current dataset.

Since Equation 1 represents the energy budget in terms of costs per unit biomass it is possible to translate this to cellular theory where the metabolic theory typically considers that total biomass and metabolic rates are the sum of individual cells: *V* = *N_c_V_c_* and *B* = *N_c_V_c_*, where denotes the cellular level and *N_c_* is the total number of cells in the body (Geoffrey B. West, Woodruff, and Brown 2002; Savage et al. 2007). Equations 1 and 2 have also been successfully used at the cellular level (Kempes, Dutkiewicz, and Follows 2012) where one can write:

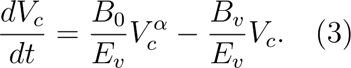

The above equations allows us to analyze the timecourse of cells in culture in two ways:

First, we can use measurements of cell volume and metabolic rate to estimate the metabolic scaling exponent. Interspecifically there are several possibilities for the connection between body mass, cell size, and cellular metabolic rate. These possibilities are bounded by a constant cell size with a cellular metabolic rate that follows *B_c_* ∝ *M*^-1/4^ to a constant cellular metabolic rate with cell sizes that follow *V_c_* ∝ *M*^1/4^ (Savage et al. 2007). From the perspective of aging it is important to consider what happens to cells after they have reached adult size and metabolic homeostasis. For example, it has been shown that humans lose mass later in life (Kuo et al. 2020; Pontzer et al. 2021). If we assume that all loss of mass comes from a decrease in cellular mass while preserving the number of cells, then we would have that 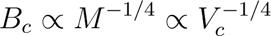, where we consider that *m_c_* ∝ *V_c_*. This result also predicts that the mass or volume specific metabolic rate of cells will follow 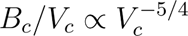 and equivalently 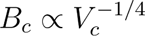. This predicts that once a consistent metabolic rate is reached in a population of cells, then any decrease in cell volume will be accompanied by an increase in metabolic rate following 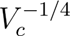, which we observe in our data **(Figure 1 & Extended Data Figure 1)**.

Second, we can use measurements of growth rate, cell size, and energy consumption to estimate the fractions of maintenance and growth **(Extended Data Figure 6A)**. Using the energetic partitioning equations of **Equation I**, we estimate that during cellular isolation from the body, fibroblasts shift the portion of their energetic budget from a lifespan average of ∼6% to ∼77% devoted to biosynthesis processes (**Extended Data Figure 6B**). We then determine the total energy usage devoted to growth and maintenance costs over the cellular lifespan (**Extended Data Figure 6C-D**). Despite total energy production increasing exponentially at the end of the cellular lifespan, the maintenance cost begins to increase gradually across the lifespan, shifting from <10% of total energy consumption early along the cellular lifespan to >90% after growth arrest (i.e., during senescence). Thus, growth costs decrease across the lifespan, with increasing maintenance costs accounting for the majority of the age-related rise in MR.

To gain insight into the origin of increased maintenance costs we measured the senescence associated secretory profile (i.e. SASP), and examine the age-related gene expresion changes using hypothesis-free gene set enrichment analysis, (**see Supporting Information v-vi, and Extended Data Figure 7-10**). As expected, aging cells from all donors exhibited a strong time-dependent elevation of dozens of measures extracellular proteins. Our high-resolution trajectories of media secreted protein levels revealed that this time-dependent upregulation of SASP proteins is exponential, in some cases increasing in concentration by 100-fold between early life to the senescent phase. As one example of a hypersecreted protein by aging cells, the most highly upregulated blood protein in aging humans, growth-derived factor 15 (GDF15) (Tanaka et al. 2020), was released 15-fold higher in aged media. Aging cells also secreted more cell-free mitochondrial DNA, which also must incur energetic costs to maintain. These data validate a global energetic repartioning away from growth and towards maintenance-related biological processes.

### Allometric predictions for interconnected rates

As noted above, a variety of molecular aging markers, including telomere shortening, are rescaled as predicted by growth rates, which is consistent with cross-species perspectives on energetic partitioning between growth and maintenance/repair assuming similar metabolic rates (**Equation 1**).

The above results verify that the basic metabolic theory holds for cell cultures. More importantly, this framework can be easily extended to systematic analyses of lifespan (Geoffrey B. West, Woodruff, and Brown 2002; Hou 2013). Across species lifespans follow *L* ∝ *M*^1/4^ (Speakman, 2005) which can be explained by considering cells from organisms of different body sizes. Since the number of cells is proportional to mass, *N_c_* ∝ *M*, the cellular metabolic rate is given by *B_c_* = *B/N_c_* ∝ *M*^-1/4^ (G. B. West, Brown, and Enquist 2001; Geoffrey B. West, Woodruff, and Brown 2002). Taken together this implies that across mammals of different size every unit of mass receives the same total energy over a lifetime since *B_c_* · *L* ∝ *M*^0^ is a constant. This also shows that cellular lifespans, *L_c_*, are inversely proportional to cell metabolic rates since 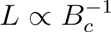 and *L_c_* ∝ *L* and thus 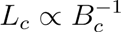. More detailed work has shown that the growth budgets in Equation 3 can be expanded to account for damage and repair costs over the life course of an organism to explain aging (Hou, 2013). We can extend these ideas to the cellular level to consider the lifespan consequences of cellular energetic shifts where lifespan should follow:

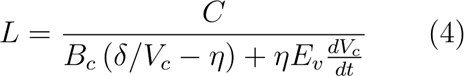

where *C* is the critical fraction of damaged organism mass, *δ* indicates how much damage is associated with a unit of metabolic expenditure, and *η* gives the fraction of maintenance metabolism dedicated to repair.

Equation 3 shows that lifespan will decrease if either *B_c_* or *dV_c_/dt* increase. More specifically, **Figure 3A** shows the map of lifespan for changes in either cellular metabolic rate or growth rate. The theory reveals a few interesting features. First, curves of constant lifespan are roughly defined by by *B_c_* proportional to *dV_c_/dt*. Second, we have shown how the cellular metabolic rates and cellular growth rates change across mammals of different body size with corresponding shifts in lifespan. Here the cellular rates and whole-organism lifespans are shifted by approximately two orders of magnitude. Third, the theory predicts that lifespan will decrease roughly proportionally for increases in *B_c_* or *dV_c_/dt* alone. This allows for a variety of experimental manipulations in culture to test these predictions. For example, *B_c_* in many experiments is roughly constant between adults and cell cultures implying that increases in growth rates should proportionally decrease lifespan following:

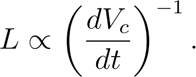

This is exactly what we observe in the transition from *in vivo* to *in vitro* resulting in a ∼203-fold reduction in effective lifespan (Figure 2D). To measure the rates of growth we deployed an epigenetic-based algorithm to count cell divisions (i.e., mitotic clock, (Youn and Wang 2018)) to fat and skin tissue samples from female donors ages 40-85 years (Grundberg et al. 2013), we calculated that on average in an adult human, fibroblasts divide on average every 537 days, while in our cell cultures, the time to divide in the early phase of the cultures is 1.44 days (**Extended Data Figure 11A-C**). Given the limited shift in cellular metabolic rate demonstrated above, this energetic repartitioning towards growth predicts a rescaling of lifespan by a factor of 322, which is comparable in scale to the observed factor of 203. This result illustrates that shifts in metabolic partition from maintenance to growth can dramatically alter cellular lifespans.

More broadly, we can alter the metabolic conditions of cell cultures to shift both cellular metabolic rates and growth rates and systematically test the predictions of Equation 3. By directly modulating metabolic rate while keeping growth rates in a similar range, we then determined if we could alter the degree of *in vitro* rescaling and rate of aging (**Extended Data Figure 12A-B)**. We used a set of established genetic, pharmacological, and environmental treatments that put the cells in either a hypermetabolic (increased MR, >150%) or hypometabolic (decreased MR, ≤27%) state (**Figure 4A-B, see methods**) (Sturm et al. 2022). All treatments reduced growth rates by 10-88% which represents a shift by less than a factor of two in comparison to the 322 fold increase from *in vivo* (**Extended Data Figure 12A**) but growth rates were not significantly correlated with MR (p=0.40, r^2^=0.04, **Extended Data Figure 12B**), suggesting that division rates and energy expenditure are not inextricably coupled, and the overall space of lifespans in **Figure 3A** can be readily explored.

**Figure 4.**
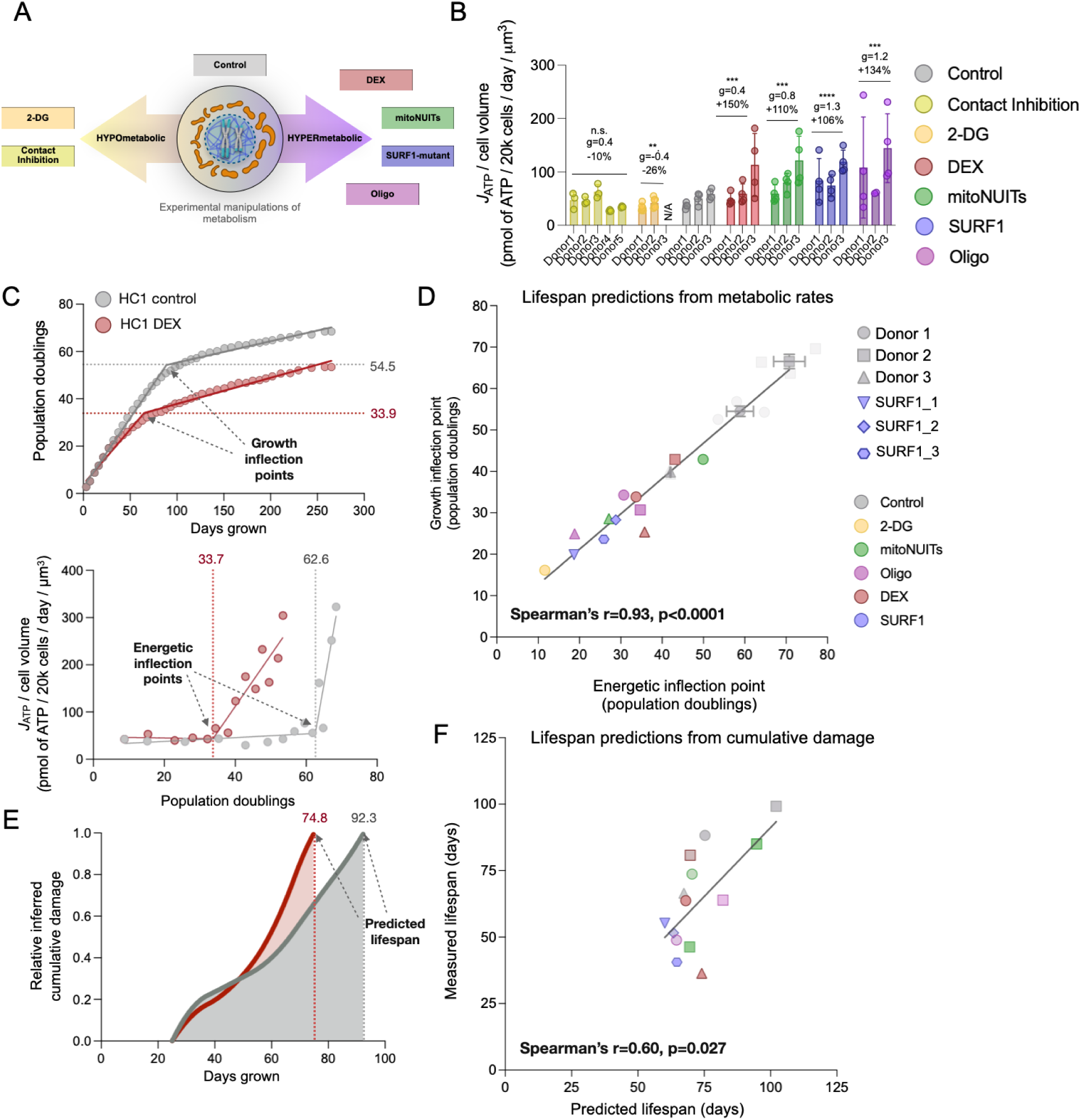
Replicative lifespan predicted by energetic, growth, and damage estimates. (**A**) MR-modulating treatments and (**B**) effects on volume-specific metabolic rates between 25-95 days grown (linear phase of growth). Each bar represents a donor, and each data point is a different timepoint. N/A indicates missing data for 2DG-treated Donor3. *Contact Inhibition* = induction of growth arrest and quiescence through contact inhibiting; *2-DG*, chronic 1mM 2-deoxy-d-glucose; *DEX*, chronic 100nM dexamethasone; *mitoNUITs,* chronic combination of mitochondrial nutrient uptake inhibitors; *SURF1-mutant,* fibroblasts from pediatric patients *SURF1* mutations; Oligo, chronic 1nM oligomycin (see *Methods* for details). (**C**) Example analysis pipeline to derive energetic and growth inflection points (healthy control 1 untreated and 100nM dexamethasone shown). Inflection points were derived using the X0 value of a two-segment linear regression. (**D**) Linear correlation between the number of doublings underwent at the energetic and growth inflection points. Error bars indicate S.E.M. across 2-3 replicate lifespan experiments and were only performed on untreated cell lines. Faded gray points are 2-3 replicate experiments of untreated cells (only mean value included in correlation). (**E**) Cumulative damage over cellular lifespan measured as a function of growth rate and maintenance costs (see Box 1.II). Estimated lifespan is measured as the inverse of the integrated rate over a fixed time (25-95 days grown). (**F**) Linear correlation between the predicted lifespan vs the measured lifespan of a cell population. Measured lifespan is defined as the number of days until the inflection point of growth is reached quantified by a two-segment linear regression. R value indicates spearman correlation. Two-tailed paired ratio t-test, * p < 0.05, ** p < 0.01. * p < 0.05, **** p < 0.0001.

In relation to the coupling of MR and the transition to senescence, despite the large shifts in division rate and Hayflick limit, we observed a remarkable temporal alignment between the inflection points at which growth arrest occurs and volume-specific MR rises at the end of life (Spearman’s r=0.93, p<0.0001), regardless of the intervention used (**Figure 4C-D**). This finding highlights the robust temporal interconnection of hypermetabolism (i.e., excess metabolic rate relative to an organism’s optimum) and cellular senescence, which is consistent with the connection between metabolic and growth rates (Kempes, Dutkiewicz, and Follows 2012). In fact, using the lifespan predictions of Equation 3, we find a significant correlation between the predicted lifespan estimated from cumulative damage over time (**Figure 4E**) to the measured inflection point of growth (Spearman’s r=0.60, p=0.027), i.e. cellular lifespan in a population of cultured cells (**Figure 4F**).

These energetic determinants of lifespan can further be correlated with established biomarkers of aging. Across all experimental conditions that either increase or decrease MR, where each datapoint represents the average of multiple donors, we found that volume-specific MR was linearly correlated with the rate of telomere erosion (p<0.001, r^2^=0.56) and with the rate of epigenetic aging (PCHorvath2, p<0.05, r^2^=0.24) (**Figure 5A-D**). Additionally, MR was linearly correlated with gene expression-derived rescaling factor (p<0.001, r^2^=0.95) but not DNAm (p=0.61, r^2^=0.04) (**Figure 5E-F**). In other words, treatments that increased MR also increased the rate at which age-related gene expression changed. Note that the hypometabolic treatments were able to slow down the rate of methylation changes but hypermetabolic treatments failed to further increase the rate of change beyond untreated cells.

**Figure 5.**
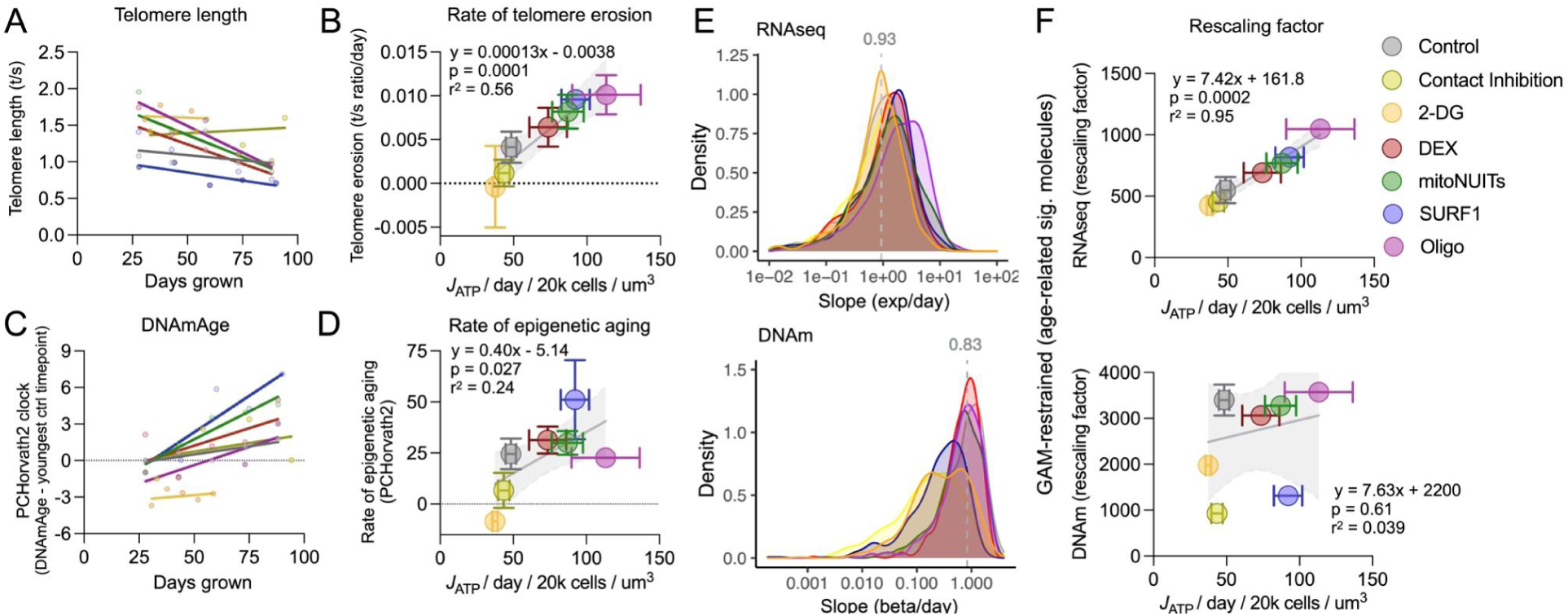
Experimental manipulation of metabolic rate rescales *in vitro* biological aging rates. (**A**) Relative telomere length trajectory (25-95 days grown, for Donor 2 and SURF1_1) used to derive the rate of telomere shortening across MR-altering treatments. (**B**) Correlation between MR and the rate of telomere erosion across all donors (error bars are S.E.M. across donors; linear regression with 95% confidence intervals). (**C**) Baseline-adjusted trajectory of DNAmAge (PCHorvath2, also known as Skin&Blood clock) for the same samples as in (C), and (**D**) correlation between MR and epigenetic aging. Frequency distribution of the slopes for the rate of change for all expressed genes (*top*) and methylated CpG sites (*bottom*) for a given MR-altering treatment. Gray line: median slope of untreated cells. (**F**) Correlation of MR with rescaling factors (*in vivo*-to-*in vitro* acceleration of molecular processes) across all age-related genes (*top*) and CpG sites (*bottom*, linear regressions with 95% confidence intervals), highlighting the strong association between volume-specific MR and age-related gene expression changes.

We further measured the proportion of speed-up attributable to cell division by partially recapitulating the multicellular environment of tissues through contact inhibition, which increases cell-cell contacts and reduces cell division rates (**Extended Data Figure 12C**). Contact inhibition reduced both RNAseq and DNAm rescaling factors by -58% and -85%, respectively. This suggests that over half of aging speedup from cell culturing was caused either by physical isolation or continuous replication, whereas the remaining 42% and 15% for RNAseq and DNAm, respectively, were dependent upon other unidentified physiological factors, such as MR.

Finally, we directly assessed the conservation of the size scaling inverse power law originally found cross-species, to metabolically-altered human fibroblasts (**Extended Data Figure 12D**). We find that despite large shifts in cell volume, up to 73% loss between treatments, the metabolic size scaling coefficient of -0.25 originally found cross-species was maintained across time and treatment groups *in vitro* (**Extended Data Figure 12E**). We, therefore, extend allometric size scaling predictions to cell culture, where smaller cells exhibit predictable increases in MR and commensurate alterations in lifespan trajectories.

## DISCUSSION

We have generated a longitudinal, high-temporal-resolution dataset of bioenergetic, cytological, and genome-wide molecular dynamics in primary fibroblasts from multiple human donors to quantify the rate of biological aging in parallel with MR. This experimental system established three main predictions based off the allometric theory of metabolism: 1) when isolated from the body, human cells exhibit an increased rate of cellular division, allocate a substantially smaller fraction of their energy budget to maintenance, and correspondingly age faster based on multiple molecular markers; 2) over the course of the cellular lifespan, primary human fibroblasts acquire a striking hypermetabolic phenotype in late stages (>100-fold increase in volume-specific MR) characterized by increased maintenance costs, and associated with increased mtDNA genome density, dramatic increases in extracellular secretion, and induction of maintenance-related transcriptional programs (e.g., autophagy); and 3) manipulating MR with mitochondria-targeted metabolic, genetic, and pharmacological perturbations predictably altered the molecular rate of aging, providing experimental evidence for the interplay of MR and aging in a human system. These findings have important implications for several unresolved questions in geroscience and medicine, which we discuss below.

### Energetic perspective of cellular aging

This work complements the replicative theory of cellular aging and emphasizes an alternative perspective based on energetics. This energetic perspective predicts the life course of cultured cells on par with measures of telomere length. Our experimental data indicate that MR-altering perturbations regulate the rate of telomere shortening and the rate of biological aging based on molecular (gene expression and DNA methylation) dynamics. Therefore, this suggests that MR and its partitioning may play a causal role in aging and lifespan regulation. Moreover, the late phase of the lifespan marked by an exponential hypermetabolic trajectory, as costly synthesis and secretion of dozens of proteins increase by orders of magnitude, could constrain growth-related activities and thus contribute to the onset of quiescence and senescence programs. This supports the notion that senescence features can be reversed and cell division restored in replicatively senescent fibroblast treated with cycloheximide to partially inhibit protein synthesis (Takauji et al. 2016). This energetic contribution to molecular aging dynamics could be in addition to, or in parallel with other factors, such as telomere length generally believed to dictate replicative potential. Our experiments rule out the contribution of varying oxygen levels, as previously reported in mitochondrial OxPhos-deficient fibroblasts (Sturm et al. 2023). Therefore, one would expect that preventing the age-related increase in MR-associated mitochondrial components could mitigate aging phenotypes, a prediction in line with the fact that mitochondria are required for the acquisition of senescent phenotypes (Correia-Melo, Marques, and Anderson 2016).

Although this system by no means captures the full physiological complexity of aging, we nonetheless show wide-scale shared age-related changes in molecular processes between *in vivo* and *in vitro* aging. These include epigenetic clocks trained in a variety of tissues, telomere shortening, global gene expression signatures, and DNA methylation at single genomic loci. The high degree of similarity among age-related trajectories in gene expression and DNAm *in vitro* and *in vivo* was the basis for our rescaling factors. All of these processes show that upon isolation and culturing, human cells exhibit a qualitatively similar but unanimous speedup of specific age-related molecular events at a rate 50-300x faster than when they existed in the human body. This is consistent with expectations from theory since lifespans are expected to be inversely proportional to either mass-specific metabolic rate (cellular metabolic rate) or cellular division rate under a fixed cellular metabolic rate (Hou 2013). Because the acceleration of molecular dynamics in cultured fibroblasts occurs even in aged or experimentally stressed cells undergoing practically no divisions over several weeks, this speedup is largely *independent of division rates,* and therefore cannot be fully explained by more rapid proliferation. Our work here illustrates that shifts in either cellular metabolic rates or how energy is partitioned between growth and maintenance will both affect lifespan. Specifically, either increasing growth rates under a fixed energy budget, or increasing cellular metabolic rates directly, both accelerate molecular aging rates.

These speedups are in line with evidence from physiological aging where similar connections to metabolic rate have been described. For example, in humans, elevated resting MR (basal metabolic rate, BMR) is a risk factor for mortality (Ruggiero et al. 2008; Jumpertz et al. 2011; Schrack et al. 2014); correspondingly, lower BMR is associated with healthier and longer lifespans (Schrack et al. 2014). Mitochondrial diseases that trigger energetically costly integrated stress responses both *in vivo* and *in vitro* among fibroblasts similarly cause a speedup of molecular and physiological processes (e.g., elevated heart rate), have elevated MR, and dramatically reduced lifespan (Sturm et al. 2023). This relationship has been further documented in cells exposed to chronic glucocorticoid signaling (Bobba-Alves et al. 2022) and obesity (Tencerova et al. 2019). Similarly, interventions such as exercise and calorie restriction, have been shown to lower basal MR through a process known as metabolic compensation (Careau et al. 2021), which may account for their health and lifespan benefits in humans (Pontzer 2018). Likewise, *C. elegans* worms follow a temporal scaling of lifespan across environmental and genetic interventions including diet, temperature, and stress (Stroustrup et al. 2016). Interestingly, this temporal scaling arises independently of any specific molecular target, thus suggesting a whole-organism control system such as metabolic rate enabling predictable stretching or shrinking of lifespan trajectories across perturbations (Stroustrup et al. 2016).

Overall, the speedup of isolated human fibroblasts mimics the cross-species relationships of size, cellular replication rates, MR, and lifespan. Our findings are also in line with the relationship between the size and metabolism of fibroblasts from different-sized dogs (Jimenez et al. 2018; Brookes and Jimenez 2021). Likewise, cross-species developmental rates, as measured by the molecular dynamics of segmentation clocks, are conserved between the stem cells isolated from humans and mice (Matsuda et al. 2020; Diaz-Cuadros et al. 2023). In cortical neurons these species-specific rates can be regulated by modulating the metabolic activity of mitochondria, again pointing to energy metabolism playing an instructive role in the rate of development and aging (Iwata et al. 2023). Moreover, amongst invertebrates, smaller organisms with higher metabolic rates have been documented to have slowed time perception (perceiving more units of time each second) (Healy et al. 2013) further linking, although indirectly, metabolic rate to the biology of time perception- both *in vivo* and *in vitro*. Our data suggests that the fundamental ability of cultured fibroblasts to generate time- or age-related molecular recalibrations (i.e., to perceive time) may be based on their metabolic rates. For a detailed discussion of potential underlying mechanisms, we refer the reader to the **Supporting Information vii**.

### Lifetime MR trajectories

Our longitudinal age-related trajectories demonstrating profound bioenergetic recalibrations across the cellular lifespan agree with recent work in a human lung fibroblast cell line (WI-38), indicating that senescent cells accumulate higher levels of OxPhos, TCA cycle, and carbon metabolism proteins and transcripts across cellular passages, in parallel with the upregulation of glycolysis in senescence (Chan et al. 2022). Senescent cells have more, not less of the energy-transformation machinery. The age-related increase in MR that we quantify across the lifespan also is in line with a recent report demonstrating, in mice, an age-related elevation in both resting MR and MR associated with a given workload (i.e., decreased metabolic efficiency) (Petr et al. 2021). In other words, in this system as in our fibroblast model, metabolic efficiency declines and the total energetic cost of living increases with age. The increased cost of living is also in keeping with some aspects of human physiology, including the increased mRNA splicing activity that increases in skeletal muscle with aging (Ubaida-Mohien et al. 2019; Ferrucci et al. 2022; Tumasian et al. 2021).

However, the age-related hypermetabolism is in contrast with resting and free-living whole-body MR in aging individuals where lifetime measures of body mass and MR appear to decrease in later life (after roughly 60 years on average) (Ruggiero et al. 2008; Schrack et al. 2014; Pontzer et al. 2021; Kitazoe et al. 2019; Kuo et al. 2020). We speculate that the contrast between our cellular system and the *in vivo* situation may reflect shared bioenergetic constraints at both the cellular and organismal levels, which are met via different strategies. In cultured cells, the adaptive strategy appears to consist of reducing (and eventually halting) the rate of cell division, as well as reducing cell volume, which should similarly decrease total metabolic demand, thereby freeing MR and a portion of the ATP budget for maintenance processes (e.g., SASP secretion). Accordingly, cells with higher MR have markedly lower Hayflick limits and slow division rates. In the aging human body, similar bioenergetic constraints and cellular recalibrations likely occur and may cause, in addition to the atrophy of some energy-consuming tissues (e.g., brain, muscles) in old age, integrated multi-system responses involving endocrine, metabolic, neural, and behavioral recalibrations that contribute to improving efficiency and decreasing MR among the organ network. This would include, for example, aversion to high-intensity activities and reduced physical activity. It should also be noted that while the mass-specific metabolic rate is maintained across adulthood, some older adults lose a significant amount of weight (Pontzer et al. 2021; Kuo et al. 2020; Yeakel, Kempes, and Redner 2018). Moreover, differences between *in vivo* and *in vitro* MR trajectories across the lifespan could also be explained by survivor bias. Old frail individuals may in fact show decreased body mass and a rise in MR (as seen in cultured cells) but die soon after or are not in sufficiently good health condition to be included in the aforementioned population studies of healthy older adults. This selection bias leaves only the “normometabolic” or “hypometabolic” individuals who successfully manage to physiologically decrease MR in order to survive into older ages. The accelerated timescales of cultured cellular lifespans may therefore reflect the late-life frailty period in human years which remains out of reach of epidemiological studies.

Following the predictions made from cross-species scaling laws established at the organismal level, this work extends allometric theory to the cellular level and establishes the interconnected speedup of energetic and molecular processes in cultured human cells. The current study provides an alternative energetic perspective of cellular aging inspired by allometric and metabolic theory, integrated with canonical hallmarks and biomarkers of aging, which opens up many new possibilities to understand the role of energetics in cellular aging.

### Limitations

Looking forward, there are a few important factors that should be considered to provide a complete picture of metabolic rescaling that occurs upon cell isolation. First, to fully integrate our results with the cross-species perspectives it is important to recognize that humans are metabolic outliers in the allometric scaling of body size and lifespan, exhibiting roughly 4x longer lifespans than most species of similar size (Pontzer et al. 2021). Correcting for this deviation and understanding its origin could be useful for future intercomparisons with mice or other model organisms. Second, comparing MR and aging rates between cultured fibroblasts and whole-body dermal skin measurements is imperfect. This comparison is limited by the heterogenous mixture of cell types typical of skin tissue (e.g., fibroblasts, macrophages, adipocytes, mast cells, Schwann cells, and stem cells), although fibroblasts are the principal cell type of the dermal layer (T. M. Brown and Krishnamurthy 2021). We further note that our *in vitro* measurements are taken from cell populations including millions of cells, such that our data reflect population-level measurement and not single cells. Our results therefore reflect comparisons among groups of cells with some of the properties of a complete tissue. This may explain why many of the aging signatures in our fibroblasts mirror whole-body cross-species expectations. Moreover, since it is not possible to precisely isolate skin from other organs, the *in vivo* MR estimates of skin from indirect calorimetry are imprecise. There are also limitations to the extracellular flux measurements on which our cellular MR rates are based (Schmidt, Fisher-Wellman, and Neufer 2021), which together, likely lead to imprecision in the *in vivo* to *in vitro* comparison of MR. Third, we note that there is no known way to directly manipulate MR without off-target effects on other biological processes. MR reflects the integrated state of the system rather than a single, modifiable process. Thus, the MR-altering conditions included in **Figure 5** have unavoidable confounding effects on multiple cellular features that should be considered in the interpretation. Nevertheless, the convergence of results across a spectrum of genetic, metabolic, and pharmacological perturbations with different targets increase our confidence in the generalizability and robustness of our interpretation. Finally, while it is reasonable to assume that energy budgets are fundamentally limiting to organisms, and thus metabolic rate should proportionally adjust all molecular clocks, the explicit mechanisms between metabolic rate and cellular aging will need to be revealed in future work.

## Methods

All *in vitro* data presented in this paper is part of the ‘Cellular Lifespan Study’ which is described in detail in (Sturm et al. 2022), including a more thorough description of all methods. A portion of these data examining the hypermetabolic and accelerated aging effects of primary OxPhos defects (SURF1 mutations and Oligo treatment) are reported in (Sturm et al. 2023), and the effects of chronic dexamethasone treatments are reported in (Bobba-Alves et al. 2022). Please see the ***Supporting Information viii*** for more elaboration of the methodology.

## Data Availability

All data can be accessed, visualized, and downloaded without restrictions at https://columbia-picard.shinyapps.io/shinyapp-Lifespan_Study/. All data visualized is downloadable as a .csv file which can further be found directly at FigShare (18441998). The unprocessed RNAseq (GSE179848) and EPIC DNA methylation array data (GSE179847) can be accessed and downloaded in full through Gene Expression Omnibus (GEO). Brightfield microscopy images can be downloaded at FigShare (18444731). Raw Seahorse assay files along with corresponding data analysis scripts can be found on FigShare (20277606), and GitHub (Cellular_Lifespan_Study/tree/main/Seahorse), respectively. *In vivo* omics data can be found on their respective databases: GTEx RNAseq (dbGAP phs000424.v8.p2) and TwinsUK DNA methylation (GEO GSE90124). The R code for the computational pipeline used to estimate rescaling factors is further available on GitHub (Cellular_Lifespan_Study/tree/main/Rescaling).

## Code Availability

Code is available at https://github.com/gav-sturm/Cellular_Lifespan_Study

## Acknowledgments

This work was supported by NIH grant R01AG066828, the Wharton Fund, and the Baszucki Brain Research Fund to M.P. C.P.K. thanks the Charities Aid Foundation of Canada (CAF) for supporting this work. Special thanks to Wallace Marshall for his conceptual insights and mentorship while drafting this manuscript. The authors are grateful to Anna Monzel for insightful input on drafts of this manuscript, and to Michio Hirano for providing the SURF1 and control cell lines.

## Author contributions

G.S. and M.P. designed the *in vitro* studies. G.S. performed cell culture and sample collection. G.S. and J.M. performed replication experiments. N.B.A. performed inflection point analysis. G.S. and R.A.T. performed age-related statistical modeling. G.S. and C.K. performed all other data analyses. C.K. developed the theory for novel interpretation of the data. G.S., C.K., and M.P. drafted the manuscript. L.F. edited the manuscript. All authors reviewed and approved the final version of this manuscript.

## Competing interests

The authors declare no competing interests.

## Supplementary files

**Supplemental File 1.** Cell-to-whole body oxygen consumption calculator

**Supplemental File 2.** RNAseq generalized additive modeling significant age-related genes

**Supplemental File 3.** DNA methylation generalized additive modeling significant age-related CpGs

**Supplemental File 4.** ShinyGO gene set enriched KEGG pathways for UPregulated RNAseq genes

**Supplemental File 5.** ShinyGO gene set enriched KEGG pathways for DOWNregulated RNAseq genes

**Supplemental File 6.** ShinyGO gene set enriched Gene Ontology processes for UPregulated RNAseq genes

**Supplemental File 7.** ShinyGO gene set enriched Gene Ontology processes for DOWNregulated RNAseq genes

**Supplemental File 8.** Gene expression heatmaps of selected pathways across cellular lifespan

## Supporting Information

### i. Age-related volume reduction and increased mtDNA density

We first describe the cytometric dynamics across the replicative lifespan (**Figure 1A**). As cells entered quiescence and senescence, the population doubling rate decreased progressively from 0.6-0.8 (first passage) to fewer than 0.01 (last passage) divisions per day. Early in life during the linear growth phase, long-lived cells maintained relatively constant cell volume. In the latter portion of the lifespan, although cells appeared larger (covering a larger surface area) when imaged in their adherent state (**Extended Data Figure 1B & 13**), optimal measurements of floating cell diameter showed that the volume of aging fibroblasts declined by 57-83% towards the end of life (average=-72.5%, p<0.0015, **Figure 1B-C**). All cell lines reached a similar lower limit of cell size, suggesting an inherent minimum biological size limit of ∼2000µm^3^ in this cell type. We further estimated the molecular density of cells across the lifespan by combining total protein, RNA, and DNA per cell, which did not reveal any stable age-related trend in cell density. (**Extended Data Figure 1C**). A parallel measure of cellular density was measured using mitochondrial DNA. Accordingly, we observed an increase in the number of mtDNA copies per cell (mtDNAcn) across the lifespan (**Figure 3D**). As cells aged, they accumulated 113-347% more copies of mtDNA (average=+178.1%, p<0.0014, **Figure 1E & Extended Data Figure 1D**). Since cell volume decreased with aging, the density of mitochondrial genome per unit of cellular volume increased by a striking 2.7-12.1-fold across the cellular lifespan (**Figure 3F**), foreshadowing substantial bioenergetic recalibration among aging cells.

### ii. Age-related cell size and shape changes

Here we find that aging fibroblasts get smaller in size as they age. On the surface, this finding contradicts the aging of human stem cells (Mitsui and Schneider 1976) where replicative age is associated with larger size. However, Lengefeld et al. (Lengefeld et al. 2021) found that aged stem cells show similar decreased growth and increased maintenance pathways with replicative aging. These differences in age-related size changes may instead reflect different strategies among different cell types, whereby stem cells rely on glycolytic-derived metabolism to maintain dormancy and proliferative potential by diluting cellular contents (Neurohr et al. 2019), as opposed to fibroblasts (both *in vitro* and *in vivo*) which rely heavily on high-growth OxPhos-derived metabolism to increase cell density in support of the epithelial structure. We also note that rather than recede towards the most thermodynamically neutral and efficient shape, the spheroid, aged primary human fibroblasts adopt a characteristic flattened morphology with substantially greater complexity and covering greater surface area than young cells (see **Extended Data Figure 13**). Greater shape complexity entails increased surface area, which incurs energetic costs associated with maintaining ionic balance across the plasma membrane, as well as energetic costs associated with producing and maintaining physical membrane deformations. This morphological response could also be indicative of energetically challenged cells trying to uptake more resources, again owing ultimately to increased maintenance costs. Thus, a number of unaccounted processes could be contributing to age-related hypermetabolism.

### iii. Aged fibroblasts exhibit an exponential rise in metabolic rate

To derive stable estimates of metabolic rates (MR) *in vitro*, we estimated total ATP synthesis rates from both oxidative phosphorylation (oxygen consumption rate, OCR) and glycolysis (extracellular acidification rate, ECAR) (Mookerjee et al. 2018)(**Extended Data Figure 1E**). By concurrently measuring MR and cell size, we derive energy expenditure per unit volume. The volume-specific MR was ∼58.1 J_ATP_ / μm_3_ in early life. All donors examined exhibited volume-specific MR within a relatively narrow range (40.4-57.2, C.V. = 12%), except for Donor 2’s short-lived fibroblasts, which exhibited a high MR (**Figure 1G**). Remarkably, small inter-individual variations in initial MR per unit volume inversely correlate with the final Hayflick limit of a given cell line, where volume-specific MR explains 67% of the variance (r^2^) in the maximal number of population doublings (**Extended Data Figure 2A**). This high correlation is on par with the many other metrics of replicative lifespan including initial telomere length of a given cell line (r^2^ = 0.73, **Extended Data Figure 2B-D**). The reduction in cell size reveals that cellular energy demand per unit of cell volume increases by up to 15-fold from early to late life (p<0.014, avg. 509% increase, **Figure 1H**). This age-related rise in MR is disproportionately due to glycolysis-derived ATP (J_ATPglyc_). Whereas young cells derive on average 39% of total J_ATP_ from glycolysis, J_ATPglyc_ contributes on average 63% of total J_ATP_ in late life (**Extended Data Figure 1F**). Also consistent with this, mitochondria-derived J_ATPox_ relative to mtDNAcn revealed that in aged fibroblasts, the mitochondrial ATP output per mtDNA copy was on average 58% lower, pointing to lower “efficiency” of each mitochondrial genome (**Extended Data Figure 1G**). To put these *in vitro* metabolic rates over the cellular lifespan in context, we next compare these rates to *in vivo* MR measures.

### iv. RNAseq and DNAm rescaling factors and RNAvelocity validation

We assessed how the rescaling factor changes depending on which portion of the cellular lifespan was used for rate estimations (**Extended Data Figure 4C-D**). Due to the longitudinal design of our *in vitro* datasets, generalized additive modeling (GAM) was performed to identify nonlinear age-related genes and DNAm markers. This restriction ensured that the same genes and CpG sites were used to quantify rescaling factors and did not have any bias toward a particular portion of lifespan. The in vivo-to-in vitro speedup indexed by rescaling factors peaked in early to mid-phase of cellular lifespan, suggesting that cells may be aging faster because they are dividing faster.

Accordingly, across both RNAseq and DNAm datasets and regardless of how age-related markers were selected, the late-phase of cellular lifespan, when division rate is lowest, showed the lowest rescaling factor, with a reduction in rescaling factors by 46 and 72% relative to early life, respectively. This result indicated that as cells approach growth arrest, they exhibit a global slowdown of age-related gene expression and methylation changes. To validate this observation, we used an entirely orthogonal computational approach and estimated the rate of unspliced to spliced mRNA by RNAvelocity (Manno et al. 2018) for each sample across the cellular lifespan (**Extended Data Figure 5**). Global splicing-related factors is downregulated with cellular aging and regulates the onset of replicative senescence (Kwon et al. 2021). RNAvelocity confirmed that the greatest period of speedup occurred in early life and subsequently decreased by an average of 60% during replicative arrest.

### v. Age-related secretion factors contributing to maintenance costs

This age-related increase in maintenance cost could be explained by the expected rise in signaling proteins released from aged quiescent cells known as the senescence-associated secretory profile (SASP) (Coppé et al. 2010). To characterize the secretory activity of aging cells across the lifespan, we designed a custom Luminex cytokine array designed based on comprehensive proteomics characterization of aging human plasma (Tanaka et al. 2018). Of the 27 cytokines detected in extracellular media, we found that compared to young cells (<50 days grown), aged cells (>150 days grown) secreted on average ∼11.4-fold higher amounts of cytokines on a per-cell basis, including several pro-inflammatory cytokines, chemokines, and proteoglycans which require translation and post-translational modifications (Payea et al. 2021) (**Extended Data Figure 7A-B**). Upregulated cytokines included the canonical pro-inflammatory cytokines IL-6 and IL-8. The age-related metabokine GDF15 was also secreted in a time-dependent manner, exponentially rising 15-fold (p<0.011) across the cellular lifespan, consistent with changes occurring in aging human plasma (**Extended Data Figure 7C**). We further measured the release of cell-free mitochondrial DNA (cf-mtDNA) across the lifespan and observed an exponential rise (+10.6-fold, p<0.0041, **Extended Data Figure 7D**). Parallel measurements of cell-free nuclear DNA (cf-nDNA) showed that the released mitochondrial-to-nuclear genome ratio was on average 8.3-fold higher in aged cells than young cells (p<0.012), indicating partially selective mtDNA release. Based on the fixed cost of protein synthesis and extracellular concentrations observed, we estimate that as few as 100 upregulated proteins could account for the dramatic increase in maintenance costs. Together with the dramatic reduction of growth rates by >95%, these data implicating elevated energetic costs to sustain the SASP validate a global energetic re-partitioning away from growth and towards maintenance-related biological processes.

### vi. Gene expression changes for synthesis and maintenance pathways

To gain further insight into the potential origin of increased maintenance costs beyond increased secretory activity, and to examine transcriptional changes associated with increased MR, we sequenced bulk RNA across the lifespan in 3 of the 5 cell lines (∼8 timepoints per cell line, **Extended Data Figure 8A**). We then performed gene set enrichment (GSE) analysis on the age-related genes identified by generalized additive modeling (nonlinear trajectories) using both KEGG (**Extended Data Figure 8B & Supplemental Files 4-5**) and gene ontology (GO) processes (**Extended Data Figure 9 & Supplemental Files 6-7**). Age-related upregulated genes belonged to several canonical maintenance pathways: ‘Lysosome’ (fdr=7.8e-9, KEGG:04142), ‘Autophagy’ (fdr=7.8e-7, KEGG:04140), ‘Mitophagy’ (fdr=3.6e-3, KEGG:04137), and ‘mTOR signaling pathway’ (fdr=2.2e-3, KEGG:04150) (Figure 4F). Similarly, enriched gene ontology (GO) processes amongst upregulated genes included ‘Autophagy’ (fdr=8.0e-9, GO:0006914), ‘Vacuole organization’ (fdr=3.8e-6, GO:0007033), and ‘Protein transport’ (fdr: 5.8e-6, GO:0015031), consistent with a hypersecretory phenotype (**Extended Data Figure 7**). Downregulated genes mainly mapped to protein synthesis pathways: ‘Ribosome biogenesis in eukaryotes’ (fdr=2.6e-15, KEGG:03008) and ‘Ribosome’ (fdr=2.6e-15, KEGG:03010) (Extended Data Figure 8B). Enriched GO processes amongst downregulated genes included: ‘Cellular biosynthetic process’ (fdr=2.6e-15, GO:0044249), ‘Gene expression’ (fdr=3.6e-19, GO:0010467), and ‘Translation’ (fdr=2.9e-15, GO:0006412) (**Extended Data Figure 9**). This hypothesis-free enrichment analysis supports our model’s predictions for age-related increase of maintenance costs and a simultaneous decrease in growth costs.

The age-related changes of ribosomal content were further examined using the transcript levels of all ribosomal subunits (**Extended Data Figure 8C**). Interestingly, ribosomal mRNA expression shows a biphasic trajectory, increasing during the stable growth phase (when cells divide at a constant rate) until the inflection point when division rate slows down, then declining rapidly upon the induction of senescence (see **Supplemental File 8** for additional cellular processes). However, in actively dividing cells, a substantial portion of the ribosomal machinery is produced to supply both daughter cells (i.e., growth-related), whereas in non-dividing cells all the ribosomal machinery can be actively devoted to (maintenance-related) protein synthesis. Therefore, to estimate the proportion of ribosomal machinery available for maintenance-related protein synthesis, we normalized transcript levels per growth rate (i.e., population doubling rate). This correction highlights a continuous, cell division-independent rise in ribosomal gene expression (**Extended Data Figure 8D**). One interpretation of this result is that it reflects the preparatory buildup of translation machinery in aging cells, commensurate with the rising maintenance costs.

We further defined the age-related gene expression changes for mtDNA-encoded genes (**Extended Data Figure 10A**). By categorizing all 37 genes into their RNA species type (mRNA, rRNA, or tRNA), we show a preferential increase in mitochondrial tRNA and rRNA genes, while mRNA are stably expressed with age (**Extended Data Figure 10B**). Because the number of mtDNA copies per cell increased by 518%, this reflects a 53% decrease of mRNA yield per mtDNA molecule (**Extended Data Figure 10C**), which could be explained by either a decrease in mtDNA transcription or loss of mRNA stability in aging cells.

### vii. Potential mechanism of speedup

The question remains as to why higher metabolic rates and cell isolation lead to shorter lifespans. Our data support both i) cumulative damage and ii) evolutionary programmed aging.

The damage mechanism of aging postulates that as cells grow they accumulate damage proportional to the total amount of metabolic activity (G. West 2018; Hou 2013). Correspondingly, an increasing percentage of a cell’s energy budget is used for repair and turnover of cellular machinery with age. This hypothesis has been justified by the scaling relationship that total metabolic expenditure over a lifespan is constant across organisms of vastly different sizes and can even be used to predict invariant cancer risk across organisms (G. B. West, Brown, and Enquist 2001; Geoffrey B. West, Woodruff, and Brown 2002; Kempes, Dutkiewicz, and Follows 2012; Kempes, West, and Pepper 2020). This perspective proposes that as metabolic rate shifts, so do all of the other cellular rates, including growth and damage, such that a single cell always experiences the same lifetime totals but reached at different rates. Here we provide a much more direct validation of the theory by showing that over the cellular lifespan there is an age-dependent metabolic switch from growth to maintenance processes.

For evolutionary considerations of aging, which need not be in conflict with the damage hypothesis, there is a long history of proposed mechanisms including: i) the inability of natural selection to operate on late-life traits, ii) programmed death as a way to benefit populations, or iii) fundamental tradeoffs between life-history traits that benefit young individuals but later detract from older individuals (Mitteldorf 2010; Ljubuncic and Reznick 2009; Gavrilov and Gavrilova 2002). In the context of our work here it is important to note that all of these mechanisms will be affected by a rescaling of the physiological clock. If there are problematic late-life traits that have not been selected against by evolution these will simply be reached more quickly, and the same would be true for programmed death. Similarly, since most life-history traits scale with metabolic rate across organisms of different sizes, it is reasonable to assume that metabolic rate would adjust the rates associated with life-history tradeoffs that produce aging. All three proposals are consistent with our finding showing a striking temporal alignment of rising metabolic costs and rapid reduction in cell division (see **Figure 6**).

Regardless of the underlying reason for cellular aging and considering that many of these mechanisms could be fundamentally interconnected and co-optimized by evolution, we speculate that cells in the human body are able to age at a significantly slower rate because of the collective partitioning of biological functions across multiple functionally specialized tissues and organs, making the overall state of living more energetically efficient. For example, in populations of syngeneic yeast, metabolite exchange where some cells produce and consume different metabolites creates a state of metabolic cooperation that extends lifespan of individual cells within the population (Correia-Melo et al. 2022). In cells and animal communities, cell growth and fitness follow a density-dependent relationship known as the Alee effect (Kramer, Berec, and Drake 2018). Imposing severe isolation by culturing human cells at very low density impedes growth and compromises survival, even of cancer cells (K. E. Johnson et al. 2019). Long-lived mammalian organisms also exhibit extensive metabolic cooperation, where specific organs are responsible for the transformation and synthesis of specific metabolites and hormones, which are supplied at no or little cost to the cells and tissues consuming them (Jang et al. 2019). Thus, various cell types operate as an integrated network of metabolically and physiologically interconnected units, from which emerges a greater state of metabolic efficiency and lower resting metabolic rate per unit of body mass (Geoffrey B. West, Woodruff, and Brown 2002). In contrast, the spatial isolation and relative uniformity of cell monocultures would represent a more metabolically stressful state, where each cell must subserve every function necessary for its own survival. Further evidence for this explanation comes from hibernating organisms, where across five orders of magnitude in mass, the cellular metabolic rate of hibernation is lower than the lower limit of MR in isolated mammalian cells (Nespolo, Mejias, and Bozinovic 2022). This notion also draws from evidence in regenerative biology where cohabiting cells within the organism behave as an interconnected collective driven by computations taking place at the whole-body level, rather than at the single-cell level (Levin 2021). This understanding implies that experimentally adding energetic constraints that mimic the *in vivo* environment, such as contact inhibition and nutrient turnover, can lower energetic pressure and allow cells to live longer.

As discussed, a few different factors could contribute to age-related hypermetabolism. Here we identify maintenance-related processes as the main driver, specifically mitochondrial content accumulation with decreased energy-production efficiency, and upregulation of autophagy and extracellular secretions. These drivers of hypermetabolism include transcriptional activation of several stress response pathways and the production and secretion of cytokines and chemokines, notably the metabokine GDF15, which is the most robustly upregulated circulating protein in the blood of aging humans (Tanaka et al. 2020). We find that the secretion of an average cytokine is roughly 1% of metabolic rate, implying that as few as 100 high-abundance secreted cytokines (Basisty et al. 2020; Tanaka et al. 2018) could dominate the cellular energy budget. Here we also report a novel senescence-associated secretory molecule, cell-free mtDNA, a signature that matches cross-sectional lifespan findings in human blood (Pinti et al. 2014). Other processes that could not be measured in this study, including molecular repair and degradation processes, ion cycling across membranes, and cell migration could also contribute to the exponential rise in energetic costs at the end of life - even in the absence of cell division.

Previous work has established an upper limit of Gibbs free energy (Yang et al. 2021; Niebel, Leupold, and Heinemann 2019), where flux redistribution allows the cell to switch between different energy pathways (i.e., respiration and fermentation) but is ultimately limited and must avoid a critical Gibbs energy dissipation rate. Our data suggest that although cells within the organism operate at only 5-10% of maximal rates, cultured cells operate much closer to this upper limit (∼50%). If this upper limit is imposed by physical constraints on metabolism, the fidelity of molecular operations, and on cell division (Morowitz 1978; DeLong et al. 2010; Lan et al. 2012; Kempes, Dutkiewicz, and Follows 2012; Kempes, Wolpert, et al. 2017; Yang et al. 2021), this could explain: i) the 258-fold reduction in lifespan of cultured cells, ii) why cells slow down rates of change as they enter quiescence (see **Extended Data Figure 4D**), and iii) why treatments that induced higher MR failed to speedup DNAm rescaling factors any further (see Figure 5H). In support of this, and congruent with our results indicating increased production and secretion of protein material in old cells, human fibroblasts in replicative senescence were brought out of senescence by mildly inhibiting protein synthesis (Takauji et al. 2016), thus freeing up a portion of the energetic budget. Thus, our results are internally consistent with a link between total energy flux or MR, and lifespan regulation.

However, recent work in rodents suggests that temperature, rather than MR, may be a more important determinant of lifespan (Zhao et al. 2022). Elevating body temperature reduces lifespan, even in the context of reduced MR. In cultured fibroblasts, although the overall temperature of the cellular system should be well-controlled at 37°C, we cannot rule out the possibility that intracellular temperature could be affected in parallel with MR. This may be particularly relevant since mitochondria are the warmest compartment of the cell, operating at temperatures estimated around 50°C (Chrétien et al. 2018). Thus, a complete mechanistic model of the MR-lifespan connection, which is beyond the scope of the current analysis, will likely need to incorporate both MR and temperature. In addition, it has also recently been shown in *C. elegans* that other factors such as stressors, as well as temperature, adjust the rate of aging (Stroustrup et al. 2016). Amazingly, organisms appear to follow a universal aging curve rescaled by these various environmental, energetic, and stress factors (Stroustrup et al. 2016), and this work should be integrated with our perspective here to understand how multiple effects adjust aging in humans beyond the basic energetic arguments.

### vii. Detailed methods

#### In vitro fibroblast cultures

Human fibroblasts were obtained from a local-clinic (IRB #AAAB0483) and commercial distributors from five healthy donors (**Table 1**) and stored in liquid nitrogen before culturing. Cells were cultured in T175 flasks (Eppendorf #0030712129) at standard 5% CO_2_ and atmospheric O_2_ at 37°C in DMEM (5.5 mM physiological glucose) supplemented with 10% FBS (Thermofisher #10437010), 50 μg/ml uridine (Sigma-Aldrich #U6381), 1% MEM non-essential amino acids (Life Technologies #11140050), 10 μM palmitate (Sigma-Aldrich #P9767), and 0.001% DMSO (treatment-matched, Sigma-Aldrich #D4540). For metabolic rate measurements in Seahorse analyzer (Agilent), cells were grown in complete XF media without pH buffers and supplemented with 5.5 mM glucose, 1 mM pyruvate, 1 mM glutamine, 50 μg/ml uridine, and 10 μM palmitate to ensure that cells have access to a variety of energetic substrates.

**Table 1.** Cell line characteristics for all donors.

| Cell line name | Received from | Source tissue | Sex | Donor age | Passage | Cat# |
| --- | --- | --- | --- | --- | --- | --- |
| Donor1 (HC1, hFB12) | Lifeline Cell Technology | Dermal breast | male | 18 | 1 | FC-0024 Lot # 03099 |
| Donor2 (HC2, hFB13) | Lifeline Cell Technology | Dermal breast | female | 18 | 1 | FC-0024 Lot # 00967 |
| Donor3 (HC3, hFB14) | Coriell Institute | Foreskin | male | 0 | 4 | AG01439 |
| Donor4 (HC5, hFB1) | Hirano Lab | Dermal upper-arm | male | 29 | 3 | NA |
| Donor5 (HC6, hFB2) | Hirano Lab | Dermal upper-arm | female | 26 | 4 | NA |
| Patient1 (SURF1-1) | Hirano Lab | Dermal upper-arm | male | 1 | 7 | NA |
| Patient2 (SURF1-2) | Hirano Lab | Dermal upper-arm | male | 11 | 5 | NA |
| Patient3 (SURF1-3) | Hirano Lab | Dermal upper-arm | female | 9 | 9 | NA |

Cells were grown for up to 265 days by repeated passaging every 5-7 days. To determine the number of cells to plate at each passage, growth rates from the previous passage were used, pre-calculating the expected number of cells needed to reach ∼90% confluency (∼3 million cells) by the next passage, ensuring relatively similar confluence at the time of harvesting for molecular analyses. Cells were never plated above 2.5 million cells to avoid plating artifacts of contact inhibition. Individual cell lines from each donor were grown until they exhibited less than one population doubling over a 30-day period, at which point the cell line was terminated, reflecting the end of the lifespan. The Hayflick limit was calculated as the total number of population doublings reached by the end of each experiment.

Repeat cellular lifespan experiments were performed three times (Study Phases II,IV,V) for Donors 1-3 under untreated conditions. In study phase V, cells were seeded to reach a lower maximum confluency of <80%.

#### Cytology

Cell counts, volume, and proportion of cell death were determined in duplicates (CV <10%) and averaged at each passage using Countess II Automated Cell Counter (Thermofisher #A27977). Cell density measures were estimated by taking the weighted sum of total extracted protein (BCA assay, Bioworld #20831001-1), RNA (RNeasy kit, Qiagen #74104), and DNA (DNeasy kit, Qiagen #69506) at each timepoint. Further, at each passage, brightfield images were taken at 10x and 20x using an inverted phase-contrast microscope (Fisher Scientific #11350119).

### Treatments

MR-altering treatments design, dose, and duration are listed in **Table 2**. Hypermetabolic treatments included chronic 1nM oligomycin (Oligo, +134% MR), chronic 100nM dexamethasone (DEX, +150% MR), chronic mitochondrial nutrient uptake inhibitors (mitoNUITs, +110% MR), and fibroblasts taken from SURF1-mutant patients (+106% MR). Hypometabolic treatments included chronic 2-deoxy-d-glucose treatment (2DG, -27% MR) and contact inhibition (-10% MR) (**Figure 4B**). Contact inhibition was induced by allowing dividing cells to fill the culture flask (i.e., confluency), which reduces the growth rate by an average of 87% (from 0.65 to 0.085 div / day). This induced a quiescent cell cycle arrest allowing the determination of time-dependent changes independent of cell division. Treatments began after 15 days of culturing post-thaw to allow for adjustment to the *in vitro* environment. Media was changed weekly and cells were placed in biohood for the same amount of time for cells undergoing standard passaging.

**Table 2.**
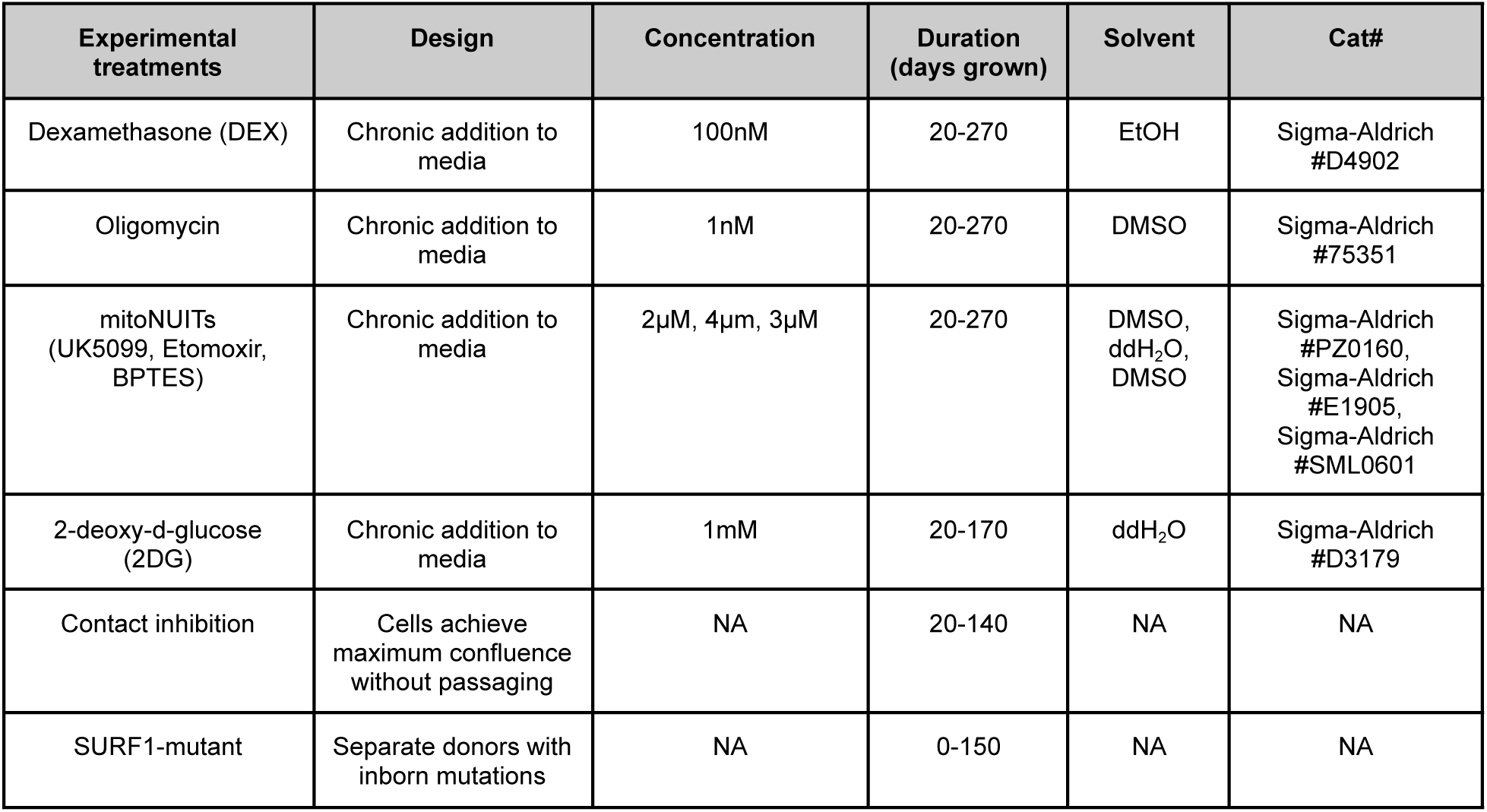
Experimental design of MR-altering treatments. *Chronic indicates treatment was reapplied at each passage

### Metabolic rates

Bioenergetic parameters of cultured fibroblasts were measured using the XFe96 Seahorse extracellular flux analyzer (Agilent) (Tan et al. 2015). Oxygen consumption rates (OCR) and extracellular acidification rates (ECAR, i.e., pH change) were measured over a confluent 20,000k cell monolayer every ∼15 days, following the manufacturer’s instructions. Respiratory states such as basal and maximum respiration rates were all determined using the MitoStress Test (Brand and Nicholls 2011) which involves the sequential additions of the ATP synthase inhibitor oligomycin (final concentration: 1 μM), the protonophore uncoupler FCCP (4 μM), and the electron transport chain Complex I and III inhibitors rotenone and antimycin A (1 μM). The final injection included Hoechst nuclear fluorescent stain (Thermofisher #62249) to allow for automatic cell counting using the Cytation1 Cell Imager (BioTek). Raw bioenergetic measurements were normalized to relative cell counts on a per-well basis.

ATP production rates from oxidative phosphorylation (OxPhos, *J*_ATPox_) and glycolysis (*J*_ATPglyc_), as well as total cellular ATP production and consumption (*J*_ATPtotal_), were estimated using the method described by Mookerjee et al. (Mookerjee et al. 2018). Briefly, the method relies on the phosphate-to-oxygen (P/O) ratios of OxPhos and glycolysis, using oxygen consumption and proton production rates (PPR) as input variables. The same constants were used for all estimations, assuming glucose as the predominant carbon source and constant coupling efficiency. Changes in substrate consumption along the lifespan would require parallel assessments of metabolic flux to resolve. This assumption should have only a minor influence on calculated ATP production rates. We further note that maximum *J*_ATP_ measures were determined by adding the *J*_ATPox_ after FCCP injection with the *J*_ATPglyc_ after oligomycin injection. We note that a separate metabolic assay would be required to assess true maximum glycolytic ATP production *in vitro*, which is beyond the scope of the current experimental capacity. This metric of maximum energetic capacity is therefore limited and should be interpreted with caution (Schmidt, Fisher-Wellman, and Neufer 2021; Mookerjee et al. 2018).

Baseline whole-body metabolic rates were obtained from indirect calorimetry in three small human cohorts reporting values consistent with literature norms (Taivassalo et al. 2003; Jeppesen et al. 2009; Rising et al. 2003). Max whole body MR (i.e., VO_2_peak) estimates were obtained from Kaminsky et al. (Kaminsky et al. 2021). An estimation of skin-specific MR was calculated using Q_O2_ values for human *in vivo* tissues (see Table 4 in (Davies 1961). Additionally, skin-specific MR was calculated using the mechanistic model: REE = Σ(K(i) × T(i)), where REE is whole body resting energy expenditure measured by indirect calorimetry and T(i) is the mass of individual organs and tissues measured by magnetic resonance imaging (Wang et al. 2010). In this estimate, skin-specific MR is determined based on Elia’s Ki values in which skin tissue Ki is estimated along with intestines, bones, and lungs, which together have Ki (12) similar to inactive skeletal muscle (13). To compare mass-specific MR *in vivo* to volume-specific MR in cultured fibroblasts, *in vivo* and *in vitro* units were converted using the attached **Supplemental File 1** (‘SF1_Cell-to-Body_Oxygen_Consumption_Calculator’). This conversion assumes an average cell mass of 10ng (Geoffrey B. West, Woodruff, and Brown 2002; Park et al. 2008).

#### Estimation of growth and maintenance costs

Energetic budgets of growth and maintenance were estimated using the mathematical growth modeling originally describing unicellular organisms (Kempes, Dutkiewicz, and Follows 2012) which incorporates the basic energetic partitioning of the Pirt model (Pirt 1965) along with metabolic scaling principles (Geoffrey B. West, Woodruff, and Brown 2002). See Equation 3 of Box 1 for the calculation of the energetic budget of cultured cells.

#### Telomere length

Relative telomere length was evaluated on the same genomic material used for other DNA-based measurements. Measurements were performed by qPCR and expressed as the ratio of telomere to single-copy gene abundance (T/S ratio), as previously described (Cawthon 2002; Lin et al. 2010). Triplicate values of T and S for each sample were averaged to calculate the T/S ratios after a Dixon’s Q test for outlier removal. The T/S ratio for each sample was measured twice.

Telomere length rates of erosion were calculated as the rate of change in T/S ratio per day grown from days 10-70. Telomere erosion rates for physiological aging were obtained from the GTEx dataset outlined in Demanelis et al. (Demanelis et al. 2020), which contains relative telomere lengths of skin (sun-exposed) from donors ages 20-70 years old.

#### DNA methylation

DNA methylation markers were measured across the genome using the Illumina EPIC microarray (Illumina, San Diego). Briefly, DNA was extracted using the DNeasy kit (Qiagen #69506) and quantified using the QUBIT broad-range kit (Thermofisher #Q32852). Extracted DNA was processed at the UCLA Neuroscience Genomic Core (UNGC) for bisulfite conversion and hybridization using the Infinium Methylation EPIC BeadChip kit. Methylation data was processed in R (v4.0.2), using the ‘minfi’ package (v1.36.0), and normalized using functional normalization (FunNorm). RCP and ComBat adjustments using the ‘sva’ package (v3.12.0) were performed to correct for probe-type and plate bias, respectively. After quality control, DNAm levels were converted to beta values for 865,817 CpG sites.

Age-related DNAm changes were estimated using generalized additive modeling on individual DNAm sites using the R package ‘mgcv’ (v1.8-31). All genes that had a p-value < 0.001 across all three cell lines were identified as significant. Genes were separated into UPregulated and DOWNregulated lists based on the direction of age-related change. The full list of significant DNAm sites can be found in **Supplemental File 3.**

#### DNA methylation clocks and related measures

Epigenetic-based clocks were calculated using DNA methylation data (Horvath and Raj 2018) (**Table 3**). We computed three clocks designed to predict the chronological age, Horvath1 (i.e. PanTissue clock) (Horvath 2013), Horvath2 (i.e. Skin&Blood clock) (Horvath et al. 2018), Hannum (Hannum et al. 2013), and PedBE (McEwen et al. 2020) clocks; two clocks designed to predict mortality, the PhenoAge (Levine et al. 2018) and GrimAge (Lu, Quach, et al. 2019) clocks; a clock to measure telomere length, DNAmTL (Lu, Seeboth, et al. 2019); a clock designed to measure mitotic age, MiAge (Youn and Wang 2018); a clock trained to predict cellular senescence, DNAmSen (Levine et al. 2019), and two DNA methylation measure of the rate of deterioration in physiological integrity, DunedinPoAm (Belsky et al. 2020), and DunedinPACE (Belsky et al. 2022). The PC-based method replaces the clock’s individual illumina probe measurements (5-500 CpGs) with the shared extracted variances among genome-wide CpGs from principal components (PC), yielding the PC-adjusted DNAmAges for each clock (Higgins-Chen et al. 2022).

**Table 3.**
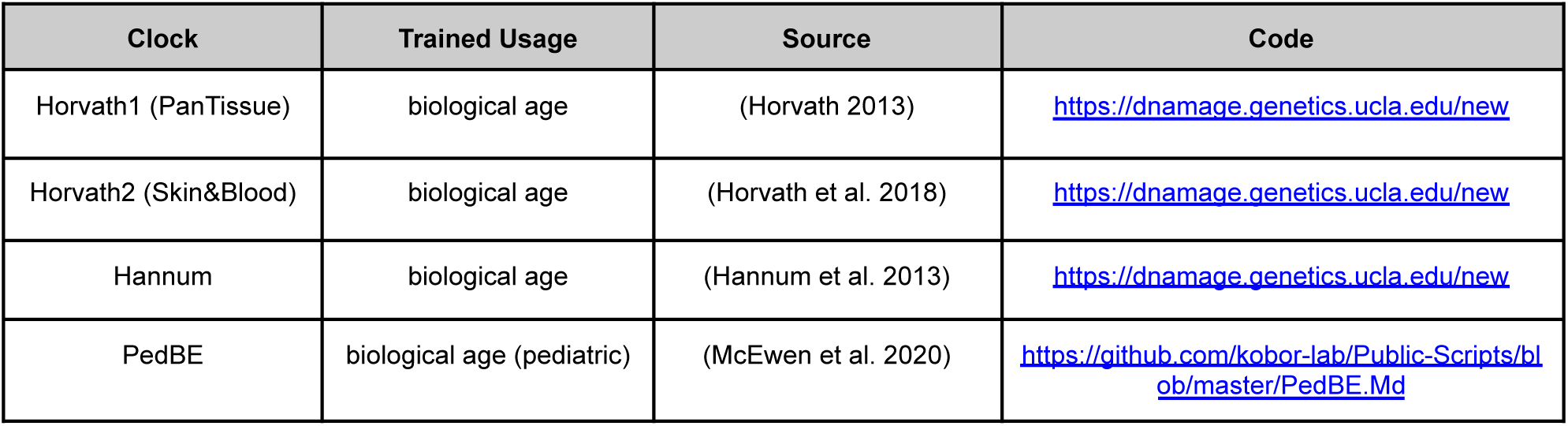
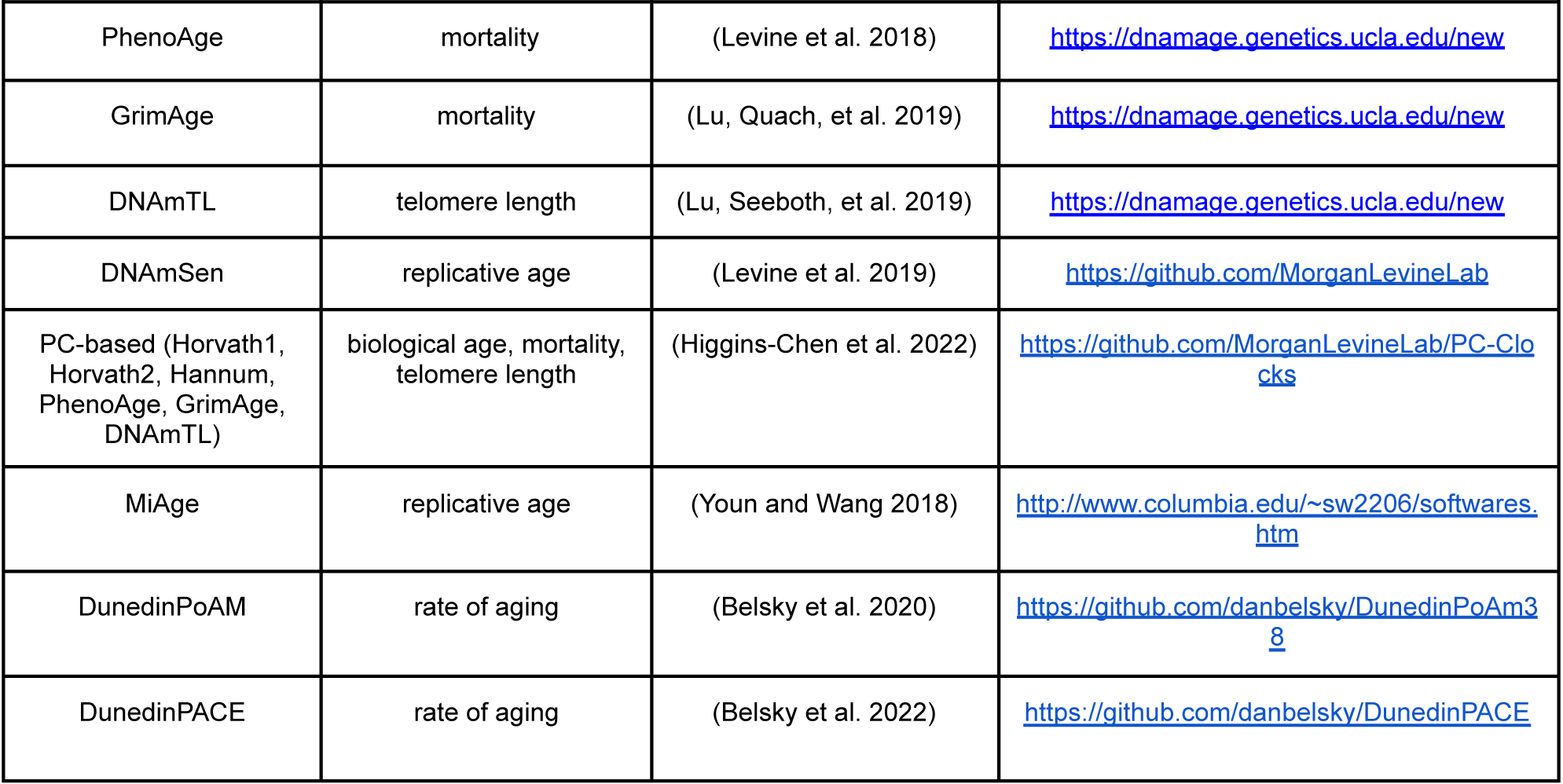
Epigenetic clocks used to estimate the rate of biological aging based on DNA methylation dynamics.

#### In vivo DNA methylation data

TwinsUK DNA methylation data was obtained through the online GEO portal (GSE90124) collected as part of the TwinsUK study (Moayyeri et al. 2013). This dataset includes 322 skin samples collected by punch tissue biopsy of healthy female individuals ages 39-83 years old. Bisulfite-converted DNA from the 322 samples were hybridized to the Illumina HumanMethylation450 BeadChip. All 450,531 CpG sites of the 450k chip were present on the EPIC chip used for *in vitro* DNA methylation.

*In vivo* growth rates were obtained using the Mitotic Age clock (Youn and Wang 2018) on DNA methylation measured across the genome of adult skin and adipose tissue. The adipose dataset is made available by the TwinsUK Adult Twin Registry (ArrayExpress E-MTAB-1866) (Grundberg et al. 2013) and provides Illumina 450K DNA methylation data of 648 adipose obtained from female European twins ages 40-85.

#### mtDNA copy number

mtDNA copy number was measured using a duplex qPCR as previously described (Picard et al. 2014). Briefly, duplex qPCR reactions with Taqman chemistry were used to simultaneously quantify mitochondrial (mtDNA, ND1) and nuclear (nDNA, B2M) amplicons. The final mtDNAcn was derived using the ΔCt method, calculated by subtracting the average mtDNA Ct from the average nDNA Ct. mtDNAcn was calculated as 2^ΔCt^ x 2, yielding the estimated number of mtDNA copies per cell. Note that mtDNA copy number was quantified on the same genomic material used for DNA methylation measurements.

#### Cell-free DNA

Cell-free mitochondrial (cf-mtDNA) and nuclear DNA (cf-nDNA) levels were measured simultaneously by qPCR on sampled media collected across the cellular lifespan. Taqman-based duplex qPCR reactions targeted mitochondrial-encoded ND1 and nuclear-encoded B2M sequences as described previously (Ware et al. 2020; Belmonte et al. 2016; Trumpff et al. 2019). Each gene assay contained two primers and a fluorescent probe and was assembled as a 20X working solution according to the manufacturer’s recommendations (Integrated DNA Technologies). Digital PCR (dPCR) of the standard curve used in cf-mtDNA/cf-nDNA assessment were measured separately using singleplex ND1 and B2M assays using a QuantStudio 3D Digital PCR System and associated reagents (Thermo Fisher, cat#A29154) according to the manufacturer’s protocol.

#### Cytokines

Two multiplex fluorescence-based arrays were custom-designed with selected cytokines and chemokines based on human age-related plasma proteins correlated with chronological age (Tanaka et al. 2018), and based on their availability on R&D custom Luminex arrays (R&D, Luminex Human Discovery Assay (33-Plex) LXSAHM-33 and LXSAHM-15, http://biotechne.com/l/rl/YyZYM7n3). Briefly, media samples were collected at each passage and stored at -80°C. Thawed samples were centrifuged at 500xg for 5 min and the supernatant collected. Media samples were run undiluted on a Luminex 200 instrument (Luminex, USA) as per the manufacturer’s instructions. Data was fitted and final values were interpolated from a standard curve in xPONENT v4.2. Of the 51 measured cytokines, 27 were detected in the media of cultured cells. Cytokine concentrations were then normalized to the number of cells counted at the time of collection to produce estimates of cytokine production on a per-cell basis.

#### Gene expression

Gene expression was estimated by RNAseq selected for functional mRNAs using a Ribo-Zero Gold extraction protocol. Briefly, total genomic RNA was isolated every ∼11 days across the cellular lifespan for control lines and selected treatments. RNA was stabilized using TRIzol (Invitrogen #15596026), stored at -80°C until extraction using an RNeasy kit (Qiagen #74104), quantified using the QUBIT high sensitivity kit (Thermofisher #Q32852), and underwent quality control on Bioanalyzer (Agilent RNA nano kit 6000, #5067-1511) and Nanodrop 2000. cDNA library preparation was performed using a Ribo-Zero Gold purification (QIAseq FastSelect -rRNA HMR Kit #334387) and NEBNext^®^ Ultra^™^ II RNA Library Prep Kit (Illumina #E7770L). cDNA was sequenced using paired-end 150bp chemistry on a HiSeq 4000 instrument (Illumina, single index, 10 samples/lane, Genewiz Inc). Sequencing depth was on average 40 million reads per sample. Sequenced reads were then aligned using the pseudoalignment tool *kallisto* v0.44.0 (Bray et al. 2016), processed using txi import (‘tximport’, v1.18.0, length-scaled TPM), and vst normalized (‘DEseq2’, v1.30.1). Growth rate normalizations were performed using the median-centered doubling rates (divisions / day) at each timepoint.

Gene Set Enrichment analysis (GSEA) was performed using the ShinyGO webtool (http://bioinformatics.sdstate.edu/go/, v0.75). Age-related changes were estimated using generalized additive modeling using the R package ‘mgcv’ (v1.8-31). All genes that had a p-value < 0.1 across all three cell lines were included in the analysis. Genes were separated into UPregulated and DOWNregulated lists based on the direction of age-related change. The full list of significant genes can be found in **Supplemental File 2.** Selected pathway gene sets (**Supplemental File 8**) were obtained from https://maayanlab.cloud/Harmonizome/. Expression values were centered to the median of the youngest control timepoints and then transformed to a log2 scale. For ribosome-related analyses, ribosomal genes were selected from the KEGG database at https://www.genome.jp/kegg/pathway/hsa/hsa03010.html.

Rates of gene splicing and degradation can be derived from the relative abundance of unspliced (nascent) and spliced (mature) mRNA quantified by RNA sequencing. These relative abundances can then be transformed into RNAvelocity, a high-dimensional vector that predicts the future state of cell identity on a timescale of hours (Manno et al. 2018). RNAvelocity has been used to track cellular lineage transitions, circadian rhythms, neurogenesis, embryogenesis development, and tissue regeneration using single-cell sequencing of tissue populations containing multiple cell types (Bergen et al. 2020; Manno et al. 2018). The computational pipeline for obtaining RNAvelocity estimates was as follows: RNAseq fastq files were aligned using STAR v2.7 (https://github.com/alexdobin/STAR/) to a pre-curated annotation of the hg19 genome (Lareau and Brenner 2015) to obtain spliced and unspliced RNA reads. RNAvelocity estimates of mapped reads were then quantified using the velocyto.py smart-seq command (https://velocyto.org/velocyto.py/) which treats each bulk RNAseq sample as if it were a single cell from a SMART-seq experiment. All further analysis of velocyto output was performed in a Jupyter notebook adapted from the scVelo pipeline (https://scvelo.readthedocs.io/VelocityBasics.html).

#### In vivo gene expression data

*In vivo* RNAseq data was obtained from the Genotype-Tissue Expression (GTEx) study (Lonsdale et al. 2013) through the online dbGaP accession #phs000424.v8.p2, under the project #27813 (Defining conserved age-related gene expression trajectories). This dataset includes 12,767 samples obtained postmortem from organ donors across 54 unique tissues from n=980 individuals (653 men, 327 women), ages 20-70 years. Data was filtered to the tissue ‘Skin - Sun Exposed (lower leg)’, yielding 520 samples (330 men, 178 women) ages 21-70 years.

#### Rescaling factors

Rescaling factors were obtained using the rates of change in global gene expression and DNA methylation markers in *in vivo* and *in vitro* data (see schematic in **Extended Data Figure 4A**). *In vivo* data was obtained from the GTEx and TwinsUK studies. Age-related markers were identified through linear modeling across the lifespan in both datasets with a cutoff of 0.1 p-value. Genes and DNAm markers were then grouped by direction of change with age (increase or decrease with age) and then the overlap of markers between the *in vivo* and *in vitro* datasets moving in the same direction with age were selected for final calculations. The rescaling factor was calculated as the ratio of the median *in vitro slopes* to the median of the *in vivo* slopes.

We note several assumptions that need to be considered when evaluating our MR-altering rescaling factor results (**Figure 5G**): i) we use independent genes and methylation sites for each treatment group that is significantly changing with age. An alternative approach of preselecting nonlinear GAM molecules and then estimating slopes off that list was attempted and showed similar results. ii) the final rescaling factor was further documented as the median of all overlapping significant molecules and excludes the rates of change in genes or sites that do not change with age. iii) we report the slopes of only genes and molecules that share the same directional change between *in vitro* and *in vivo* datasets. An alternative approach of measuring our rescaling factors on the union of both *in vitro* and *in vivo* age-related genes and sites was performed and showed minimal difference to our final findings. iv) our analysis assumes that dermal fibroblasts are comparable to biopsies of skin samples, which contain a heterogeneous mixture of fibroblasts, epidermis, and epithelial cells. v) For *in vitro* treatments, the derived slopes for each molecule were taken from the same time period of 25-95 days grown. This portion was selected to ensure a) at least 3 timepoints were included for each donor, b) that the treatment had fully taken effect (treatment started at 15 days), and c) that there were matching timepoints with close to equal distributions of samples in each treatment group. We also note that the treatment 2DG was only performed on 2 donors (Donors 3 & 4) and the SURF1-mutant cells involved comparing entirely different donors with an inherited genetic mutation(s) that could have had developmental effects.

#### Rescaling factor modeling for In vivo data

*In vivo* rates of change were estimated using four parallel approaches: i) linear regression, ii) linear mixed effects, iii) min-max confidence intervals, and iv) permutation modeling. Linear regression was performed using R (v) built-in lm function. Linear mixed effects modeling was performed using the ‘lmer’ function from the R package ‘lme4’ (v1.1) with a fixed effect for age and random effects for type of collection (postmortem or organ donor), donor relationship (i.e. twin, sibling, etc.), and sex of the donor. Significance values were determined using an ANOVA to a null model with the same random effects. Min-max confidence interval slope estimation was determined by taking the minimum and maximum of the 95% intervals at the first and last timepoint of a linear regression of the *in vivo* data. The slope was then estimated as the maximum absolute linear slope from either the minimum of the first timepoint to the maximum of the last timepoint or vice versa. This approach ensures the fastest possible slope in 95% of sampling. Permutation modeling entailed performing linear regression across time on a random 50 samples of either the full 520 (for RNAseq) or 322 (DNAm), bootstrapped 1000 times. To account for the high variance of *in vivo* data which likely overestimates rescaling factors (i.e., the rate of change), we subsampled the top 10% of the fastest slopes of the resulting distribution of which the median was used for the final rescaling factor for each gene/methylation site.

#### Statistical Analyses

All statistical analyses were performed using GraphPad Prism (v9.0) or RStudio (v1.3.1056) using R (v4.0.2). Comparisons of end of life to early life were performed using two-tailed paired ratio t-tests using the minimum and maximum values across the lifespan. If no clear maximum or minimum value was present in the trajectory, then the average of the first or last three timepoints was used. Data visualization and statistical analyses were generated in R (‘ggplot2’, v3.3.5) and Prism 9.

## Notes

https://columbia-picard.shinyapps.io/shinyapp-Lifespan_Study/

https://github.com/gav-sturm/Cellular_Lifespan_Study

https://figshare.com/articles/dataset/Lifespan_Study_Data/18441998

https://www.nature.com/articles/s41597-022-01852-y

https://www.nature.com/articles/s42003-022-04303-x

https://www.nature.com/articles/s43587-022-00248-2

### Competing Interest Statement

The authors have declared no competing interest.

### Summary of Updates

Reorganized the main text and figures to simplify the progression of the manuscript.

