## Extended figures 1-13 for "Accelerating the clock: Interconnected speedup of energetic and molecular dynamics during aging in cultured human cells"

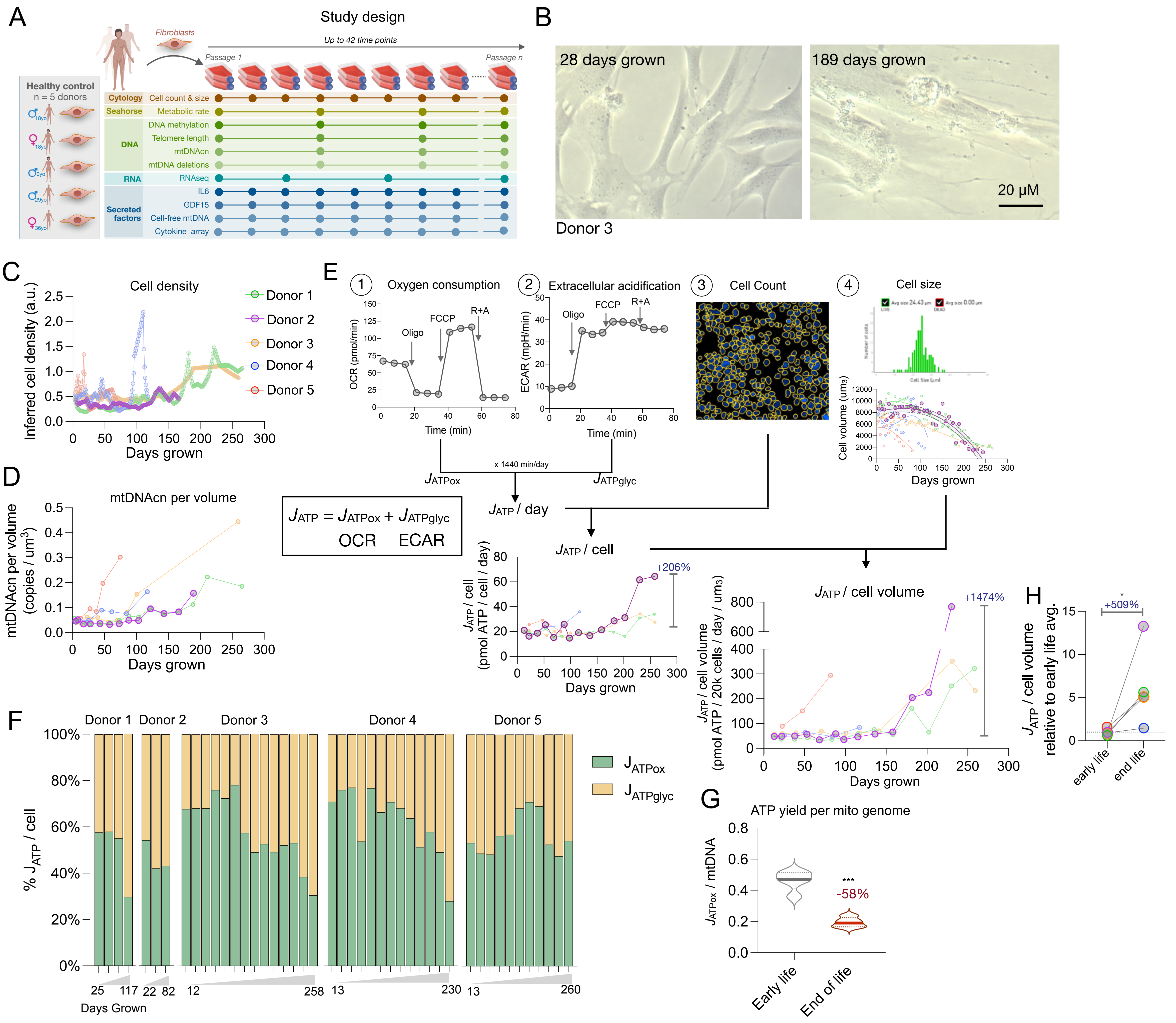


**Extended Data Figure 1. Study design and derivation of metabolic rates** (**A**) Schematic of Cellular Lifespan Study design showing repeat measures collected as cells are passaged until replicative exhaustion. (**B**) Bright-field images of Donor 5 fibroblasts as young (left panel) and aged (right panel) morphology. Note how young cells have a spindle shapes that align side-by-side , while aged cells flatten with spider web-like protrusions. (**C**) Schematic of metabolic rate (MR) measurements derived from parameters measures using the Seahorse XFe96. (G1-2) Raw oxygen consumption rate (OCR) and extracellular acidification rate (ECAR) are converted into OxPhos-derived ($J_{ATPox}$) and glycolysis-derived ($J_{ATPglyc}$) ATP production rates, respectively, and combined into total ATP production rate ($J_{ATPtotal}$). MR measures are then normalized to the number of cells in each measurement (E3), and the average cell volume at each passage (E4). Applied to longitudinal trajectories in each donor, this yields the total ATP consumption/production rate per cell volume ($J_{ATP-Total}$ /cell volume) and closest available estimate of mass-specific metabolic rates. (**D**) Indirect measure of cell density across cellular lifespan. Values are determined as the weighted sum of DNA, RNA, and protein mass per cell at a given timepoint. (**E**) Mitochondrial DNA per unit volume across the cellular lifespan. (**F**) Stacked barplot showing % of the rate of ATP production derived from OxPhos ($J_{ATPox}$, green) and ATP derived from glycolysis ($J_{ATPglyc}$, yellow). Blue triangle on x-axis indicates the age of the cells in days grown (increasing left-to-right). (**G**) Energetic efficiency of young and aged cells, measured as the ATP yield per a mitochondrial genome. Paired t-test, Hedge's G.

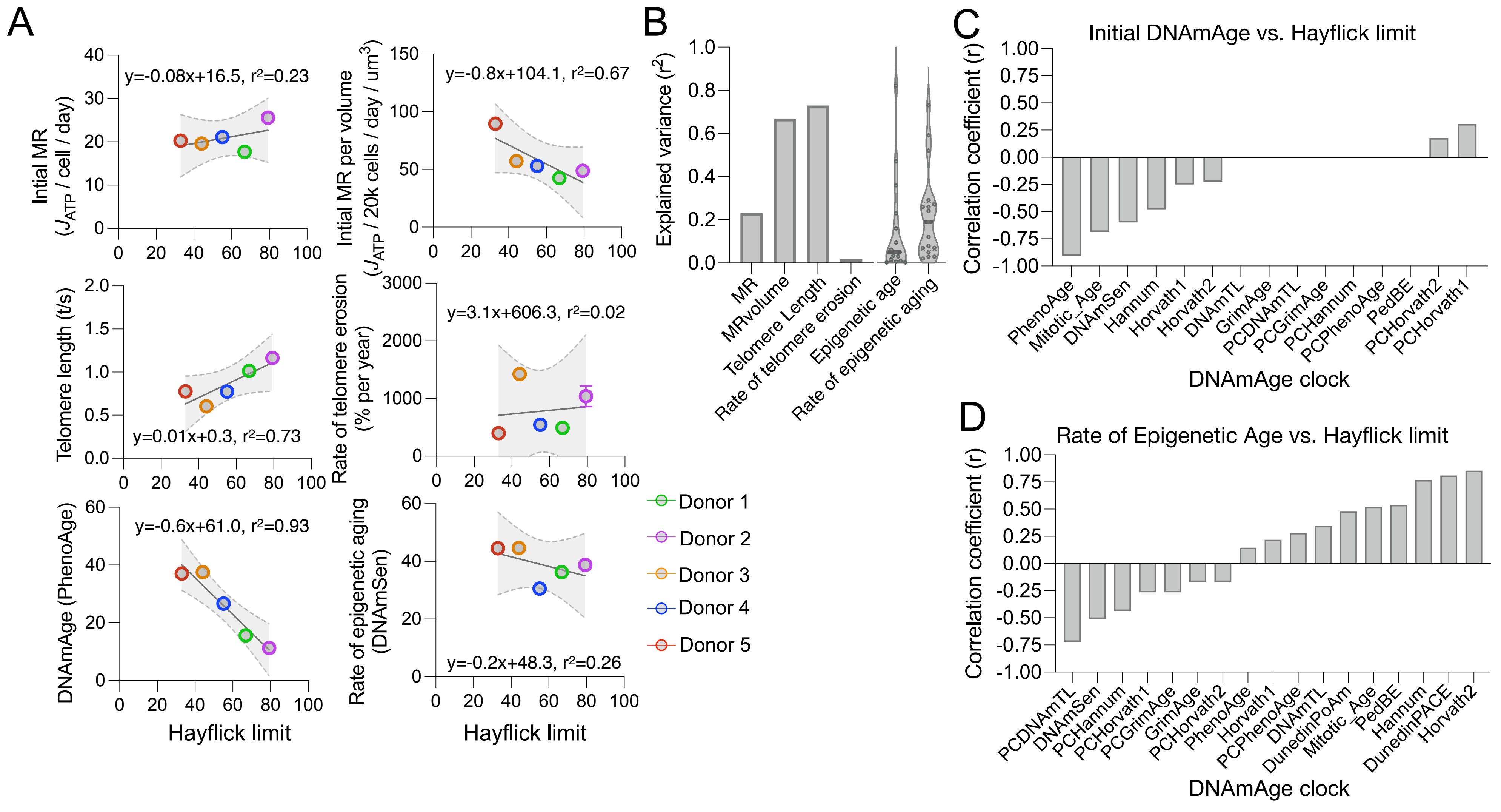


**Extended Data Figure 2. Prediction of hayflick limit from MR and aging measures.** (**A**) Correlation of hayflick limit (n=5 cell lines with metabolic rate; per cell (top-left) and per cell volume (top-right) (10-70days, 2-5 timepoints/cell line), telomere length; initial y-intercept (middle-left), rate of telomere length (middle-right) (10-70days, 4 timepoints/cell line), DNAmAge; initial y-intercept (bottom-left), and rate of epigenetic aging (bottom-right) (10-70days, 4-5 timepoints/cell line) (**B**) Explained variance in hayflick for each parameter. Epigenetic age and rate values are shown as a violin plot with each point indicating a different DNAmAge clock. (**C**) Correlation coefficient for each DNAmAge clock initial epigenetic age (y-intercept 10-70 days grown, 4-5 timepoints/cell line) with hayflick limit (n=5 cell lines). (**D**) Correlation coefficient for each clock's rate of epigenetic aging (slope, 10-70 days grown, 4-5 timepoints/cell line) with hayflick limit (n=5 cell lines). Note, Dunedin clocks are measures of the rate of aging and therefore were calculated using the mean of each donor for selected timepoints (10-70 days grown, 4-5 timepoints/cell line).

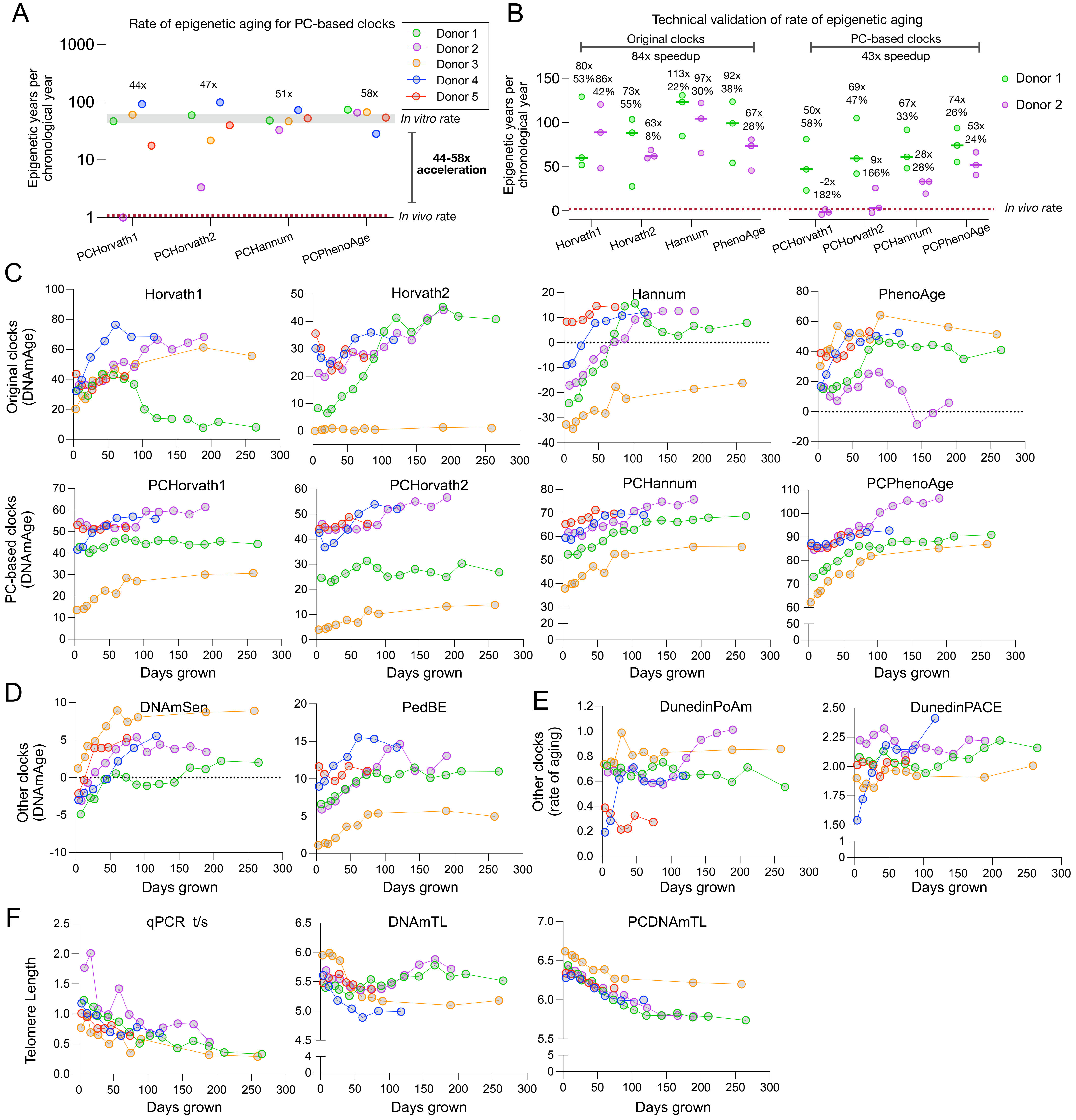


**Extended Data Figure 3. Rate of epigenetic aging of cultured fibroblasts.** (**A**) Rate of epigenetic aging for cultured fibroblasts as measured by several PC-based epigenetic clocks. Rates are defined as the slope of the linear fit line from days 10-70 of growth in culture. Red line indicates the *in vivo* rate of epigenetic aging of 1 biological year per a chronological year. (**B**) Repeat experiments measuring the variability in the rate of epigenetic aging for Donor 3 (male, green) and 4 (female, purple). N = 3 repeat experiments. Horizontal bar represents the mean of the rates, noted with average speed up and coefficient of variation per group. (**C**) DNAmAge over cellular lifespan for each original clock (top-panel) and PC-based clock (bottom-panel). Horvath1 i.e. PantTissue clock, Horvath2 i.e. Skin&Blood clock. (**D**) Other DNAmAge clocks specialized for cultured cells. (**E**) Dunedin clocks used as blood biomarker for the pace of aging. (**F**) Telomere length estimates using qPCR-based T/S ratio (left-panel), DNAmTL (middle-panel), and PC-based DNAmTL (right-panel).

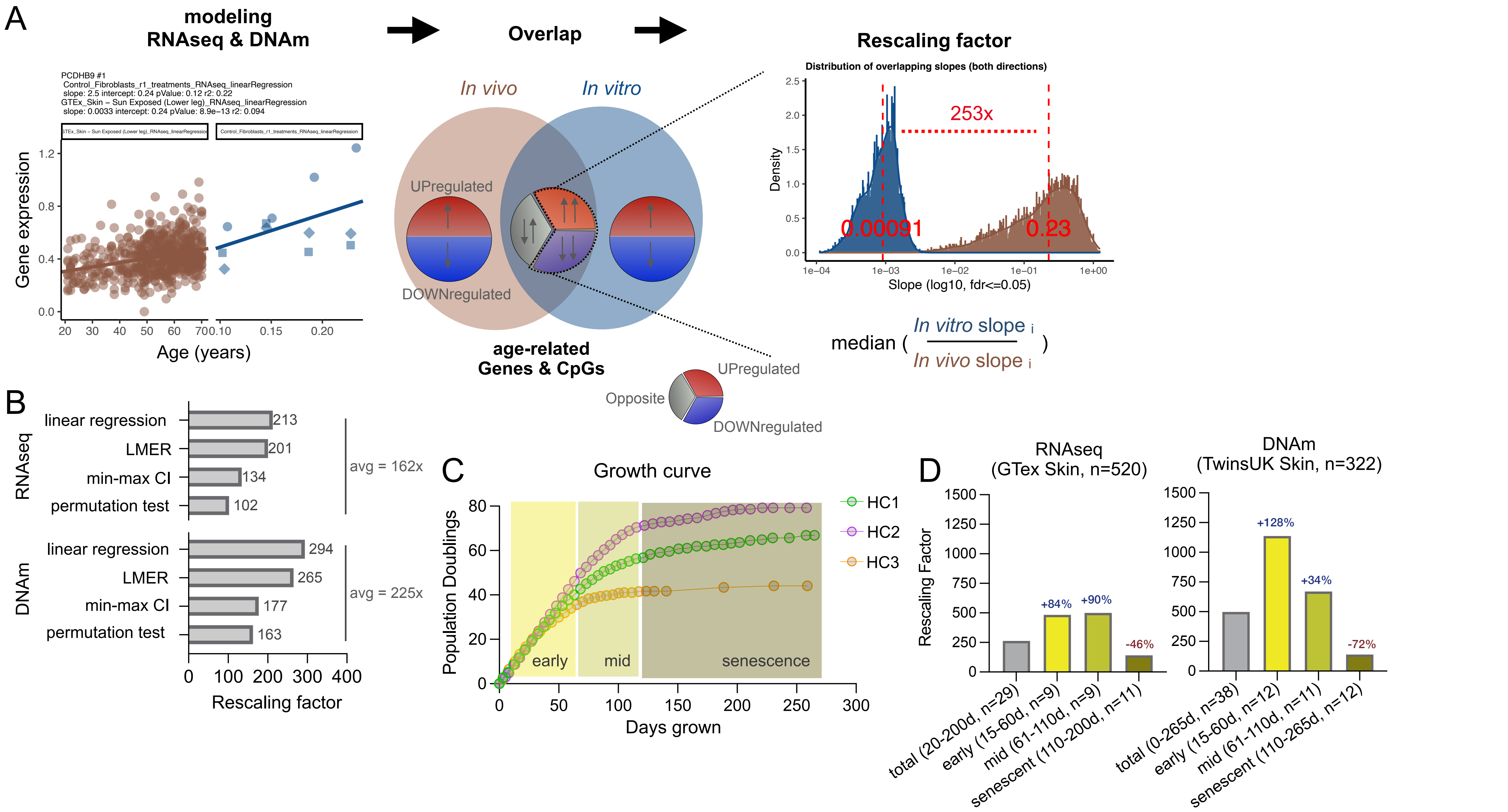


**Extended Data Figure 4. Derivation of methylation and transcription rescaling factor between *in vivo* to *in vitro* systems.** (**A**) Computational pipeline for deriving rescaling factors: Age-related gene expression and DNA methylation markers are selected from a linear modeling across lifespan in both *in vivo* and *in vitro* datasets. Genes and DNAm sites are than grouped by their direction of a change with age (UP or DOWN) and then the overlap of shared directional genes between the *in vivo* and *in vitro* datasets are selected. Finally, the rescaling factor is calculated as the ratio of the median *in vitro slopes* to the median of the *in vivo* slopes. (**B**) Rescaling factor based on different modeling techniques of the *in vivo* data. Note, all *in vitro* slopes were obtained using a linear model. (**C**) Growth curves of three untreated cell lines (Donor3-5). Color blocks indicate sections of lifespan operationalized as linear, mid, and senescent growth phases. (**D**) Comparison of rescaling factor depending on the portion of the cellular lifespan at which the slope was calculated. Genes and DNAm marker were selected from the same set of age-related molecules as identified by a full-lifespan nonlinear generalized additive modeling (GAM).

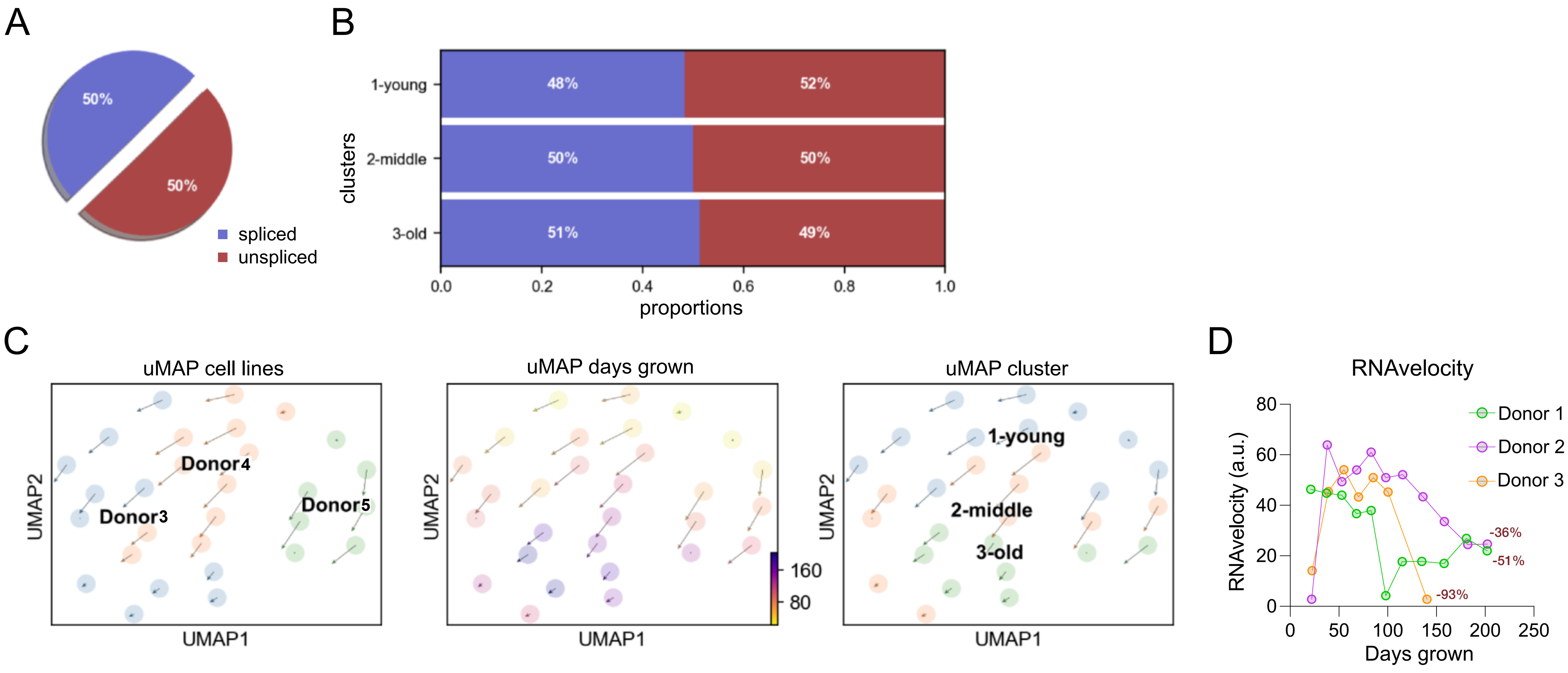


**Extended Data Figure 5. RNAvelocity across cellular lifespan.** (**A-D**) Rate of transcriptional events calculated as using ratios of splice : unspliced mRNA as outlined in La Manno et al. (REF. Nature, 2018) (**A**) Pi chart of number of splice to unspliced mRNA. (**B**) Splice ratio grouped by age of cells. Young = 0-60 days, middle = 61-110 days, old = 110-200 days. (**C**) Uniform manifold approximation and projection (uMAP) applied for dimension reduction of gene expression data shown by donor (left-panel), days grown (middle-panel) and clustered age group (right-panel). Length of arrows at each point indicate the RNAvelocity measured using stochastic modeling. (**D**) RNAvelocity rates across cellular lifespan confirming the slow-down of transcriptional events as cells age *in vitro.* Percent values are the final timepoint / average of the first three timepoints.

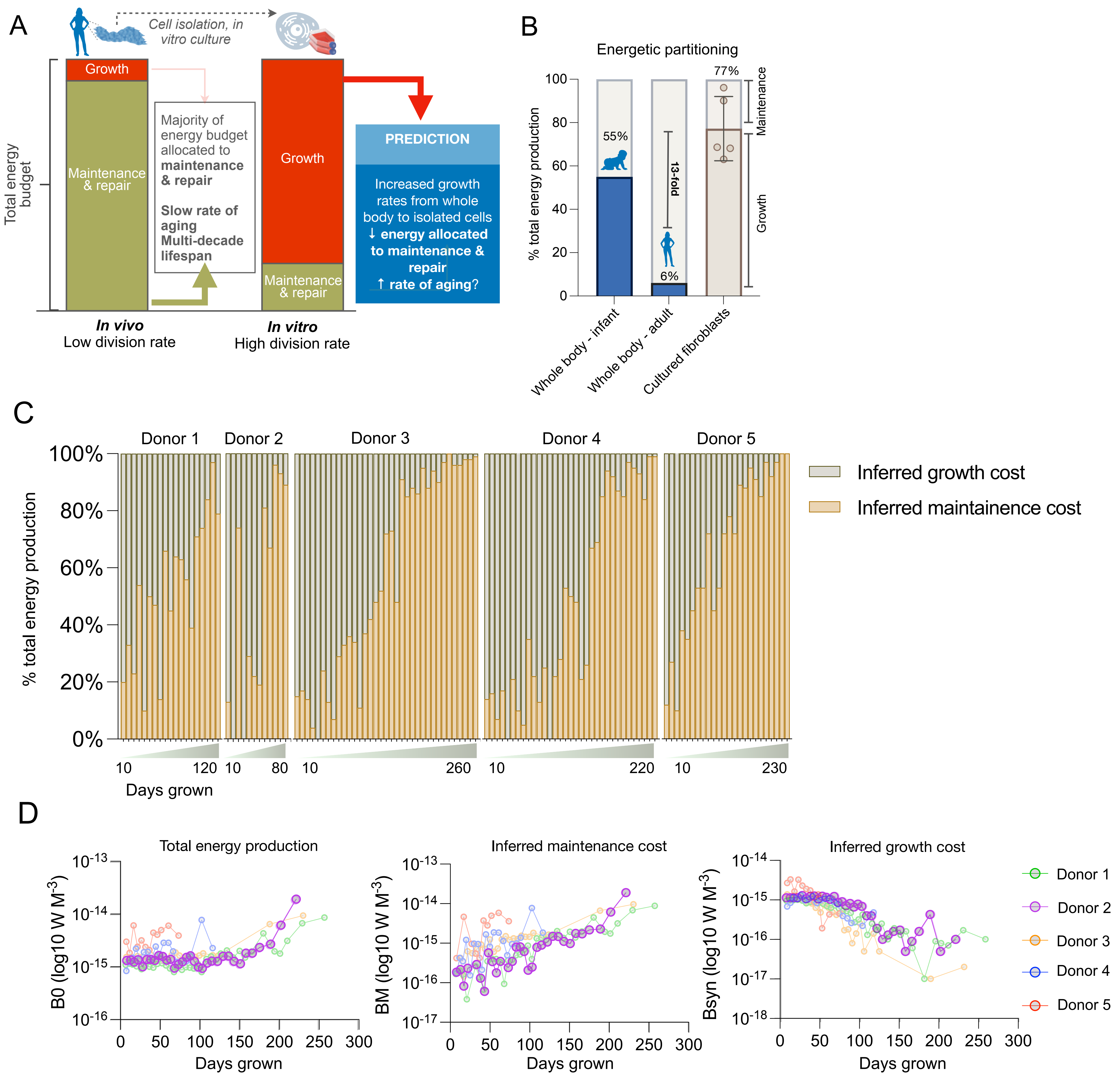


**Extended Data Figure 6. Replicative lifespan predicted by energetic, growth, and damage estimates.** (**A**) Illustrative framework for metabolic partitioning of cells isolated from human tissue and placed in culture. Upon isolation, cellular growth rate increases, and the amount of energy devoted to maintenance and repair processes decrease and consequently the rate of aging increases. (**B**) Empirical energetic partitioning of maintenance and growth costs in whole body infants and adults, and cultured fibroblasts. Cellular rates are quantified during the linear growth phase (days 10-70, 4-5 timepoints per cell line) . (**C**) Fractions of total energy production devoted to maintenance and growth costs across the cellular lifespan in culture. Note the reduction in growth-related costs as cell division rates approach zero at the end of the lifespan. All values are interpolated from a piecewise first-order model. (**D**) Inferred total energy production (B0, left-panel), maintenance costs (middle-panel), and growth costs (right-panel) across the cellular lifespan (see equation 1 for derivation).

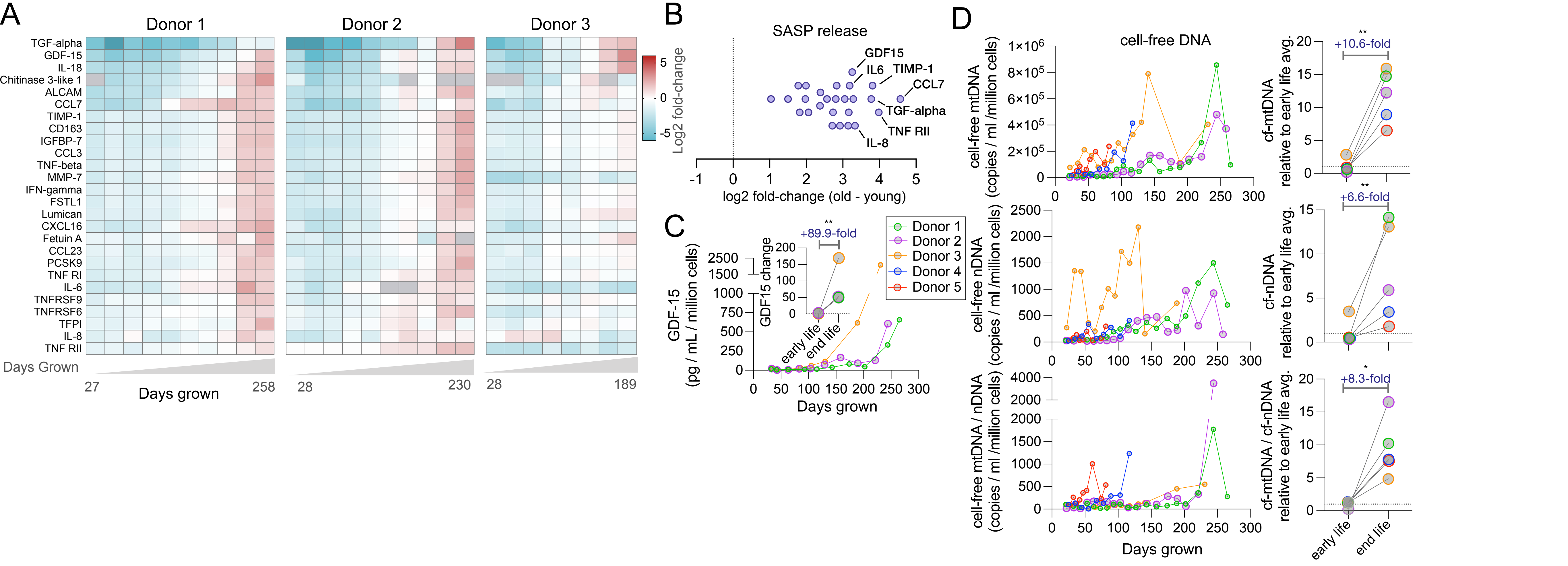


**Extended Data Figure 7. Age-related release of secretion factors.** (**A**) Extracellular cytokine concentrations released on a per-cell basis, measured on two multiplex arrays across the cellular lifespan of Donors 1-3. Values are $Log_2$ median-centered for each cytokine; samples with undetectable values are shown as grey cells. (**B**) Median $log_2$ fold changes in extracellular levels of each cytokine from young to old fibroblasts, showing up to >20-fold upregulation for all cytokines. (**C**) Extracellular GDF15 across the cellular lifespan for Donors 1-3, with (*inset*) fold change relative to the early life average. (**D**) Cell-free mtDNA (*top*), nDNA (*middle*) and mtDNA/nDNA ratio (*bottom*) across the cellular lifespan. (*Right*) Summary plots show fold change relative to early life average for Donors 1-5. Two-tailed paired ratio t-test, * $p < 0.05$, ** $p < 0.01$.

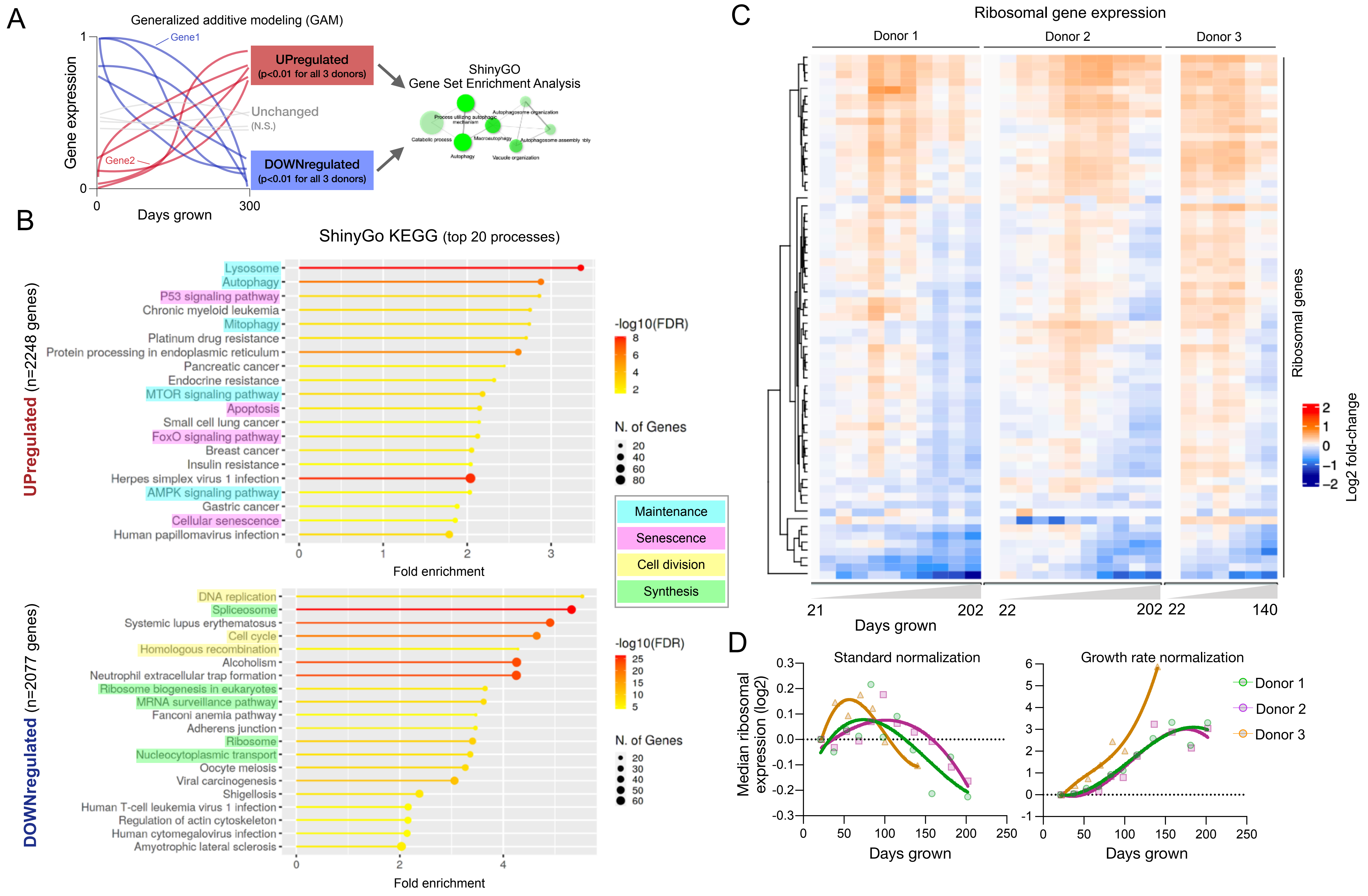


**Extended Data Figure 8. Age-related upregulation in maintenance and synthesis pathways.** (**A**) Analytic strategy to identify overrepresented biological processes from RNAseq data where age-related genes (significant nonlinear change across the lifespan, generalized additive modeling, n=3 Donors, 7-11 timepoints/cell line) are grouped by the direction of change and analyzed in ShinyGo (v0.75). (**B**) Top 20 KEGG processes for both upregulated and downregulated genes. Color indicates fold enrichment, dot size indicates the number of genes enriched in each process. Highlighted processes are categorized by shared biological function. (**C**) Lifespan trajectories of ribosomal subunit gene expression (RNAseq) centered to the median of the youngest control timepoints. (**D**) Median expression of all ribosomal subunits using standard normalized expression (*left*, data fitted with third order polynomials) and adjusted for population doubling rate (i.e., growth rate in cell divisions per day, *right*), highlighting the disproportionate expression of biosynthesis machinery relative to growth rates.

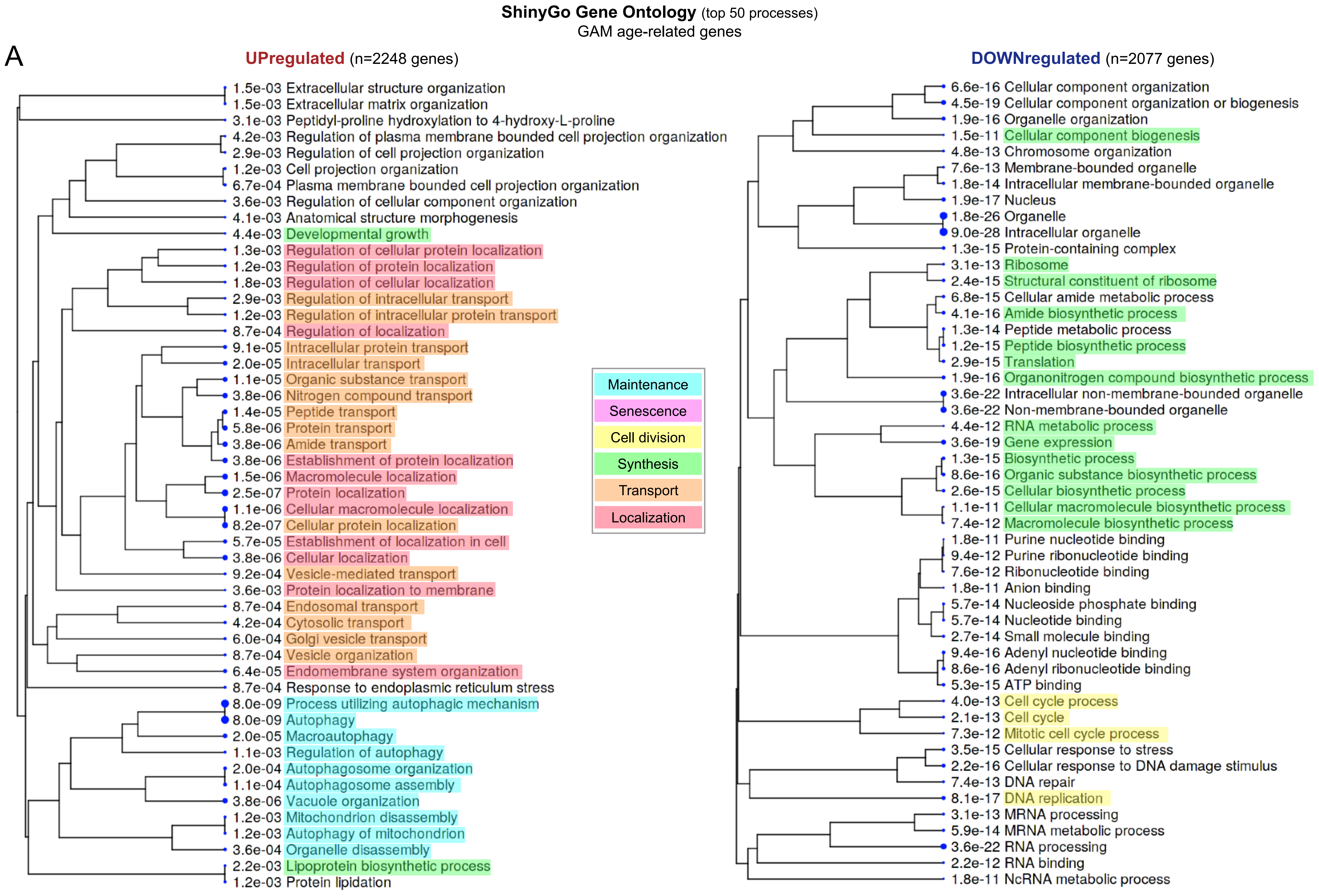


**Extended Data Figure 9. Gene set enrichment of age-related gene expression.** (**A**) Hierarchal clustering of enriched Gene Ontology pathways of age-related genes identified with generalized additive modeling. Highlighted processes are categorized by shared biological function. Only top 50 processes shown (see Supplemental Files 6 & 7 for full list).

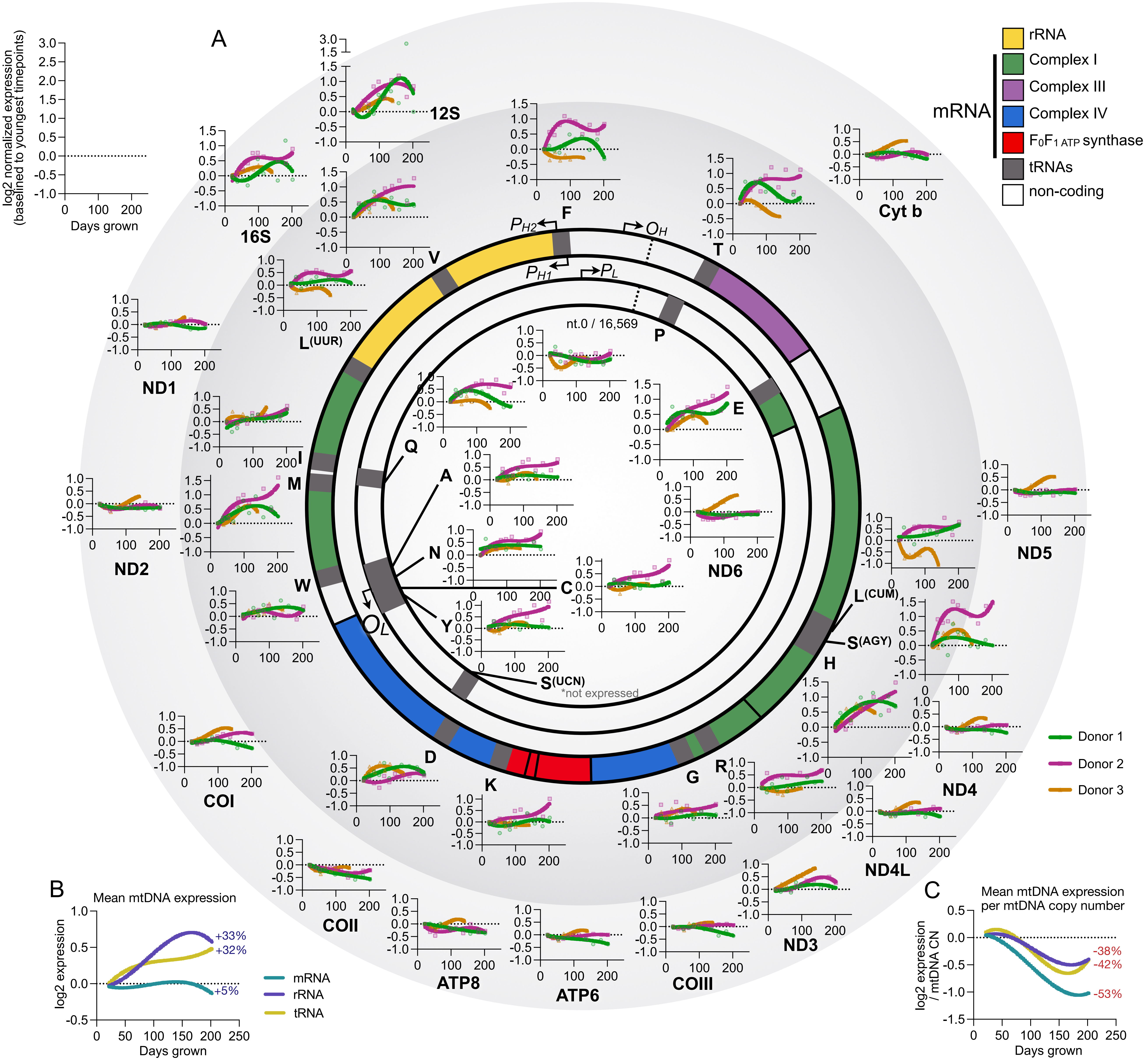

**Extended Data Figure 10. mtDNA gene expression of coding genes and tRNAs.** Circular mtDNA map with its 37 genes annotated. Graphs show normalized expression values (log2 fold-change relative to median of youngest timepoints) for 3 healthy donors across the cellular lifespan. The inner ring induces all transfer RNAs (tRNAs); the outer ring includes ribosomal and messenger RNA (rRNA and mRNA) genes. Graph height is scaled to the min/max of a given RNA. (**B**) Mean RNA species expression across the cellular lifespan. Values are the mean of log2 normalized expression baselined to youngest timepoints for the different RNA species (mRNA n=13, rRNA n=2, tRNA=22 genes). (**B**) Mean RNA species expression normalized to mtDNA copy number across the cellular lifespan. Copy number values are interpolated to match expression timepoints using generalized additive modeling (k=6).

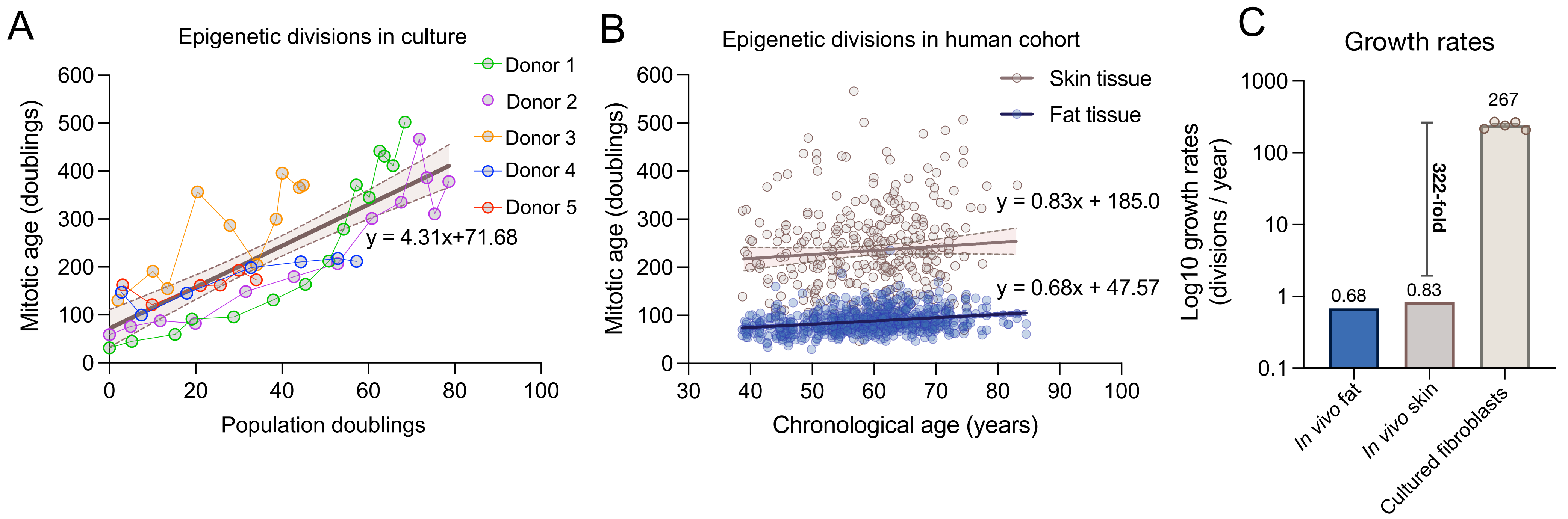


**Extended Data Figure 11. Growth rates of human fibroblasts.** (**A**) Epigenetic-derived cellular divisions (Mitotic age) as a function of empirically measured division in cultured fibroblasts (n=51 timepoints across 5 donor lines). Fit line represents linear regression with 95% confidence intervals across all timepoints. (**B**) Mitotic age measured in fat (n=648) and skin (n=322) tissue samples across a population of aging female twins (TwinsUK study). Fit line represents linear regression with 95% confidence intervals for each tissue. (**C**) Derived growth rate for *in vivo* fat and skin tissue compared to cultured fibroblasts. Rates are expressed as log 10 cellular divisions per year.

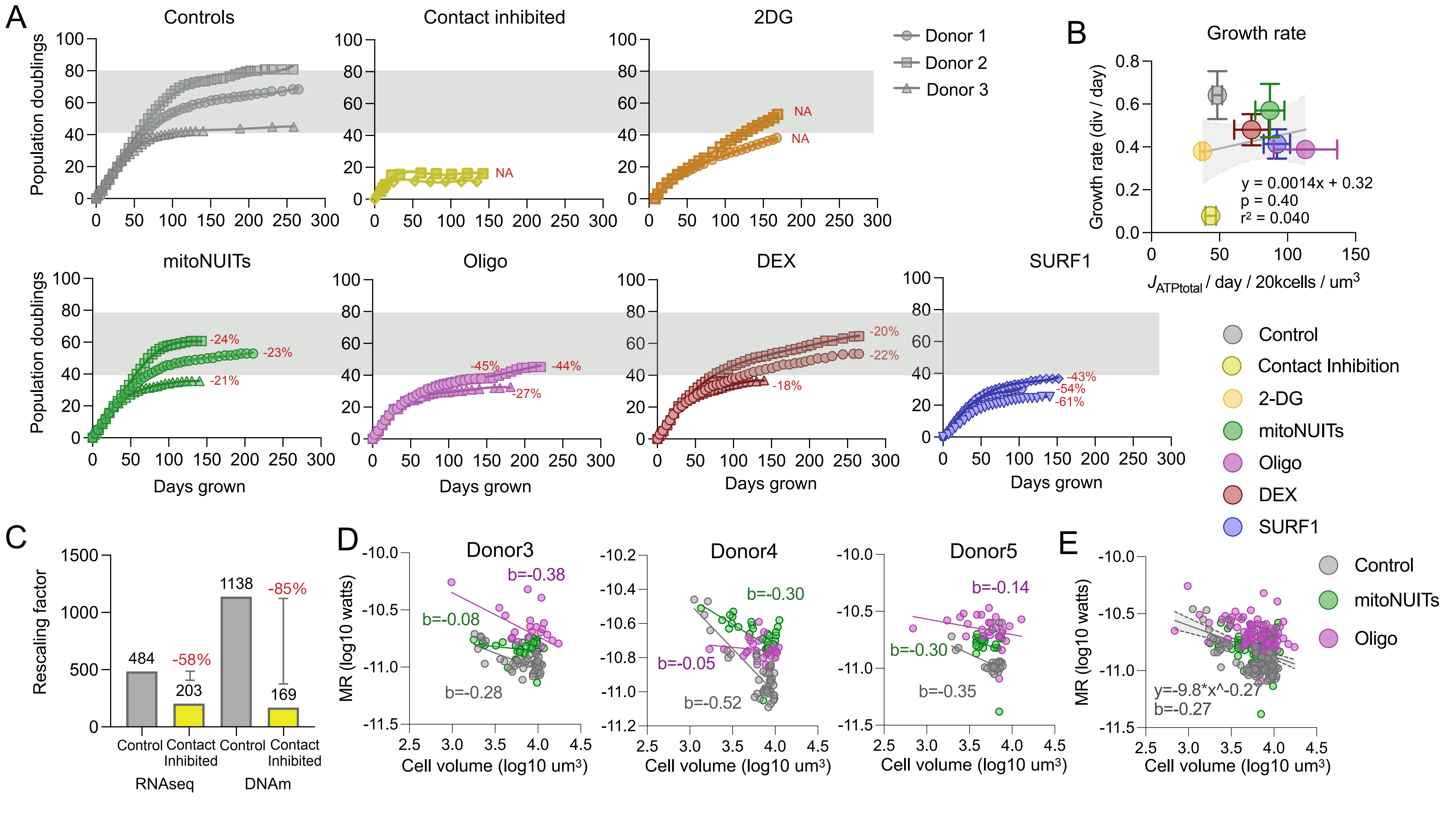


**Extended Data Figure 12. Alteration of metabolic rate with pharmacological, genetic, and environmental perturbations.** (**A**) Growth curves of each perturbation. Gray bar indicated range of the hayflick limit for untreated controls. 2-3 donors per condition. 2DG had limited growth curve due to experimental artifact. (**B**) Growth rate for each perturbation correlated with MR per volume. Growth rates are calculated between 15-60 days of growth. Gray line indicates linear regression with faded gray region indicated range of error. (**C**) Rescaling factor deceleration from contact inhibition for RNAseq and DNA methylation data. (**D-E**) Allometric scaling of cell volume to metabolic rate (MR) for selected treatment groups (controls, mitoNUITs, oligo) with time matched sampling. Treatments were selected to ensure matching timepoints across lifespan. Scaling coefficient for each cell line (D) and as for all aggregated data (E). B value indicates metabolic scaling coefficient which is -0.25 across species.

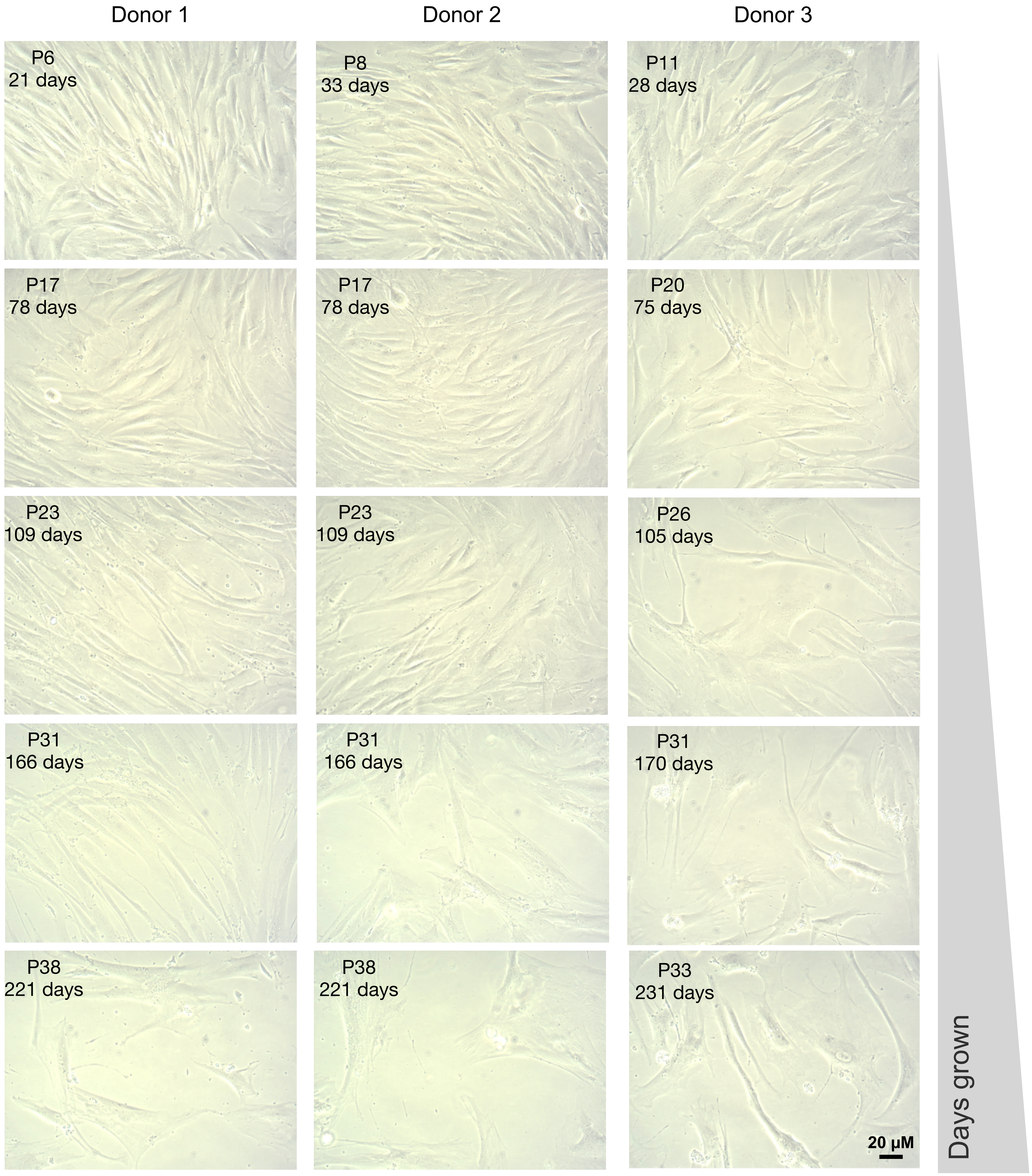


**Extended Data Figure 13. Bright-field microscopy images of fibroblasts across replicative lifespan.** Phase contrast bright field images of cultured fibroblasts across cellular lifespan. Images were taken with a 20k objective after 5-7 days of growth.
