## Supplemental files 1-8 for "Accelerating the clock: Interconnected speedup of energetic and molecular dynamics during aging in cultured human cells": SF8_Heatmaps_Aging_Controls_RNAseq.pdf

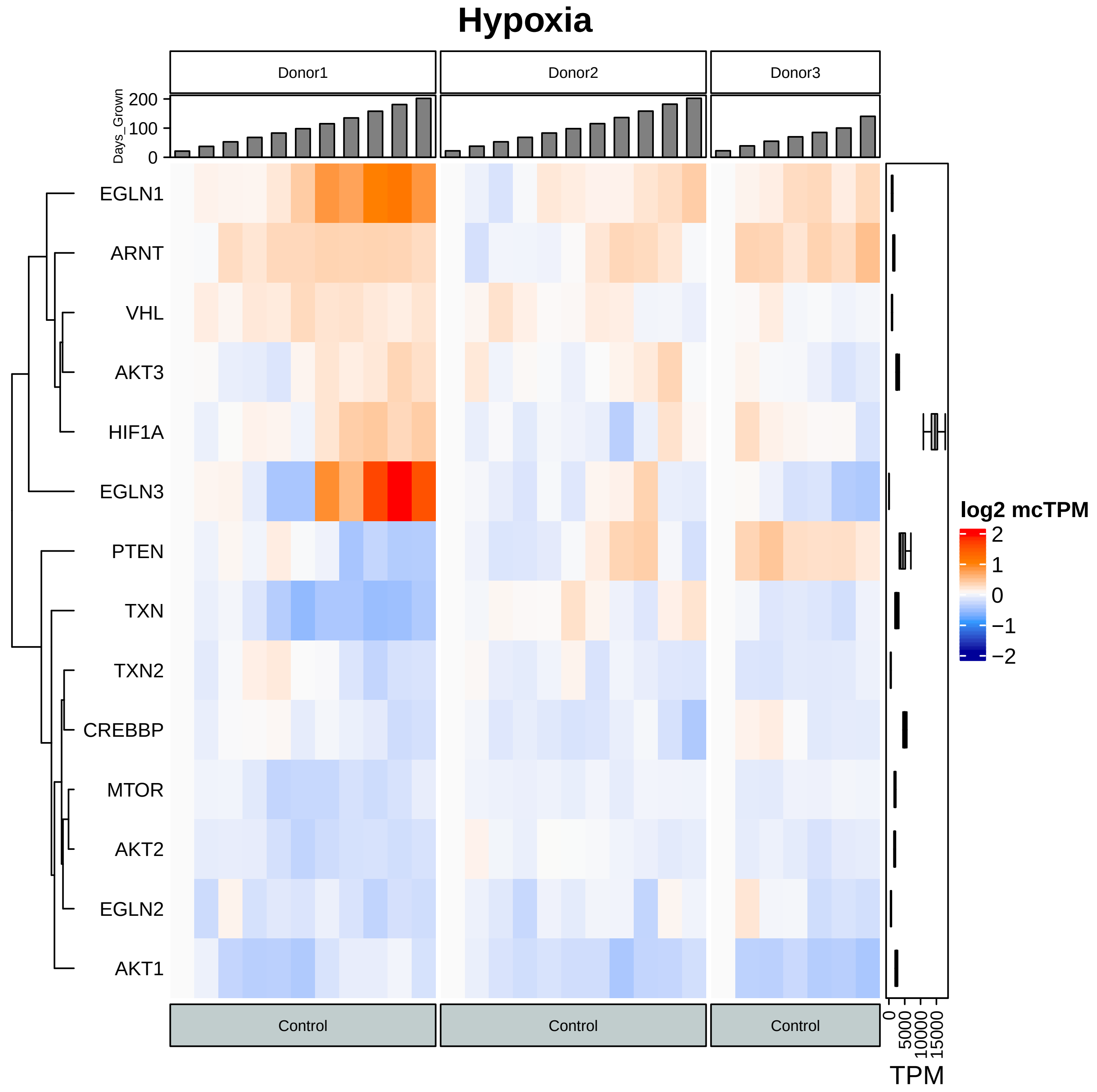

Hypoxia
Donor1
Donor2
Donor3
Days_Grown
200
100
0
EGLN1
ARNT
VHL
AKT3
HIF1A
EGLN3
PTEN
TXN
TXN2
CREBBP
MTOR
AKT2
EGLN2
AKT1
log2 mcTPM
2
1
0
−1
−2
Control
Control
Control
0
5000
10000
15000
TPM

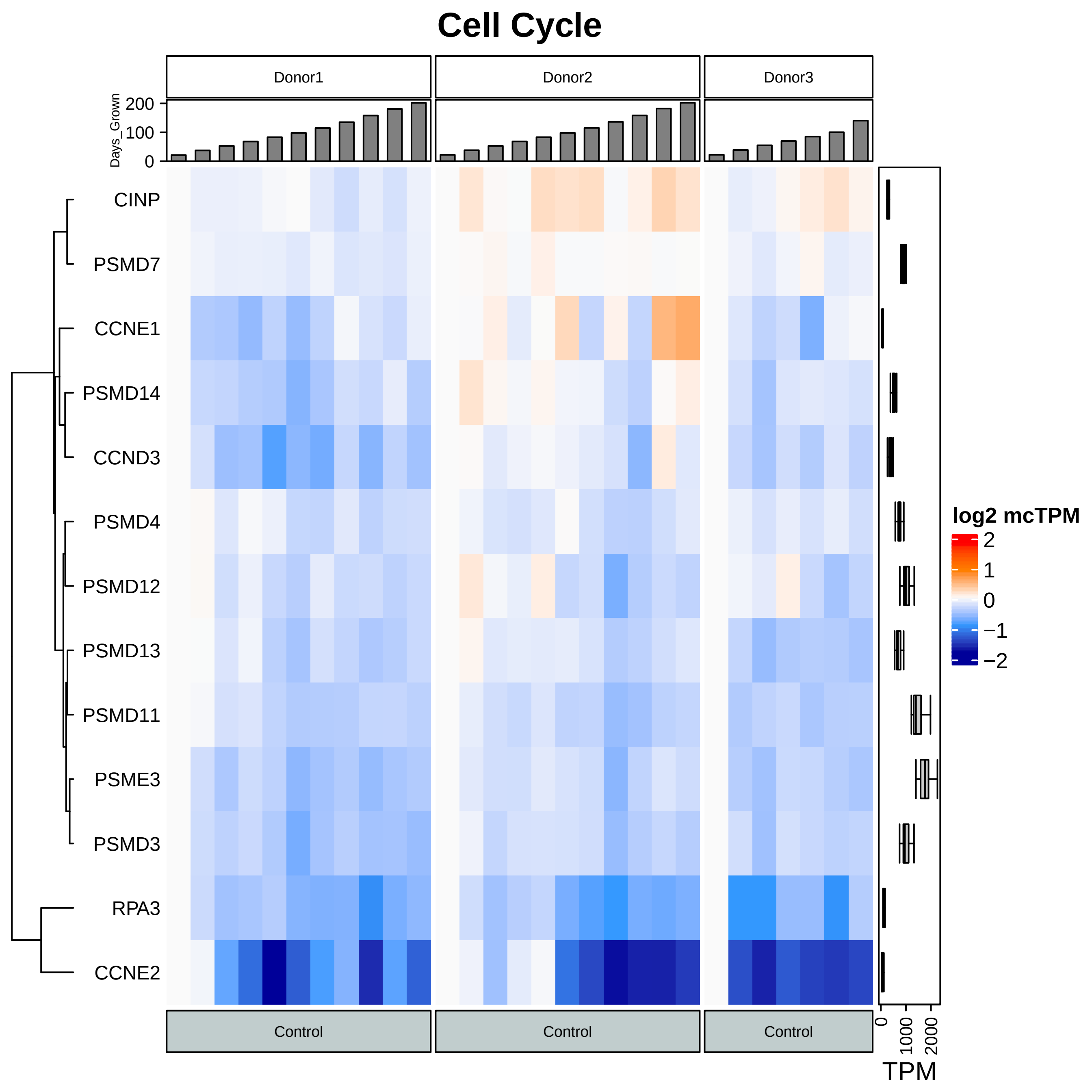
Cell Cycle
Donor1
Donor2
Donor3
Days_Grown
200
100
0
CINP
PSMD7
CCNE1
PSMD14
CCND3
PSMD4
PSMD12
PSMD13
PSMD11
PSME3
PSMD3
RPA3
CCNE2
log2 mcTPM
2
1
0
−1
−2
Control
Control
Control
0
1000
2000
TPM

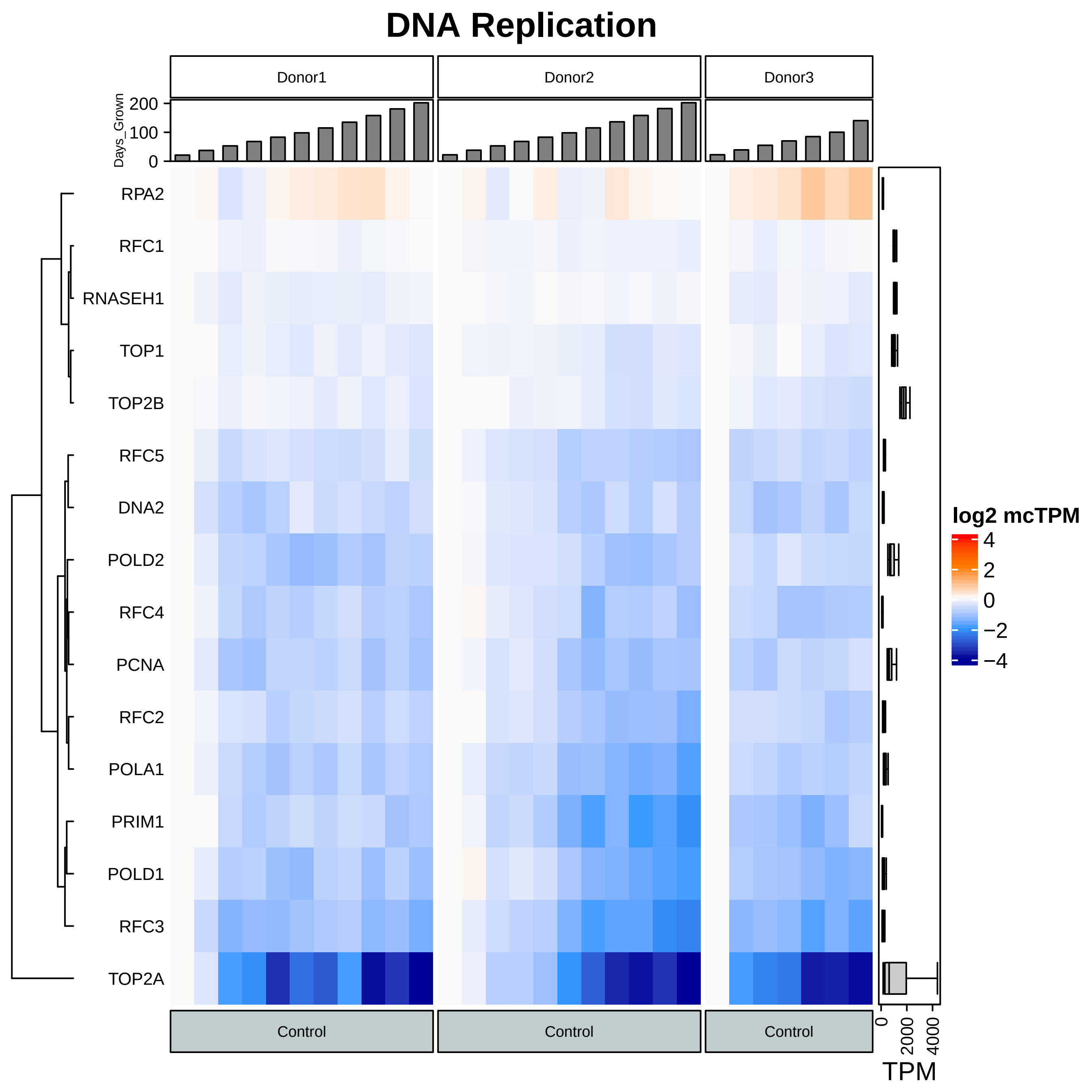
DNA Replication
Donor1
Donor2
Donor3
Days_Grown
200
100
0
RPA2
RFC1
RNASEH1
TOP1
TOP2B
RFC5
DNA2
POLD2
RFC4
PCNA
RFC2
POLA1
PRIM1
POLD1
RFC3
TOP2A
log2 mcTPM
4
2
0
−2
−4
Control
Control
Control
0
2000
4000
TPM

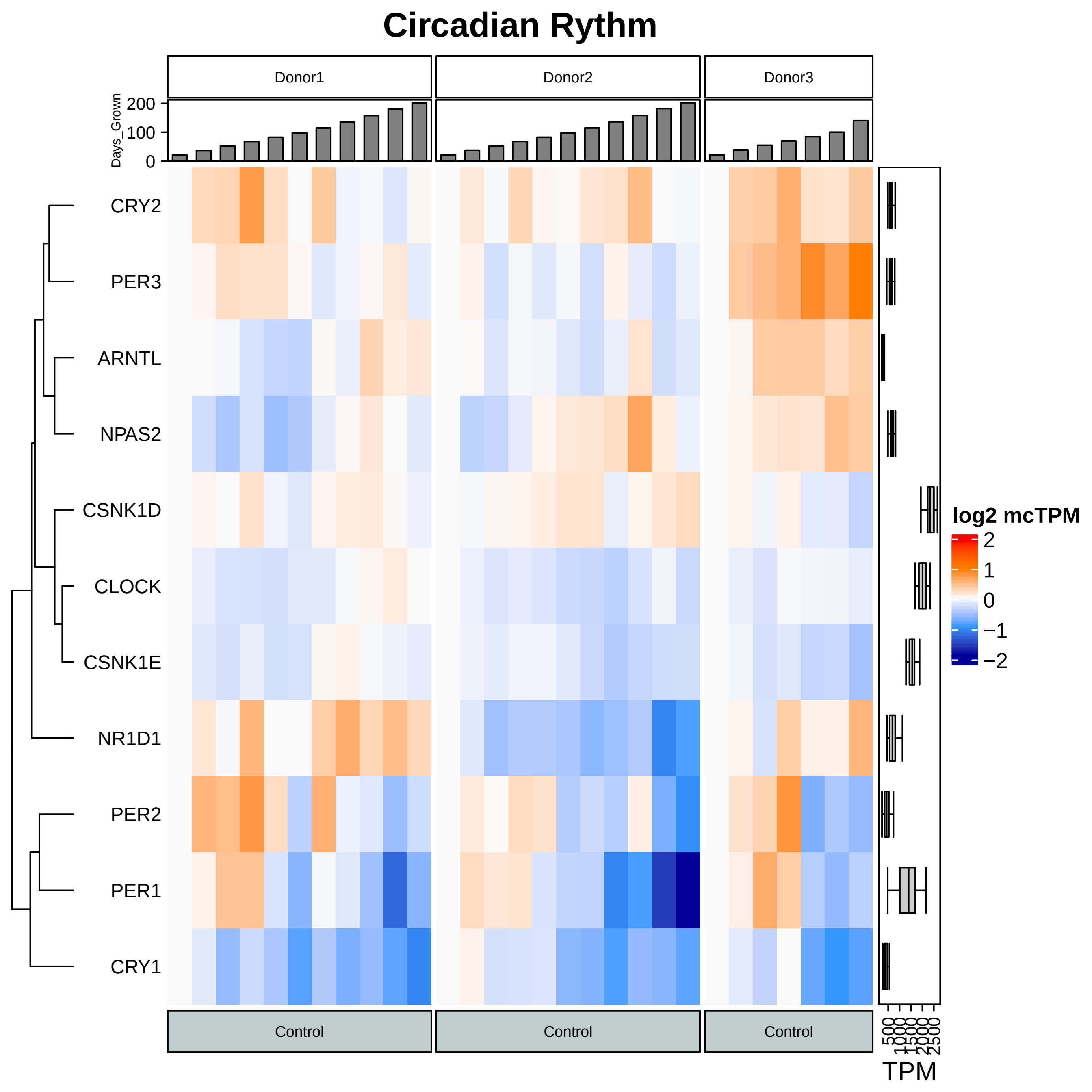
Circadian Rythm
Donor1
Donor2
Donor3
Days_Grown
200
100
0
CRY2
PER3
ARNTL
NPAS2
CSNK1D
CLOCK
CSNK1E
NR1D1
PER2
PER1
CRY1
log2 mcTPM
2
1
0
−1
−2
Control
Control
Control
500
1000
1500
2000
2500
TPM

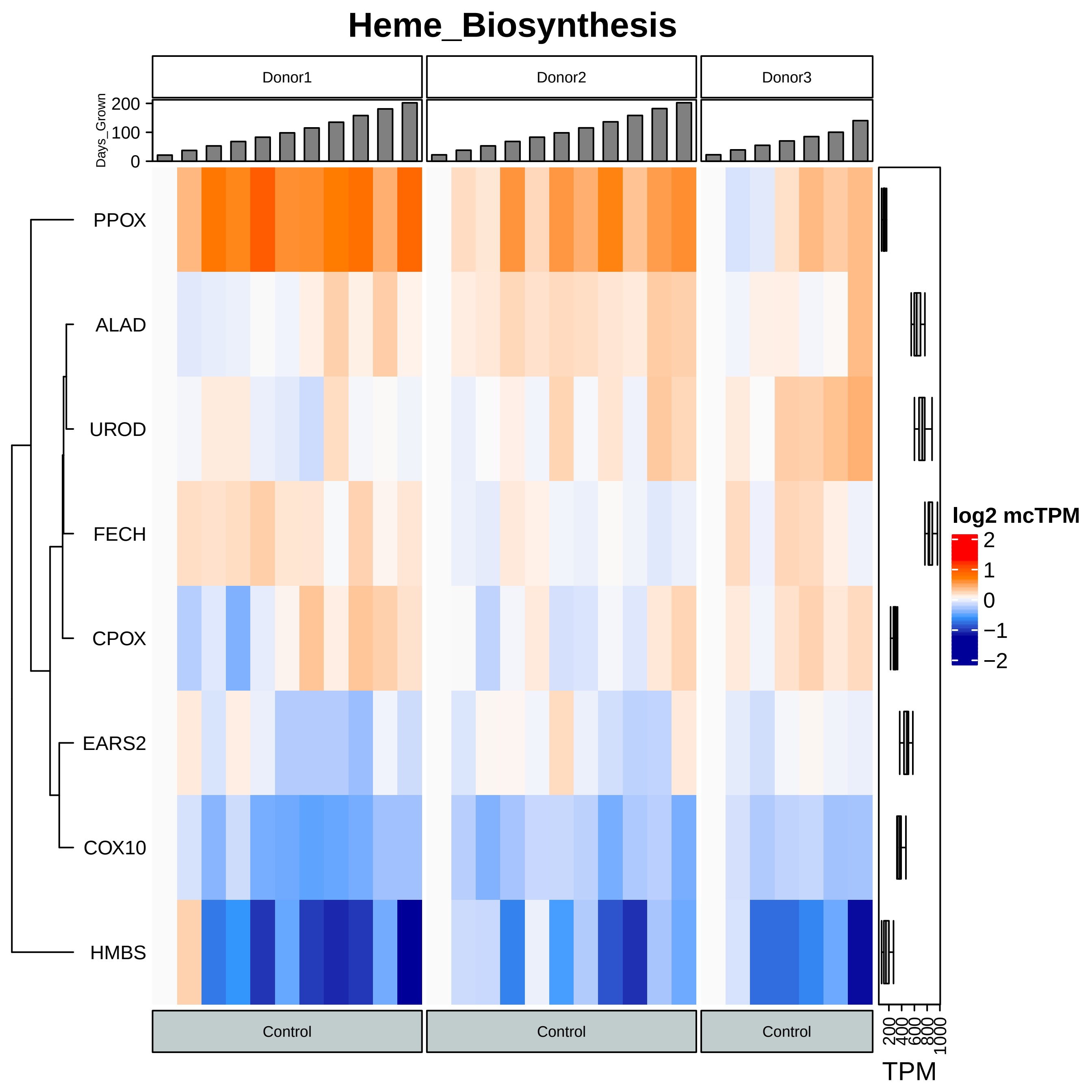
Heme_Biosynthesis
Donor1
Donor2
Donor3
Days_Grown
200
100
0
PPOX
ALAD
UROD
FECH
CPOX
EARS2
COX10
HMBS
log2 mcTPM
2
1
0
−1
−2
Control
Control
Control
200
400
600
800
1000
TPM

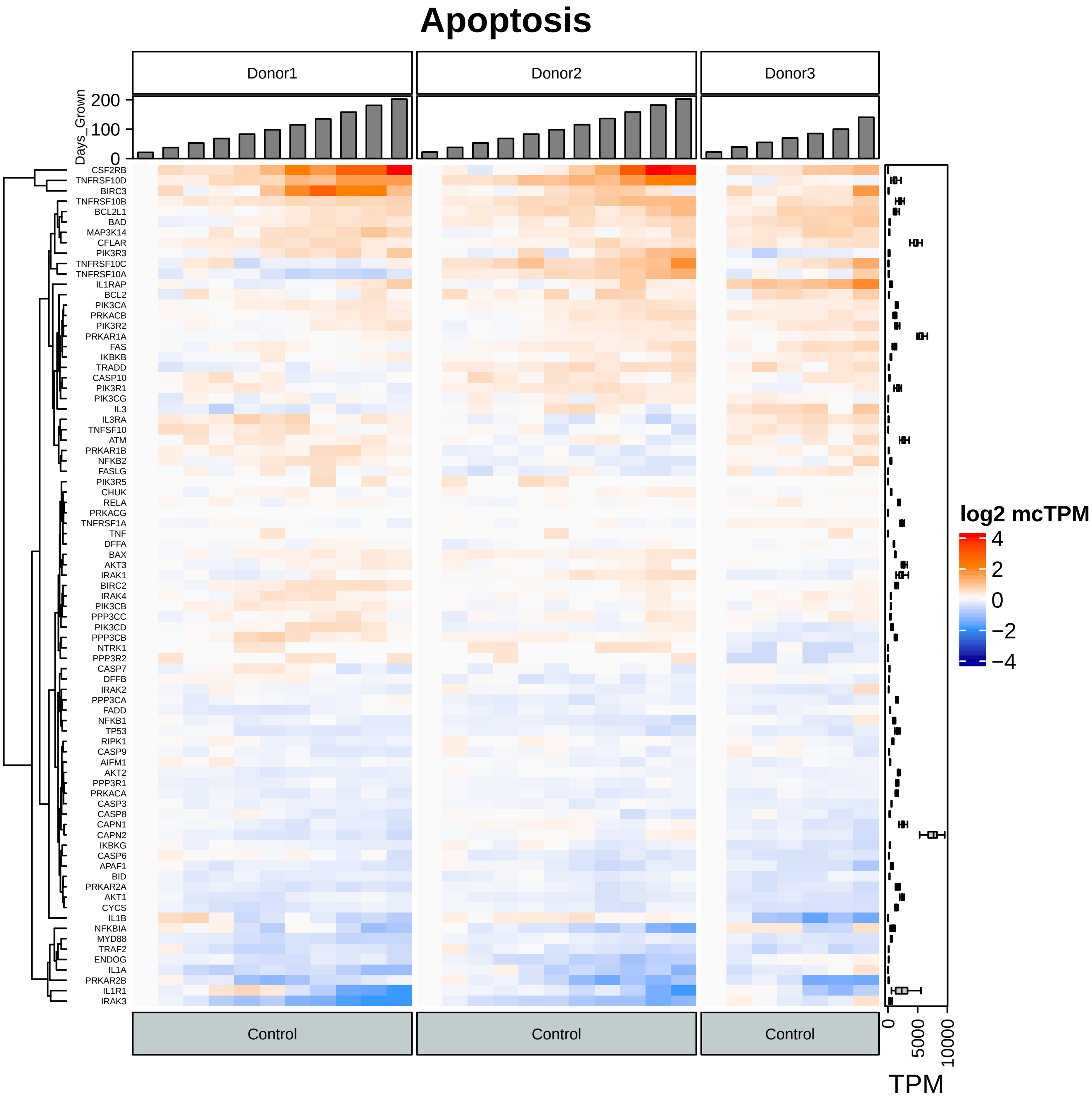
Apoptosis
Donor1
Donor2
Donor3
Days_Grown
200
100
0
CSF2RB
TNFRSF10D
BIRC3
TNFRSF10B
BCL2L1
BAD
MAP3K14
CFLAR
PIK3R3
TNFRSF10C
TNFRSF10A
IL1RAP
BCL2
PIK3CA
PRKACB
PIK3R2
PRKAR1A
FAS
IKBKB
TRADD
CASP10
PIK3R1
PIK3CG
IL3
IL3RA
TNFSF10
ATM
PRKAR1B
NFKB2
FASLG
PIK3R5
CHUK
RELA
PRKACG
TNFRSF1A
TNF
DFFA
BAX
AKT3
IRAK1
BIRC2
IRAK4
PIK3CB
PPP3CC
PIK3CD
PPP3CB
NTRK1
PPP3R2
CASP7
DFFB
IRAK2
PPP3CA
FADD
NFKB1
TP53
RIPK1
CASP9
AIFM1
AKT2
PPP3R1
PRKACA
CASP3
CASP8
CAPN1
CAPN2
IKBKG
CASP6
APAF1
BID
PRKAR2A
AKT1
CYCS
IL1B
NFKBIA
MYD88
TRAF2
ENDOG
IL1A
PRKAR2B
IL1R1
IRAK3
log2 mcTPM
4
2
0
−2
−4
Control
Control
Control
0
5000
10000
TPM

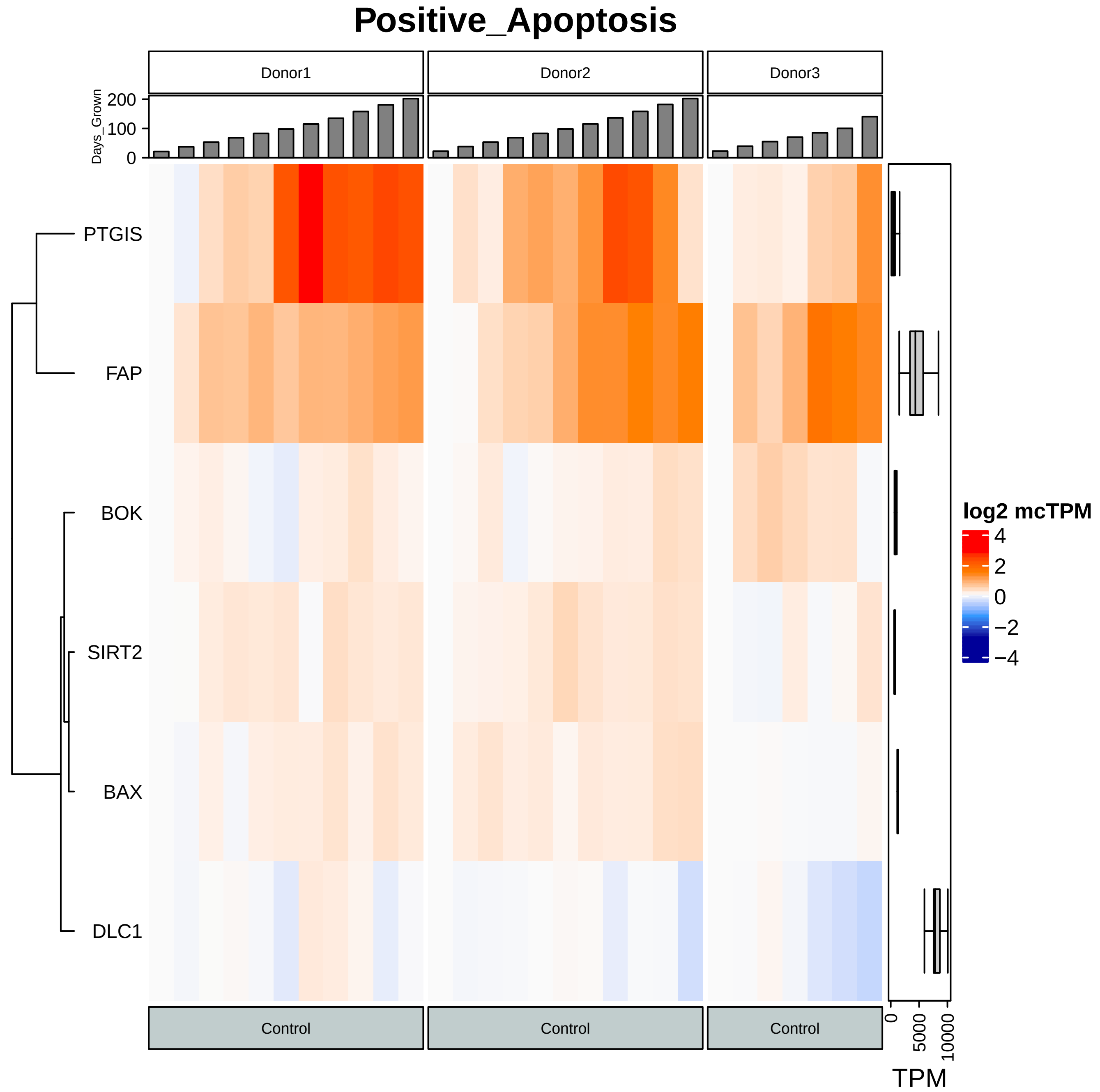

Positive_Apoptosis
Donor1
Donor2
Donor3
Days_Grown
200
100
0
PTGIS
FAP
BOK
SIRT2
BAX
DLC1
log2 mcTPM
4
2
0
−2
−4
Control
Control
Control
0
5000
10000
TPM

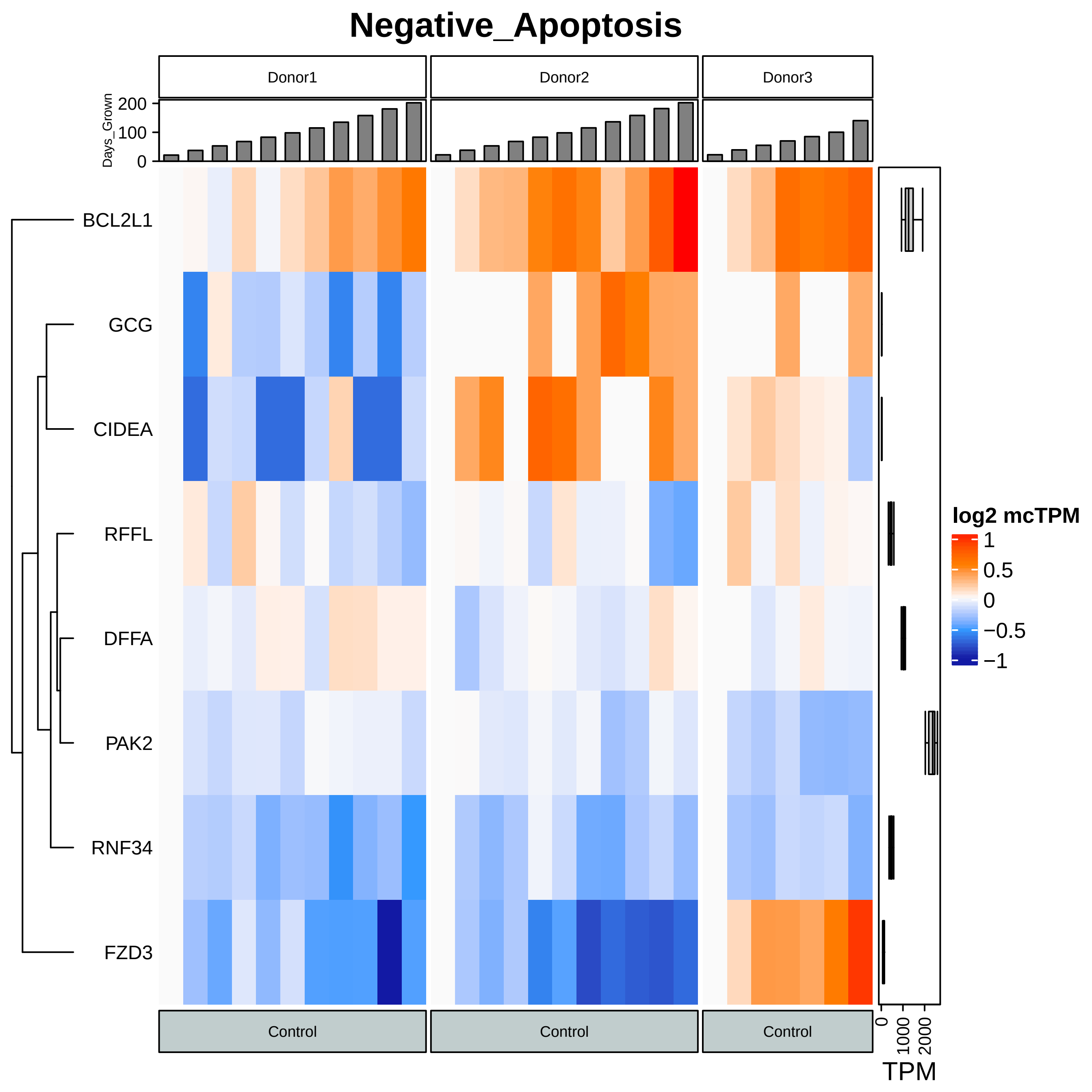

Negative_Apoptosis
Donor1
Donor2
Donor3
Days_Grown
0
100
200
BCL2L1
GCG
CIDEA
RFFL
DFFA
PAK2
RNF34
FZD3
Control
Control
Control
log2 mcTPM
1
0.5
0
−0.5
−1
0
1000
2000
TPM

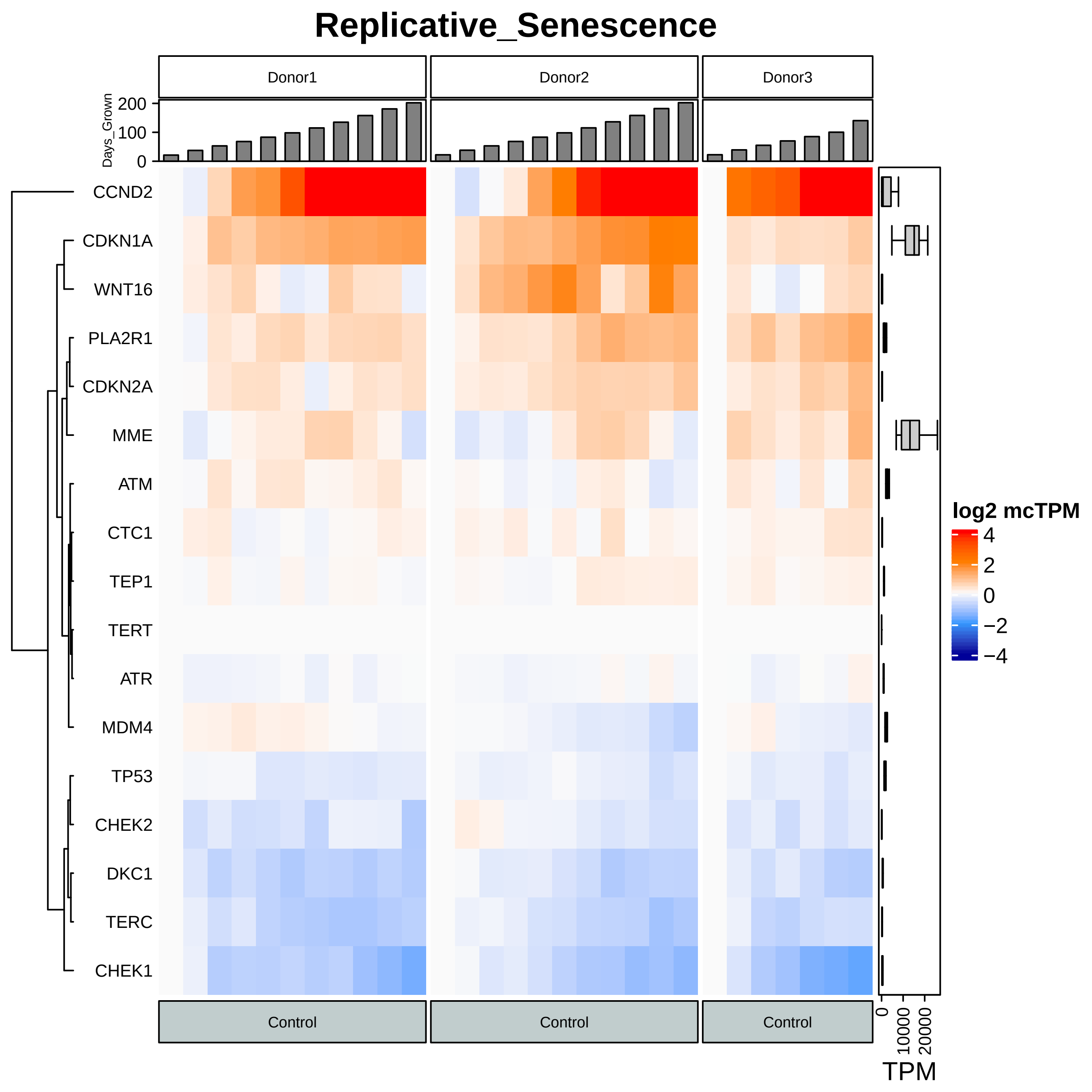
Replicative_Senescence
Donor1
Donor2
Donor3
Days_Grown
200
100
0
CCND2
CDKN1A
WNT16
PLA2R1
CDKN2A
MME
ATM
CTC1
TEP1
TERT
ATR
MDM4
TP53
CHEK2
DKC1
TERC
CHEK1
Control
Control
Control
log2 mcTPM
4
2
0
−2
−4
0
10000
20000
TPM

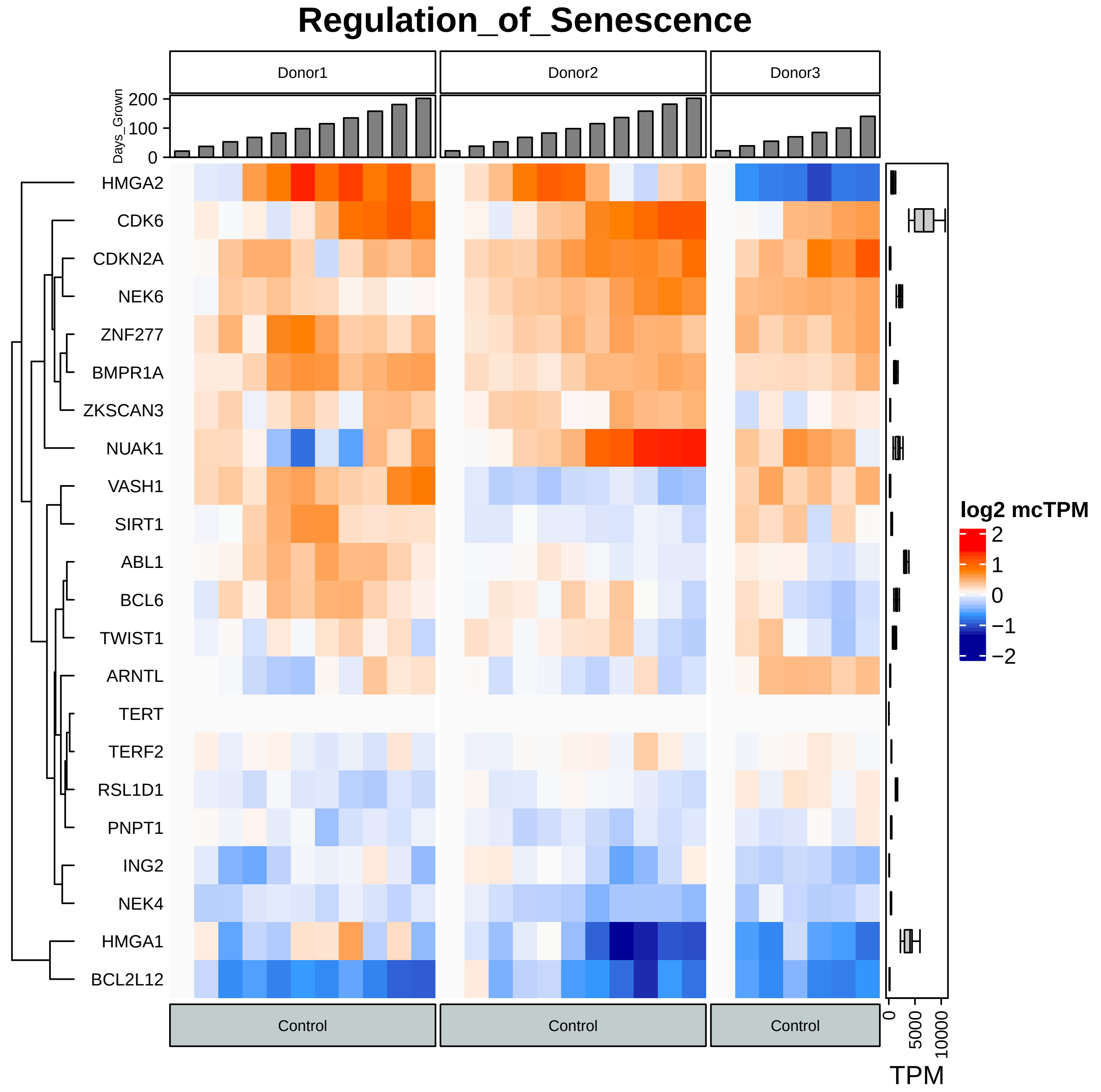

Regulation_of_Senescence
Donor1
Donor2
Donor3
Days_Grown
200
100
0
HMGA2
CDK6
CDKN2A
NEK6
ZNF277
BMPR1A
ZKSCAN3
NUAK1
VASH1
SIRT1
ABL1
BCL6
TWIST1
ARNTL
TERT
TERF2
RSL1D1
PNPT1
ING2
NEK4
HMGA1
BCL2L12
Control
Control
Control
log2 mcTPM
2
1
0
−1
−2
0
5000
10000
TPM

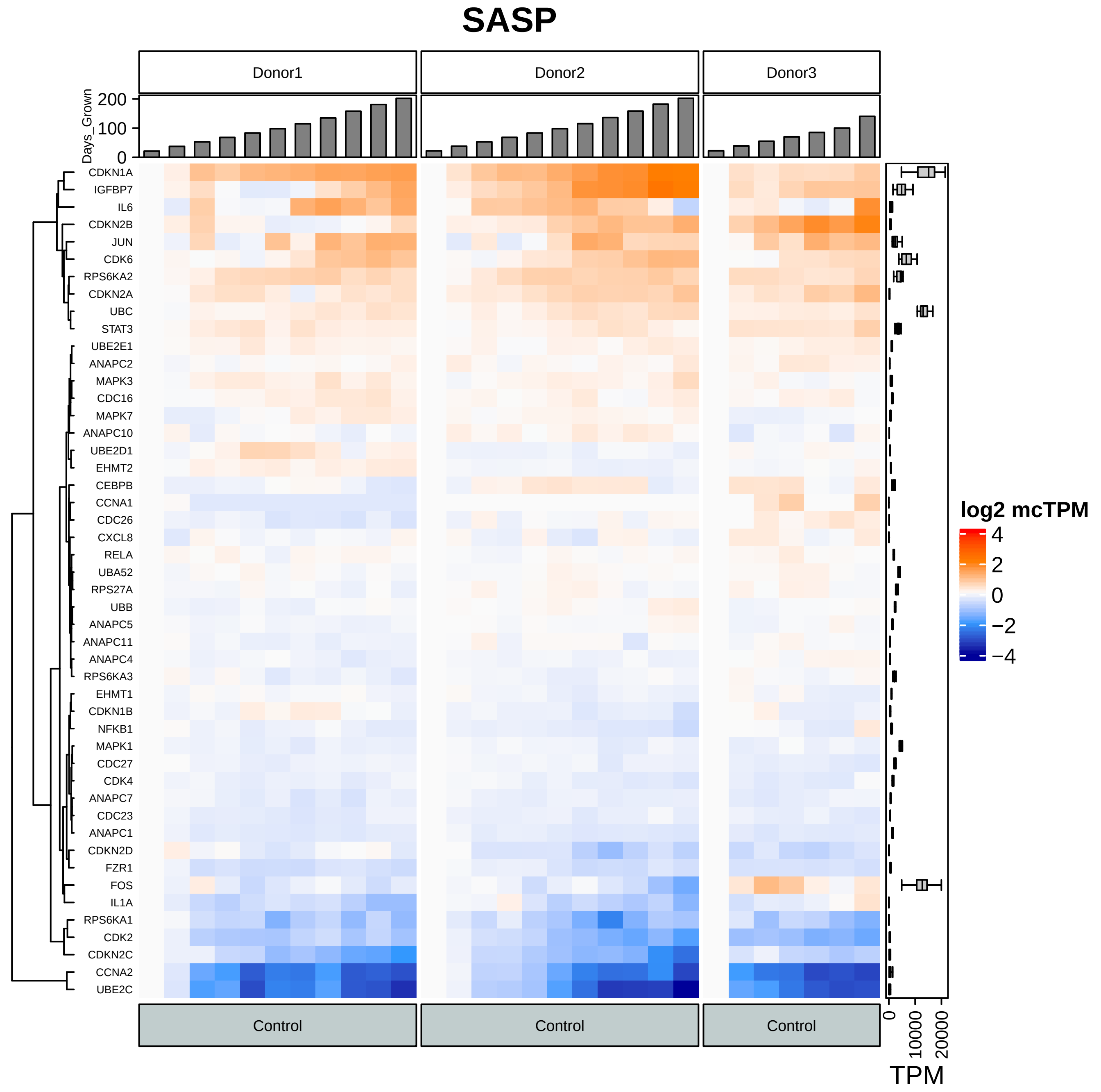
SASP
Donor1
Donor2
Donor3
Days_Grown
200
100
0
CDKN1A
IGFBP7
IL6
CDKN2B
JUN
CDK6
RPS6KA2
CDKN2A
UBC
STAT3
UBE2E1
ANAPC2
MAPK3
CDC16
MAPK7
ANAPC10
UBE2D1
EHMT2
CEBPB
CCNA1
CDC26
CXCL8
RELA
UBA52
RPS27A
UBB
ANAPC5
ANAPC11
ANAPC4
RPS6KA3
EHMT1
CDKN1B
NFKB1
MAPK1
CDC27
CDK4
ANAPC7
CDC23
ANAPC1
CDKN2D
FZR1
FOS
IL1A
RPS6KA1
CDK2
CDKN2C
CCNA2
UBE2C
Control
Control
Control
log2 mcTPM
4
2
0
−2
−4
0
10000
20000
TPM

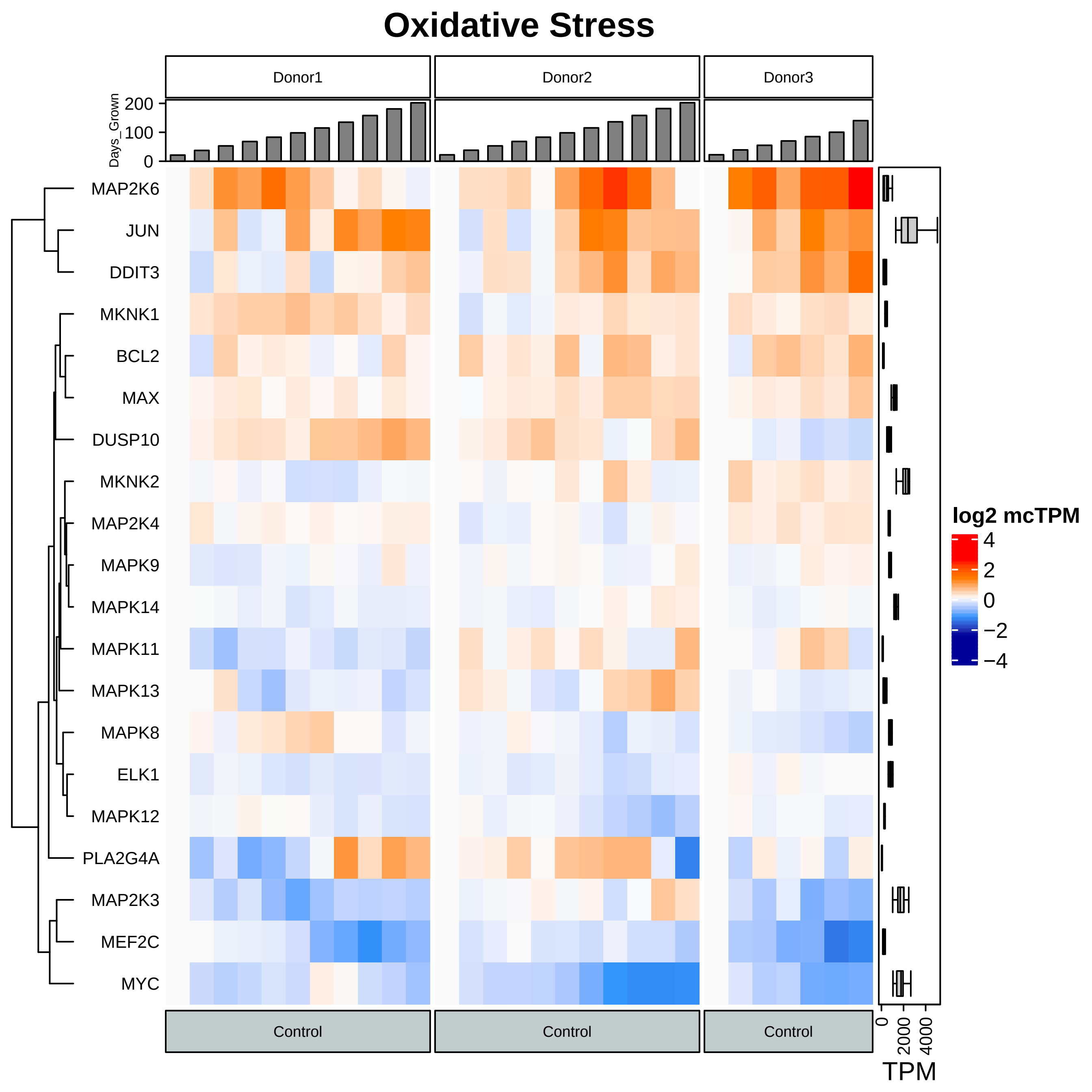

Oxidative Stress
Donor1
Donor2
Donor3
Days_Grown
200
100
0
MAP2K6
JUN
DDIT3
MKNK1
BCL2
MAX
DUSP10
MKNK2
MAP2K4
MAPK9
MAPK14
MAPK11
MAPK13
MAPK8
ELK1
MAPK12
PLA2G4A
MAP2K3
MEF2C
MYC
log2 mcTPM
4
2
0
−2
−4
Control
Control
Control
0
2000
4000
TPM

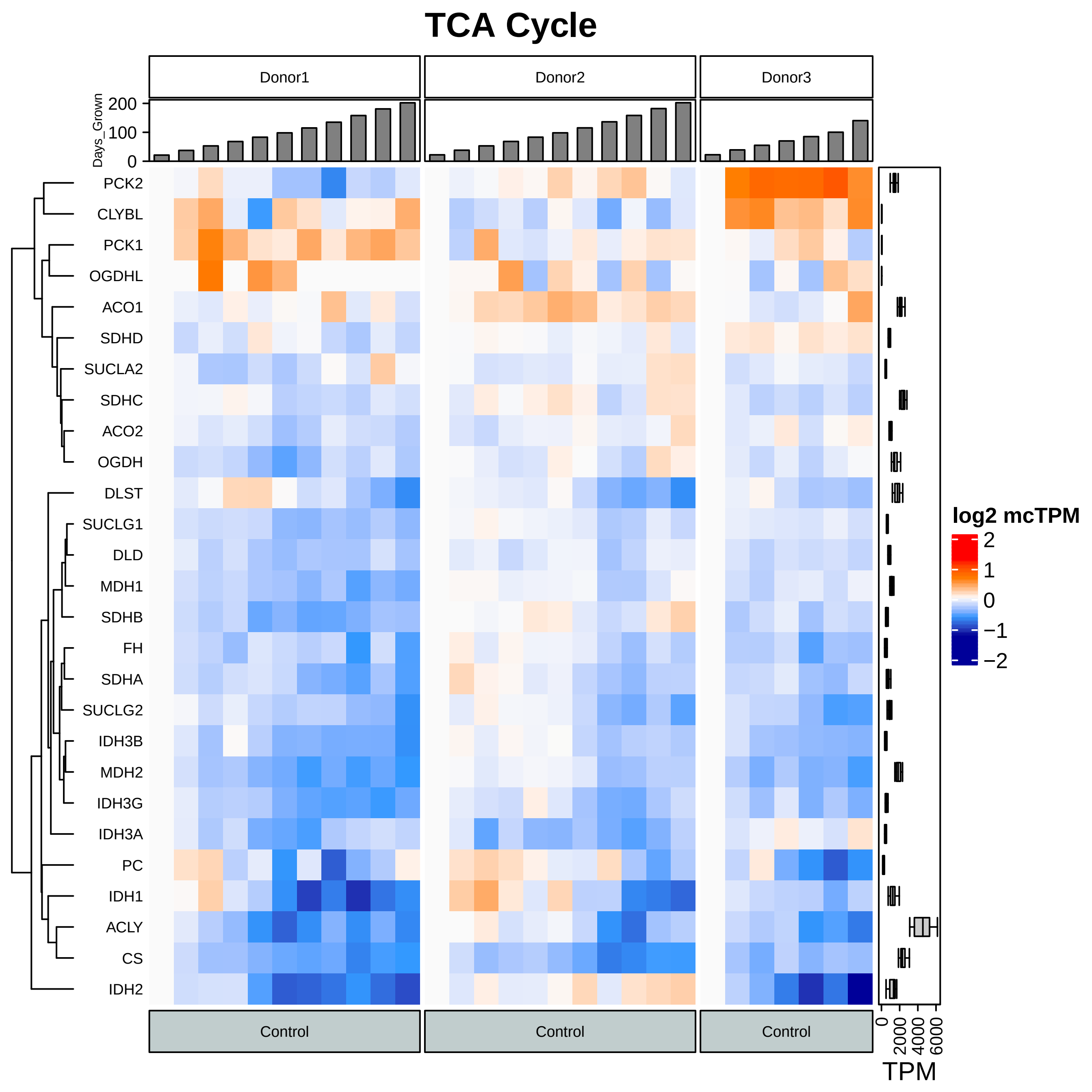
TCA Cycle
Donor1
Donor2
Donor3
Days_Grown
0
100
200
PCK2
CLYBL
PCK1
OGDHL
ACO1
SDHD
SUCLA2
SDHC
ACO2
OGDH
DLST
SUCLG1
DLD
MDH1
SDHB
FH
SDHA
SUCLG2
IDH3B
MDH2
IDH3G
IDH3A
PC
IDH1
ACLY
CS
IDH2
Control
Control
Control
log2 mcTPM
2
1
0
−1
−2
0
2000
4000
6000
TPM

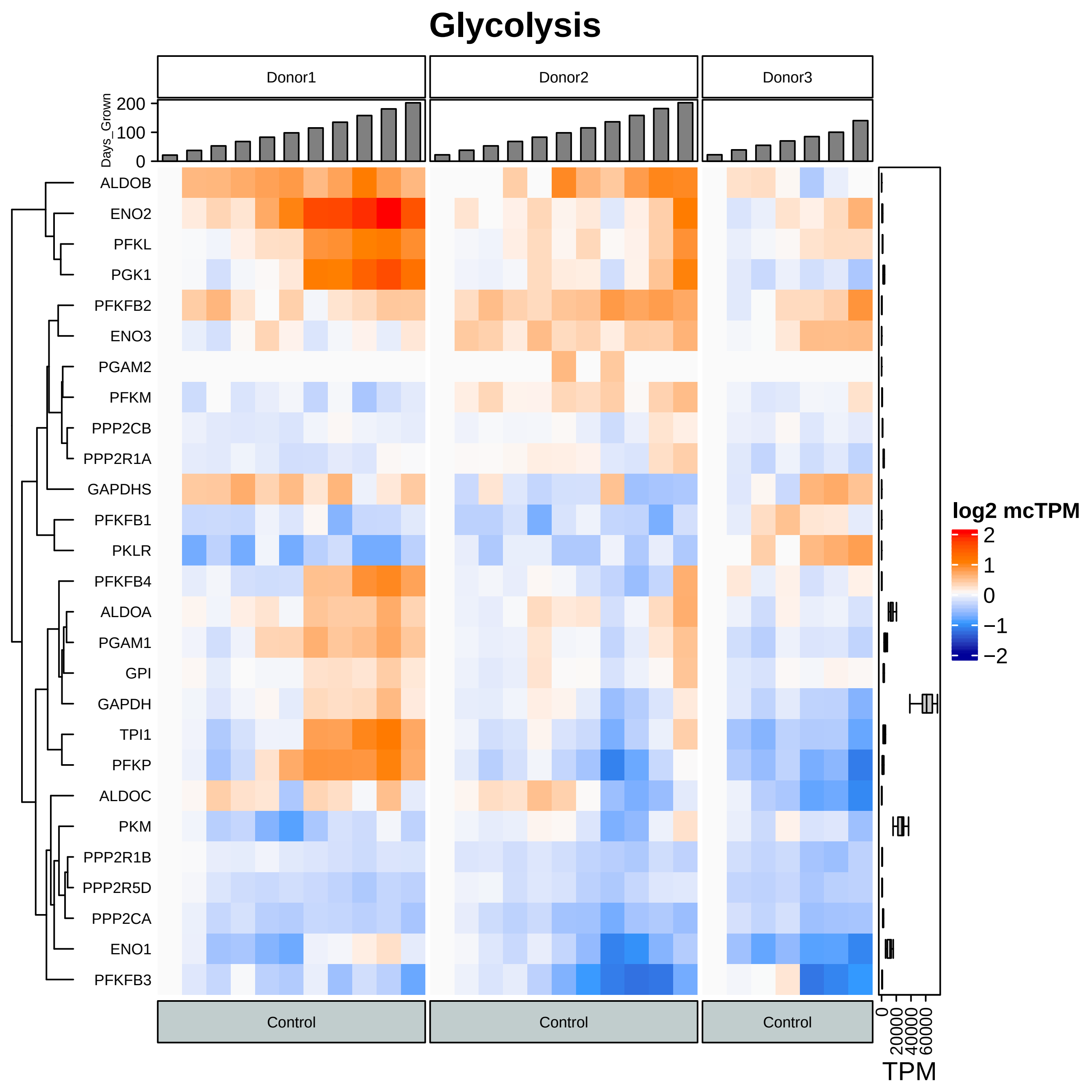
Glycolysis
Donor1
Donor2
Donor3
Days_Grown
200
100
0
ALDOB
ENO2
PFKL
PGK1
PFKFB2
ENO3
PGAM2
PFKM
PPP2CB
PPP2R1A
GAPDHS
PFKFB1
PKLR
PFKFB4
ALDOA
PGAM1
GPI
GAPDH
TPI1
PFKP
ALDOC
PKM
PPP2R1B
PPP2R5D
PPP2CA
ENO1
PFKFB3
log2 mcTPM
2
1
0
−1
−2
Control
Control
Control
0
20000
40000
60000
TPM

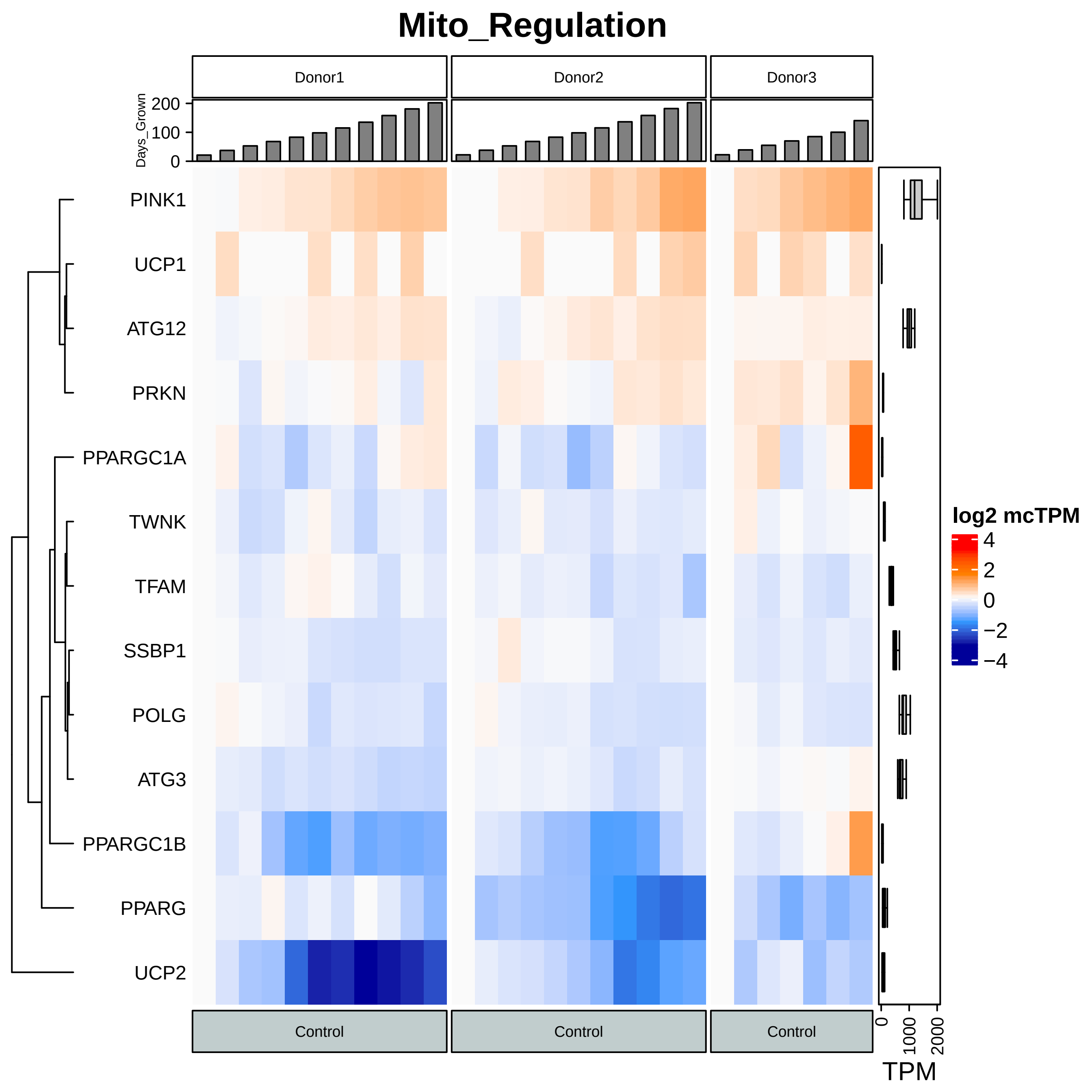

Mito_Regulation
Donor1
Donor2
Donor3
Days_Grown
0
100
200
PINK1
UCP1
ATG12
PRKN
PPARGC1A
TWNK
TFAM
SSBP1
POLG
ATG3
PPARGC1B
PPARG
UCP2
log2 mcTPM
4
2
0
−2
−4
Control
Control
Control
0
1000
2000
TPM

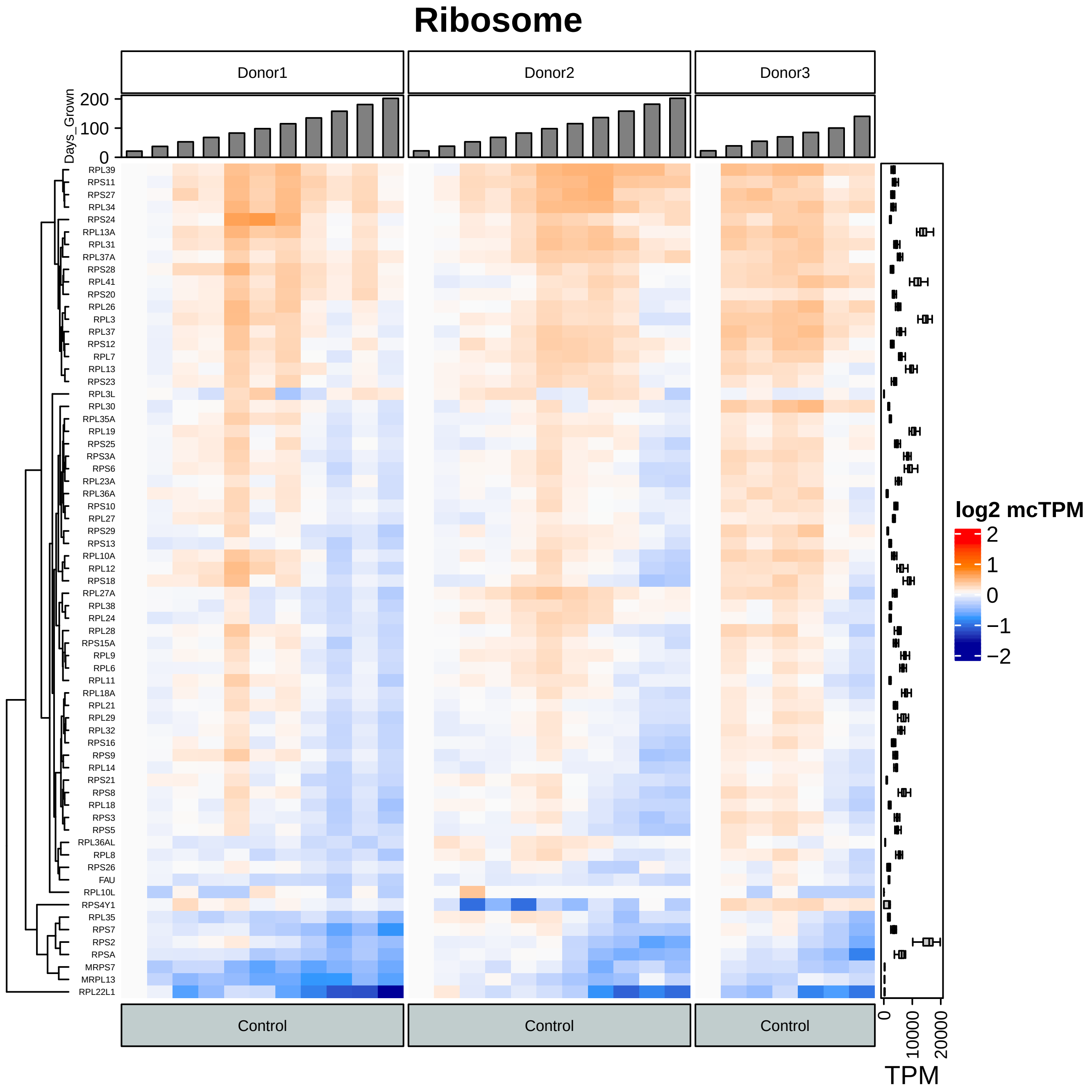
Ribosome
Donor1
Donor2
Donor3
Days_Grown
200
100
0
RPL39
RPS11
RPS27
RPL34
RPS24
RPL13A
RPL31
RPL37A
RPS28
RPL41
RPS20
RPL26
RPL3
RPL37
RPS12
RPL7
RPL13
RPS23
RPL3L
RPL30
RPL35A
RPL19
RPS25
RPS3A
RPS6
RPL23A
RPL36A
RPS10
RPL27
RPS29
RPS13
RPL10A
RPL12
RPS18
RPL27A
RPL38
RPL24
RPL28
RPS15A
RPL9
RPL6
RPL11
RPL18A
RPL21
RPL29
RPL32
RPS16
RPS9
RPL14
RPS21
RPS8
RPL18
RPS3
RPS5
RPL36AL
RPL8
RPS26
FAU
RPL10L
RPS4Y1
RPL35
RPS7
RPS2
RPSA
MRPS7
MRPL13
RPL22L1
log2 mcTPM
2
1
0
−1
−2
Control
Control
Control
0
10000
20000
TPM

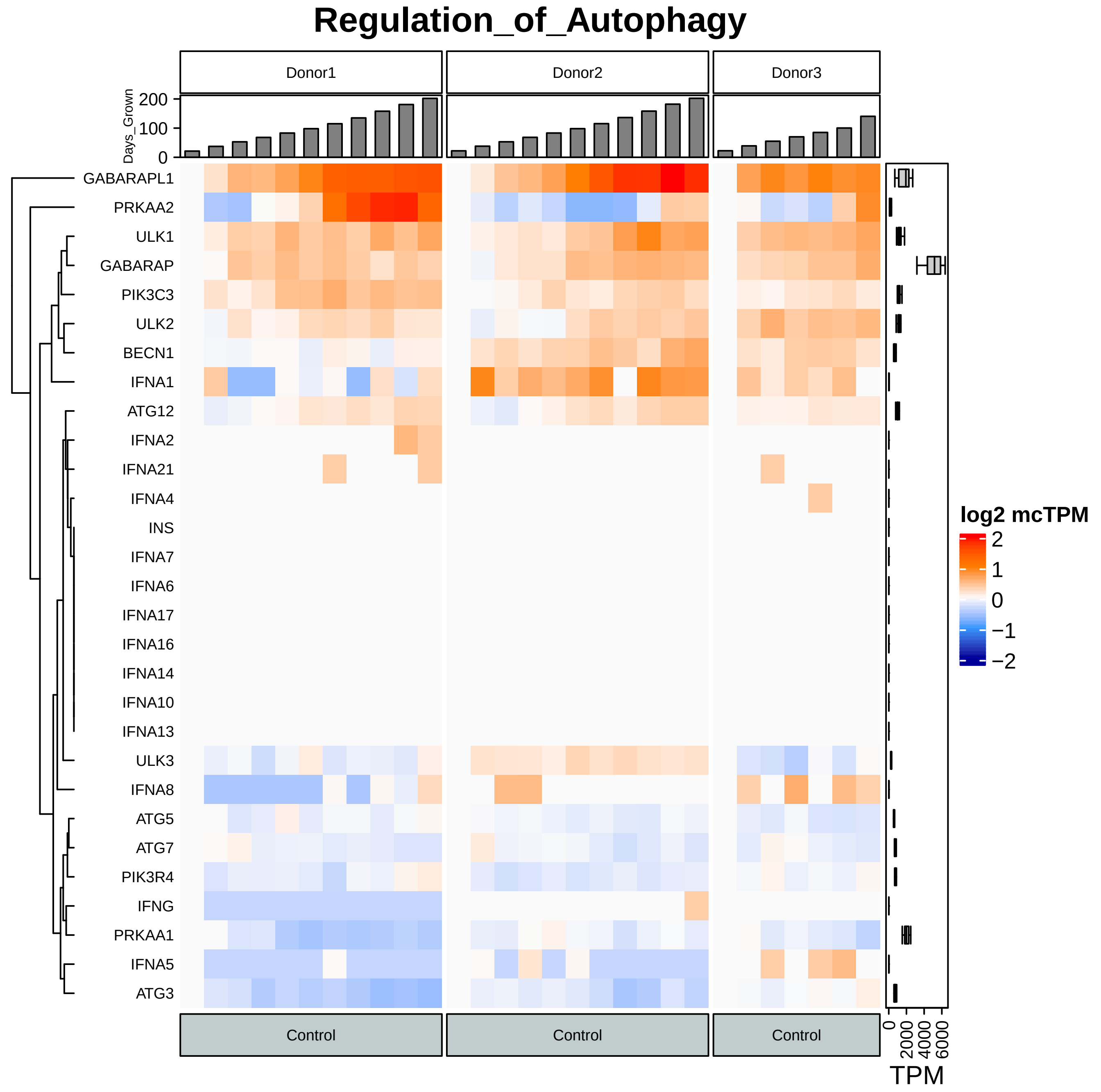

Regulation_of_Autophagy
Donor1
Donor2
Donor3
Days_Grown
200
100
0
GABARAPL1
PRKAA2
ULK1
GABARAP
PIK3C3
ULK2
BECN1
IFNA1
ATG12
IFNA2
IFNA21
IFNA4
INS
IFNA7
IFNA6
IFNA17
IFNA16
IFNA14
IFNA10
IFNA13
ULK3
IFNA8
ATG5
ATG7
PIK3R4
IFNG
PRKAA1
IFNA5
ATG3
log2 mcTPM
2
1
0
−1
−2
Control
Control
Control
0
2000
4000
6000
TPM

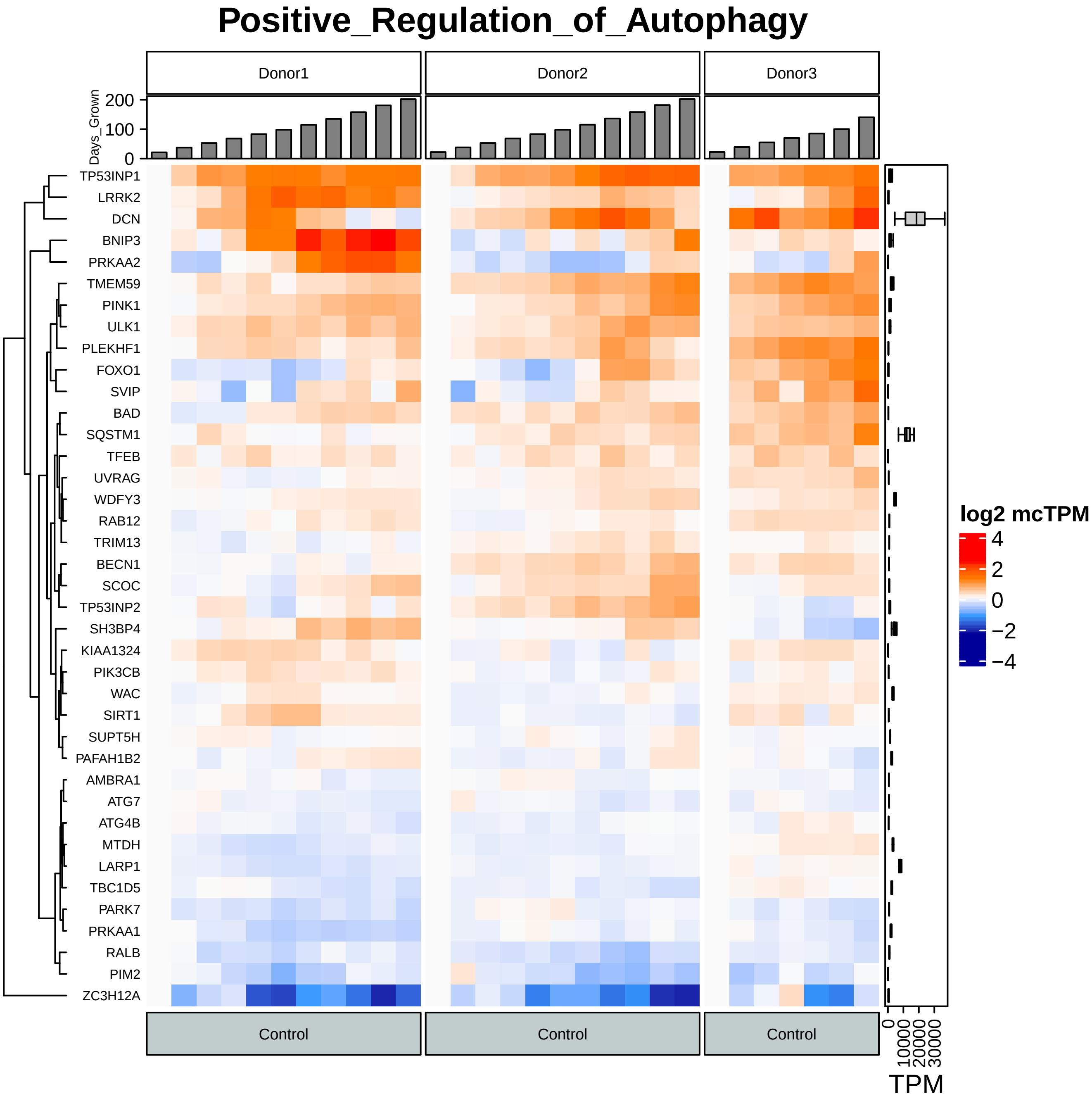
Positive_Regulation_of_Autophagy
Donor1
Donor2
Donor3
Days_Grown
200
100
0
TP53INP1
LRRK2
DCN
BNIP3
PRKAA2
TMEM59
PINK1
ULK1
PLEKHF1
FOXO1
SVIP
BAD
SQSTM1
TFEB
UVRAG
WDFY3
RAB12
TRIM13
BECN1
SCOC
TP53INP2
SH3BP4
KIAA1324
PIK3CB
WAC
SIRT1
SUPT5H
PAFAH1B2
AMBRA1
ATG7
ATG4B
MTDH
LARP1
TBC1D5
PARK7
PRKAA1
RALB
PIM2
ZC3H12A
log2 mcTPM
4
2
0
−2
−4
Control
Control
Control
0
10000
20000
30000
TPM

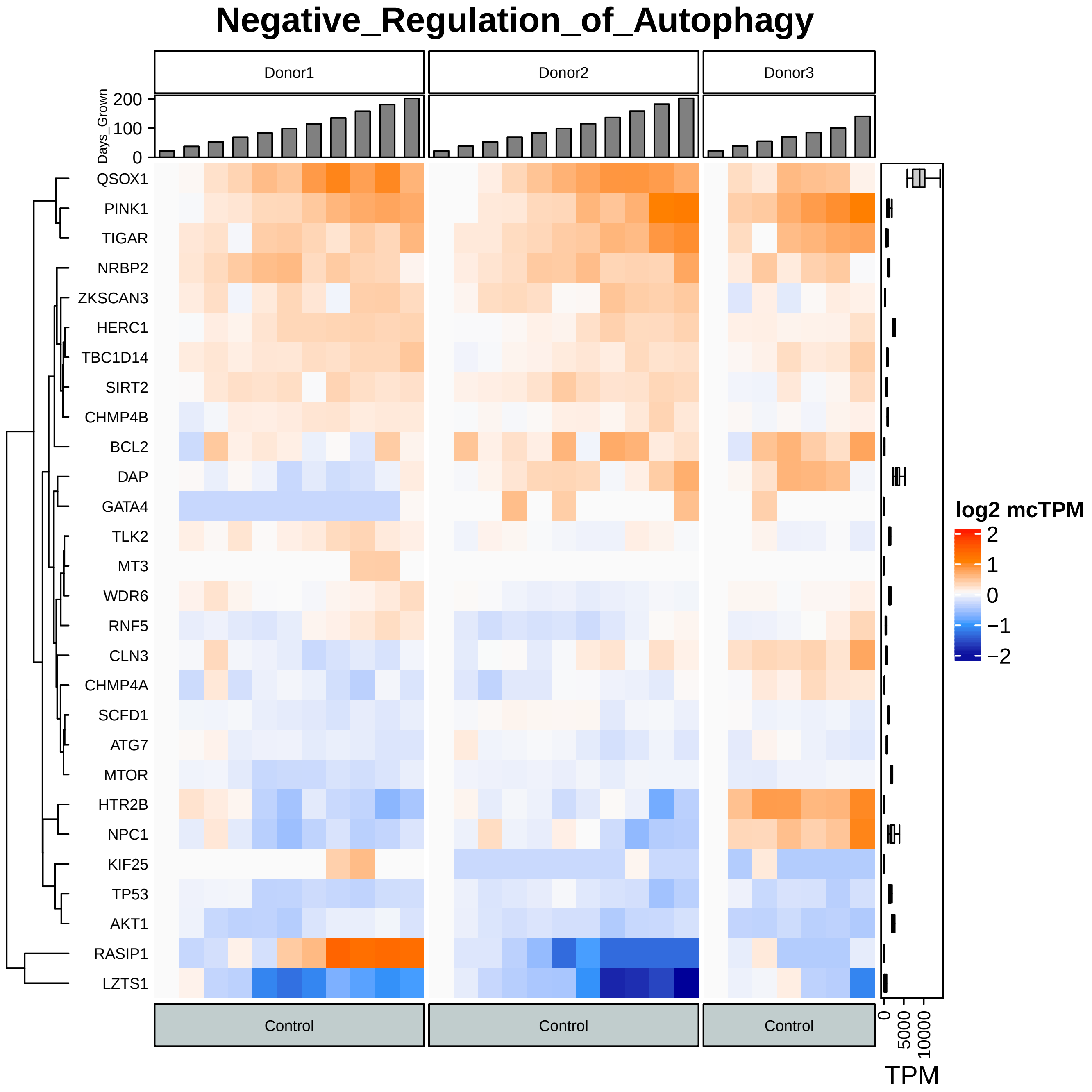
Negative_Regulation_of_Autophagy
Donor1
Donor2
Donor3
Days_Grown
200
100
0
QSOX1
PINK1
TIGAR
NRBP2
ZKSCAN3
HERC1
TBC1D14
SIRT2
CHMP4B
BCL2
DAP
GATA4
TLK2
MT3
WDR6
RNF5
CLN3
CHMP4A
SCFD1
ATG7
MTOR
HTR2B
NPC1
KIF25
TP53
AKT1
RASIP1
LZTS1
log2 mcTPM
2
1
0
−1
−2
Control
Control
Control
0
5000
10000
TPM

### Contact_Inhibition

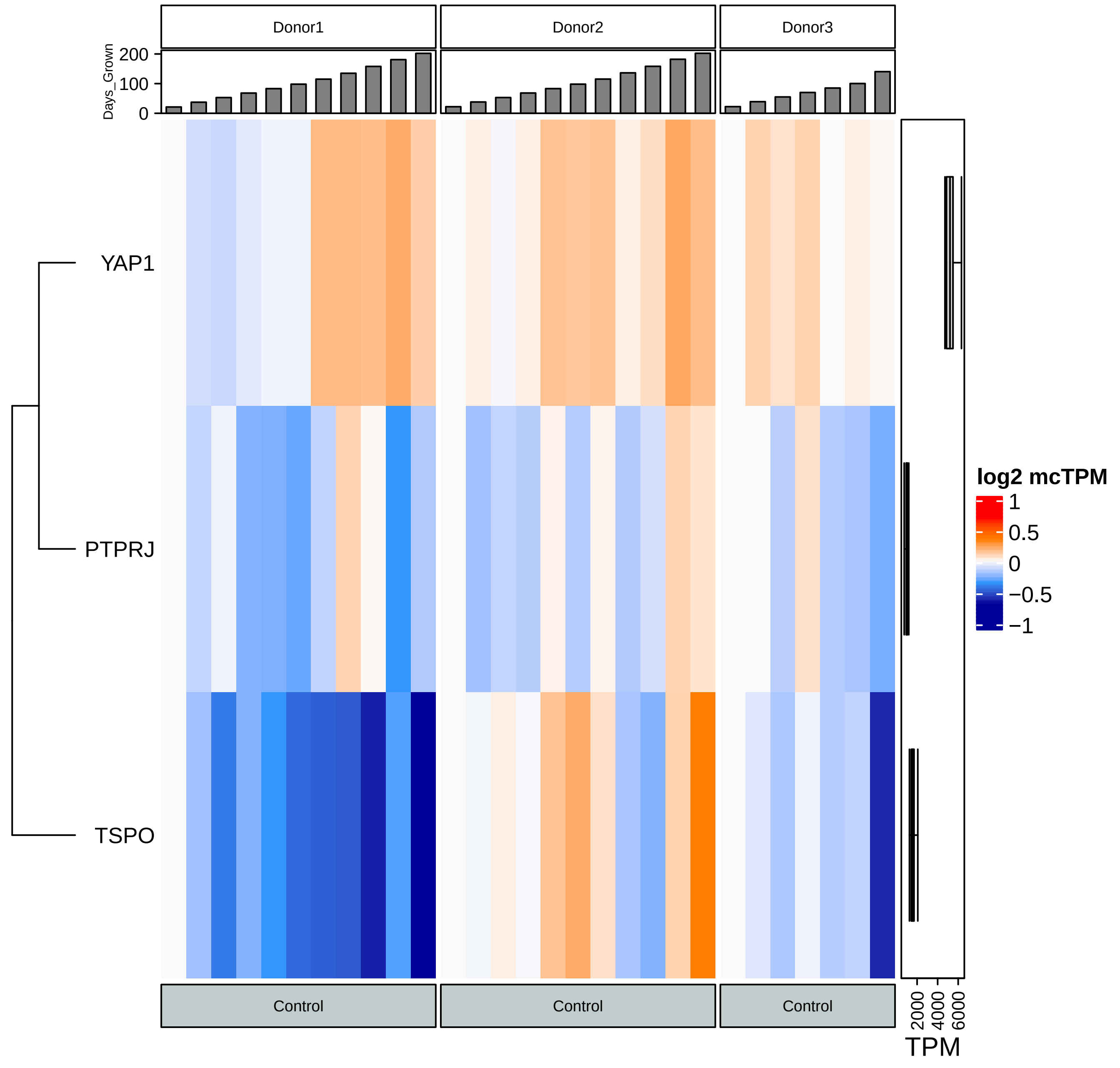

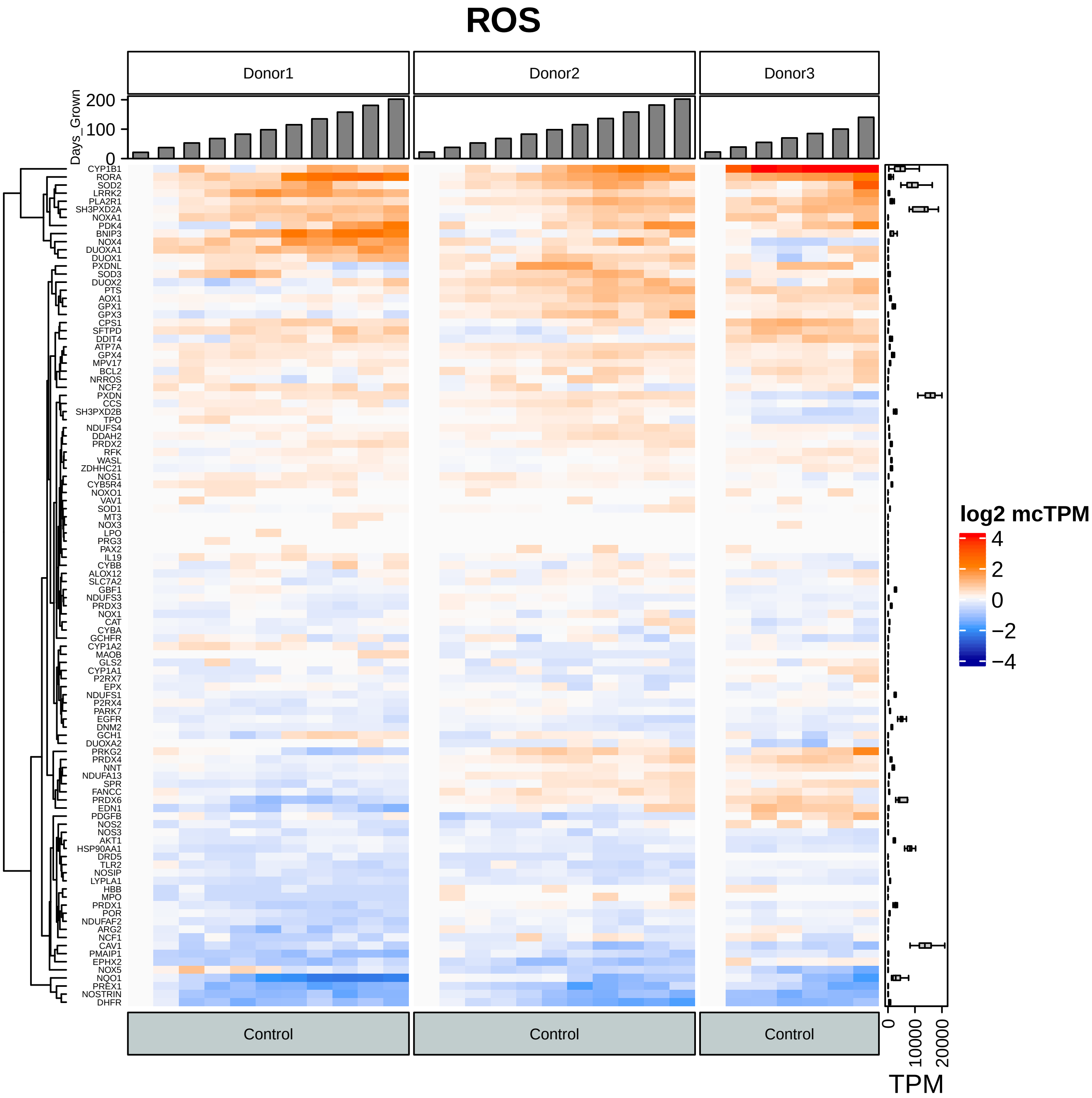
ROS
Donor1
Donor2
Donor3
Days_Grown
0
100
200
CYP1B1
RORA
SOD2
LRRK2
PLA2R1
SH3PXD2A
NOXA1
PDK4
BNIP3
NOX4
DUOXA1
DUOX1
PXDNL
SOD3
DUOX2
PTS
AOX1
GPX1
GPX3
CPS1
SFTPD
DDIT4
ATP7A
GPX4
MPV17
BCL2
NRROS
NCF2
PXDN
CCS
SH3PXD2B
TPO
NDUFS4
DDAH2
PRDX2
RFK
WASL
ZDHHC21
NOS1
CYB5R4
NOXO1
VAV1
SOD1
MT3
NOX3
LPO
PRG3
PAX2
IL19
CYBB
ALOX12
SLC7A2
GBF1
NDUFS3
PRDX3
NOX1
CAT
CYBA
GCHFR
CYP1A2
MAOB
GLS2
CYP1A1
P2RX7
EPX
NDUFS1
P2RX4
PARK7
EGFR
DNM2
GCH1
DUOXA2
PRKG2
PRDX4
NNT
NDUFA13
SPR
FANCC
PRDX6
EDN1
PDGFB
NOS2
NOS3
AKT1
HSP90AA1
DRD5
TLR2
NOSIP
LYPLA1
HBB
MPO
PRDX1
POR
NDUFAF2
ARG2
NCF1
CAV1
PMAIP1
EPHX2
NOX5
NQO1
PREX1
NOSTRIN
DHFR
log2 mcTPM
4
2
0
−2
−4
Control
Control
Control
0
10000
20000
TPM

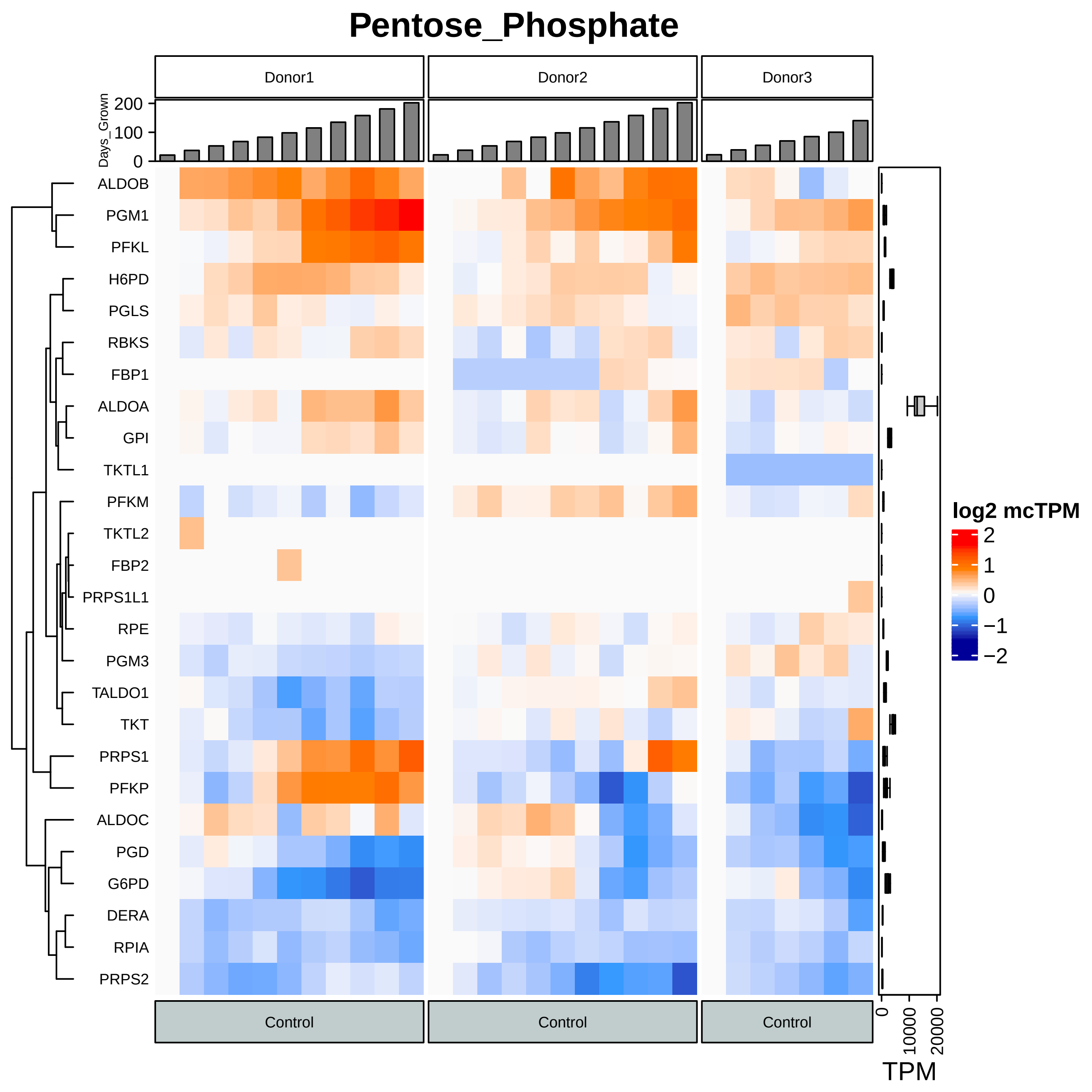
Pentose_Phosphate
Donor1
Donor2
Donor3
Days_Grown
200
100
0
ALDOB
PGM1
PFKL
H6PD
PGLS
RBKS
FBP1
ALDOA
GPI
TKTL1
PFKM
TKTL2
FBP2
PRPS1L1
RPE
PGM3
TALDO1
TKT
PRPS1
PFKP
ALDOC
PGD
G6PD
DERA
RPIA
PRPS2
log2 mcTPM
2
1
0
−1
−2
Control
Control
Control
0
10000
20000
TPM

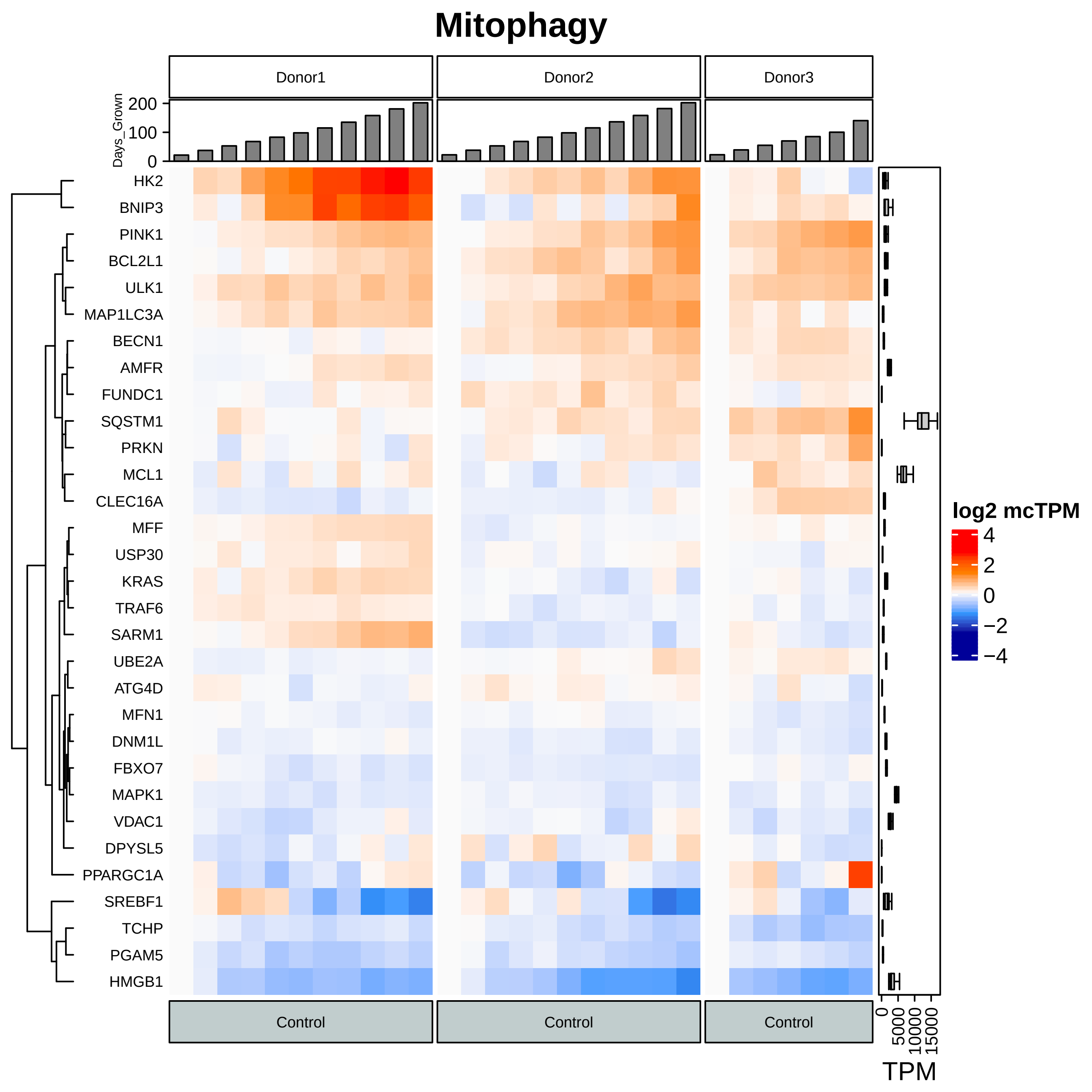

Mitophagy
Donor1
Donor2
Donor3
Days_Grown
200
100
0
HK2
BNIP3
PINK1
BCL2L1
ULK1
MAP1LC3A
BECN1
AMFR
FUNDC1
SQSTM1
PRKN
MCL1
CLEC16A
MFF
USP30
KRAS
TRAF6
SARM1
UBE2A
ATG4D
MFN1
DNM1L
FBXO7
MAPK1
VDAC1
DPYSL5
PPARGC1A
SREBF1
TCHP
PGAM5
HMGB1
log2 mcTPM
4
2
0
−2
−4
Control
Control
Control
0
5000
10000
15000
TPM

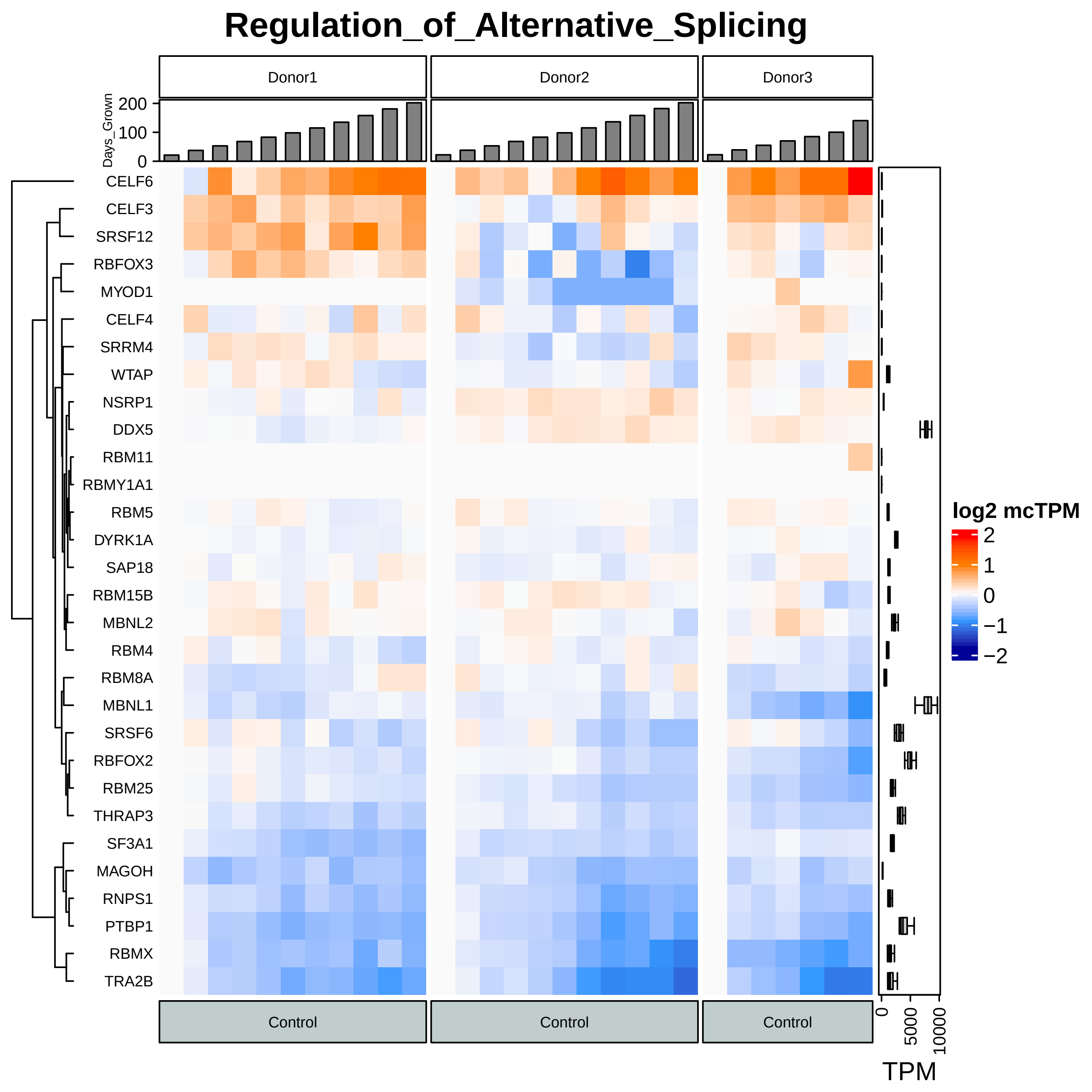
Regulation_of_Alternative_Splicing
Donor1
Donor2
Donor3
Days_Grown
200
100
0
CELF6
CELF3
SRSF12
RBFOX3
MYOD1
CELF4
SRRM4
WTAP
NSRP1
DDX5
RBM11
RBMY1A1
RBM5
DYRK1A
SAP18
RBM15B
MBNL2
RBM4
RBM8A
MBNL1
SRSF6
RBFOX2
RBM25
THRAP3
SF3A1
MAGOH
RNPS1
PTBP1
RBMX
TRA2B
log2 mcTPM
2
1
0
−1
−2
Control
Control
Control
0
5000
10000
TPM

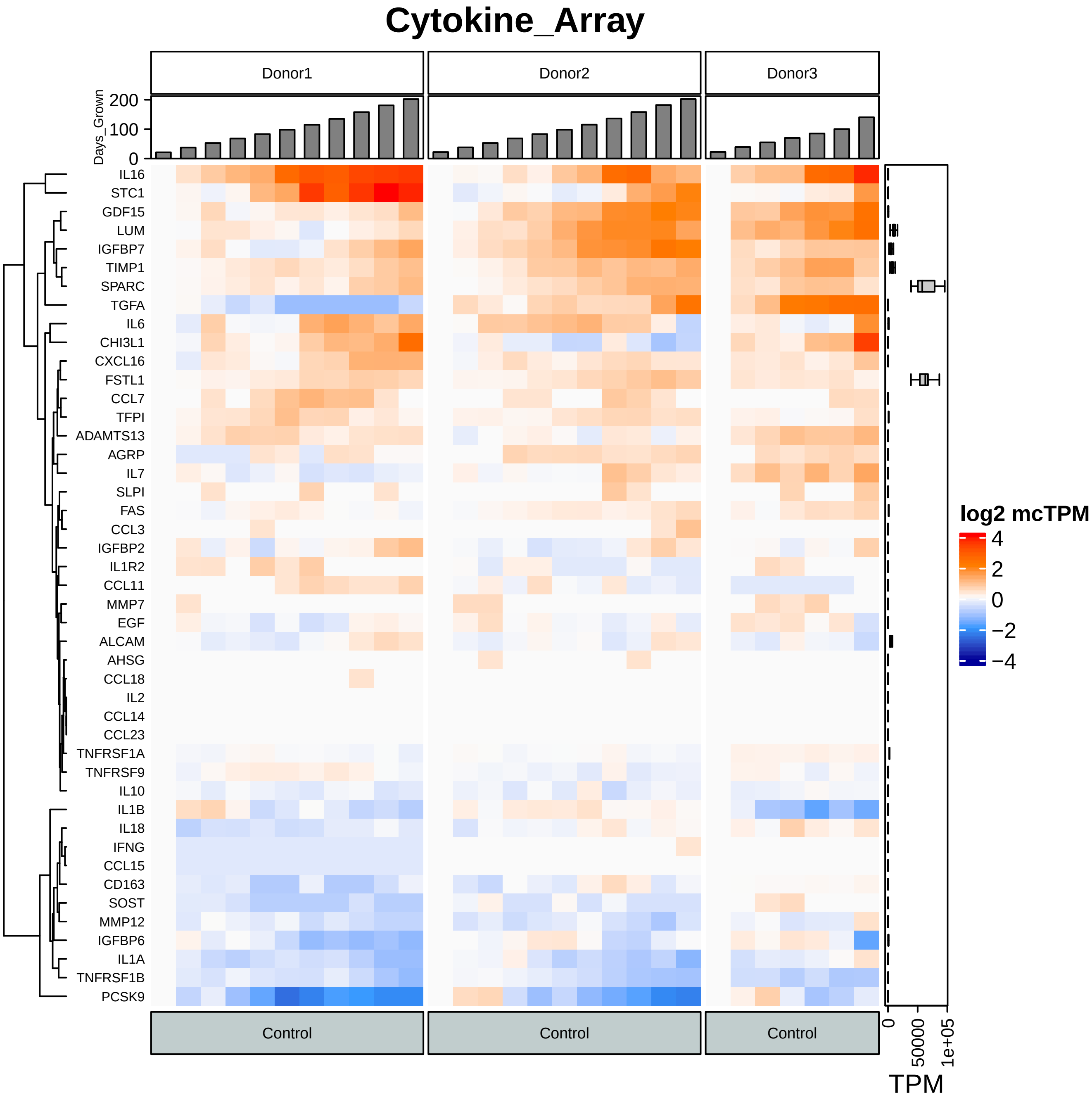
Cytokine_Array
Donor1
Donor2
Donor3
Days_Grown
200
100
0
IL16
STC1
GDF15
LUM
IGFBP7
TIMP1
SPARC
TGFA
IL6
CHI3L1
CXCL16
FSTL1
CCL7
TFPI
ADAMTS13
AGRP
IL7
SLPI
FAS
CCL3
IGFBP2
IL1R2
CCL11
MMP7
EGF
ALCAM
AHSG
CCL18
IL2
CCL14
CCL23
TNFRSF1A
TNFRSF9
IL10
IL1B
IL18
IFNG
CCL15
CD163
SOST
MMP12
IGFBP6
IL1A
TNFRSF1B
PCSK9
Control
Control
Control
log2 mcTPM
4
2
0
−2
−4
0
50000
1e+05
TPM

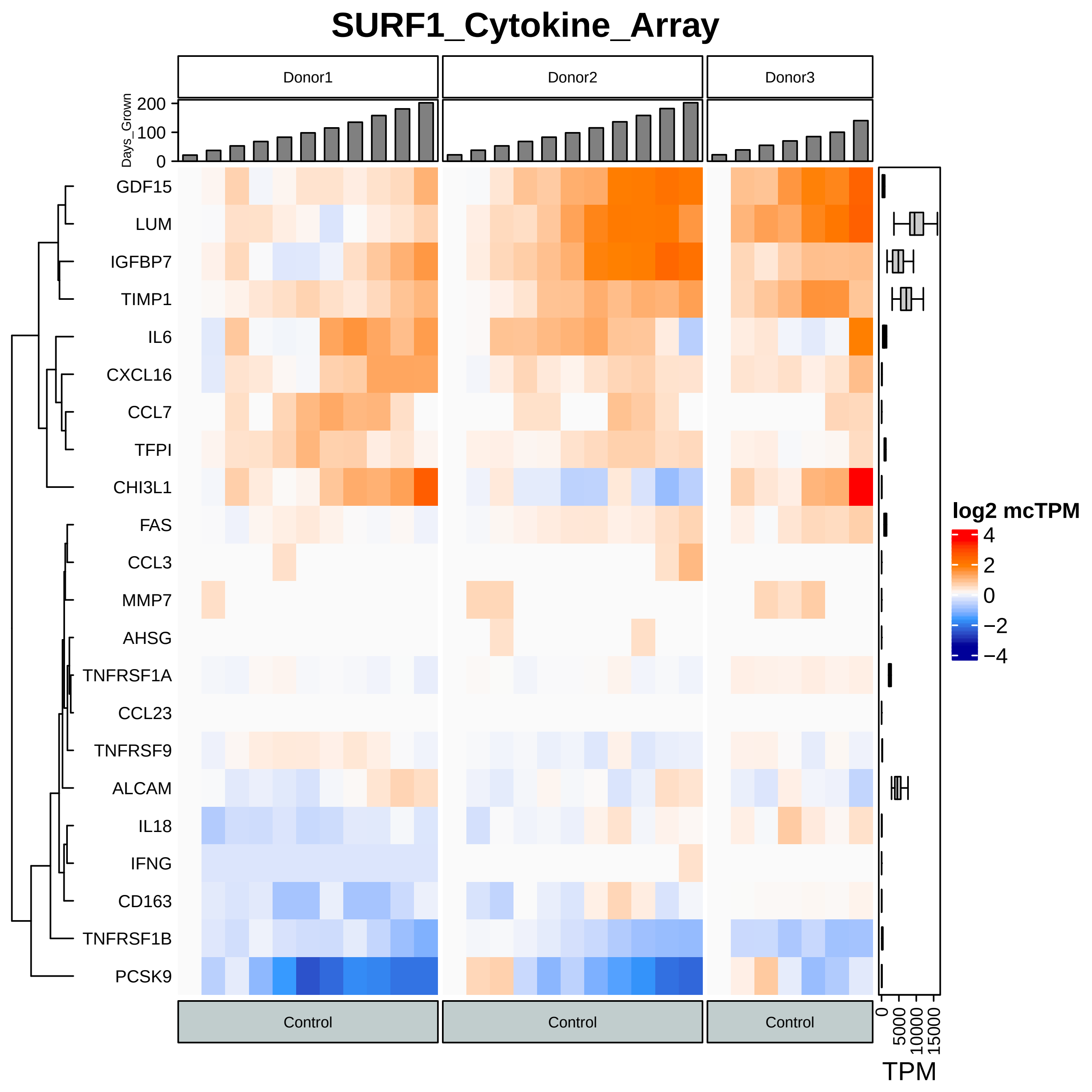
SURF1_Cytokine_Array
Donor1
Donor2
Donor3
Days_Grown
200
100
0
GDF15
LUM
IGFBP7
TIMP1
IL6
CXCL16
CCL7
TFPI
CHI3L1
FAS
CCL3
MMP7
AHSG
TNFRSF1A
CCL23
TNFRSF9
ALCAM
IL18
IFNG
CD163
TNFRSF1B
PCSK9
Control
Control
Control
log2 mcTPM
4
2
0
−2
−4
0
5000
10000
15000
TPM

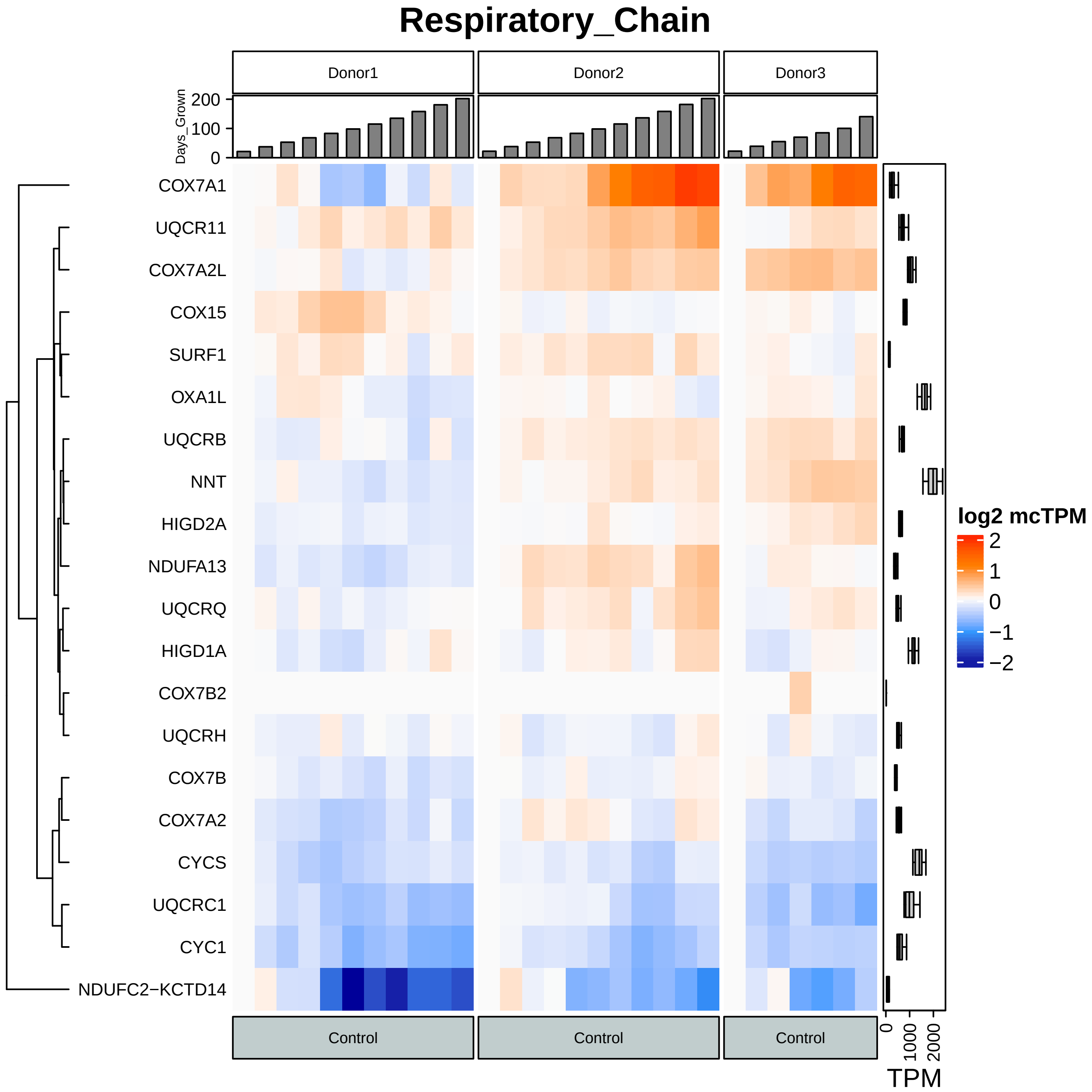

Respiratory_Chain
Donor1
Donor2
Donor3
Days_Grown
200
100
0
COX7A1
UQCR11
COX7A2L
COX15
SURF1
OXA1L
UQCRB
NNT
HIGD2A
NDUFA13
UQCRQ
HIGD1A
COX7B2
UQCRH
COX7B
COX7A2
CYCS
UQCRC1
CYC1
NDUFC2−KCTD14
log2 mcTPM
2
1
0
−1
−2
Control
Control
Control
0
1000
2000
TPM

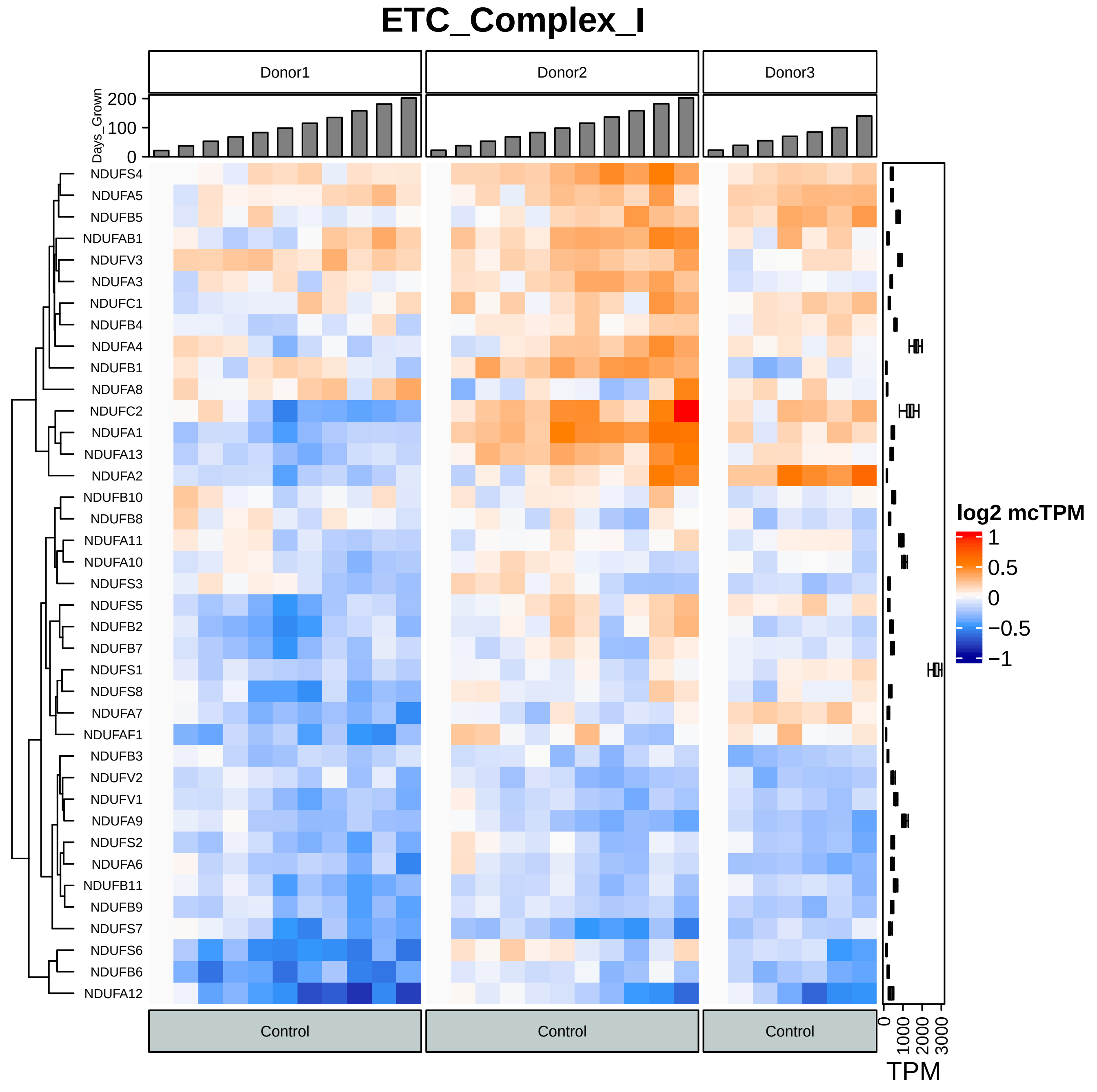
ETC_Complex_I
Donor1
Donor2
Donor3
Days_Grown
200
100
0
NDUFS4
NDUFA5
NDUFB5
NDUFAB1
NDUFV3
NDUFA3
NDUFC1
NDUFB4
NDUFA4
NDUFB1
NDUFA8
NDUFC2
NDUFA1
NDUFA13
NDUFA2
NDUFB10
NDUFB8
NDUFA11
NDUFA10
NDUFS3
NDUFS5
NDUFB2
NDUFB7
NDUFS1
NDUFS8
NDUFA7
NDUFAF1
NDUFB3
NDUFV2
NDUFV1
NDUFA9
NDUFS2
NDUFA6
NDUFB11
NDUFB9
NDUFS7
NDUFS6
NDUFB6
NDUFA12
Control
Control
Control
log2 mcTPM
1
0.5
0
−0.5
−1
0
1000
2000
3000
TPM

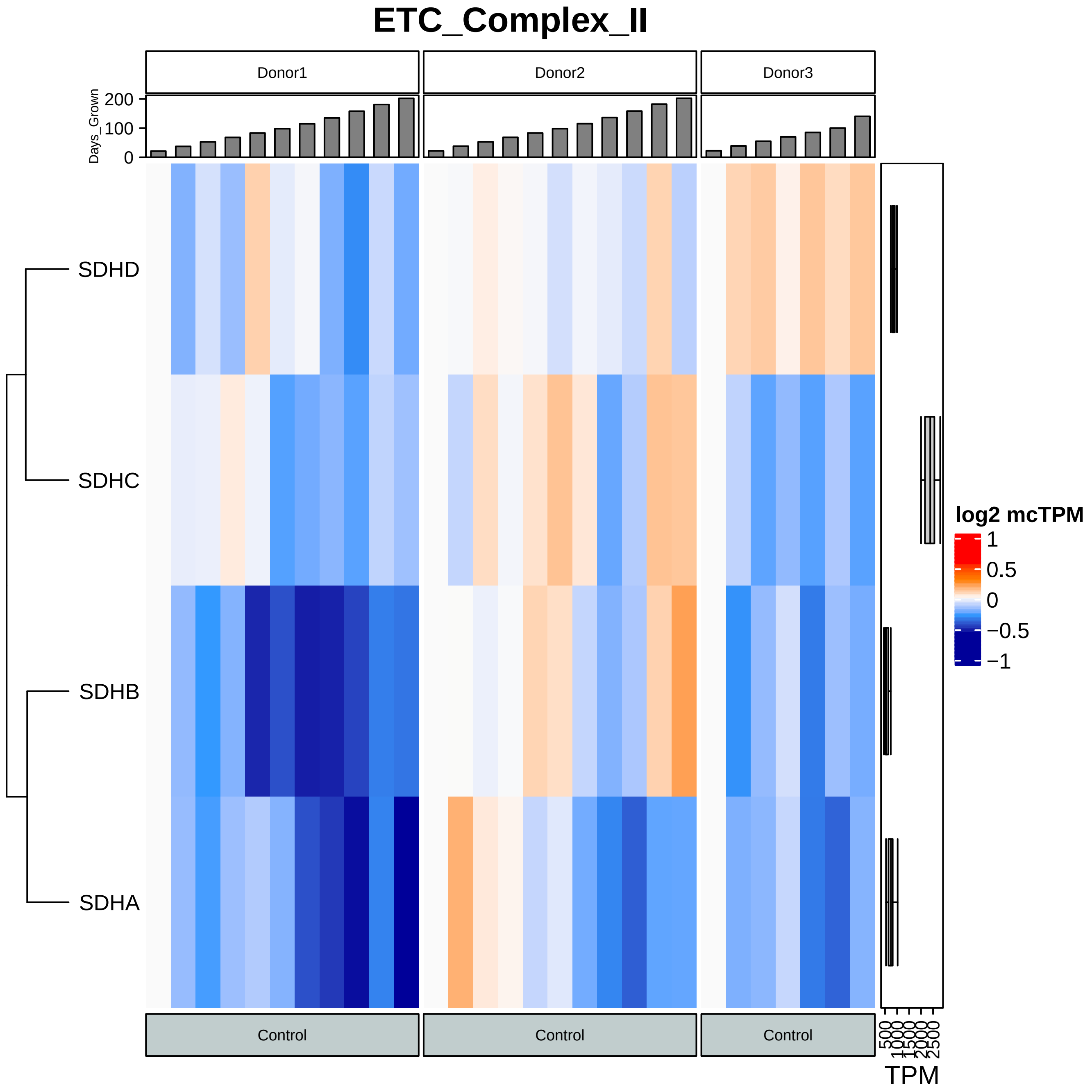
ETC_Complex_II
Donor1
Donor2
Donor3
Days_Grown
200
100
0
SDHD
SDHC
SDHB
SDHA
Control
Control
Control
log2 mcTPM
1
0.5
0
−0.5
−1
500
1000
1500
2000
2500
TPM

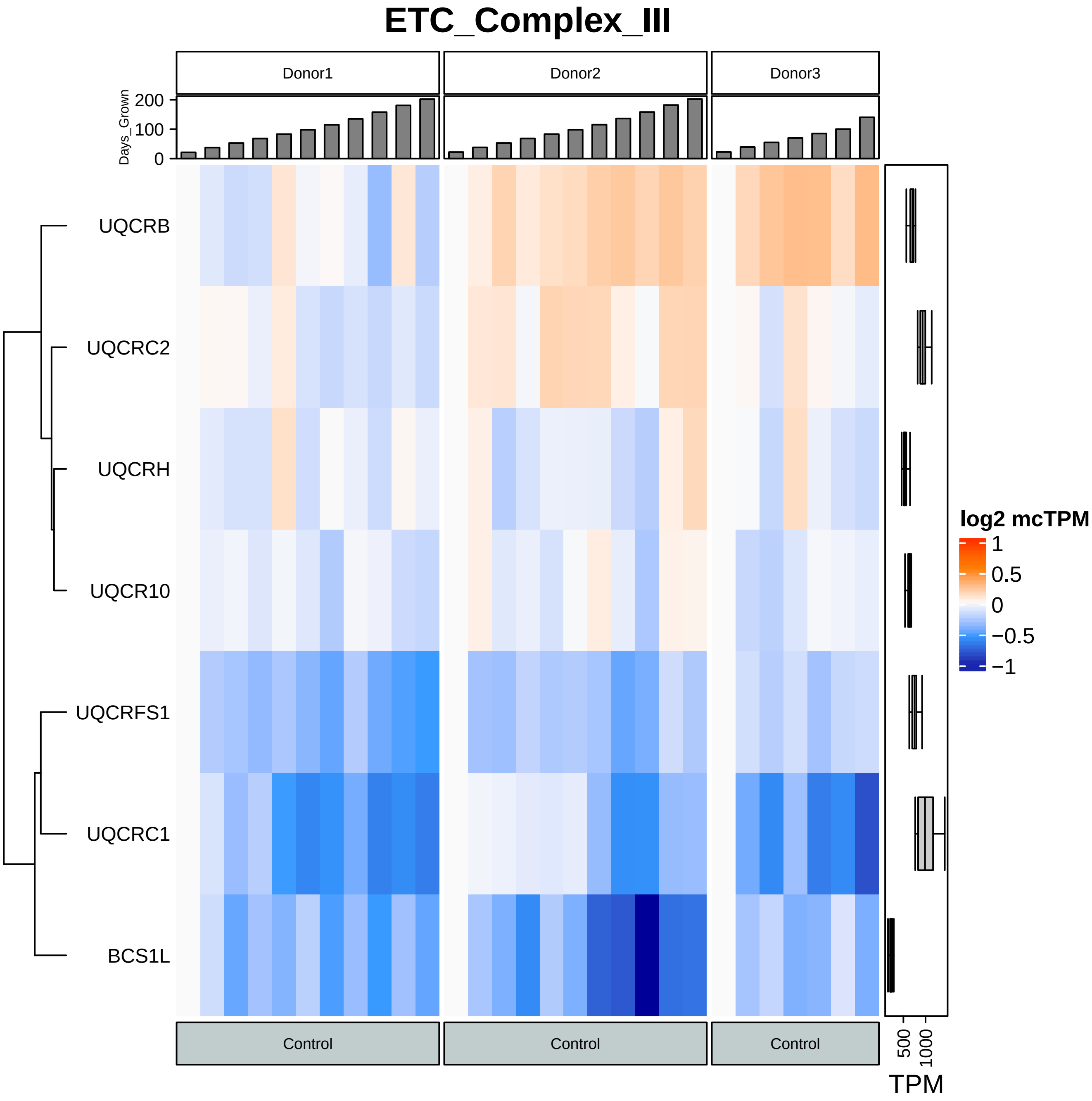
ETC_Complex_III
Donor1
Donor2
Donor3
Days_Grown
200
100
0
UQCRB
UQCRC2
UQCRH
UQCR10
UQCRFS1
UQCRC1
BCS1L
Control
Control
Control
log2 mcTPM
1
0.5
0
−0.5
−1
500
1000
TPM

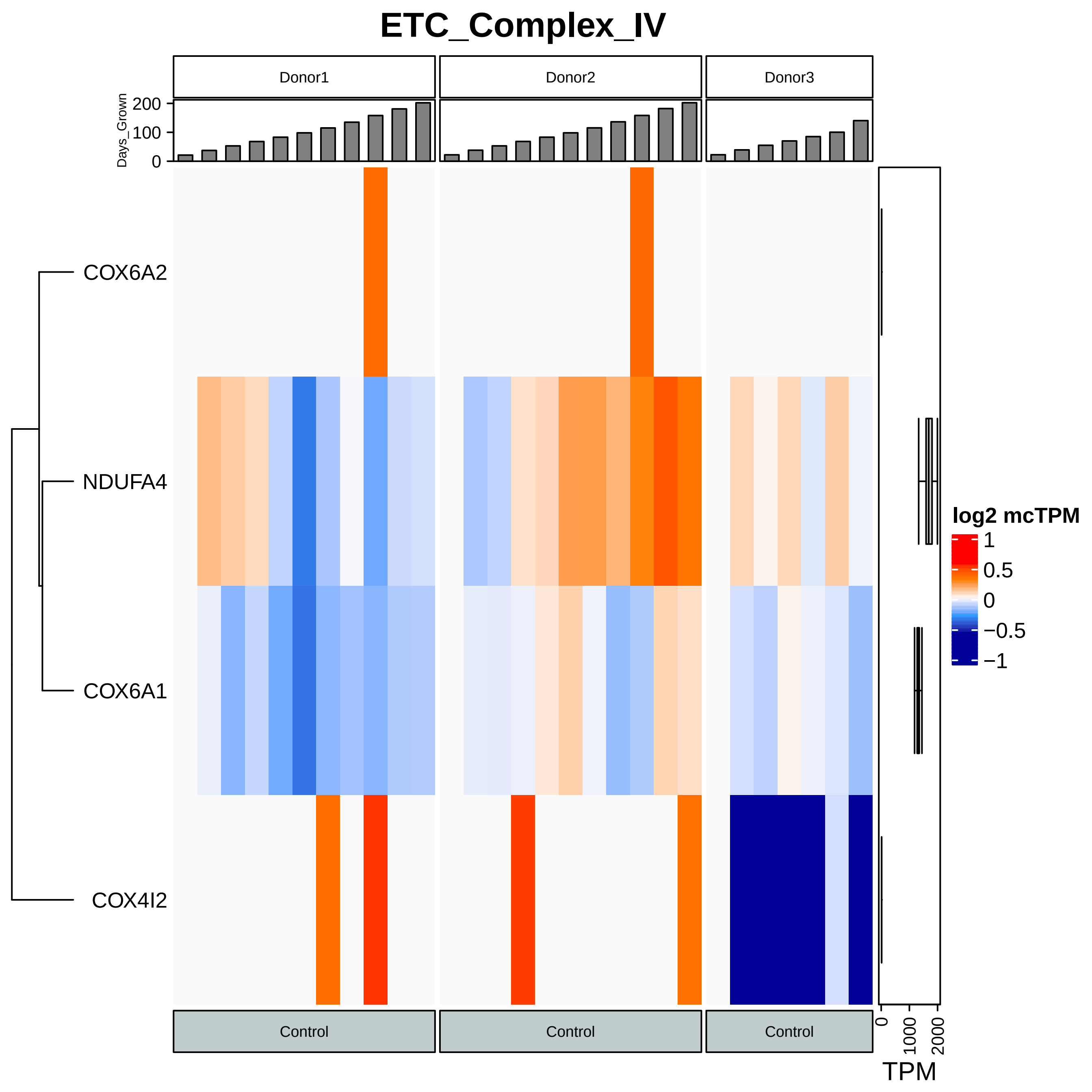

ETC_Complex_IV
Donor1
Donor2
Donor3
Days_Grown
0
100
200
COX6A2
NDUFA4
COX6A1
COX4I2
log2 mcTPM
1
0.5
0
−0.5
−1
Control
Control
Control
0
1000
2000
TPM

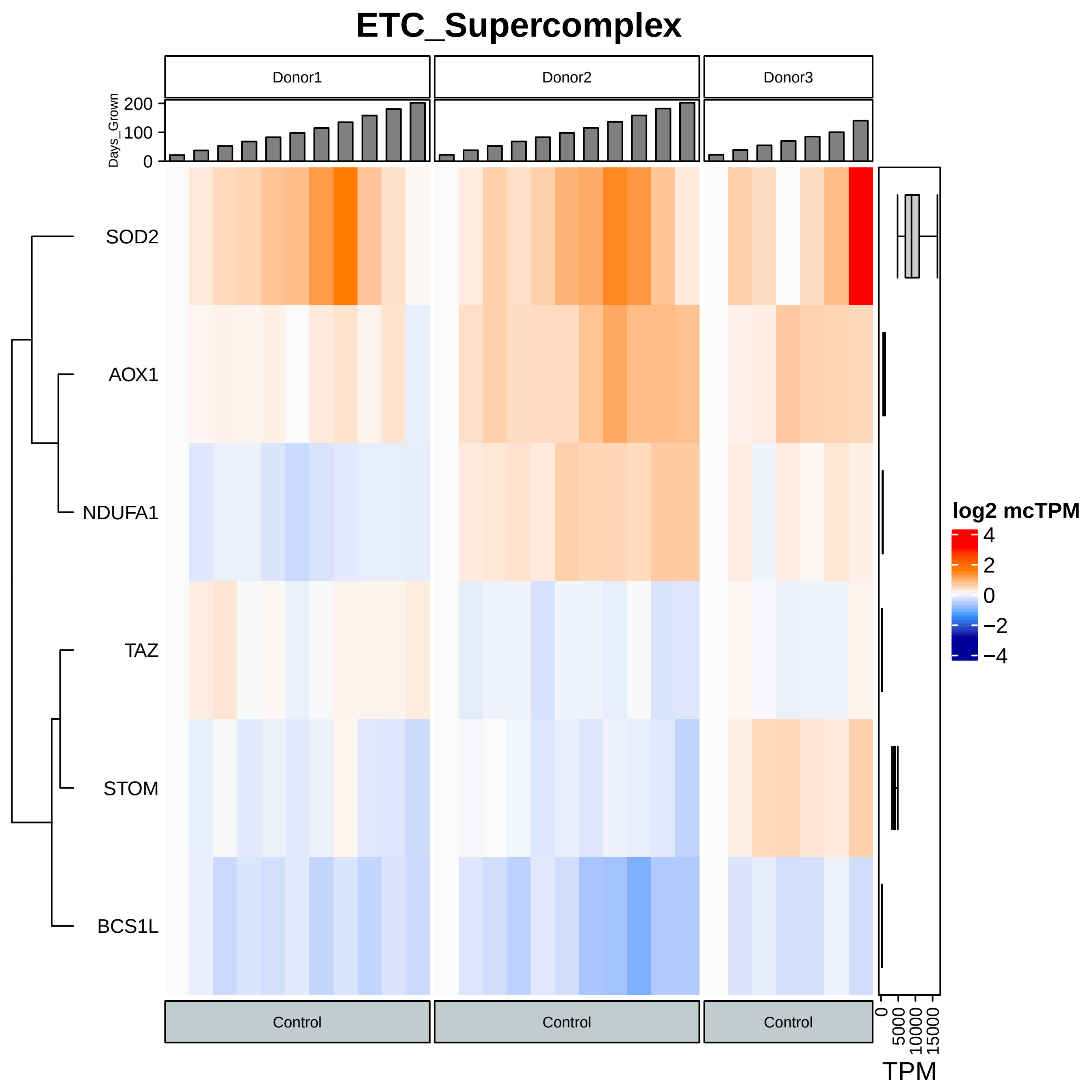

ETC_Supercomplex
Donor1
Donor2
Donor3
Days_Grown
200
100
0
SOD2
AOX1
NDUFA1
TAZ
STOM
BCS1L
log2 mcTPM
4
2
0
−2
−4
Control
Control
Control
0
5000
10000
15000
TPM

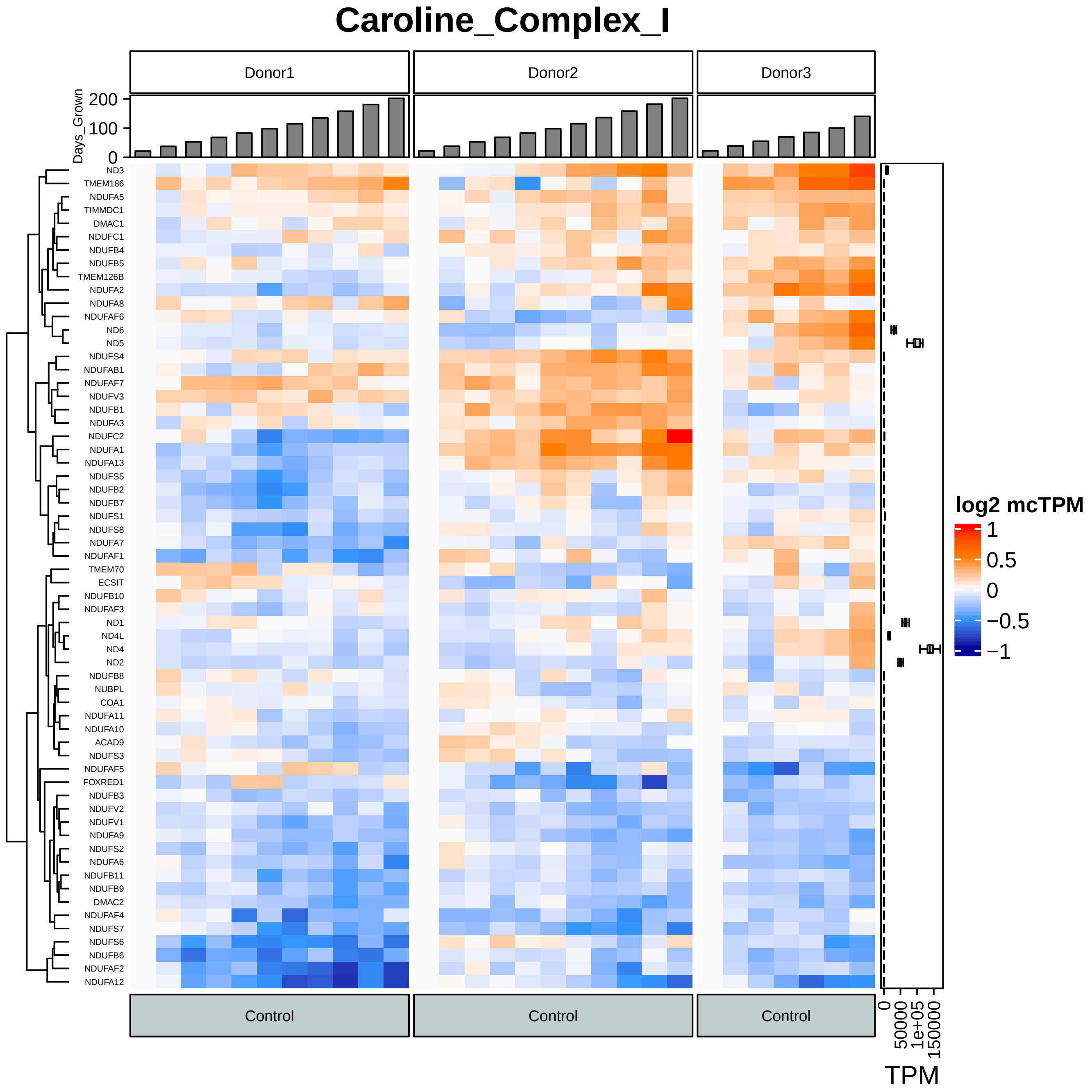

Caroline_Complex_I
Donor1
Donor2
Donor3
Days_Grown
200
100
0
ND3
TMEM186
NDUFA5
TIMMDC1
DMAC1
NDUFC1
NDUFB4
NDUFB5
TMEM126B
NDUFA2
NDUFA8
NDUFAF6
ND6
ND5
NDUFS4
NDUFAB1
NDUFAF7
NDUFV3
NDUFB1
NDUFA3
NDUFC2
NDUFA1
NDUFA13
NDUFS5
NDUFB2
NDUFB7
NDUFS1
NDUFS8
NDUFA7
NDUFAF1
TMEM70
ECSIT
NDUFB10
NDUFAF3
ND1
ND4L
ND4
ND2
NDUFB8
NUBPL
COA1
NDUFA11
NDUFA10
ACAD9
NDUFS3
NDUFAF5
FOXRED1
NDUFB3
NDUFV2
NDUFV1
NDUFA9
NDUFS2
NDUFA6
NDUFB11
NDUFB9
DMAC2
NDUFAF4
NDUFS7
NDUFS6
NDUFB6
NDUFAF2
NDUFA12
log2 mcTPM
1
0.5
0
−0.5
−1
Control
Control
Control
0
50000
1e+05
150000
TPM

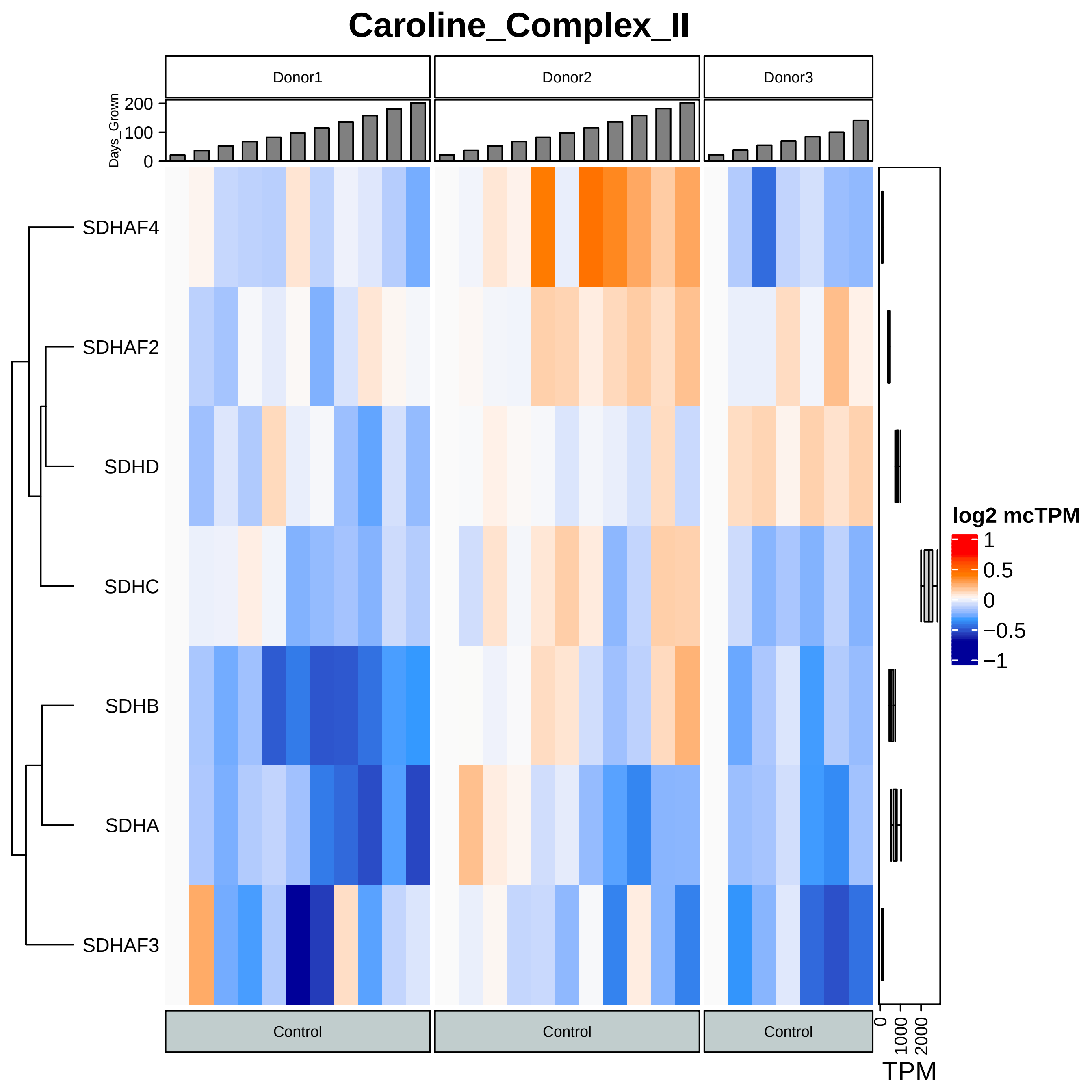

Caroline_Complex_II
Donor1
Donor2
Donor3
Days_Grown
200
100
0
SDHAF4
SDHAF2
SDHD
SDHC
SDHB
SDHA
SDHAF3
log2 mcTPM
1
0.5
0
−0.5
−1
Control
Control
Control
0
1000
2000
TPM

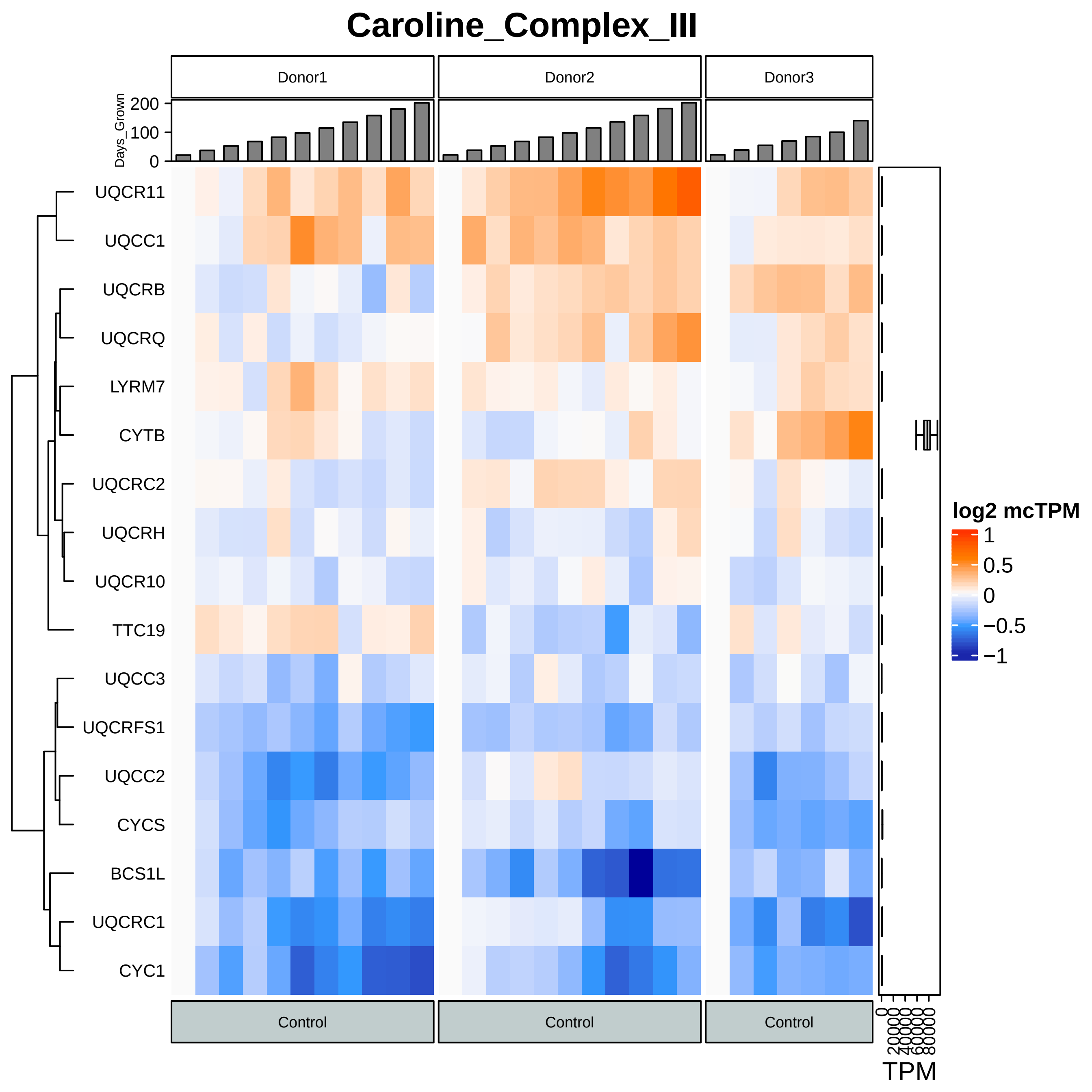

Caroline_Complex_III
Donor1
Donor2
Donor3
Days_Grown
200
100
0
UQCR11
UQCC1
UQCRB
UQCRQ
LYRM7
CYTB
UQCRC2
UQCRH
UQCR10
TTC19
UQCC3
UQCRFS1
UQCC2
CYCS
BCS1L
UQCRC1
CYC1
Control
Control
Control
log2 mcTPM
1
0.5
0
−0.5
−1
0
20000
40000
60000
80000
TPM

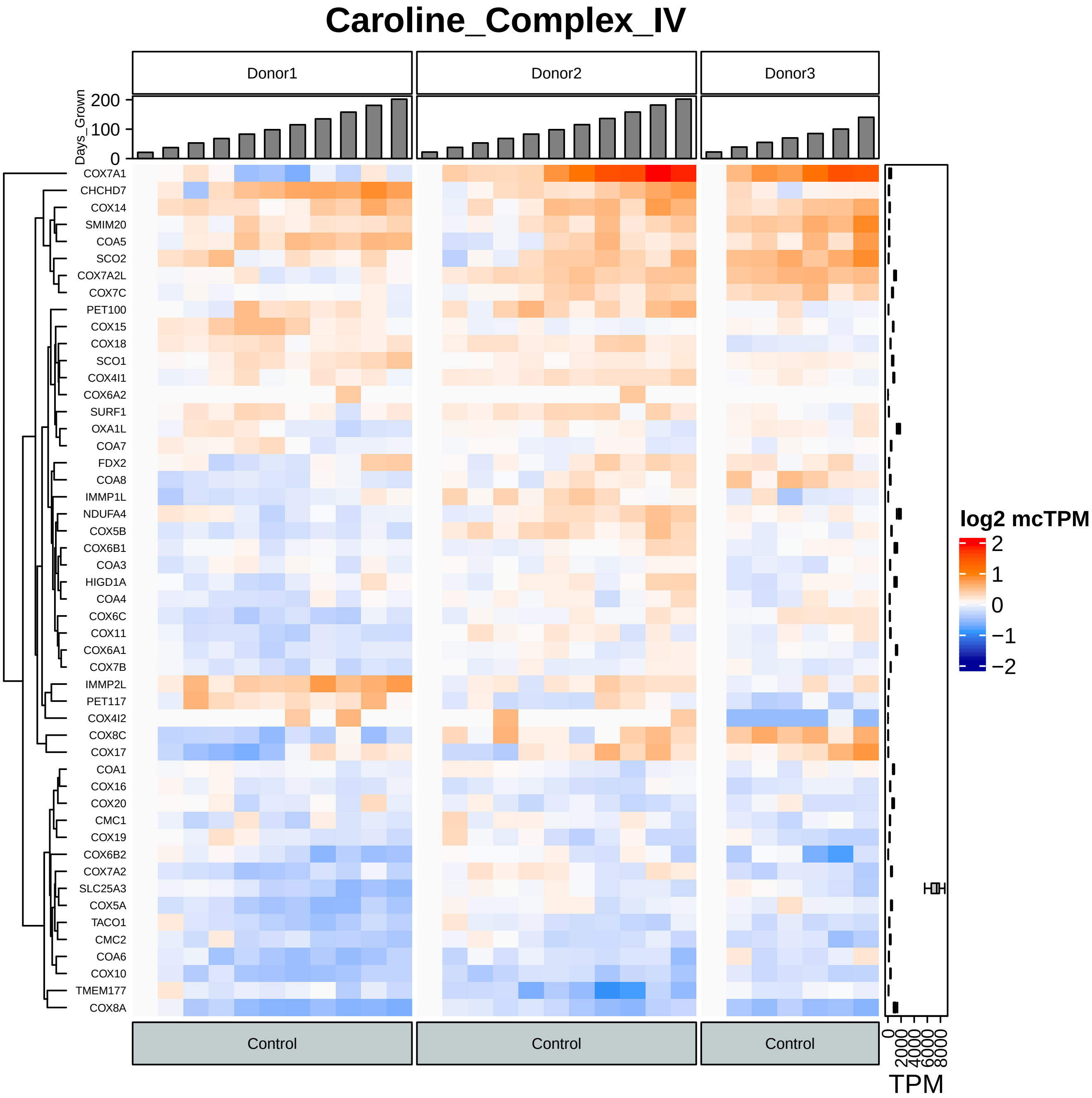

Caroline_Complex_IV
Donor1
Donor2
Donor3
Days_Grown
200
100
0
COX7A1
CHCHD7
COX14
SMIM20
COA5
SCO2
COX7A2L
COX7C
PET100
COX15
COX18
SCO1
COX4I1
COX6A2
SURF1
OXA1L
COA7
FDX2
COA8
IMMP1L
NDUFA4
COX5B
COX6B1
COA3
HIGD1A
COA4
COX6C
COX11
COX6A1
COX7B
IMMP2L
PET117
COX4I2
COX8C
COX17
COA1
COX16
COX20
CMC1
COX19
COX6B2
COX7A2
SLC25A3
COX5A
TACO1
CMC2
COA6
COX10
TMEM177
COX8A
log2 mcTPM
2
1
0
−1
−2
Control
Control
Control
0
2000
4000
6000
8000
TPM

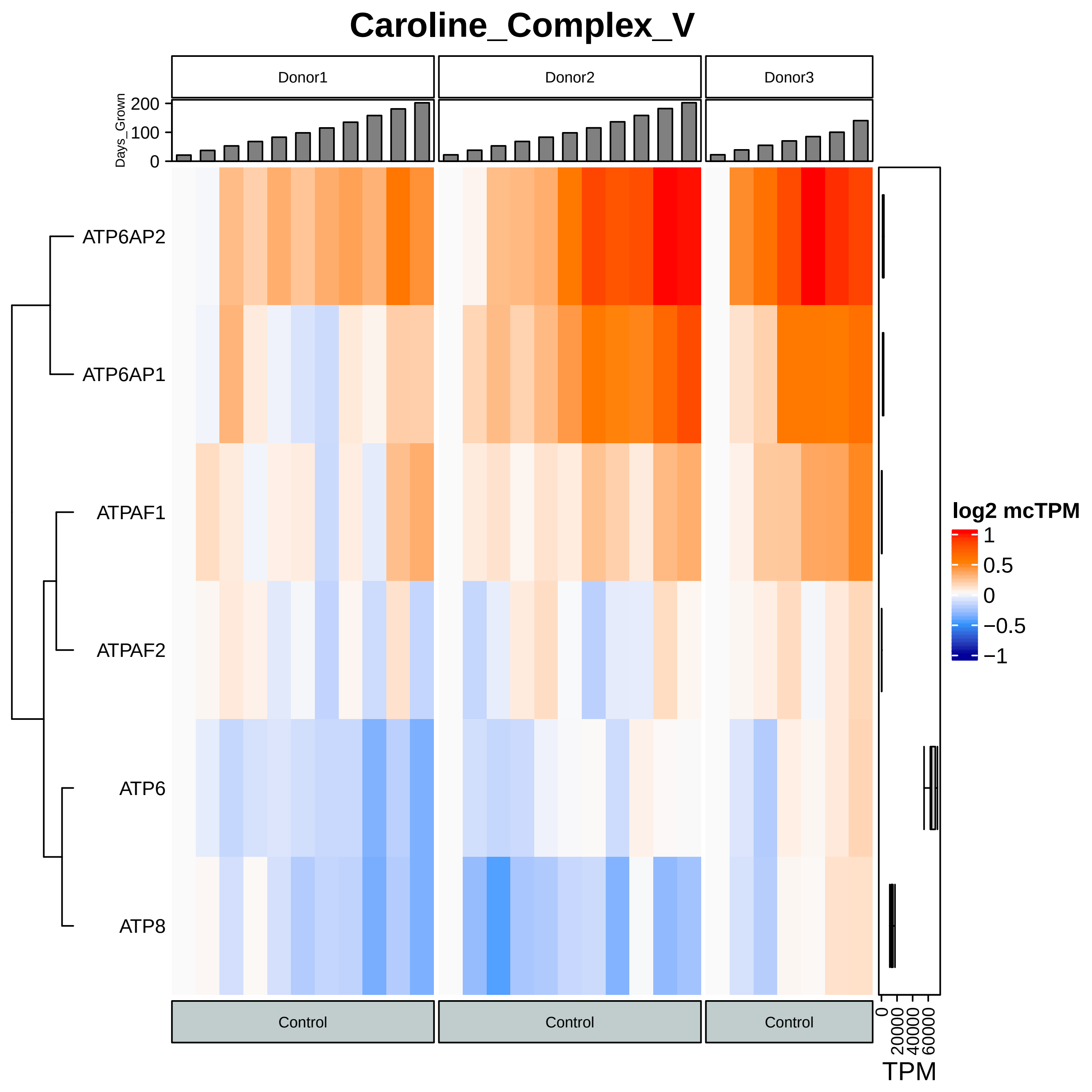

Caroline_Complex_V
Donor1
Donor2
Donor3
Days_Grown
200
100
0
ATP6AP2
ATP6AP1
ATPAF1
ATPAF2
ATP6
ATP8
log2 mcTPM
1
0.5
0
−0.5
−1
Control
Control
Control
0
20000
40000
60000
TPM

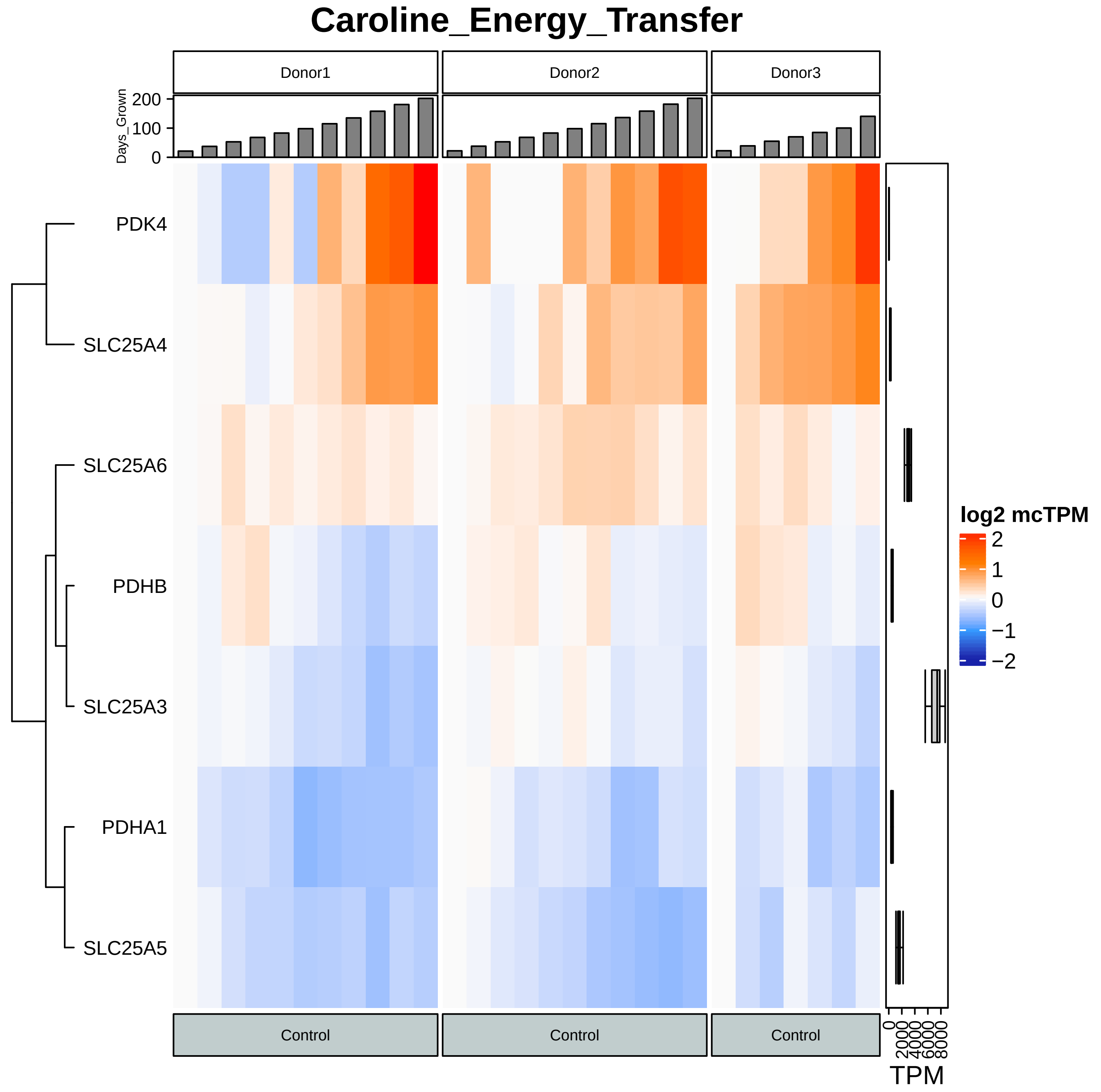
Caroline_Energy_Transfer
Donor1
Donor2
Donor3
Days_Grown
200
100
0
PDK4
SLC25A4
SLC25A6
PDHB
SLC25A3
PDHA1
SLC25A5
log2 mcTPM
2
1
0
−1
−2
Control
Control
Control
0
2000
4000
6000
8000
TPM

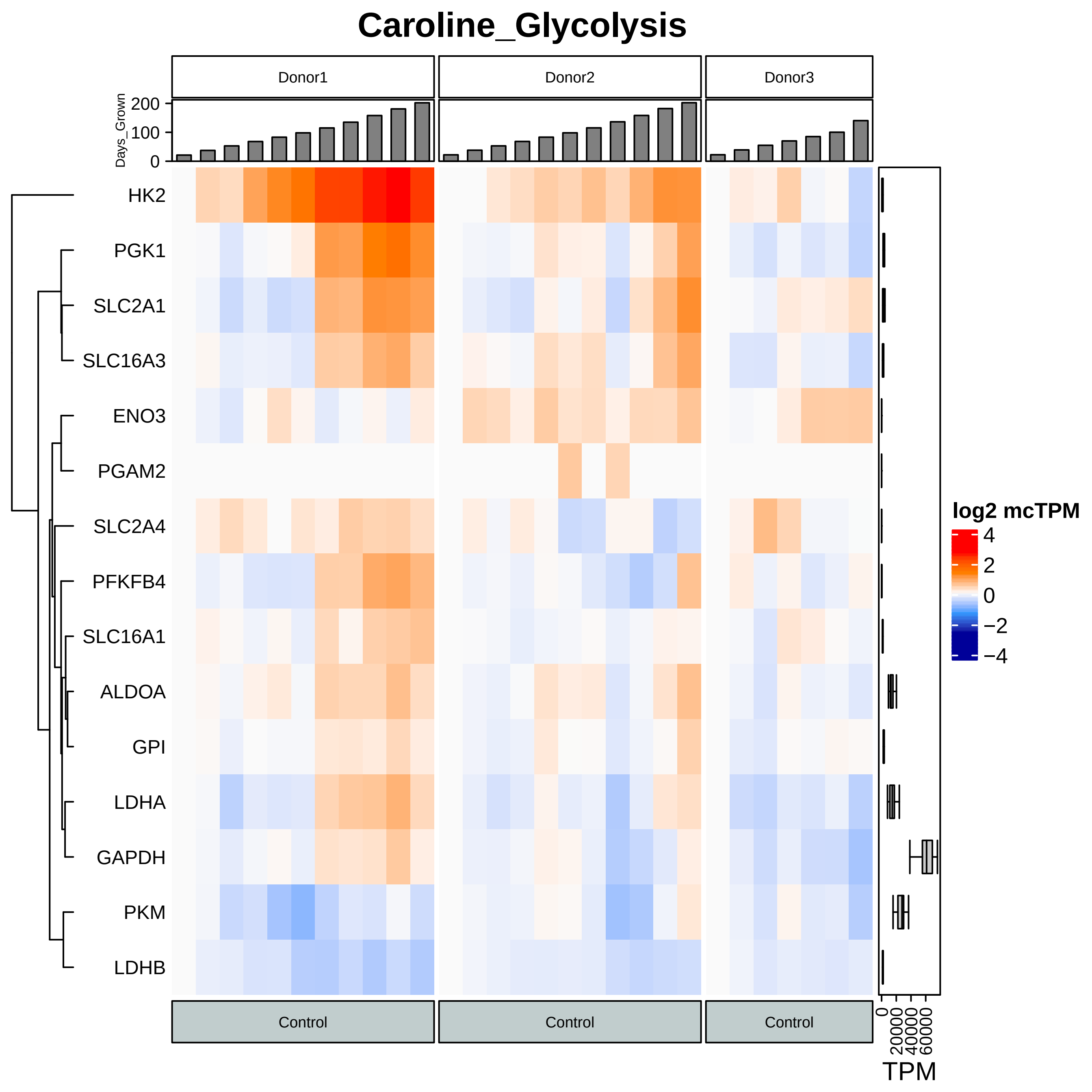

Caroline_Glycolysis
Donor1
Donor2
Donor3
Days_Grown
200
100
0
HK2
PGK1
SLC2A1
SLC16A3
ENO3
PGAM2
SLC2A4
PFKFB4
SLC16A1
ALDOA
GPI
LDHA
GAPDH
PKM
LDHB
log2 mcTPM
4
2
0
−2
−4
Control
Control
Control
0
20000
40000
60000
TPM

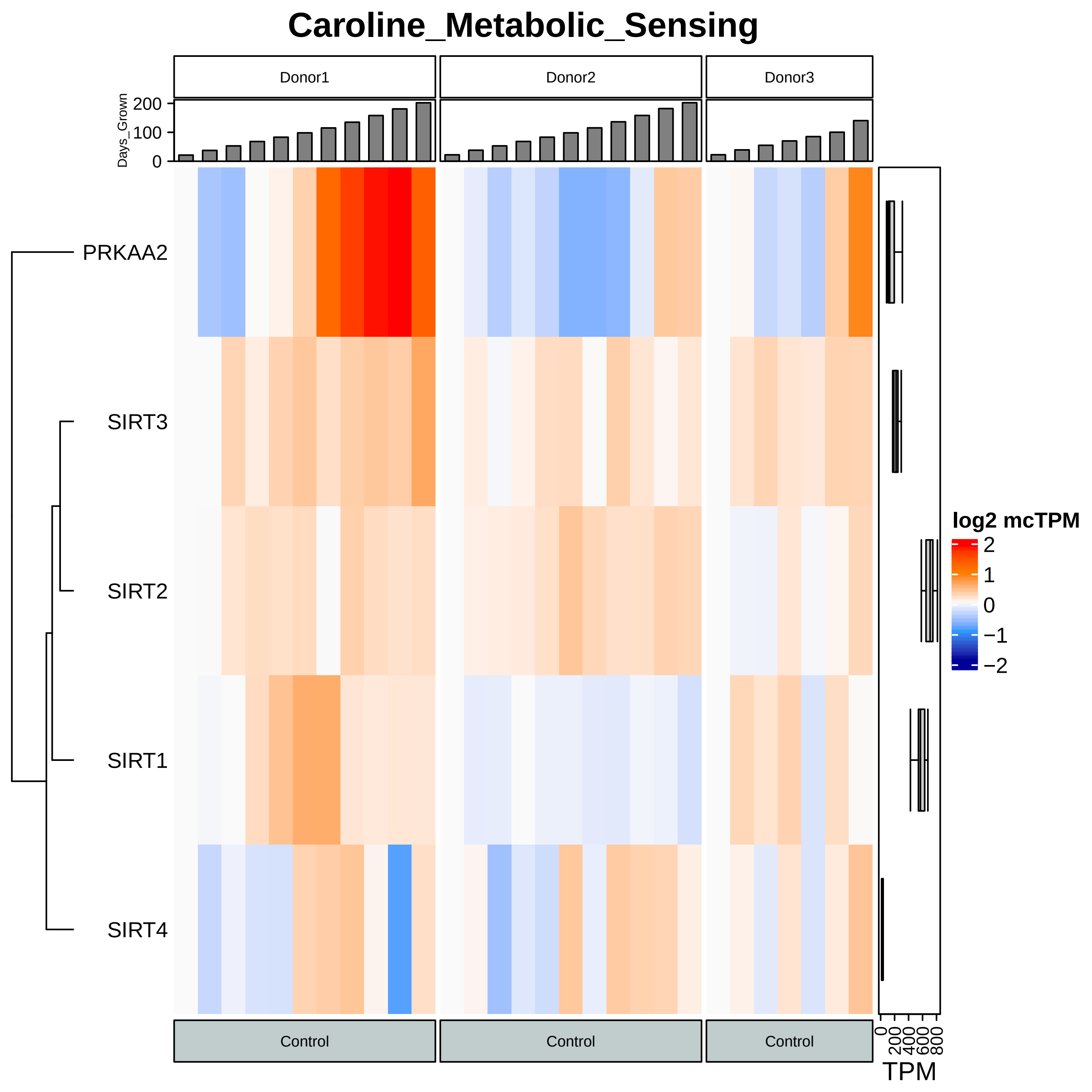

Caroline_Metabolic_Sensing
Donor1
Donor2
Donor3
Days_Grown
200
100
0
PRKAA2
SIRT3
SIRT2
SIRT1
SIRT4
Control
Control
Control
log2 mcTPM
2
1
0
−1
−2
0
200
400
600
800
TPM

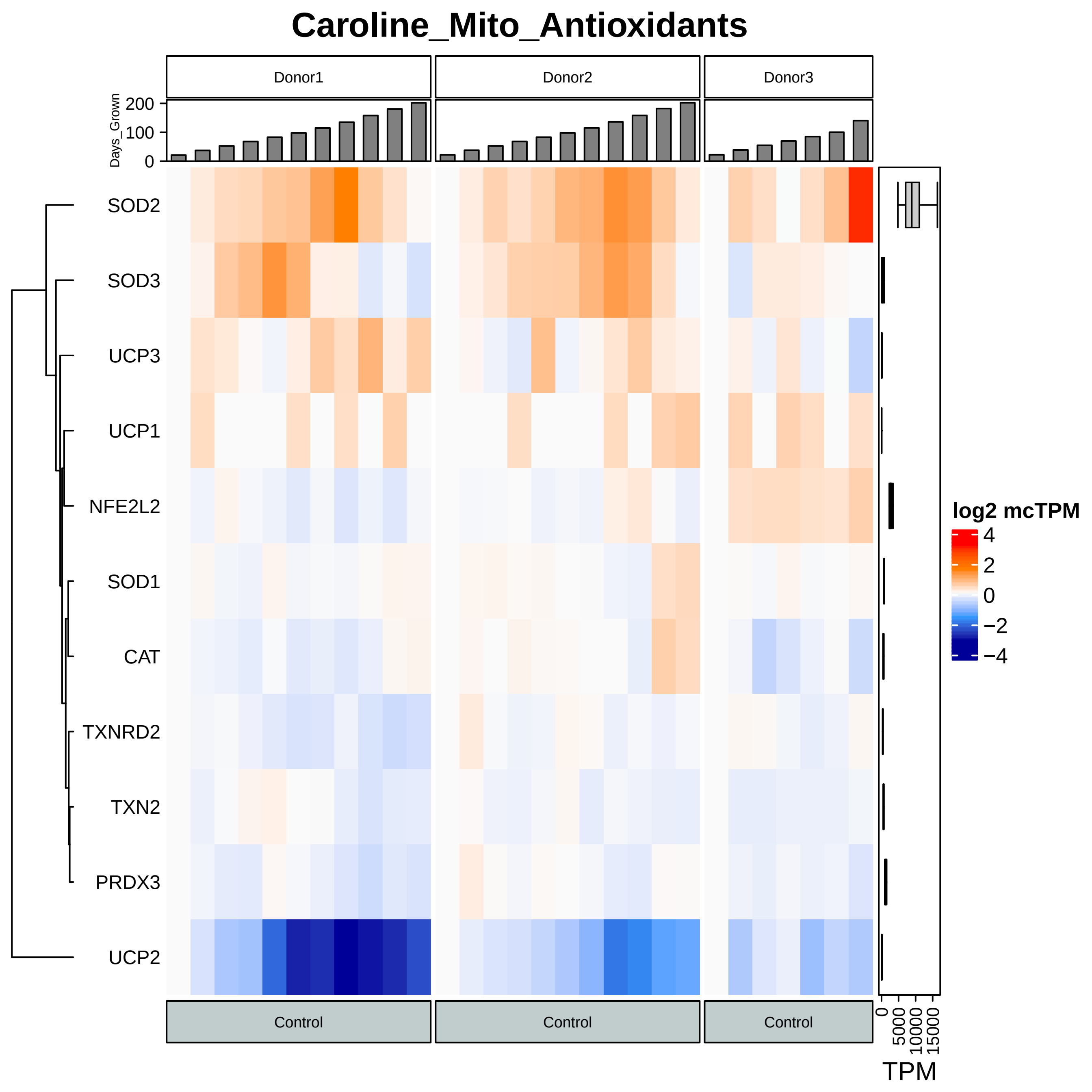

Caroline_Mito_Antioxidants
Donor1
Donor2
Donor3
Days_Grown
0
100
200
SOD2
SOD3
UCP3
UCP1
NFE2L2
SOD1
CAT
TXNRD2
TXN2
PRDX3
UCP2
Control
Control
Control
log2 mcTPM
4
2
0
−2
−4
0
5000
10000
15000
TPM

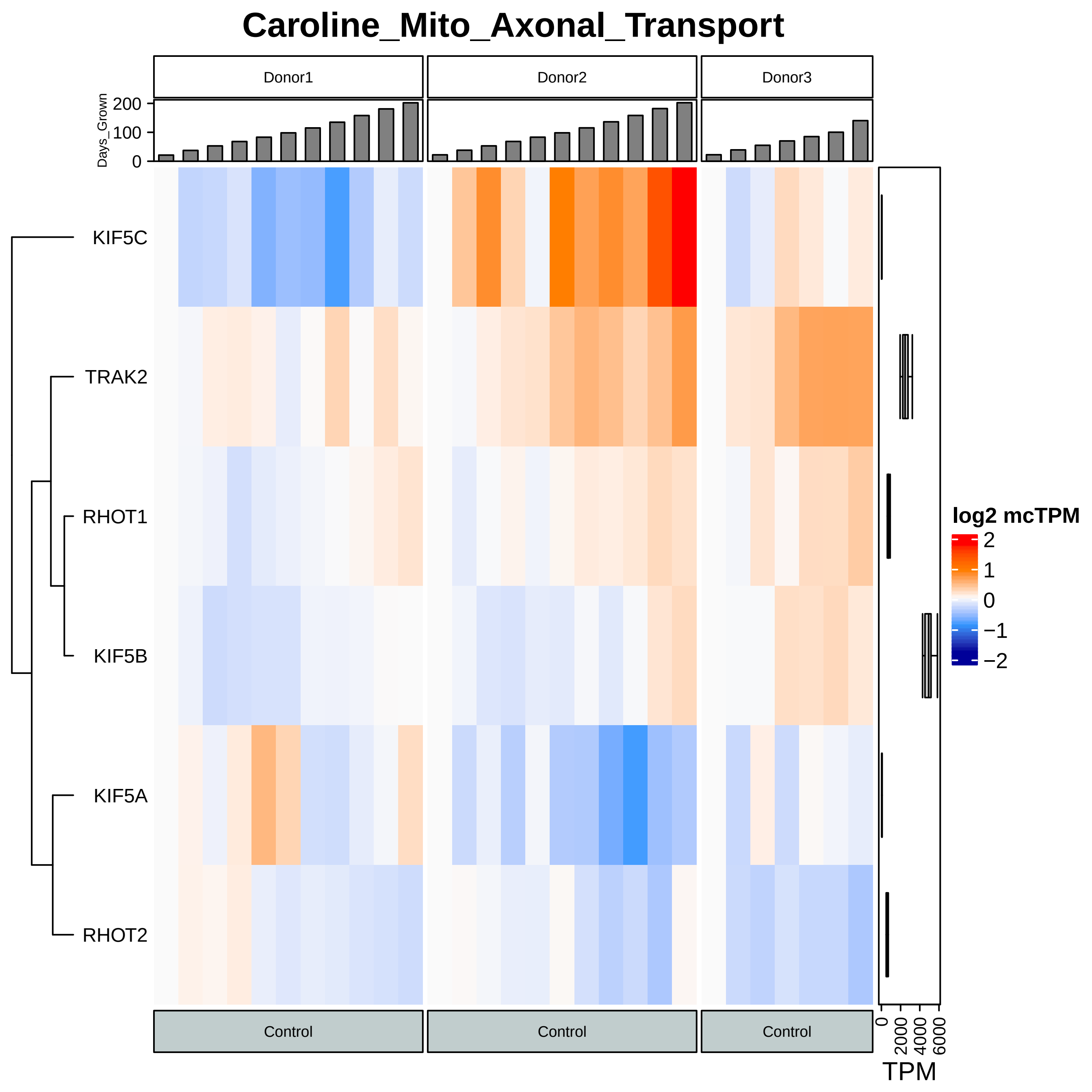

Caroline_Mito_Axonal_Transport
Donor1
Donor2
Donor3
Days_Grown
200
100
0
KIF5C
TRAK2
RHOT1
KIF5B
KIF5A
RHOT2
Control
Control
Control
log2 mcTPM
2
1
0
−1
−2
0
2000
4000
6000
TPM

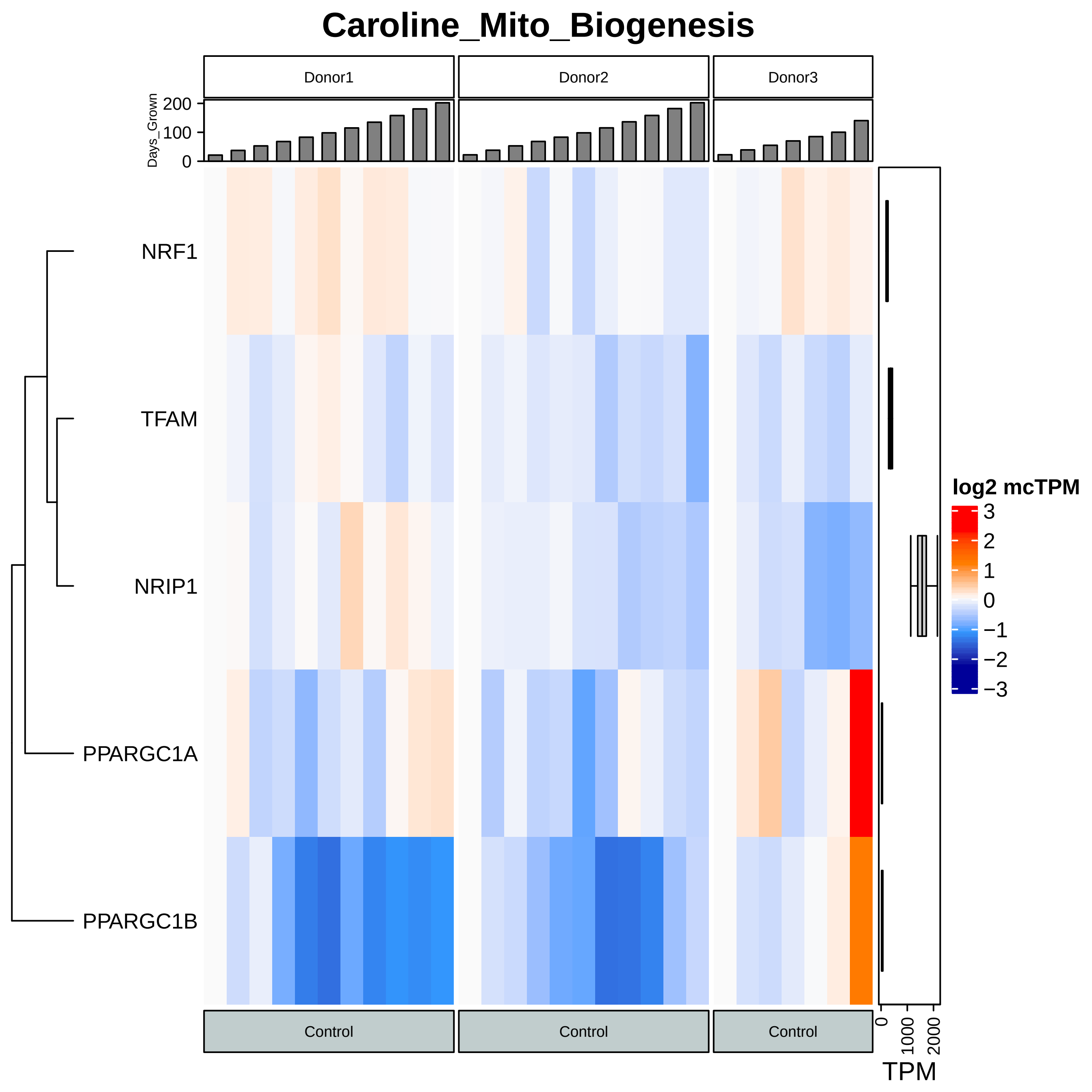

Caroline_Mito_Biogenesis
Donor1
Donor2
Donor3
Days_Grown
200
100
0
NRF1
TFAM
NRIP1
PPARGC1A
PPARGC1B
Control
Control
Control
0
1000
2000
TPM
log2 mcTPM
3
2
1
0
−1
−2
−3

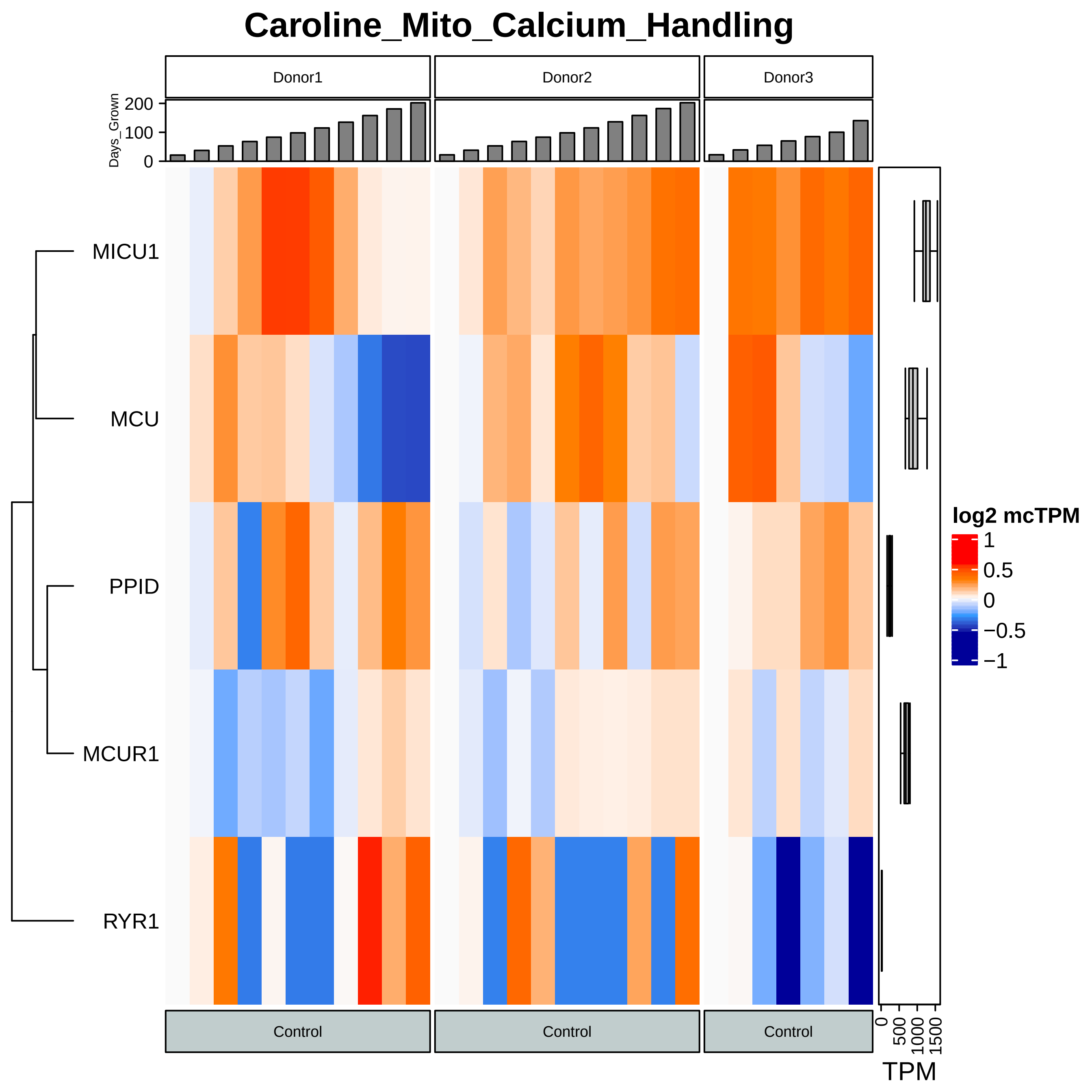
Caroline_Mito_Calcium_Handling
Donor1
Donor2
Donor3
Days_Grown
200
100
0
MICU1
MCU
PPID
MCUR1
RYR1
log2 mcTPM
1
0.5
0
−0.5
−1
Control
Control
Control
0
500
1000
1500
TPM

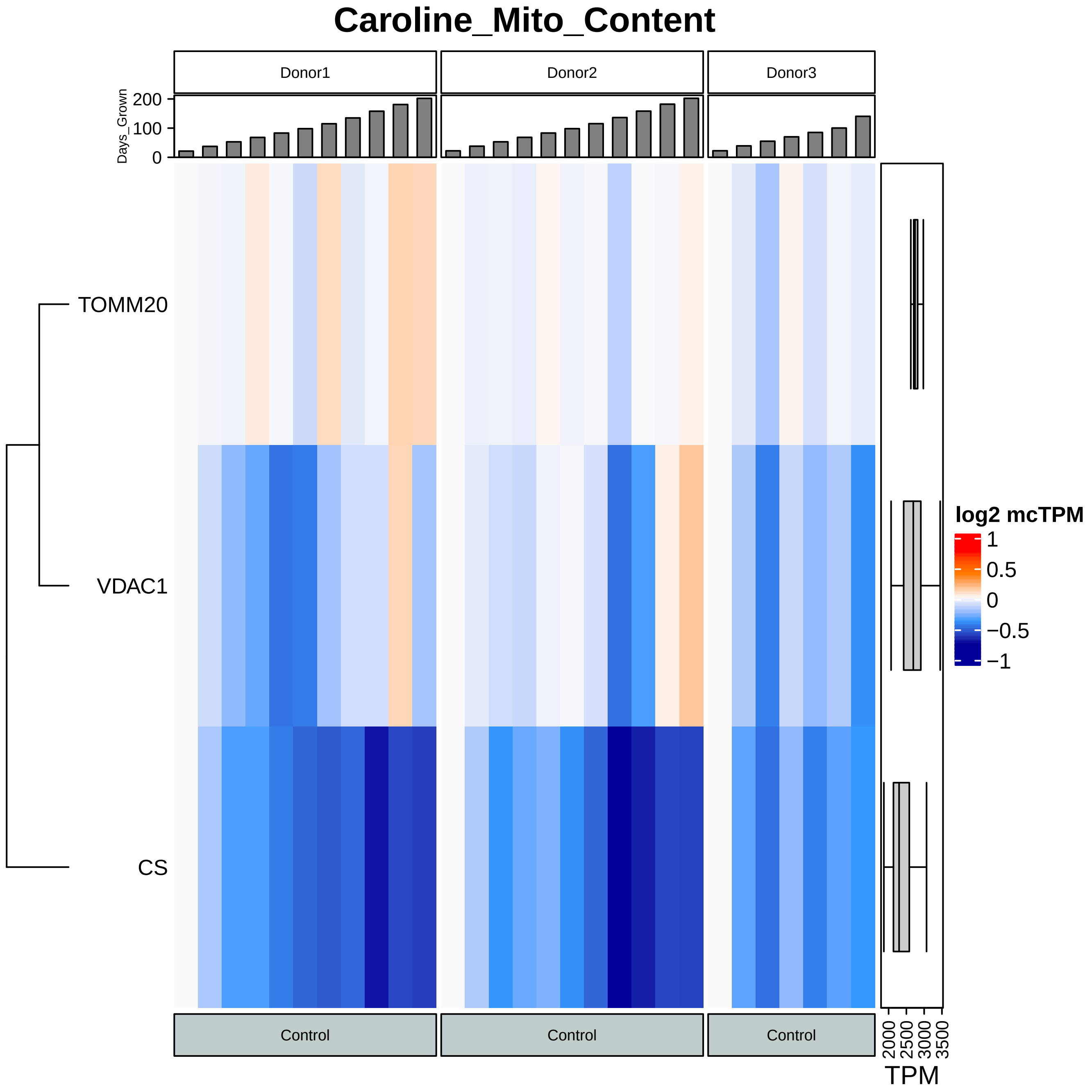

Caroline_Mito_Content
Donor1
Donor2
Donor3
Days_Grown
200
100
0
TOMM20
VDAC1
CS
log2 mcTPM
1
0.5
0
−0.5
−1
Control
Control
Control
2000
2500
3000
3500
TPM

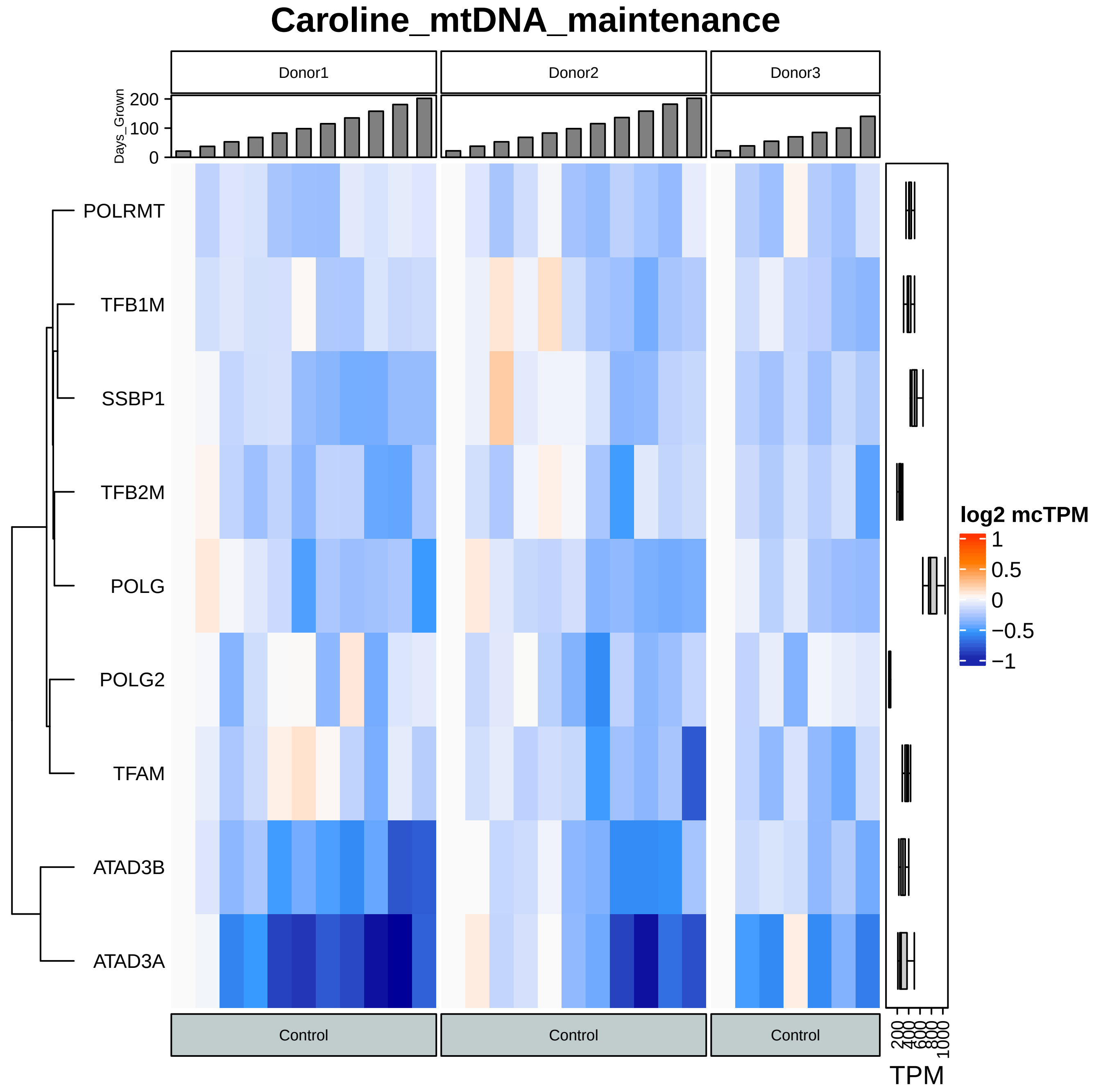

Caroline_mtDNA_maintenance
Donor1
Donor2
Donor3
Days_Grown
200
100
0
POLRMT
TFB1M
SSBP1
TFB2M
POLG
POLG2
TFAM
ATAD3B
ATAD3A
Control
Control
Control
log2 mcTPM
1
0.5
0
−0.5
−1
200
400
600
800
1000
TPM

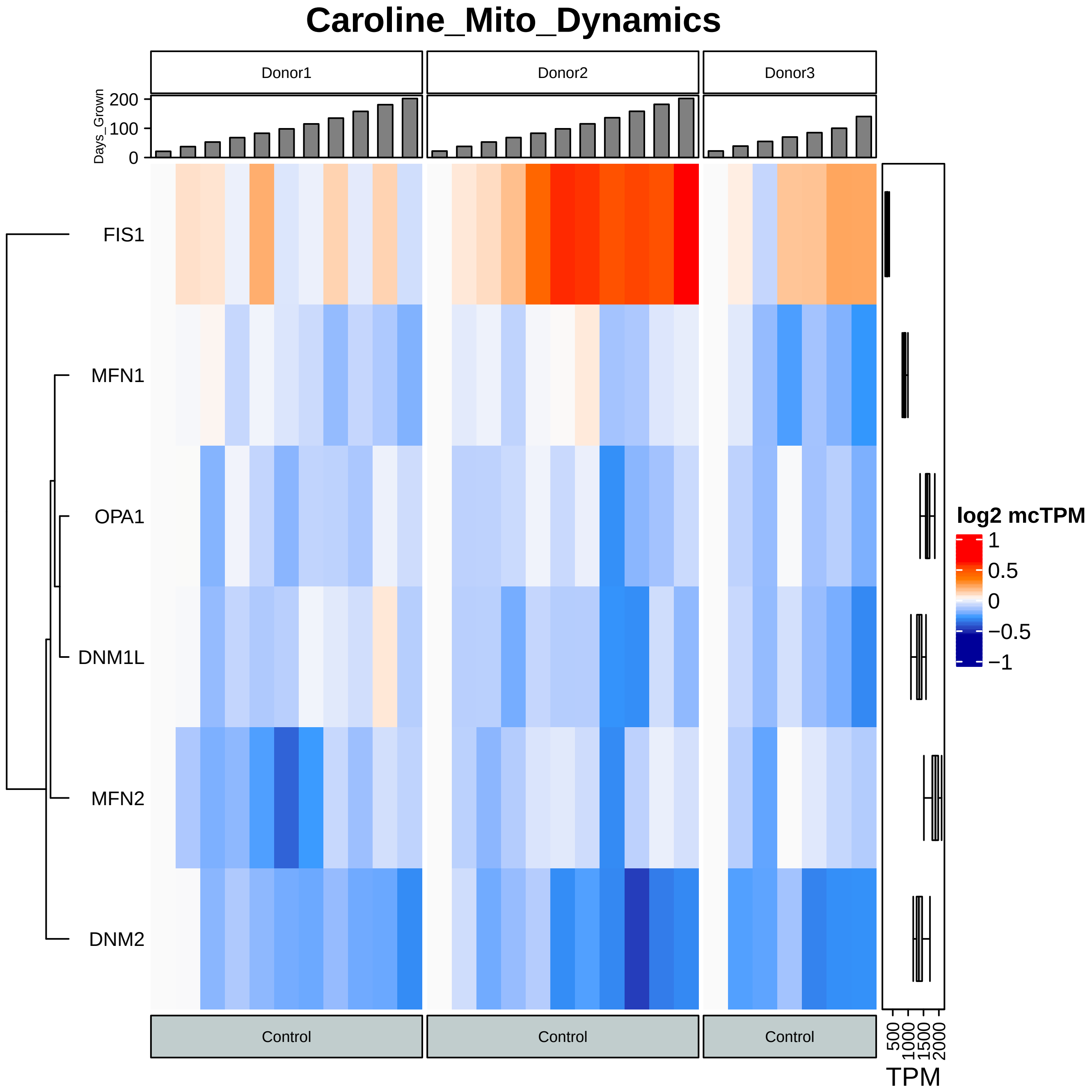

Caroline_Mito_Dynamics
Donor1
Donor2
Donor3
Days_Grown
200
100
0
FIS1
MFN1
OPA1
DNM1L
MFN2
DNM2
Control
Control
Control
log2 mcTPM
1
0.5
0
−0.5
−1
500
1000
1500
2000
TPM

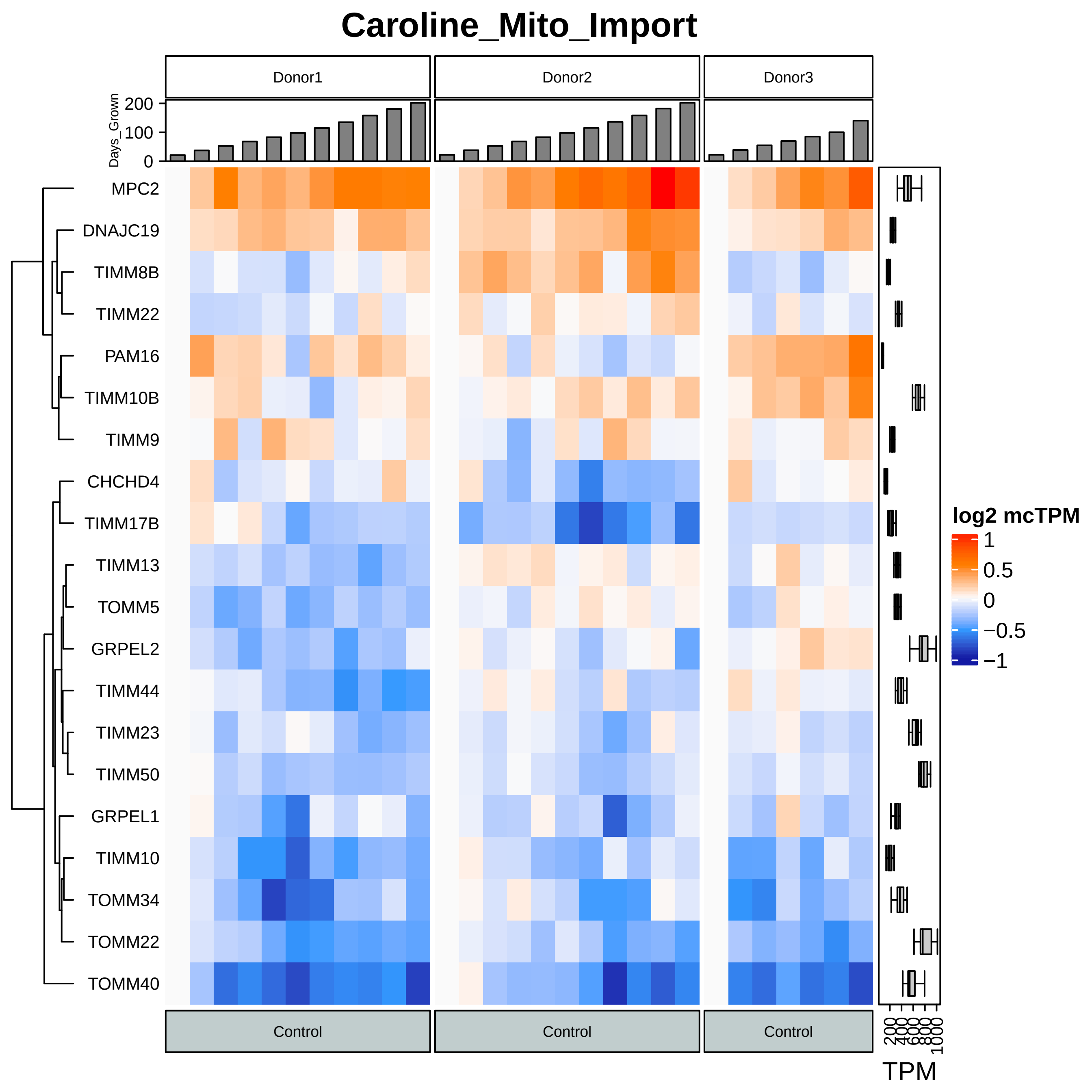

Caroline_Mito_Import
Donor1
Donor2
Donor3
Days_Grown
200
100
0
MPC2
DNAJC19
TIMM8B
TIMM22
PAM16
TIMM10B
TIMM9
CHCHD4
TIMM17B
TIMM13
TOMM5
GRPEL2
TIMM44
TIMM23
TIMM50
GRPEL1
TIMM10
TOMM34
TOMM22
TOMM40
Control
Control
Control
log2 mcTPM
1
0.5
0
−0.5
−1
200
400
600
800
1000
TPM

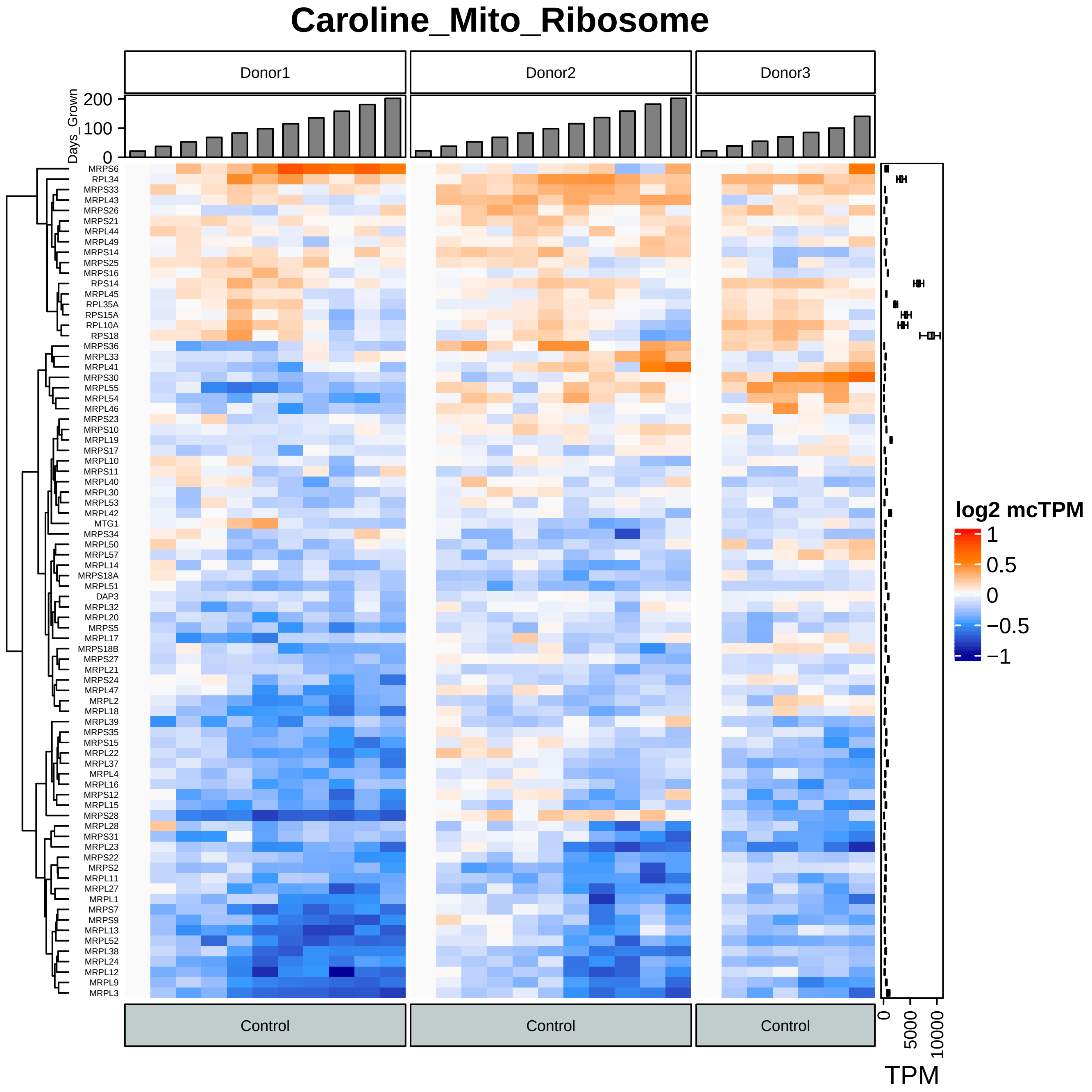

Caroline_Mito_Ribosome
Donor1
Donor2
Donor3
Days_Grown
200
100
0
MRPS6
RPL34
MRPS33
MRPL43
MRPS26
MRPS21
MRPL44
MRPL49
MRPS14
MRPS25
MRPS16
RPS14
MRPL45
RPL35A
RPS15A
RPL10A
RPS18
MRPS36
MRPL33
MRPL41
MRPS30
MRPL55
MRPL54
MRPL46
MRPS23
MRPS10
MRPL19
MRPS17
MRPL10
MRPS11
MRPL40
MRPL30
MRPL53
MRPL42
MTG1
MRPS34
MRPL50
MRPL57
MRPL14
MRPS18A
MRPL51
DAP3
MRPL32
MRPL20
MRPS5
MRPL17
MRPS18B
MRPS27
MRPL21
MRPS24
MRPL47
MRPL2
MRPL18
MRPL39
MRPS35
MRPS15
MRPL22
MRPL37
MRPL4
MRPL16
MRPS12
MRPL15
MRPS28
MRPL28
MRPS31
MRPL23
MRPS22
MRPS2
MRPL11
MRPL27
MRPL1
MRPS7
MRPS9
MRPL13
MRPL52
MRPL38
MRPL24
MRPL12
MRPL9
MRPL3
Control
Control
Control
0
5000
10000
TPM
log2 mcTPM
1
0.5
0
−0.5
−1

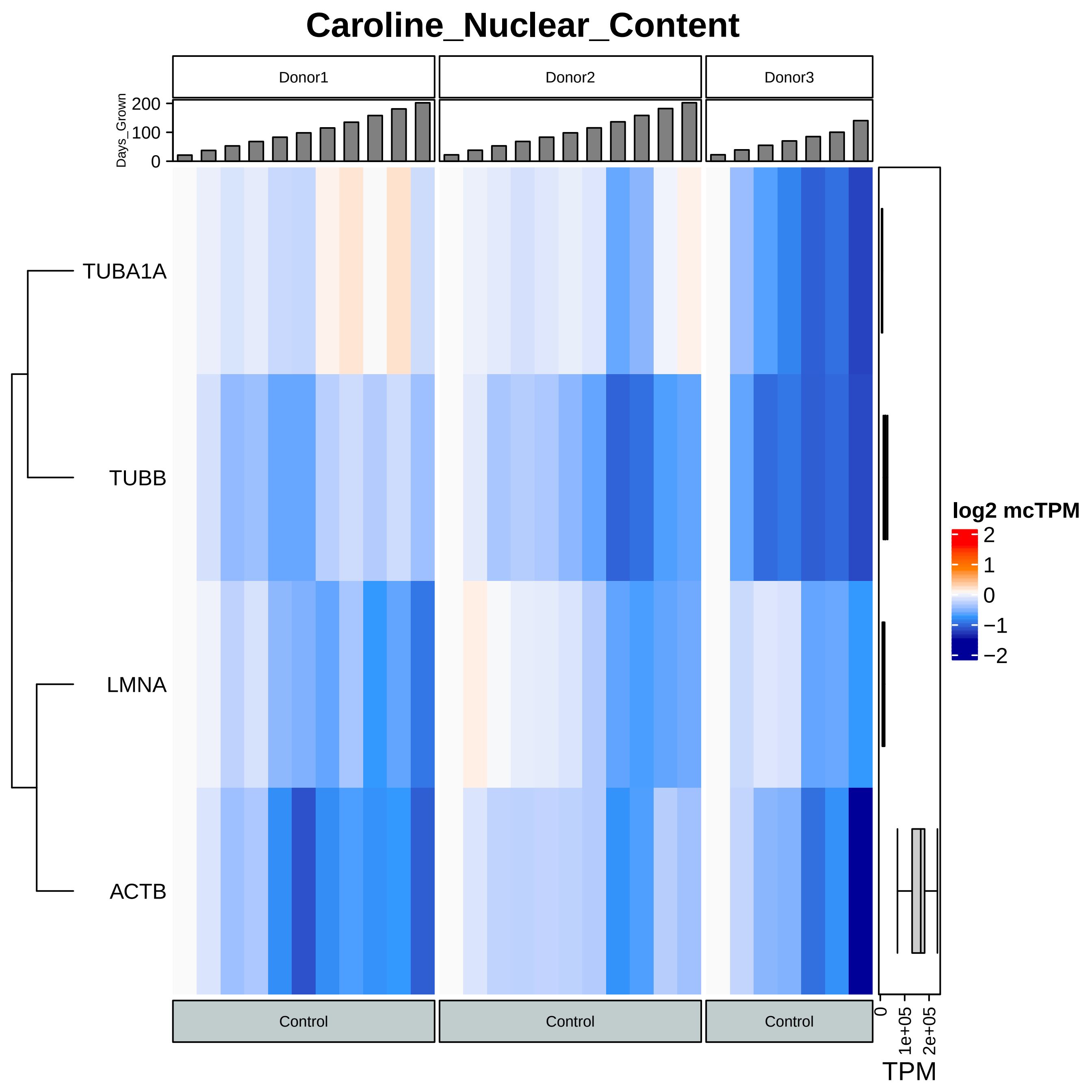
Caroline_Nuclear_Content
Donor1
Donor2
Donor3
Days_Grown
200
100
0
TUBA1A
TUBB
LMNA
ACTB
Control
Control
Control
0
1e+05
2e+05
TPM
log2 mcTPM
2
1
0
−1
−2

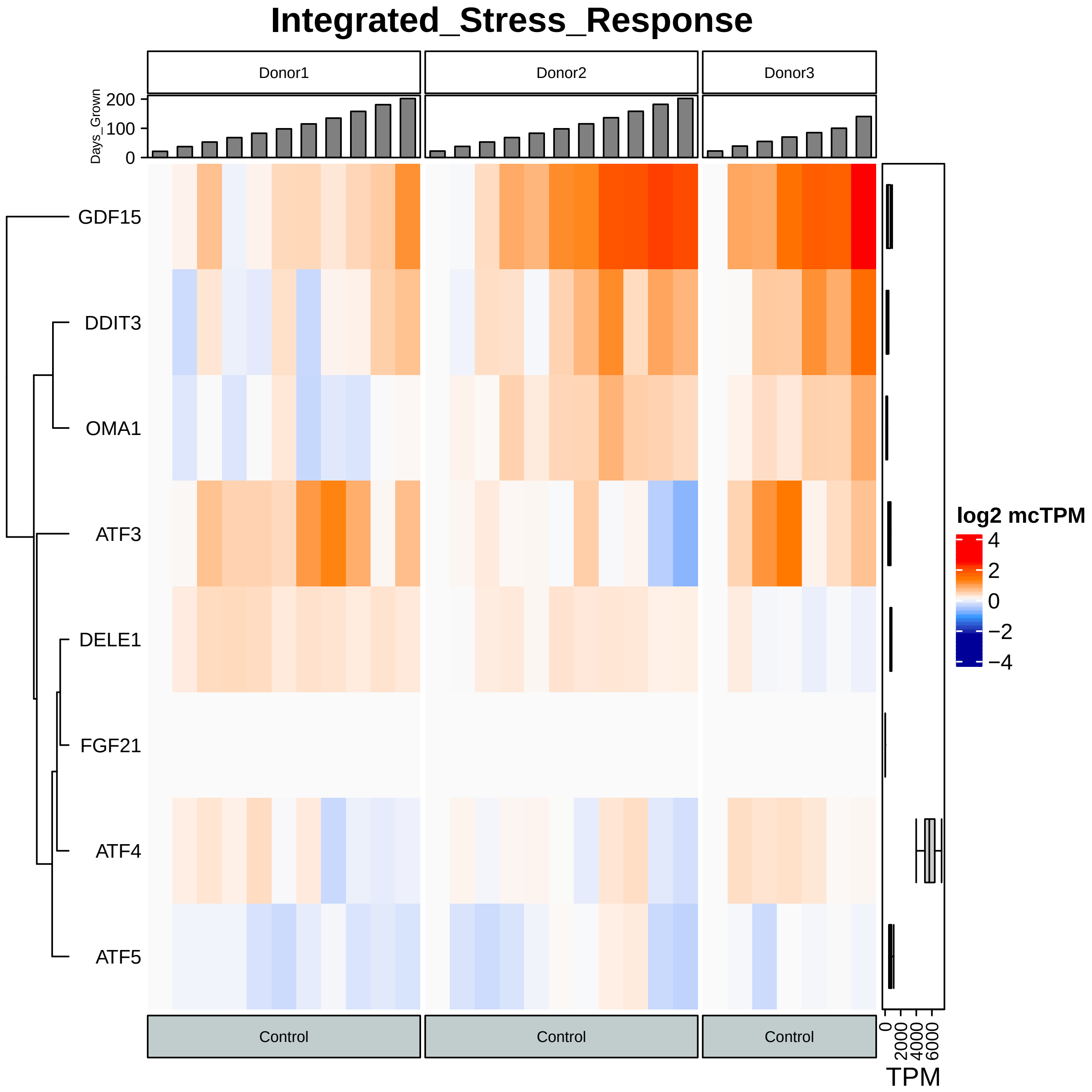

Integrated_Stress_Response
Donor1
Donor2
Donor3
Days_Grown
200
100
0
GDF15
DDIT3
OMA1
ATF3
DELE1
FGF21
ATF4
ATF5
Control
Control
Control
log2 mcTPM
4
2
0
−2
−4
0
2000
4000
6000
TPM

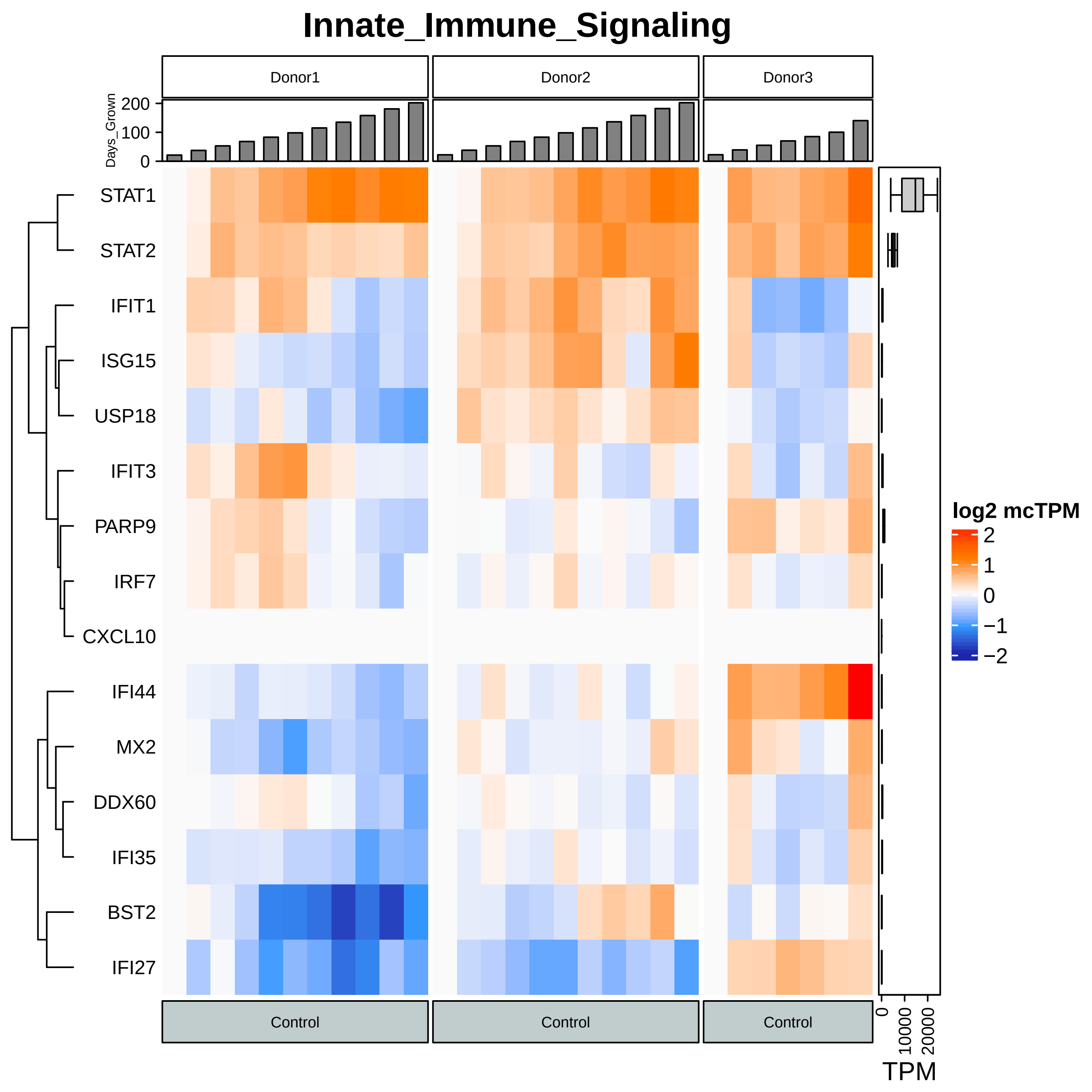
Innate_Immune_Signaling
Donor1
Donor2
Donor3
Days_Grown
200
100
0
STAT1
STAT2
IFIT1
ISG15
USP18
IFIT3
PARP9
IRF7
CXCL10
IFI44
MX2
DDX60
IFI35
BST2
IFI27
Control
Control
Control
0
10000
20000
TPM
log2 mcTPM
2
1
0
−1
−2

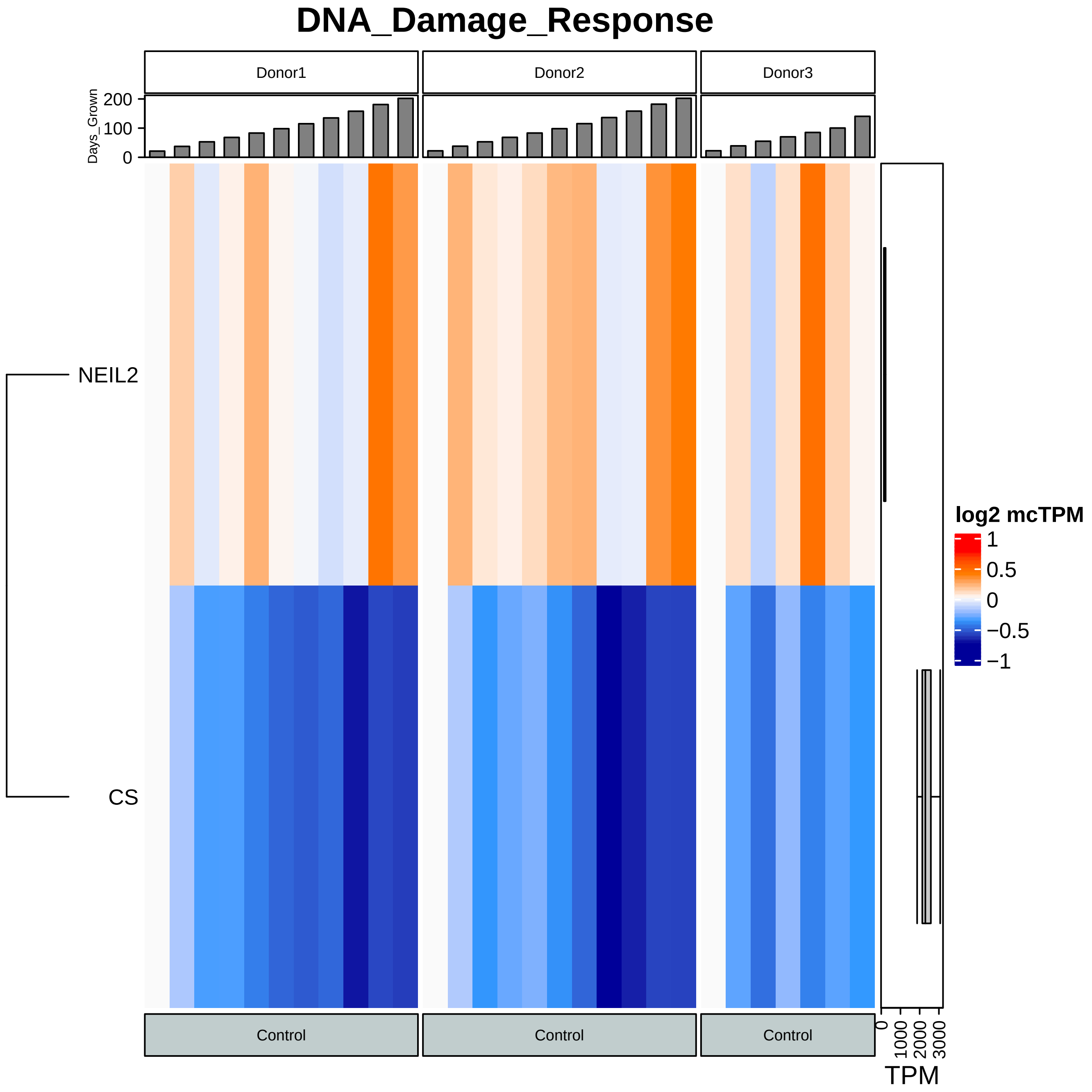

DNA_Damage_Response
Donor1
Donor2
Donor3
Days_Grown
200
100
0
NEIL2
CS
log2 mcTPM
1
0.5
0
−0.5
−1
Control
Control
Control
0
1000
2000
3000
TPM

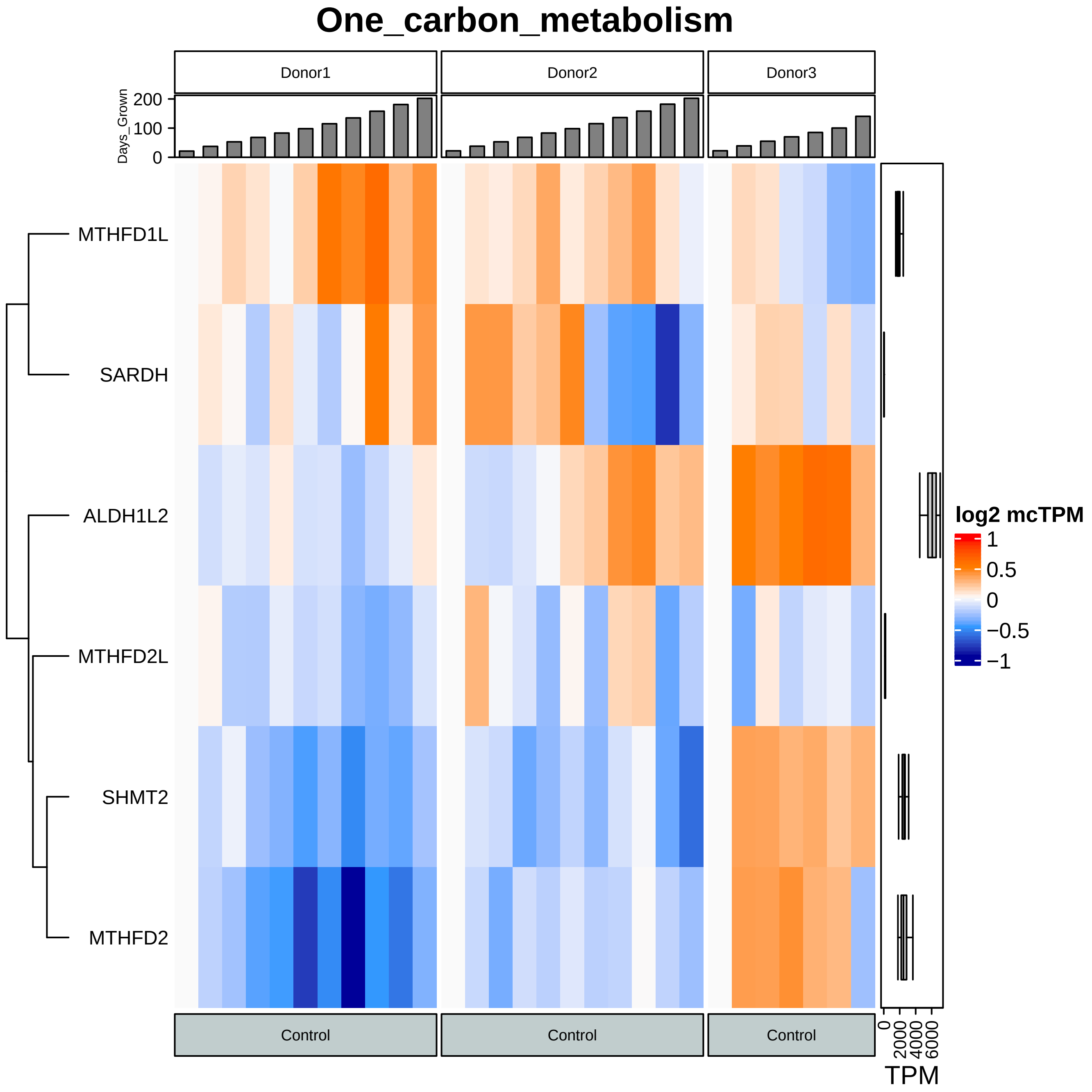

One_carbon_metabolism
Donor1
Donor2
Donor3
Days_Grown
200
100
0
MTHFD1L
SARDH
ALDH1L2
MTHFD2L
SHMT2
MTHFD2
log2 mcTPM
1
0.5
0
−0.5
−1
Control
Control
Control
0
2000
4000
6000
TPM

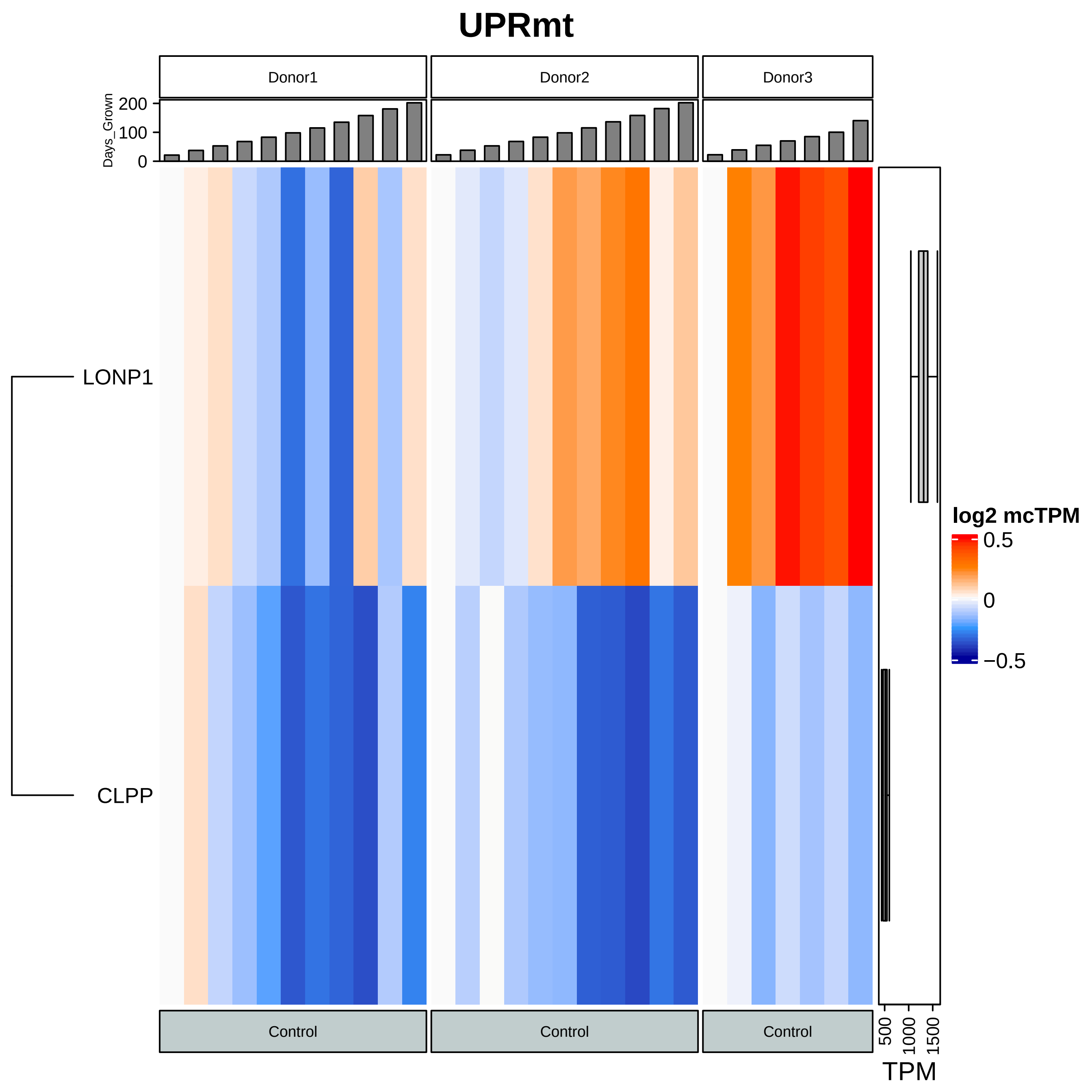
UPRmt
Donor1
Donor2
Donor3
Days_Grown
0
100
200
LONP1
CLPP
log2 mcTPM
0.5
0
−0.5
Control
Control
Control
500
1000
1500
TPM

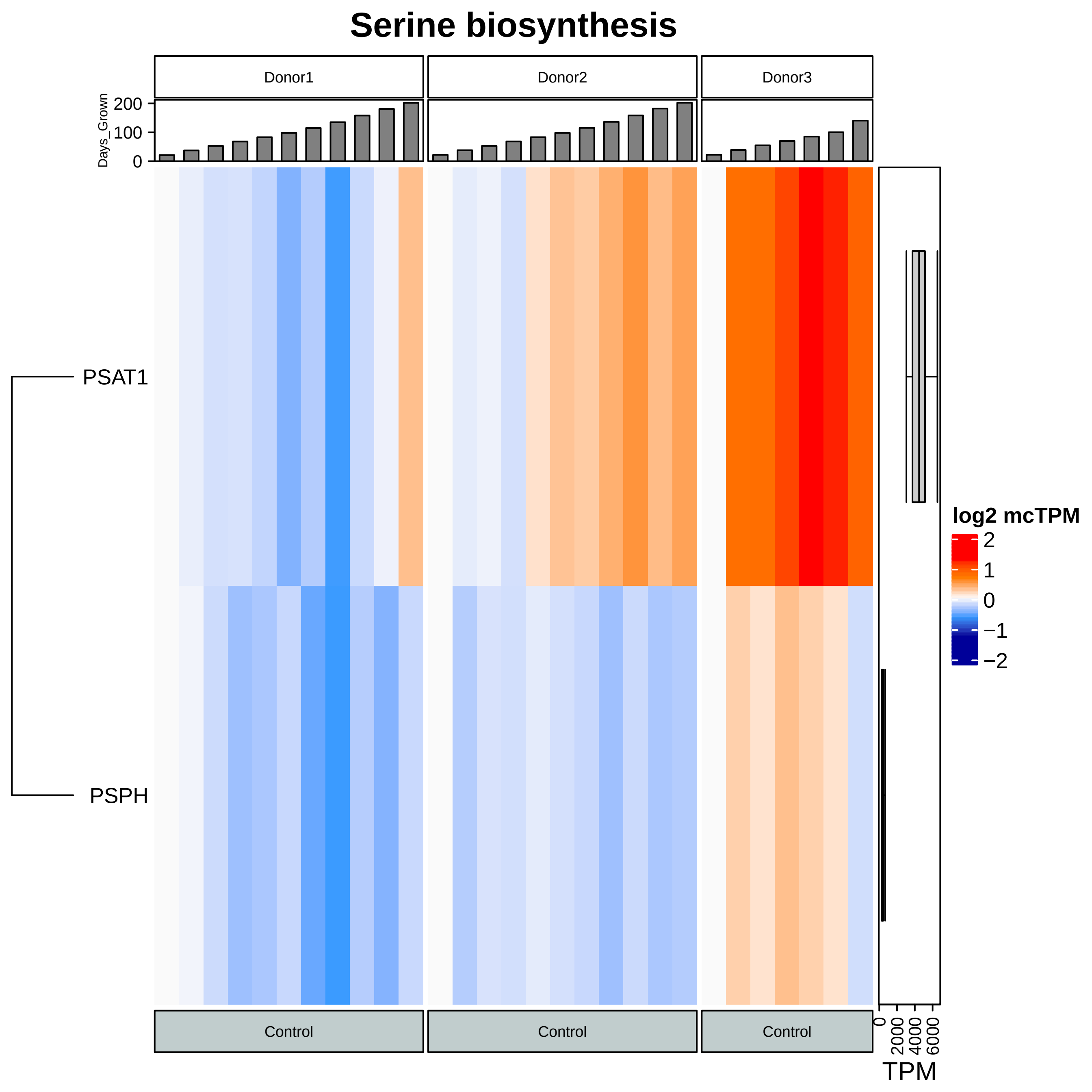
Serine biosynthesis
Donor1
Donor2
Donor3
Days_Grown
200
100
0
PSAT1
PSPH
log2 mcTPM
2
1
0
−1
−2
Control
Control
Control
0
2000
4000
6000
TPM

### Transsulfuration

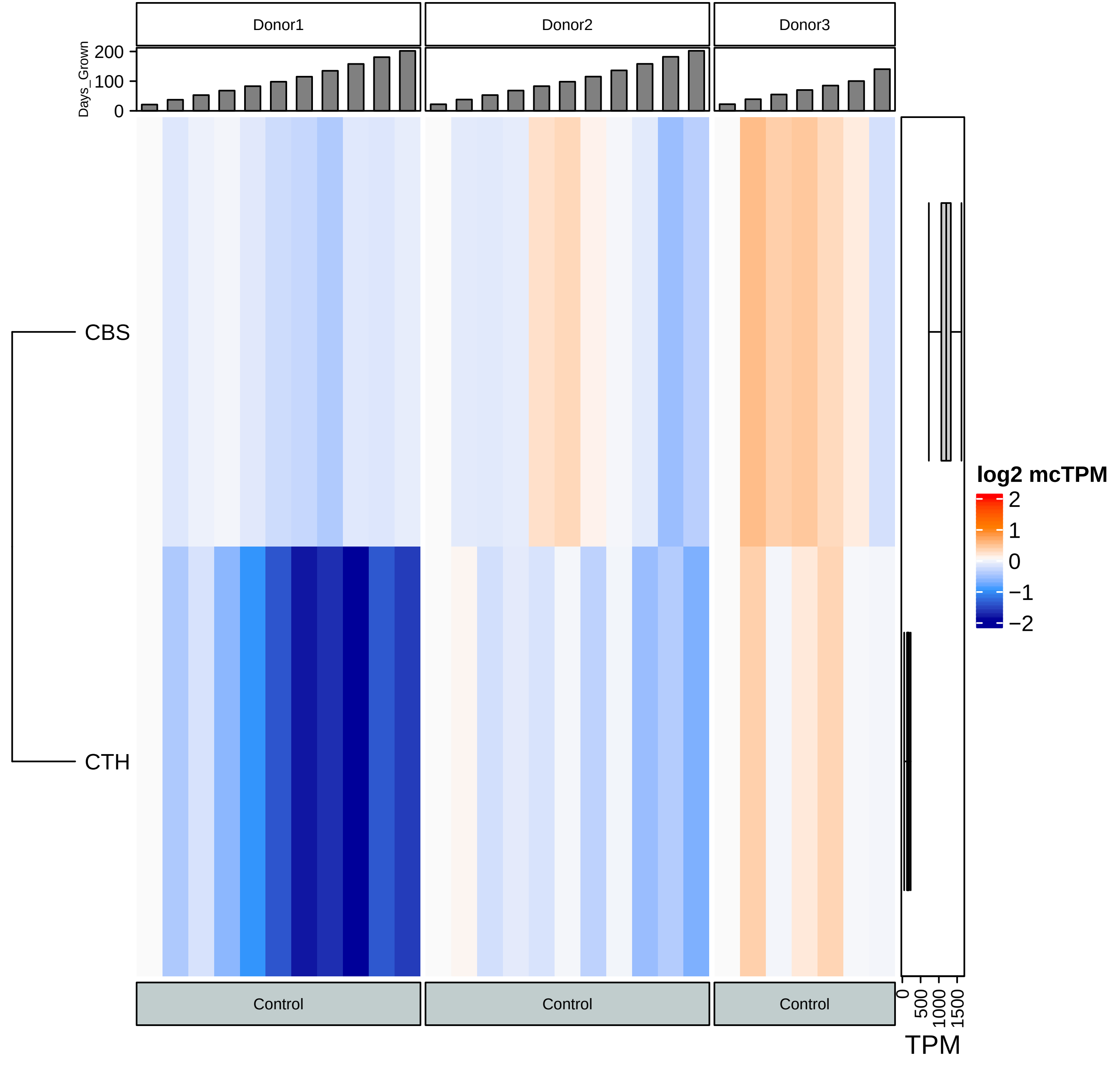

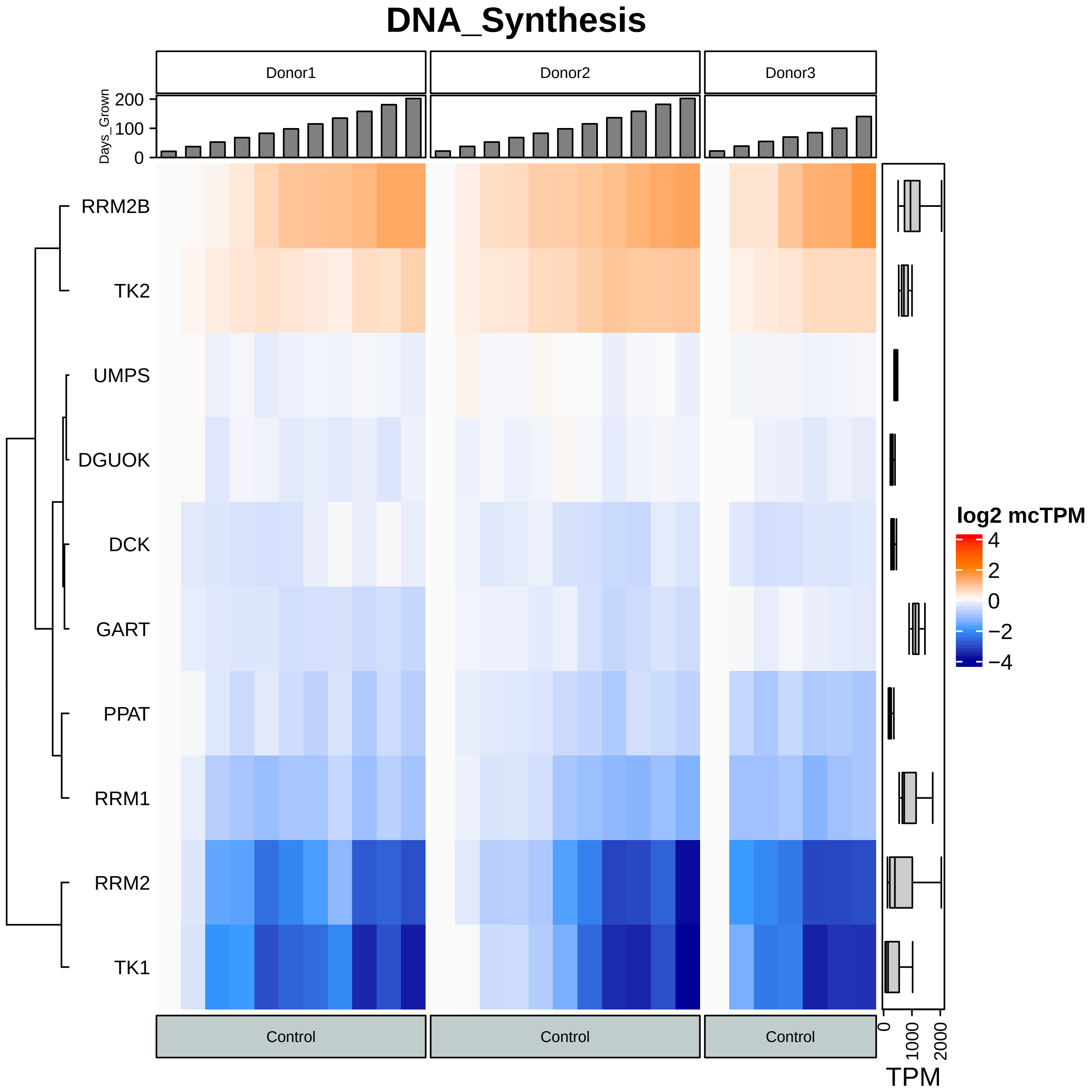
DNA_Synthesis
Donor1
Donor2
Donor3
Days_Grown
200
100
0
RRM2B
TK2
UMPS
DGUOK
DCK
GART
PPAT
RRM1
RRM2
TK1
log2 mcTPM
4
2
0
−2
−4
Control
Control
Control
0
1000
2000
TPM

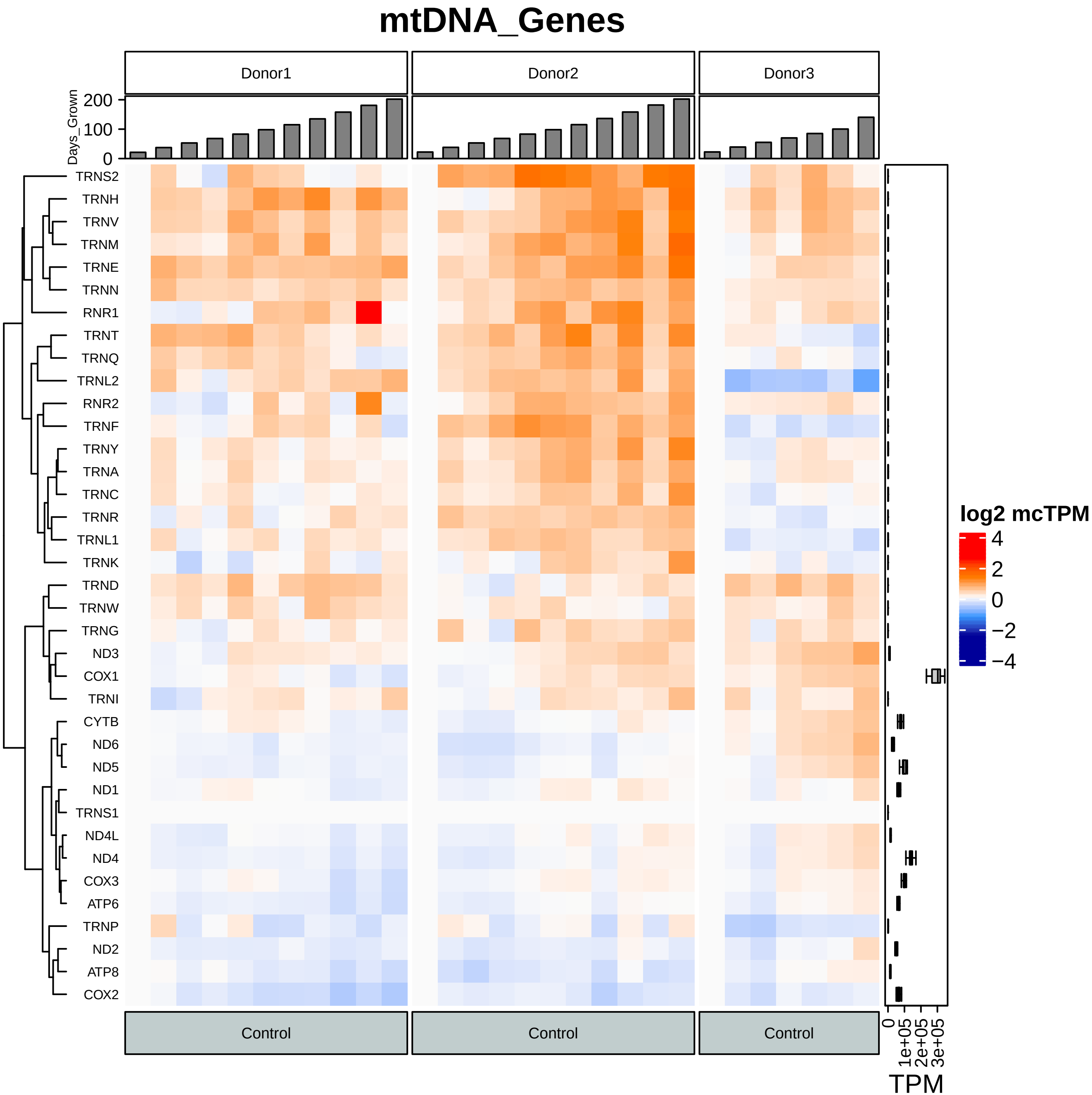

mtDNA_Genes
Donor1
Donor2
Donor3
Days_Grown
200
100
0
TRNS2
TRNH
TRNV
TRNM
TRNE
TRNN
RNR1
TRNT
TRNQ
TRNL2
RNR2
TRNF
TRNY
TRNA
TRNC
TRNR
TRNL1
TRNK
TRND
TRNW
TRNG
ND3
COX1
TRNI
CYTB
ND6
ND5
ND1
TRNS1
ND4L
ND4
COX3
ATP6
TRNP
ND2
ATP8
COX2
log2 mcTPM
4
2
0
−2
−4
Control
Control
Control
0
1e+05
2e+05
3e+05
TPM

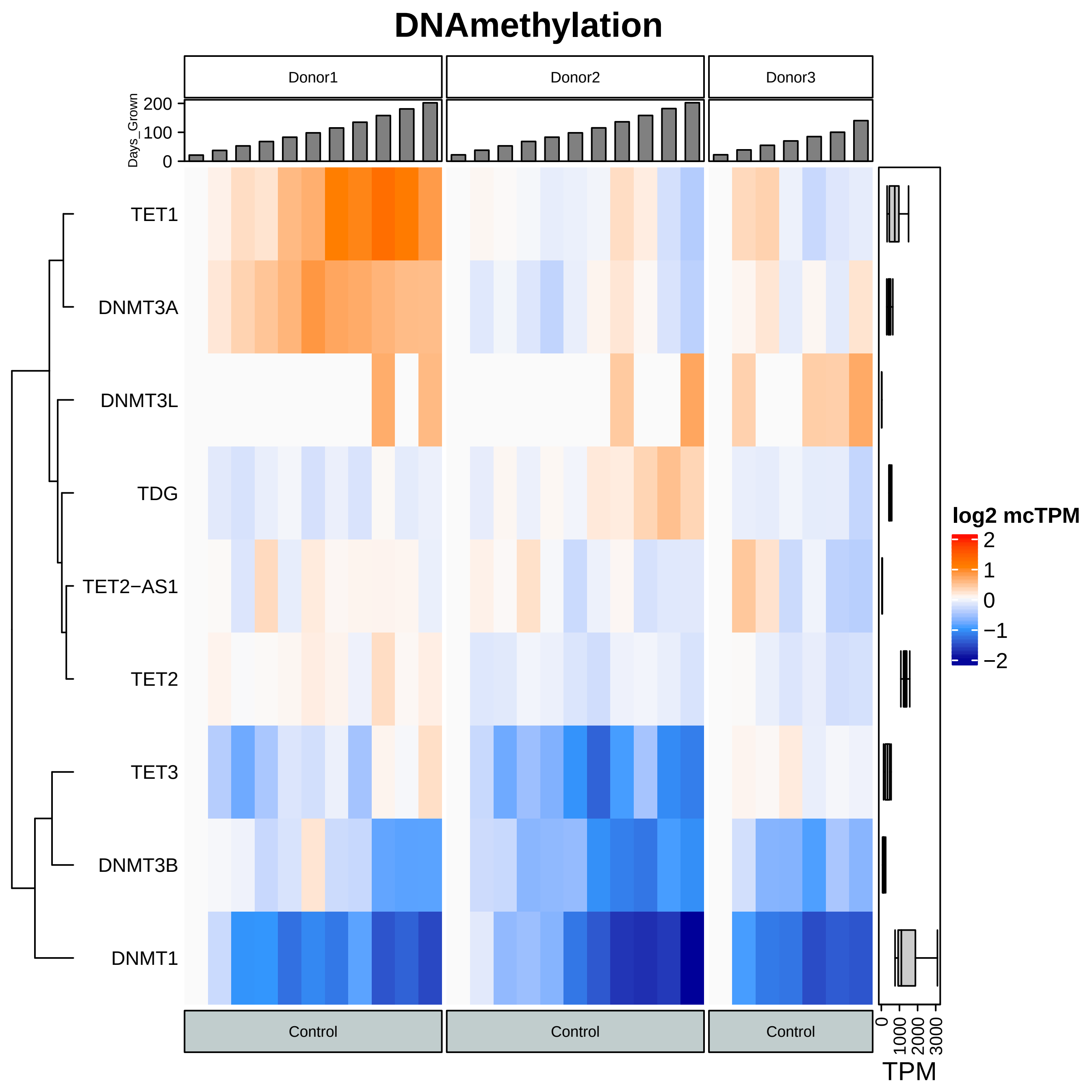
DNAmethylation
Donor1
Donor2
Donor3
Days_Grown
200
100
0
TET1
DNMT3A
DNMT3L
TDG
TET2−AS1
TET2
TET3
DNMT3B
DNMT1
log2 mcTPM
2
1
0
−1
−2
Control
Control
Control
0
1000
2000
3000
TPM

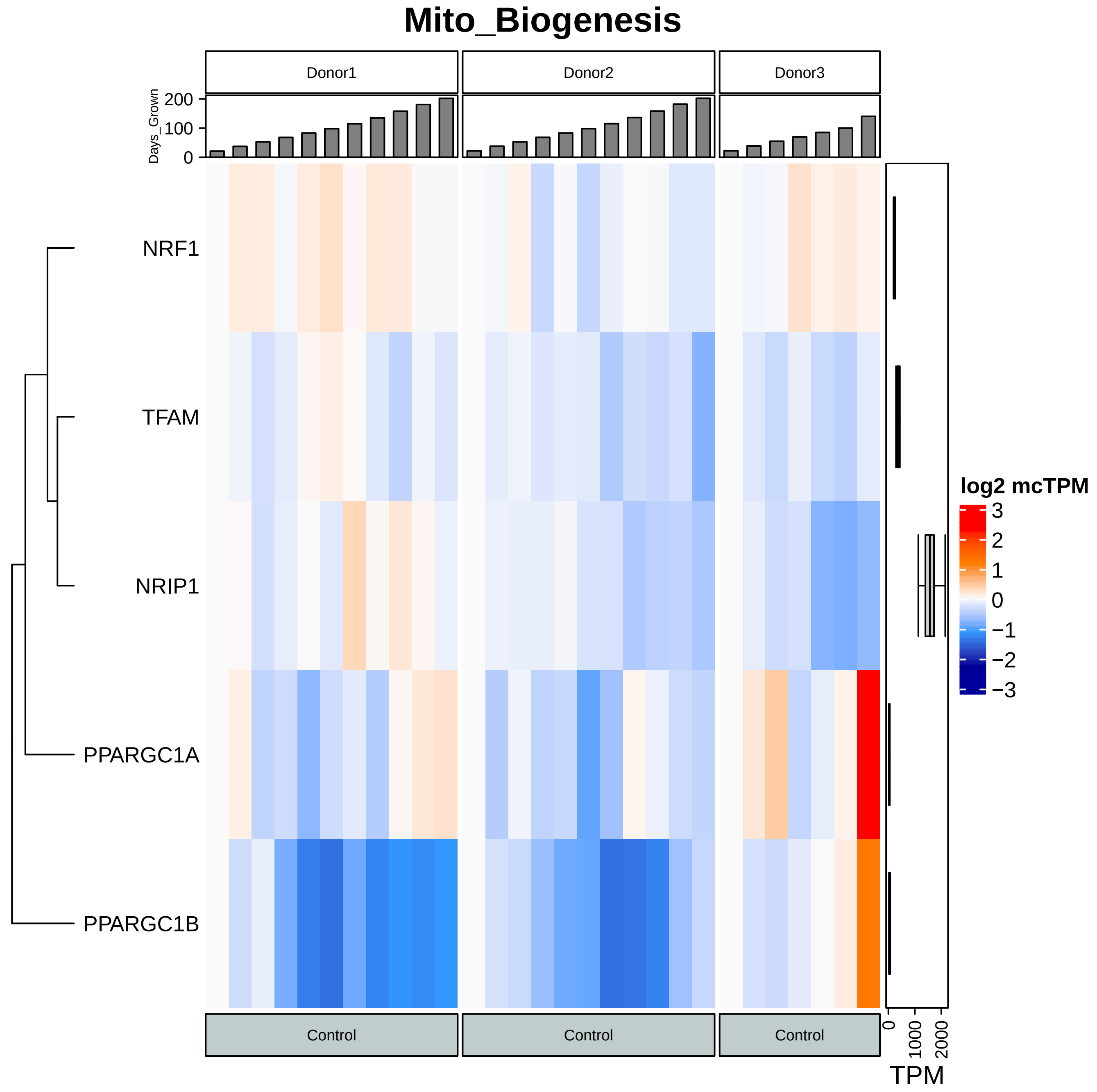

Mito_Biogenesis
Donor1
Donor2
Donor3
Days_Grown
200
100
0
NRF1
TFAM
NRIP1
PPARGC1A
PPARGC1B
log2 mcTPM
3
2
1
0
−1
−2
−3
Control
Control
Control
0
1000
2000
TPM

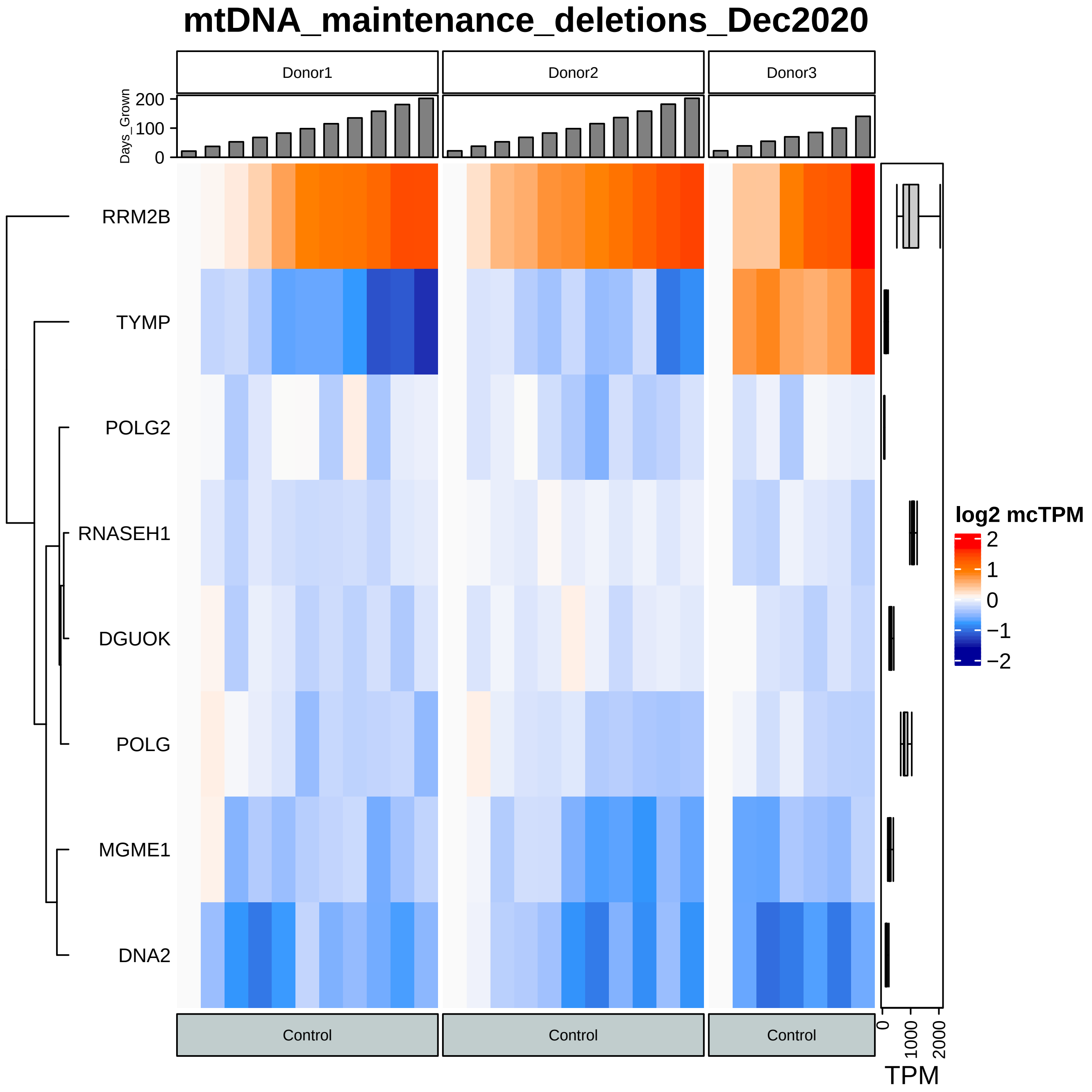
mtDNA_maintenance_deletions_Dec2020
Donor1
Donor2
Donor3
Days_Grown
200
100
0
RRM2B
TYMP
POLG2
RNASEH1
DGUOK
POLG
MGME1
DNA2
Control
Control
Control
0
1000
2000
TPM
log2 mcTPM
2
1
0
−1
−2

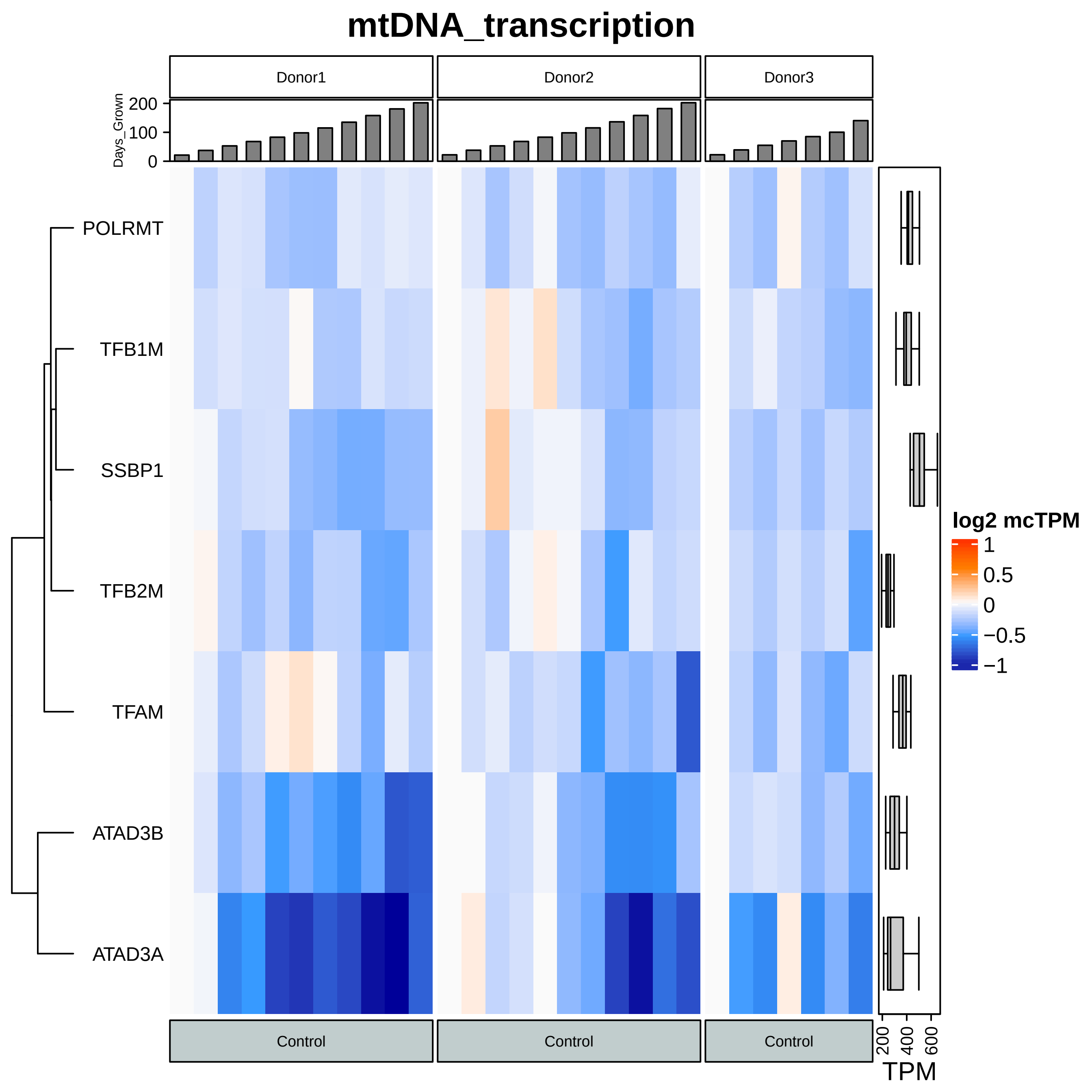

mtDNA_transcription
Donor1
Donor2
Donor3
Days_Grown
200
100
0
POLRMT
TFB1M
SSBP1
TFB2M
TFAM
ATAD3B
ATAD3A
log2 mcTPM
1
0.5
0
−0.5
−1
Control
Control
Control
200
400
600
TPM

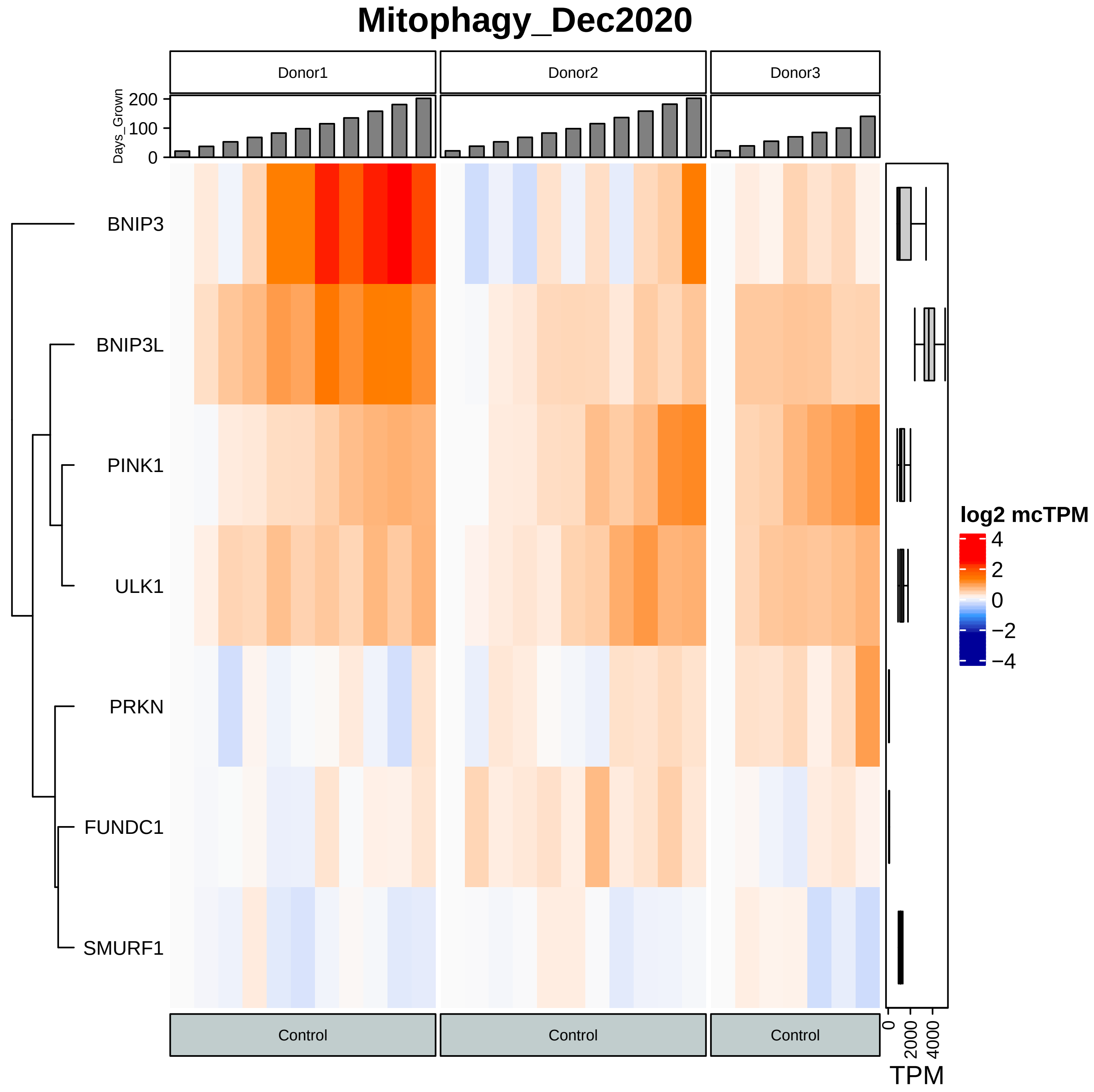

Mitophagy_Dec2020
Donor1
Donor2
Donor3
Days_Grown
200
100
0
BNIP3
BNIP3L
PINK1
ULK1
PRKN
FUNDC1
SMURF1
log2 mcTPM
4
2
0
−2
−4
Control
Control
Control
0
2000
4000
TPM

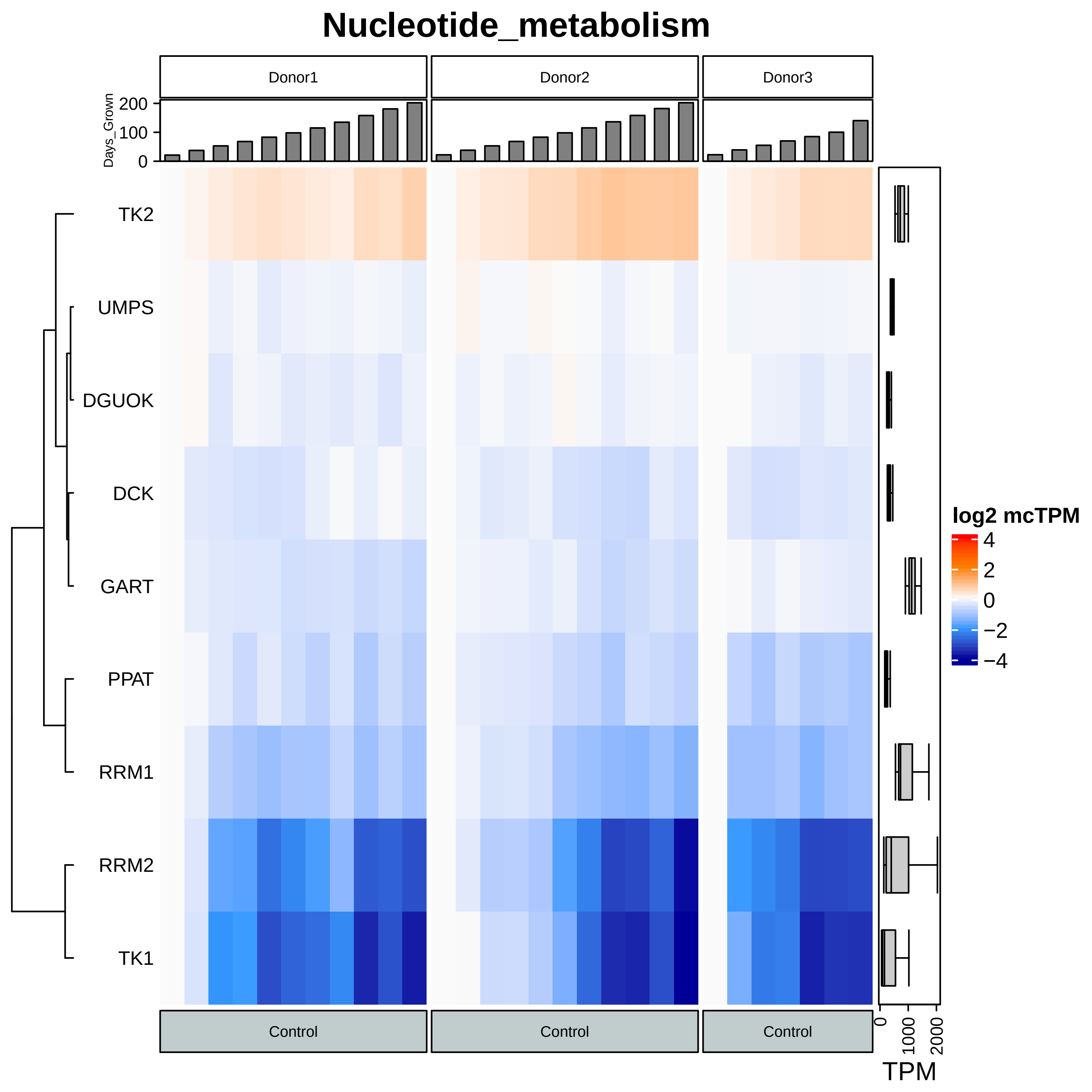
Nucleotide_metabolism
Donor1
Donor2
Donor3
Days_Grown
200
100
0
TK2
UMPS
DGUOK
DCK
GART
PPAT
RRM1
RRM2
TK1
Control
Control
Control
log2 mcTPM
4
2
0
−2
−4
0
1000
2000
TPM
